# The Erythropoietin Receptor Stimulates Rapid Cycling and Formation of Larger Red Cells During Mouse and Human Erythropoiesis

**DOI:** 10.1101/2020.11.30.404780

**Authors:** Daniel Hidalgo, Jacob Bejder, Ramona Pop, Kyle Gellatly, S. Maxwell Scalf, Anna E. Eastman, Jane-Jane Chen, Lihua Julie Zhu, Jules A.A.C. Heuberger, Shangqin Guo, Mark J. Koury, Nikolai Baastrup Nordsborg, Merav Socolovsky

## Abstract

Erythroid terminal differentiation entails cell divisions that are coupled to progressive decreases in cell size. EpoR signaling is essential for the survival of erythroid precursors, but it is unclear whether it has other functions in these cells. Here we endowed mouse precursors that lack the EpoR with survival signaling, finding that this was sufficient to support their differentiation into enucleated red cells, but that the process was abnormal. Precursors underwent fewer and slower cell cycles and yet differentiated into smaller red cells. Surprisingly, EpoR further accelerated cycling of early erythroblasts, the fastest cycling cells in the bone marrow, while simultaneously increasing their cell size. EpoR-mediated formation of larger red cells was independent of the established pathway regulating red cell size by iron through Heme-regulated eIF2α kinase (HRI). We confirmed the effect of Epo on red cell size in human volunteers, whose mean corpuscular volume (MCV) increased following Epo administration. This increase persisted after Epo declined and was not the result of increased reticulocytes. Our work reveals a unique effect of EpoR signaling on the interaction between the cell cycle and cell growth. Further, it suggests new diagnostic interpretations for increased red cell volume, as reflecting high Epo and erythropoietic stress.

## Introduction

Red cell formation (erythropoiesis) is continuous throughout life, replenishing senescent red cells. Red cell production rate is accelerated up to ten fold in the erythropoietic stress response, meeting increased demand in anemia, bleeding or hypoxic stress. Indeed, anemia is a common manifestation of nutritional deficiencies, malaria, chronic disease, cancer or hereditary hemoglobinopathies, accounting for 8.8% of all disability globally in 2010, with the highest burden afflicting women and children under five^1^. Erythropoietin (Epo) is the principal and essential regulator of erythropoietic rate in both the basal state and in the response to stress. Epo acts through its receptor, EpoR, a transmembrane type I cytokine receptor^2^. EpoR is first expressed in the earliest erythroid-committed progenitors and peaks in colony-forming-unit-erythroid (CFU-e) progenitors, a stage that undergoes amplification cell divisions^3–6^ (**Extended Data Fig 1**). EpoR becomes essential, however, only with the onset of erythroid terminal differentiation (ETD)^7^, a process that starts at the end of the CFU-e stage with the sharp induction of erythroid gene transcription^6^. During ETD, erythroblasts undergo 3 to 5 maturational cell divisions in which they become smaller, express hemoglobin and ultimately enucleate to form reticulocytes. The latter lose intracellular organelles and mature into biconcave red cells.

The essential function of EpoR is exerted early in ETD. It rescues late CFU-e, proerythroblasts and basophilic erythroblasts (here collectively termed ‘early erythroblasts’) from apoptosis^8,9^, a principal mechanism of erythropoietic rate regulation^10,11^. EpoR expression is downregulated in late erythroblasts, which no longer depend on EpoR signaling for survival^4,12,13^ (**Extended Data Fig 1**). *Epo^-/-^* or *Epor^-/-^* mice die on embryonic day 13 (E13) as a result of severe anemia^7,14,15^. Their fetal livers, the site of hematopoiesis at mid-gestation, contain early erythroid progenitors including CFU-e, but are entirely devoid of cells undergoing ETD^7,14,15^.

The absolute dependence of early erythroblasts on EpoR signaling for survival makes it challenging to identify other essential functions of EpoR in these cells. Key open questions include a role for EpoR in cell cycle regulation. Although early reports suggested that Epo does not alter the erythroblast cell cycle^16^, EpoR signaling induces some cell cycle genes in these cells^17^, and is essential for the cycling of Epo-dependent cell lines^18,19^ and of CFU-e in culture^20^. EpoR also promotes cycling in yolk-sac - derived primitive erythroblasts during early embryonic development^21^. These findings raise the possibility that EpoR may be required for the cycling of adult-type erythroblasts and that this function may contribute to the erythropoietic stress response.

A second open question is whether EpoR is required for induction of erythroid genes. EpoR and similar cytokine receptors do not instruct lineage choice and are instead required for essential permissive functions^22–25^. It is not clear, however, whether these include signals that facilitate erythroid gene transcription. EpoR signaling was shown to phosphorylate GATA1, a key erythroid transcriptional regulator, but the broad impact of this on GATA1 function is not clear^26^.

To address these gaps, we developed a genetic system that allowed us to examine essential nonsurvival functions of EpoR signaling in erythroid differentiation. We rescued mouse *Epor^-/-^* fetal liver erythroid progenitors from apoptosis by transduction with the anti-apoptotic protein bcl-x_L_, and compared their ensuing differentiation with that of *Epor^-/-^* progenitors that were rescued by reintroduction of the EpoR. We found that the bcl-x_L_ survival signal, in the absence of any EpoR signaling, was sufficient to allow expression of the erythroid transcriptional program and formation of enucleated red cells. However, key ETD features were abnormal. First, erythroblasts underwent slower and fewer cell cycles, generating far fewer red cells, showing that EpoR is essential for the rapid cycling of early erythroblasts. We confirmed this using a mouse transgenic for a fluorescent reporter of cell cycle length, finding that Epo stimulation shortened cell cycle length *in vivo* in early erythroblasts, cells that are already amongst the fastest cycling cells in the bone marrow^27,28^. Second, we found that, unexpectedly, despite its stimulation of rapid erythroblast cycling, EpoR signaling increases cell size in both erythroblasts and red cells. This contrasts with the well-established reverse relationship between the number of cell divisions in ETD and red cell size^29–32^. Using mice doubly deleted for both EpoR and HRI, we found that EpoR regulation of red cell size is also independent of the well described iron and heme-regulated pathway^33–35^. We confirmed these findings in healthy human volunteers that were administered Epo, finding an increased MCV that persisted long after Epo and reticulocyte levels returned to baseline. Our work reveals novel functions for the EpoR, including its likely modulation of red cell size during hypoxia, anemia and other high-Epo erythroid stress syndromes. These mechanistic insights will aid diagnostic interpretation of red cell size abnormalities commonly found in the clinic.

## Results

### Non-survival EpoR signals are essential for normal erythroid differentiation

Erythroid differentiation in the *Epor^-/-^* fetal liver is arrested at the CFU-e stage, which leads to the death of the *Epor^-/-^* embryo on embryonic day 13 (E13)^7,14,15,21^. *Epor^-/-^* fetal liver CFU-e can be rescued *in vitro* by transduction with EpoR or a similar cytokine receptor^7,23^. Here we asked whether transducing *Epor^-/-^* CFU-e with bcl-x_L_, an anti-apoptotic transcriptional target of EpoR signaling^36–39^, would be sufficient to support erythroid differentiation. As control, we transduced *Epor^-/-^* cells from the same fetal livers with the EpoR. The use of bicistronic retroviral expression vectors allowed us to track transduced cells (**Fig 1a**).

**Figure 1.**
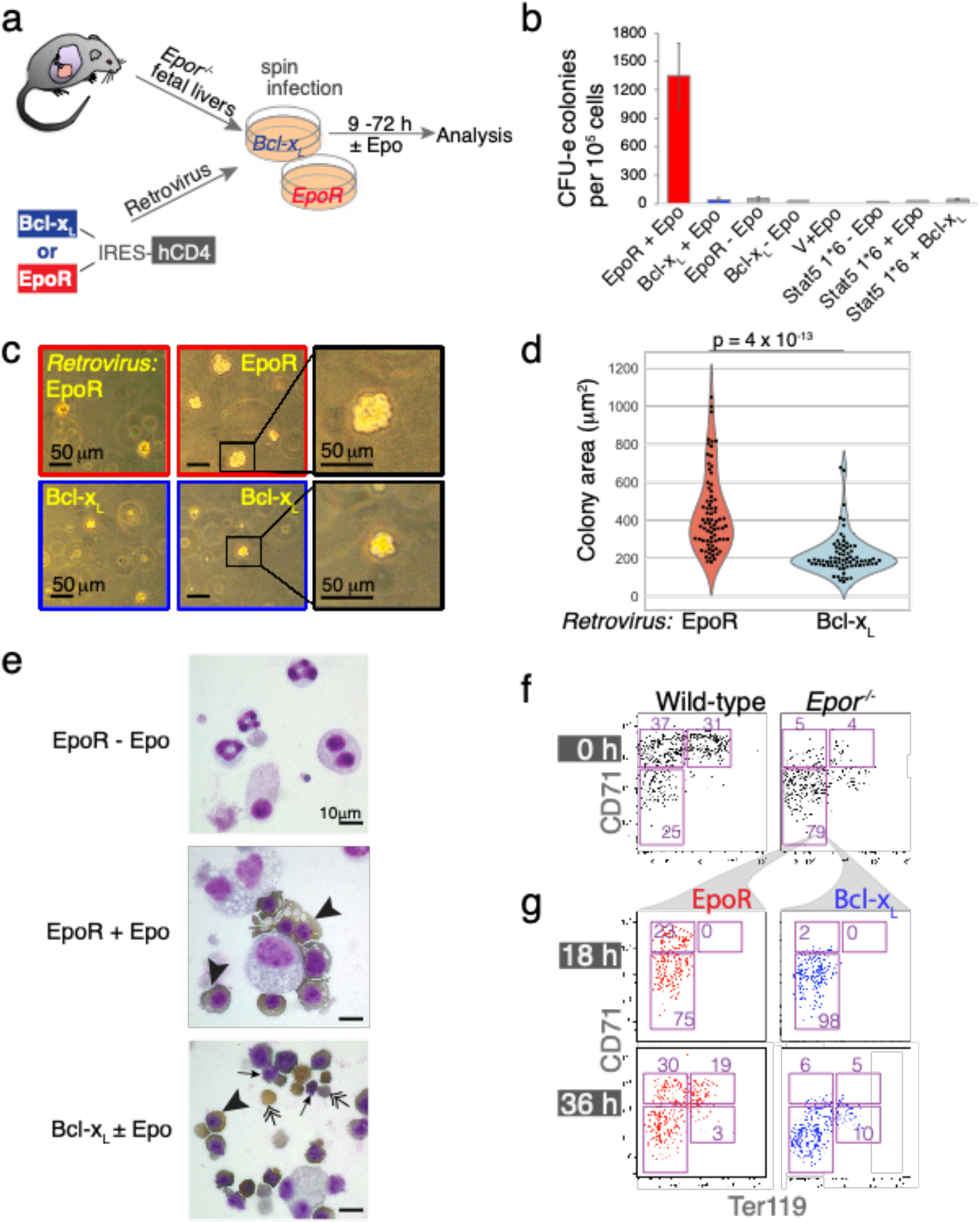
Abnormal ETD in the absence of EpoR signaling. **a** Experimental design. E12.5 *Epor^-/-^* fetal livers were transduced with bicistronic retroviral vectors encoding either Bcl-x_L_ or EpoR, linked by an internal ribosomal entry site (IRES) to human CD4 (hCD4) or GFP reporters. Transduced cells differentiated *in vitro* into red cells over the ensuing 72 hours. **b** *Epor^-/-^* CFU-e colonies, scored 48 hours following transduction with either EpoR or Bcl-x_L_. Epo was added to the medium where indicated. *Epor^-/-^* fetal liver cells were also transduced with retroviral vectors encoding the following: ‘empty’ vector (‘V’), constitutively active Stat5 (Stat5 1*6), or doubly transduced with both Bcl-x_L_ and Stat5 1*6. Data pooled from 3 independent experiments. Only CFU-e colonies of a size comparable to those of wild-type colonies were scored. **c** Representative colonies from an experiment as in ‘**b**’ **d** Colony area occupied by each of 75 colonies for each genotype (EpoR-*Epor^-/-^* or Bcl-x_L_-*Epor^-/-^*). Data pooled from 3 independent experiments as in ‘**b**’. **e** Cytospin preparations of transduced *Epor^-/-^* fetal liver cells cultured in liquid medium for 36 hours, in the presence or absence of Epo as indicated. Cells were stained for hemoglobin with diaminobenzidine (brown stain, arrowheads) and counter-stained with Giemsa. Representative of 4 independent experiments. Double-headed arrows point at enucleated red cells; arrows point at pyrenocytes (extruded nuclei). The micrograph in the bottom panel is representative of cultures both in the presence or absence of Epo. **f, g** Flow cytometric CD71/Ter119 profiles of freshly harvested *Epor^-/-^* and wild-type littermate fetal livers (**f**), and of *Epor^-/-^* fetal liver cells 18 and 36 hours post transduction and culture in Epo-containing medium (**g**).

We plated the transduced cells in methylcellulose semi-solid medium and examined their ability to give rise to CFU-e colonies (**Fig 1b**). As expected, *Epor^-/-^* cells transduced with empty vector failed to give rise to any colonies, whereas EpoR-transduced *Epor^-/-^* cells (EpoR-*Epor^-/-^*) generated CFU-e colonies in an Epo-dependent manner. Bcl-x_L_-transduced *Epor^-/-^* cells (Bcl-x_L_-*Epor^-/-^*) failed to give rise to CFU-e colonies of the usual size and appearance (**Fig 1b**). Close inspection showed, however, that they generated a similar number of much smaller cell clusters, which were not scored as CFU-e colonies in Figure 1b. These smaller clusters contained fewer than16 hemoglobinized cells, compared with 16 to 32 cells in EpoR-supported CFU-e colonies (**Fig 1c**). The area occupied by Bcl-x_L_-*Epor^-/-^* colonies was approximately half that of EpoR-*Epor^-/-^* CFU-e colonies (439 ± 208 μm^2^ *versus* 217 ± 106 μm^2^, mean ± SD for EpoR-*Epor^-/-^* v. Bcl-x_L_-*Epor^-/-^*, respectively; n = 75 colonies for each genotype; p = 3.6 × 10^-13^, **Fig 1d**), suggesting approximately 3-fold difference in colony volume due to fewer and smaller cells (see below). Co-transduction of *Epor^-/-^* cells with both Bcl-x_L_ and a constitutively active form of Stat5, a transcription factor activated by EpoR signaling, was also not sufficient to support the formation of normally-sized *Epor^-/-^* CFU-e colonies (**Fig 1b**).

Liquid cultures of both Bcl-x_L_-*Epor^-/-^* with and without Epo and EpoR-*Epor^-/-^* erythroblasts with Epo contained hemoglobinized cells by 36 hours post-transduction, while EpoR-*Epor^-/-^* erythroblasts without Epo did not (**Fig 1e**). However, differentiation of Bcl-x_L_-*Epor^-/-^* erythroblasts appeared to be accelerated, with cultures containing smaller and morphologically more mature erythroblasts including many fully differentiated and enucleated cells; there were few if any enucleated cells in EpoR-*Epor^-/-^* erythroblasts cultured with Epo at this time (**Fig 1e**).

Differentiation abnormalities of Bcl-x_L_-*Epor^-/-^* erythroblasts were also evident from flow cytometric analysis. The earliest progenitors express only low or medium levels of CD71, encoded by the transferrin receptor (*Tfrc*). In wild-type progenitors, the transition from the CFU-e stage to ETD is marked by sharp upregulation of CD71, followed by the upregulation of Ter119, a marker specific for ETD erythroblasts and red cells^6,27,40^. We previously found that freshly explanted *Epor^-/-^* fetal livers lack CD71^high^ cells, and do not express Ter119, reflecting their developmental arrest at the onset of ETD (the small number of Ter119^+^ cells in the *Epor^-/-^* fetal liver are transient yolk-sac-derived primitive erythroblasts^40^, **Fig 1f**). Here we found that transduction of *Epor^-/-^* fetal liver cells with EpoR allowed them to resume the expected sequence of cell surface marker expression, with rapid upregulation of CD71 by 18 hours post infection, followed by the upregulation of Ter119 by 36 hours (**Fig 1g**). By contrast, Bcl-x_L_-*Epor^-/-^* cells failed to upregulate CD71 at any point of the culture although they did upregulate Ter119 (**Fig 1g**).

Thus, our initial analysis showed that, when rescued from apoptosis by Bcl-x_L_, *Epor^-/-^* progenitors can differentiate into hemoglobinized, enucleated red cells in the absence of additional EpoR signals. However, the ETD they undergo has a number of abnormalities. Bcl-x_L_-*Epor^-/-^* erythroblasts failed to upregulate CD71, and their differentiation appeared to be accelerated, generating fewer and smaller red cells.

### Erythroblasts undergo fewer and slower cell cycles in the absence of EpoR signaling

CFU-e express the receptor tyrosine kinase Kit and the Interleukin-3 (IL3) receptor, both of which increase CFU-e number^23,41,42^. The addition of stem cell factor (SCF, the Kit ligand) and IL3 to the media increased the overall yield of transduced *Epor^-/-^* fetal liver cells, but the difference in cell number between Bcl-x_L_-*Epor^-/-^* and EpoR-*Epor^-/-^* erythroblasts remained (**Extended Data Fig 2a**). We modified our transduction protocol to make use of this improvement in yield, culturing freshly transduced *Epor^-/-^* progenitors for 15 hours in SCF and IL3 before transitioning the cells to an Epo-containing medium for the remainder of differentiation. Since SCF and IL3 also promote the growth of myeloid cells, all subsequent analysis was performed on cells that were both negative for non-erythroid lineage markers and positive reporters of transduction (hCD4 and/or GFP, **Extended Data Fig 2b, Fig 2a**).

**Figure 2.**
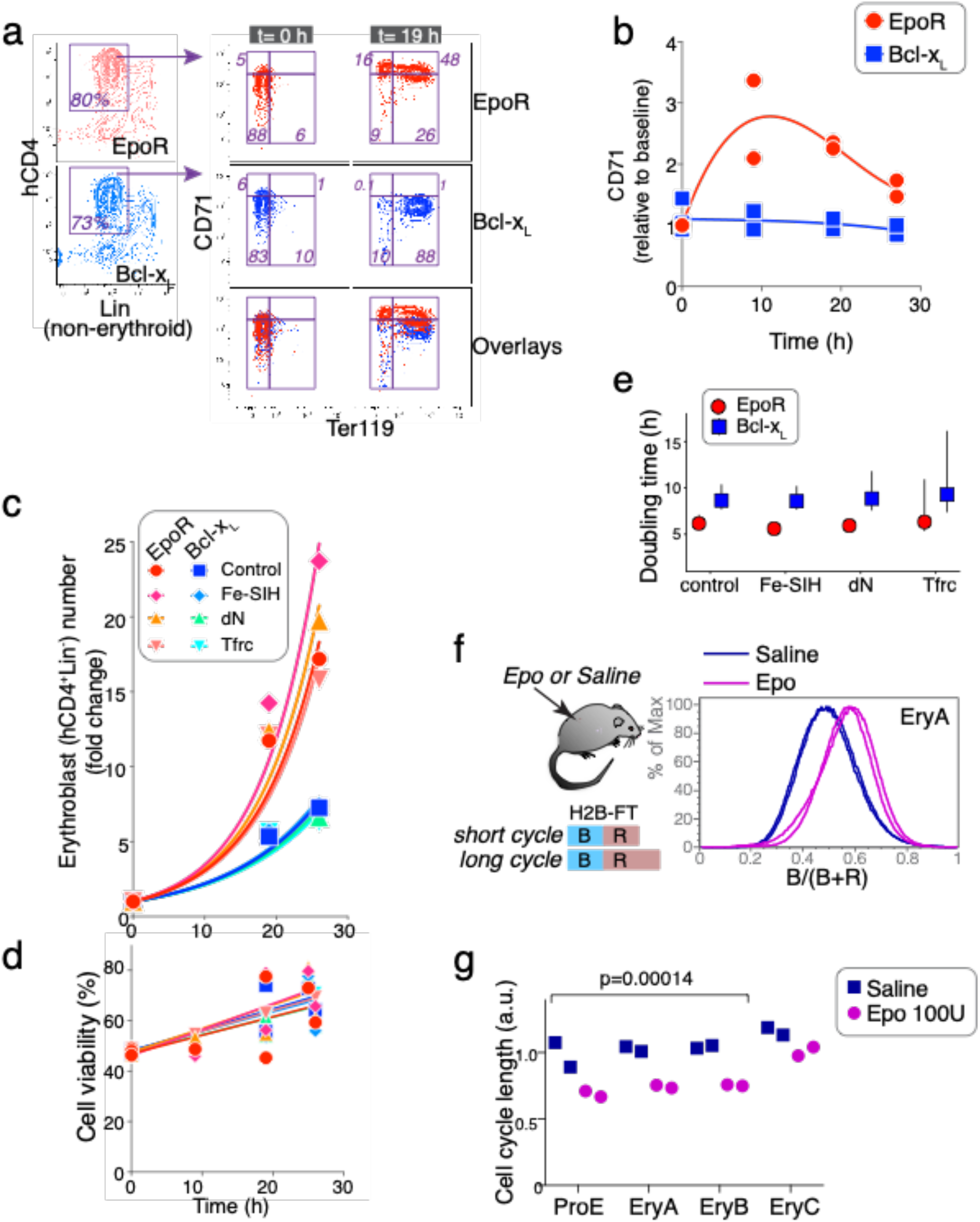
EpoR stimulates shorter cell cycle in early erythroblasts *in vitro* and *in vivo*. **a** The EpoR is required for CD71 upregulation. *Epor^-/-^* fetal livers were transduced with either EpoR or Bcl-x_L_ retroviral vectors carrying the hCD4 reporter (see Extended Data Fig 2b for experimental design). Transduced erythroid cells (hCD4^+^Lin^-^) were examined for expression of CD71 and Ter119. **b** Time course of CD71 expression following EpoR or Bcl-x_L_ retroviral transduction as in ‘a’. MFI; median fluorescence intensity, expressed relative to t=0. Data from two independent experiments. **c** Growth of *Epor^-/-^* fetal liver transduced with either Bcl-x_L_ or EpoR. Viable hCD4^+^Lin^-^ cells were counted at the indicated times. Fe-SIH (10 μM) or deoxynucleosides (dN, 0.7 μM) were added to the medium as indicated. ‘Tfrc’ cells were doubly transduced with both *Tfrc*, and either *Epor* or Bcl-x_L_. Data are pooled for each set of conditions from n=4 independent experiments (‘control’ growth curves in which cells were transduced with either EpoR or Bcl-x_L_ but without additional manipulation of iron or dN were included in every experiment). Data are expressed relative to cell number at t=0, and were fit with exponential curves (R^2^ values ranging between 0.8 and 0.94, least squares fit). **d** Cell viability, expressed as the fraction (%) of trypan blue negative cells, for the same set of experiments shown in ‘c’. **e** Cell doubling times ± 95% confidence intervals, calculated from the fitting of exponential growth curves to the data in ‘e’. **f** Cell cycle shortening in early erythroblasts *in vivo*. Mice transgenic for a fusion of histone H2B and a fluorescence-timer protein (H2B-FT) were injected with either saline or Epo (100 U) at 0 and 24 h. Bone-marrow was analyzed at 48 h. H2B-FT fluoresces blue (‘B’) for 1-2 h immediately following synthesis, and matures into a long-lived red fluorescent protein (‘R’). The blue to red fluorescence ratio in the cell (expressed as *B/(B+R))* is a function of cell cycle length^28^. Shown are histograms of *B/(B+R)* in live EryA erythroblasts (Ter119^high^CD71^high^FSC^high^). Histogram overlays are for 2 mice injected with saline and 2 mice injected with Epo. **g** Relative cell cycle lengths for the 4 mice analyzed in ‘f’, for each of the indicated erythroblast maturation stages: ProE (Ter119^med^CD71), EryA (Ter119^high^CD71^high^FSC^high^), EryB (Ter119^high^CD71^high^FSC^low^), EryC (Ter119^high^CD71^low^FSC^low^). p-value is for a paired *t* test, pairing the average values for Epo-injected and Saline injected mice for each of the early erythroblast stages (ProE and EryA/B). Cell cycle length was calculated as R/B.

Pre-incubation with SCF and IL3 did not ameliorate the abnormalities of Bcl-x_L_-*Epor^-/-^* erythroblast differentiation. In particular, these cells failed to upregulate CD71 throughout differentiation (**Fig 2a, b**). We examined the possibility that these abnormalities were the result of over-expression of Bcl-x_L_, rather than the absence of EpoR signaling, by including an additional control in which *Epor^-/-^* progenitors were co-transduced with both EpoR and Bcl-x_L_, each linked to a distinct reporter (**Extended Data Fig 2c**). These doubly-transduced progenitors were indistinguishable from cells transduced with only the EpoR, indicating that the lower cell number of Bcl-x_L_-*Epor^-/-^* erythroblasts and their failure to upregulate CD71 were not the result of Bcl-x_L_ over-expression, but rather, of absent EpoR signaling (**Extended Data Fig 2d-g**).

CD71 is the transferrin receptor, critical for importing iron-bound transferrin into erythroid cells for heme synthesis. Iron deficiency leads to microcytic anemia. We therefore examined whether iron deficiency might account for the abnormal differentiation of Bcl-x_L_-*Epor^-/-^* erythroblasts, by co-transducing *Epor^-/-^* progenitors with *Tfrc*, in addition to either Bcl-x_L_ or EpoR (**Fig 2c-e**). As an alternative approach, we added iron-loaded ferric-salicylaldehyde isonicotinoyl hydrazone (Fe-SIH) to the culture medium of both Bcl-x_L_-*Epor^-/-^* and EpoR-*Epor^-/-^* erythroblasts. SIH is a cell-membrane-permeable synthetic iron chelate, which, when pre-loaded with iron, will deliver iron intracellularly for heme synthesis, bypassing defects in Tfrc iron transport in erythroid cells^43^. Neither of these approaches altered the proliferative defect of Bcl-x_L_-*Epor^-/-^* erythroblasts (**Fig 2c, e**). The viability of all erythroblasts was high with no significant difference between Bcl-x_L_-*Epor^-/-^* and EpoR-*Epor^-/-^* erythroblasts (**Fig 2d**), suggesting that the proliferative deficit of Bcl-x_L_-*Epor^-/-^* erythroblasts is the result of fewer cell divisions. In the first 26 hours of culture, the difference in growth rate for the Bcl-x_L_-*Epor^-/-^* and EpoR-*Epor^-/-^* cultures indicated a substantial difference in doubling time, of 6.1 h for EpoR-*Epor^-/-^* erythroblasts, compared with 8.6 h for Bcl-x_L_-*Epor^-/-^* erythroblasts (95% confidence intervals: 5.6 to 7.1 h and 7.7 to 10.4 h, respectively; **Fig 2c, e**). A cycle length of 6 hours for EpoR-*Epor^-/-^* is in good agreement with our recent direct measurement of a 6 hour cell cycle duration in early erythroblasts *in vivo*^27^, and with an independent measurement using a fluorescent timer protein that found early erythroblasts to have the shortest cell cycle amongst bone marrow hematopoietic progenitors^28^.

In addition to its role in heme synthesis, iron has multiple cellular functions, including its requirement in ribonucleotide reductase (RNR) catalysis of deoxyribonucleotide synthesis. Depletion of the intracellular iron pool can therefore lead to inhibition of both DNA synthesis and cell growth^44^. We supplemented the culture medium with deoxyribonucleosides (dN), which bypass RNR via the deoxyribonucleoside kinase salvage pathway^45^. This too had little effect on the proliferative defect of Bcl-x_L_-*Epor^-/-^* erythroblasts (**Fig 2c, e**). Taken together, in the absence of EpoR signaling, erythroblasts fail to upregulate CD71 and also undergo fewer and longer cell divisions. The rescue of intracellular iron or the exogenous supply of deoxyribonucleosides does not rescue these deficits.

### Epo administration shortens cell cycle duration in early erythroblasts *in vivo*

To test whether EpoR stimulation altered cell cycle length *in vivo*, we used a recently-engineered mouse expressing a live-cell fluorescent reporter of cell cycle speed^28^. Specifically, the mouse expresses a transgene encoding histone H2B fused to a fluorescent timer protein (H2B-FT, **Fig 2f**). H2B-FT fluoresces blue when first synthesized but matures over 1-2 hours into a red fluorescent protein. Due to its short half-life, the level of blue fluorescent H2B-FT in the cell is minimally affected by cell cycle length. In contrast, the red-fluorescent H2B-FT accumulates to higher levels in cells of long cycles due to its extensive stability. The ratio of blue to red fluorescence was found to be a sensitive indicator of cell cycle length *in vivo* in diverse cell types including bone marrow progenitors^28^.

We injected H2B-FT transgenic mice with either Epo (100 U) or saline once daily for 2 days, and analyzed bone marrow at 48 h. We found a clear shift in the blue to red fluorescence ratio, indicating significant cell cycle shortening, in all bone marrow early erythroblast subsets in Epo injected mice (**Fig 2f, g** and **Extended Data Fig 3**). These data are consistent with our results in the fetal liver (**Fig 2c, e**), together indicating that Epo stimulation increases cell cycle speed in early erythroblasts.

### Slower S phase in the absence of EpoR signaling, partially rescued by iron supplementation

We recently found that the onset of ETD in the fetal liver is closely linked to cell cycle remodeling events that radically shorten the cell cycle, from an average of 15 hours in BFU-e and CFU-e, to 6 hours in early erythroblasts^6,27,40^. The short 6 hour cycle includes a shortened S phase, lasting only 4 hours^27^. Our findings here show that EpoR signaling is required for the short cycle of early erythroblasts (**Fig 2e, g**) and led us to ask whether EpoR was required for the shortening only of G1, or whether it also played a role in S phase shortening. The shortening of G1 by cytokine receptor signaling is well documented^46–50^. However, to our knowledge, there are no reports of cytokine or growth factor signaling altering the speed and duration of S phase.

To examine this, we pulsed cultures of EpoR-*Epor^-/-^* and Bcl-x_L_-*Epor^-/-^* erythroblasts with bromodeoxyuridine (BrdU), a nucleoside analog that is incorporated into DNA during S phase, and analyzed the cells 30 min following the pulse. The fraction of cells that are labeled with an anti-BrdU antibody indicates the proportion of cells in S phase at the time of the pulse. Further, the amount of BrdU incorporated into S phase cells during the 30 minutes pulse, as measured by the BrdU mean fluorescence intensity (MFI) of S phase cells, indicates the intra-S phase rate of DNA synthesis, which is inversely related to S phase duration^27^. We found that, in the first 10 hours of ETD, BrdU MFI in S phase cells was 50% higher in EpoR-*Epor^-/-^* compared with Bcl-x_L_-*Epor^-/-^* erythroblasts, suggesting that EpoR signaling increases intra-S phase DNA synthesis rate and shortens S phase duration by 50% (**Fig 3a, b**).

**Figure 3.**
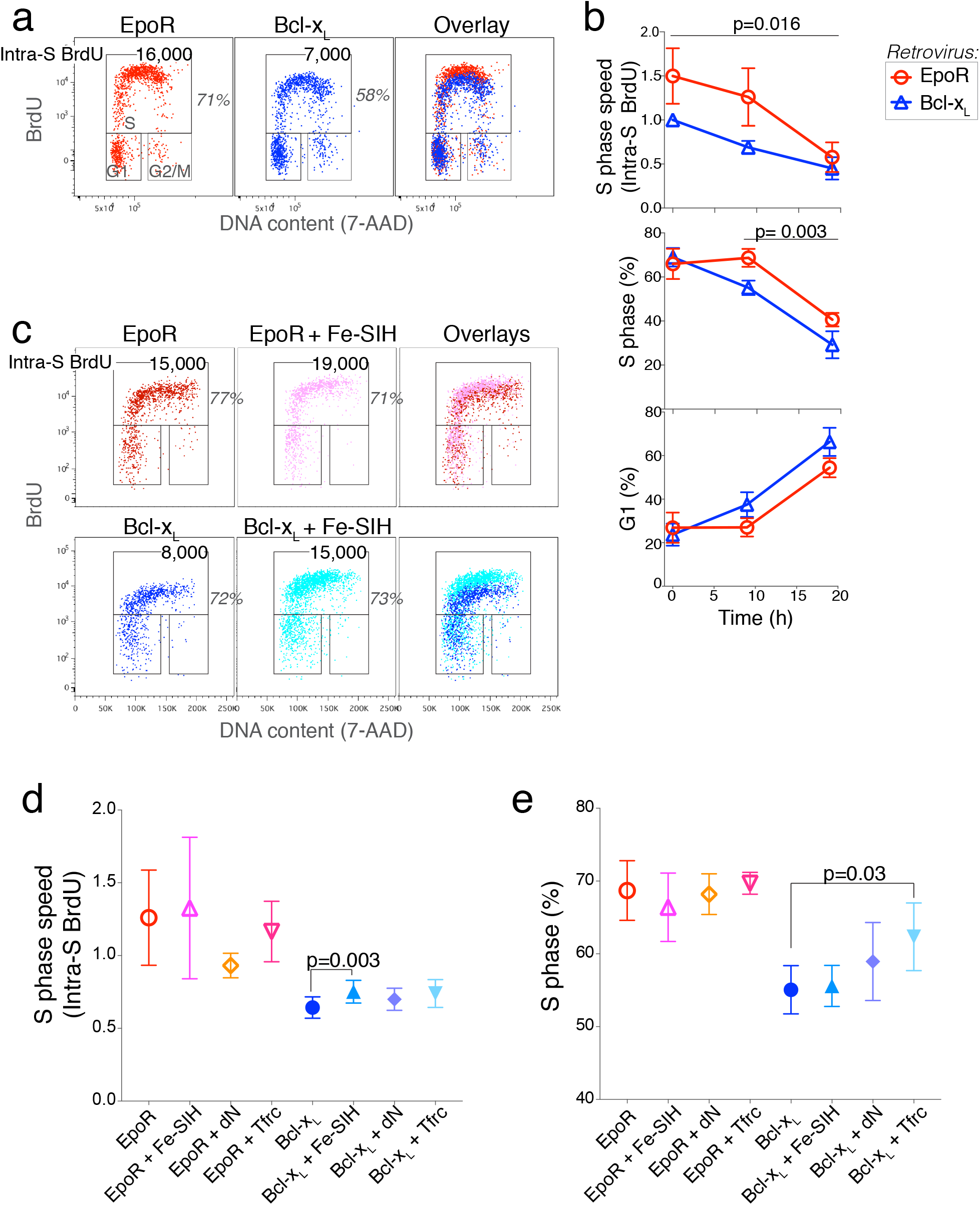
EpoR regulates the speed of S phase. **a** Cell cycle analysis of *Epor^-/-^* fetal liver cells transduced with either EpoR or Bcl-x_L_ and cultured as in Fig 2b. Cells were pulsed with BrdU for 30 min at t=9 h and were immediately harvested for analysis. The fraction (%) of erythroblasts (hCD4^+^Lin^-^) in S phase is indicated, as is S phase speed, measured as the intra-S phase rate of BrdU incorporation (BrdU MFI within the S phase gate). **b** Summary of cell cycle status and S phase speed, as measured by intra-S phase BrdU incorporation in EpoR or Bcl-x_L_–transduced *Epor^-/-^* fetal liver cells. Data is pooled from 6 independent experiments similar to ‘a’. In all cases, cells were pulsed with BrdU for 30 minutes prior to harvesting for analysis. Data are mean ± sem. Intra-S phase BrdU (MFI) is expressed as the ratio to BrdU MFI of Bcl-x_L_-transduced fetal liver cells at t=0 in each experiment. Significance p values are paired *t* test, pairing EpoR and Bcl-x_L_–transduced cells for each time point across all experiments (upper panel), and for t=9 and t=19 h in all experiments (middle and lower panels). **c** Effect of the cell-permeable iron carrier, Fe-SIH (10 mM) on S phase speed. Experiment and cell cycle analysis as in ‘b’. Cells were harvested at t=9 h. **d, e** Summary of S phase speed (d) and cell cycle status (e) in EpoR and Bcl-x_L_-transduced *Epor^-/-^* fetal liver cells at t=9 h, experimental design as in Fig 2b, and ‘a’ to ‘c’ above. S phase speed is expressed relative to the speed at t=0 in each experiment. Shown are the effects of adding Fe-SIH or dN to the medium, or of doubly transducing cells with both Bcl-x_L_ and Tfrc. Data are mean ± sem for n = 4 independent experiments each for Fe-SIH and dN, and n = 3 for Tfrc. All experiments also had *Epor^-/-^* fetal liver cells transduced with EpoR and with Bcl-x_L_ without additional additives or transductions.

If the slowing of S phase alone could account for the increased cell cycle length of Bcl-x_L_-*Epor^-/-^* erythroblasts, S phase would constitute a larger fraction of the total cell cycle duration. However, the fraction of Bcl-x_L_-*Epor^-/-^* erythroblasts in S phase was actually somewhat lower, with a corresponding increase in the fraction of cells in G1 (**Fig 3b**), and little change in the fraction of cells in G2 or M (not shown). These observations suggest that, in the absence of EpoR signaling, both S and G1 phases lengthen.

We examined the potential roles of iron and nucleotides in the slower S phase of Bcl-x_L_-*Epor^-/-^* erythroblasts. Supplementing the culture medium with Fe-SIH increased S phase speed modestly in both Bcl-x_L_-*Epor^-/-^* and EpoR-*Epor^-/-^* erythroblasts (**Fig 3c**). Across 4 experiments, the increase in S phase speed in Bcl-x_L_-*Epor^-/-^* erythroblasts was statistically significant but accounted for only 10% of the difference between Bcl-x_L_-*Epor^-/-^*and EpoR-*Epor^-/-^* erythroblasts (relative S phase speed of **0.68 ± 0.07** for Bcl-x_L_-*Epor^-/-^*, **1.2 ± 0.3** for EpoR-*Epor^-/-^*, and **0.75 ± 0.07** for Bcl-x_L_-*Epor^-/-^* supplemented with Fe-SIH, Fig 3d). There was no rescue of S phase speed in Bcl-x_L_-*Epor^-/-^* erythroblasts by either the addition of deoxyribonucleosides nor double transduction with both Bcl-x_L_ and Tfrc (**Fig 3d**), although there was a small increase in the number of cells in S phase in the latter (**Fig 3e**). Taken together, these results indicate that EpoR is essential for accelerating both G1 and S phases of the shortened cell cycles in early ETD, largely, but not exclusively, via mechanisms independent of iron and the nucleotide pool.

### Premature induction of the CDK inhibitor p27^KIP1^ in the absence of EpoR signaling

We compared gene expression in EpoR-*Epor^-/-^* and Bcl-x_L_-*Epor^-/-^* erythroblasts during the 48-hour differentiation time course, using RT-qPCR (**Extended Data Fig 4**). ETD markers *Slc4a1* (*Band3*) and *Hbb1* were induced similarly in both cell types. There were no significant differences in the expression of transcription factors, with the possible exception of Tal1, whose levels were 30% lower in Bcl-x_L_-*Epor^-/-^* (p<0.005, **Extended Data Fig 4**).

Among cell cycle regulators, we found few differences. The clear exception was the CDK inhibitor p27^KIP1^ (encoded by *Cdkn1b*), previously shown to be induced towards the end of ETD, as part of the mechanism that slows the cycle and leads to mitotic exit prior to enucleation^51–54^. Here we found that p27^KIP1^ was induced prematurely in Bcl-x_L_-*Epor^-/-^* compared with EpoR-*Epor^-/-^* cells, with levels 2.3 and 2.6-fold higher at 19 and 24 hours of culture (p<0.005, **Extended Data Fig 4**). The premature induction of p27^KIP1^ might explain the slower and fewer cycles of Bcl-x_L_-*Epor^-/-^* cells (**Fig 2, 3**), and may also contribute to the accelerated maturation of Bcl-x_L_-*Epor^-/-^* erythroblasts (**Fig 1e** and further analysis below). We also found a small but significant expression difference in a second member of the CIP/KIP family, p57^KIP2^ (encoded by *Cdkn1c*). p57^KIP2^ is expressed in CFU-e cells, and undergoes rapid downregulation at the onset of ETD, precipitating the concurrent shortening of the cycle^27^. Here we found that, in the absence of EpoR, p57^KIP2^ downregulation at the onset of ETD was largely intact. These findings indicate that p57^KIP2^ downregulation and EpoR signaling each drive cell cycle shortening in early ETD by largely independent mechanisms.

### Imaging flow cytometry shows *Epor^-/-^* erythroblasts and reticulocytes are smaller

Erythroid maturational cell divisions are coupled to loss in cell size. This is supported by the findings that drugs, nutritional deficiencies or genetic perturbations that reduce the number of cell divisions lead to the formation of larger red cells (macrocytosis^29–32,55^). Therefore, we expected that the fewer cell divisions of Bcl-x_L_-*Epor^-/-^* erythroblasts would result in larger size for these cells. Instead, they appeared to be smaller (**Fig 1e**). To address this question quantitatively, we measured cell and nuclear size in several thousand EpoR-*Epor^-/-^* and Bcl-x_L_-*Epor^-/-^* erythroblasts by imaging flow cytometry, at 20 and 46 hours of differentiation (**Fig 4a, b**). We calibrated the measured cell areas by comparing them with those of beads of known diameter (**Extended Data Fig 5a**). We found that both cell and nuclear size were significantly smaller in Bcl-x_L_-*Epor^-/-^* erythroblasts, at both time points (at t= 46 h, cell diameter was 7.5 ± 0.6 μm and 6.7 ± 0.7 μm for EpoR-*Epor^-/-^* and Bcl-x_L_-*Epor^-/-^* erythroblasts, respectively, mean ± sem, p = 0.001). Although Bcl-x_L_-*Epor^-/-^* erythroblasts express significantly lower CD71 (**Fig 2a, b**), the addition of Fe-SIH to the culture did not alter their smaller cell or nuclear size (**Fig 4b**).

**Figure 4.**
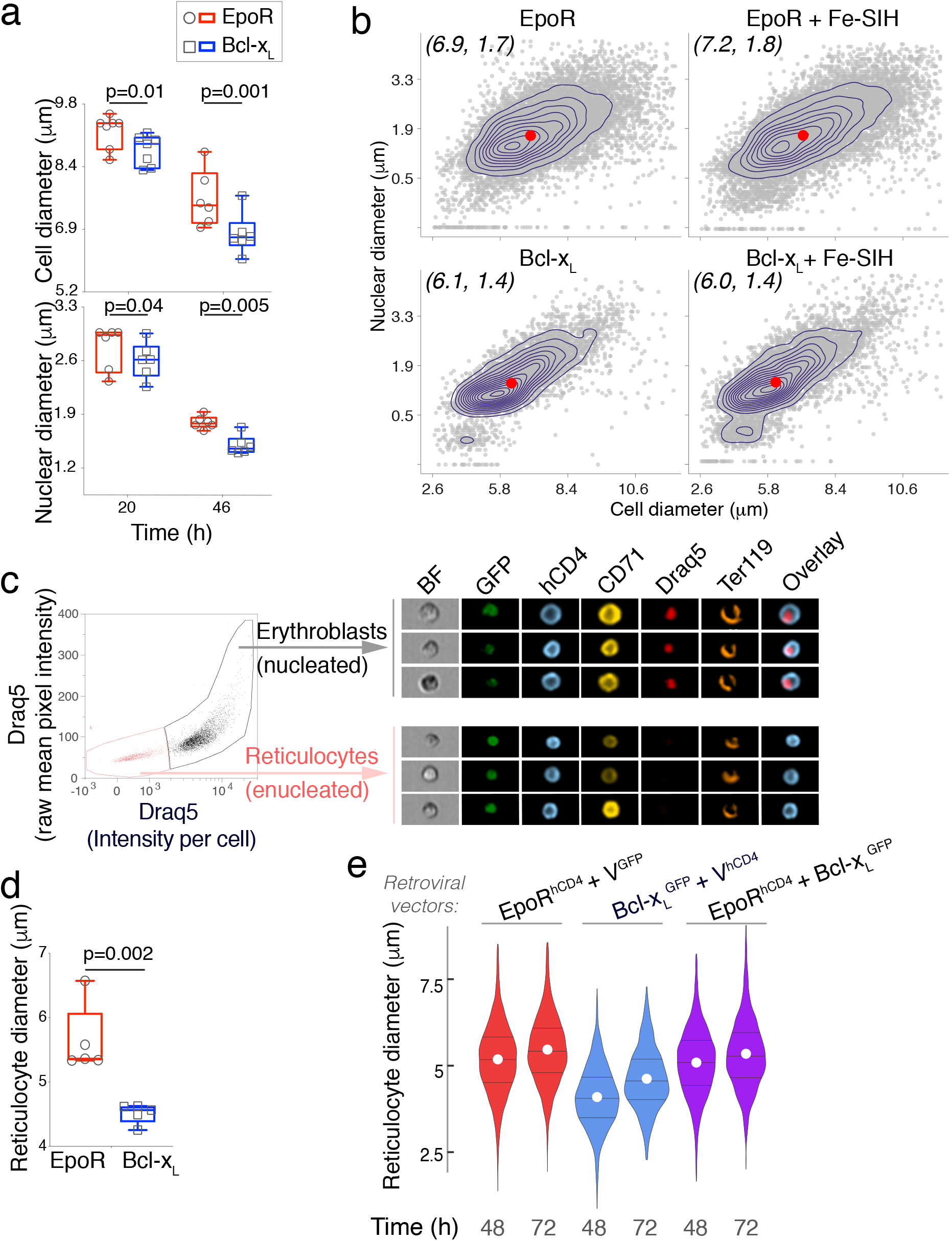
Smaller erythroblasts that differentiate into smaller reticulocytes in the absence of EpoR. **a** Cell and nuclear diameter of hCD4^+^Lin^-^ erythroblasts, measured by imaging flow cytometry. Experiment as in Fig 2b. Polystyrene beads of known diameters were used for calibration (see methods, **Extended Data Figure 4a**). Datapoints are population medians for individual samples, with 50,000 cells imaged per sample. Box and whiskers mark the 25^th^ to 75^th^ percentiles and min to max values, respectively, with the median indicated. Data pooled from 7 independent experiments, p-values are from 2-tailed paired *t*-tests, pairing EpoR and Bcl-x_L_–transduced cells in each experiment. **b** A representative experiment as in ‘a’, showing individual sample contour plots overlaid on scatter plots (each dot is one cell), of nuclear diameter vs. cell diameter. The effect of adding Fe-SIH to the culture medium is also shown. Data are hCD4^+^Lin^-^ erythroblasts at 48h post transduction, **c** Distinguishing erythroblasts from reticulocytes using imaging flow cytometry, with the nuclear dye Draq5. The analysis was performed on Ter119^+^ cells. Representative images are shown, from cultures of *Epor^-/-^* fetal liver cells that were doubly transduced with bicistronic retroviral vectors encoding GFP and hCD4 reporters (see **Extended Data Fig 1a**), at 48 hours post transduction. **d** Reticulocyte cell diameter in cultures of EpoR-Epor^-/-^ or Bcl-x_L_-*Epor^-/-^* at 48 h post transduction, identified as in ‘c’. Data are population medians from 5 independent experiments. Box and whiskers as in ‘a’. **e** Reticulocyte diameters in cultures of *Epor^-/-^* fetal liver cells that were doubly-transduced with bicistronic vectors carrying GFP and hCD4 reporters (**Extended Data Fig 1a**). These vectors were either ‘empty’ (V^GFP^, V^hCD4^) or encoded either Bcl-x_L_ or EpoR (Bcl-x_L_^GFP^, EpoR^hCD4^). Violin lines mark the 25^th^, 50^th^ and 75^th^ percentile with a white circle marking the mean. Representative of two independent experiments.

We asked whether the smaller size of Bcl-x_L_-*Epor^-/-^* erythroblasts could reflect an accelerated process of differentiation. If at any given time of the culture Bcl-x_L_-*Epor^-/-^* erythroblasts were smaller only as a result of being at a more advanced maturation stage, they should give rise to normally-sized enucleated reticulocytes, albeit at an earlier time. However, imaging flow-cytometry showed that Bcl-x_L_-*Epor^-/-^* reticulocytes were significantly smaller (5.6 ± 0.5 μm vs. 4.5 ± 0.15 μm for respective diameters of EpoR-*Epor^-/-^* vs. Bcl-x_L_-*Epor^-/-^* reticulocytes, mean ± sem, p = 0.002, **Fig 4c, d**).

It remained possible that the smaller size of Bcl-x_L_-*Epor^-/-^* erythroblasts and reticulocytes is the result of over-expression of Bcl-x_L_, rather than absent EpoR signaling. To address this, we doubly transduced *Epor^-/-^* fetal liver cells with both EpoR and Bcl-x_L_. We used the the Bcl-x_L_-linked GFP and the EpoR-linked hCD4 fluorescence reporters to quantify expression and ensured that all comparisons were made between cells expressing similar levels of each retroviral vector (**Extended Data Fig 5b-d**). We found that erythroblasts and reticulocytes transduced with both EpoR and Bcl-x_L_ were similar in size to those transduced with only the EpoR, and significantly larger than those transduced with only Bcl-x_L_ (**Fig 4e; Extended Data Fig 5c, d**). Therefore, Bcl-x_L_ over-expression is not the cause of the smaller size of Bcl-x_L_-*Epor^-/-^* erythroblasts and reticulocytes. Taken together, these observations suggested a dependence of erythroblast and reticulocyte size on EpoR signaling, that appeared to be independent of iron.

### EpoR regulation of red cell size is independent of HRI

In addition to the number of maturational cell divisions, red cell size is regulated by iron status. HRI is activated by heme deficiency and mediates the formation of smaller, hypochromic red cells^56^. It phosphorylates eukaryotic initiation factor 2α (eIF2α), inhibiting translation of globins and most other cellular transcripts. Our results showed that Bcl-x_L_-*Epor^-/-^* erythroblasts and reticulocytes were smaller and failed to upregulate CD71 (Tfrc), the principal iron transporter. Although iron supplementation did not appear to rescue the smaller size of Bcl-x_L_-*Epor^-/-^* erythroblasts (**Fig 4b**), it remained possible that intracellular iron delivery was somehow incomplete.

To determine definitively the relevance of the iron/heme/HRI pathway to the smaller size of Bcl-x_L_-*Epor^-/-^* erythroblasts, we generated mice that were doubly deleted for both *Epor* and *Hri* (**Fig 5a**). The *Epor^-/-^ Hri^-/-^* mice were phenotypically similar to *Epor^-/-^* mice, dying at mid-gestation as a result of severe anemia. We rescued both singly (*Epor^-/-^*) and doubly-deleted (*Epor^-/-^ Hri^-/-^*) fetal liver cells in parallel, by transduction with either Bcl-x_L_ or EpoR (**Fig 5b-e**). Bcl-x_L_-transduced erythroblasts were indeed larger on the *Epor^-/-^ Hri^-/-^* genetic background than on the *Epor^-/-^* background. However, EpoR-transduced erythroblasts were also larger on the *Epor^-/-^ Hri^-/-^* background, compared with the *Epor^-/-^* background. Further, for a given genetic background, either *Epor^-/-^ Hri^-/-^* or *Epor^-/-^* the difference in size between Bcl-x_L_ and EpoR-rescued cells remained (**Fig 5b-e**). These results are in agreement with the known role of HRI as a negative regulator of erythroblast size. In addition, they clearly show that EpoR signaling regulates cell size independently of the HRI pathway, since, even in the absence of HRI, EpoR signaling promotes larger erythroblasts (**Fig 5b-d**) and reticulocytes (**Fig 5e**).

**Figure 5.**
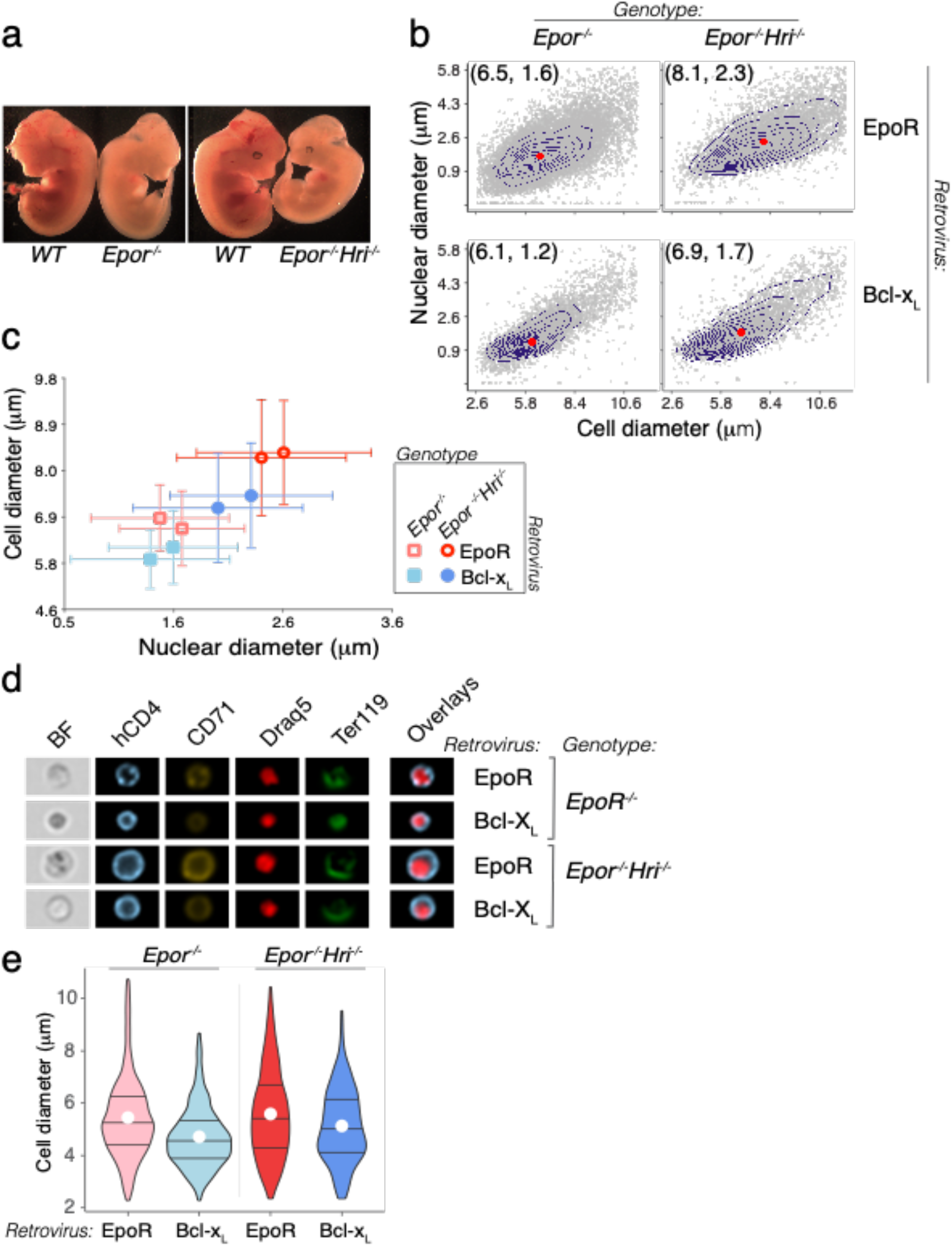
EpoR regulates cell size independently of HRI. **a** *Epor^-/-^* and doubly-deleted *Epor^-/-^ Hri^-/-^* E12.5 embryos with wild-type littermates **b** Cell and nuclear diameters in fetal livers from either *Epor^-/-^* or *Epor^-/-^ Hri^-/-^* embryos, transduced with either EpoR or Bcl-x_L_, at t=48h post transduction. Individual sample contour plots are overlaid on scatter plots (each dot is one cell). **c** Summary data for cell and nuclear area, for two independent experiments as in ‘b’, each containing all 4 genotype/retrovirus combinations. Data are mean ± SD for each cell population. Each transduced population consisted of pooled fetal liver cells from either *Epor^-/-^* or *Epor^-/-^ Hri^-/-^* embryos. **d** Imaging flow cytometry of representative Lin^-^hCD4^+^Ter119^+^ erythroblasts from each of the genotype/retroviral combinations at t= 48 h. **e** *Epor^-/-^* and *Epor^-/-^ Hri^-/-^* Reticulocyte cell diameter, from cultures transduced with either EpoR or Bcl-x_L_. Representative of 2 experiments. Violin lines mark the 25^th^, 50^th^ and 75^th^ percentile with a white circle marking the mean.

### Several EpoR signaling pathways are implicated in the regulation of cell size

The EpoR activates three principal signaling pathways: ras/MAP kinase, Stat5, and phosphatidylinositol 3-kinase (PI3K)^57,58^. Neonatal mice hypomorphic for Stat5 have microcytic anemia^59^. Here we found that, similarly, circulating red cells from E13.5 *Stat5*-deficient embryos are smaller than wild-type littermates (**Extended Data Fig 6a-b**). Therefore, Stat5 signaling contributes to the regulation of cell size by EpoR. To determine whether other pathways are implicated, we examined the effects of U0126, a MEK1- and MEK2-specific inhibitor^60^, and of LY 294002, a specific inhibitor of PI3K^61^, on differentiating fetal liver cells. We found that PI3K inhibition significantly decreased the size of early (‘S2’) and late (‘S3) erythroblasts and reticulocytes. By contrast, inhibition of MEK1/2 had no significant effect on reticulocyte size, though it appeared to result in larger early erythroblasts (**Extended Data Fig 6c, d**). Therefore, it is likely that cell size regulation by EpoR is the integrated result of multiple signaling pathways.

### Accelerated maturation in the absence of EpoR, assessed independently of cell size

Cell size is frequently used as a convenient indicator of maturational stage during ETD (^62–65^). Our initial impression was that Bcl-x_L_-*Epor^-/-^* erythroblasts completed their maturation sooner than EpoR-*Epor^-/-^* erythroblasts (**Fig 1e g**). However, the finding that cell size is consistently smaller in Bcl-x_L_-*Epor^-/-^* erythroblasts at all maturational stages, including reticulocyte, made cell size an unreliable indicator of maturational stage differences between Bcl-x_L_-*Epor^-/-^* and EpoR-*Epor^-/-^* erythroblasts. We therefore assessed maturation using two alternative measures.

First, we looked at the level of expression of Ter119, a cell surface marker that emerges with the onset of ETD and whose expression increases with maturation^62–65^. We found that Ter119 levels increased more rapidly in Bcl-x_L_-*Epor^-/-^* erythroblasts, reaching significantly higher levels than in EpoR-*Epor^-/-^* erythroblasts at 48 h (**Extended Data Fig 7a**). Second, we looked at nuclear offset, a quantitative morphological measure of nuclear eccentricity that is independent of cell size and that was previously used to assess erythroblasts prior to enucleation^65^ (**Extended Data Fig 7b-d**). Here we found that nuclear offset increases continuously throughout ETD. Early in ETD, when the nucleus is large, the geometrical centers (centroids) of the cell and the nucleus are close. With increasing maturation, the nucleus undergoes gradual condensation, and the distance between the cell and nuclear centroids, known as the ‘delta centroid’, gradually increases. The nuclear offset is the ratio of this distance to the cell diameter, making it independent of cell size (**Extended Data Fig 7b**). We found that nuclear offset increased earlier in Bcl-x_L_-*Epor^-/-^* erythroblasts, with the difference between Bcl-x_L_-*Epor^-/-^* and EpoR-*Epor^-/-^* erythroblasts peaking at 48 h (p=0.02) **(Extended Data Fig 7c, d**). Taken together, the timing of both Ter119 expression and nuclear offset suggest that EpoR signaling prolongs erythroblast maturation.

### Epo increases cell size and prolongs erythroid maturation over a wide concentration range

We next asked whether the effect of EpoR signaling on cell size is sensitive to Epo concentration. We harvested wild-type fetal liver CFU-e (‘S0’ in **Fig 6a**^40^) and plated them in the presence of a range of Epo concentrations. We followed their differentiation as they sequentially upregulate CD71 and Ter119^40^. By 48 h, the cultures contained a mixture of erythroblasts at various maturational stages (S1 to S3, **Fig 6a**) as well as reticulocytes. We found that cell size was increased in an Epo-concentration-dependent manner, with larger cells at every stage of differentiation, including reticulocytes (**Fig 6b, Extended Data Fig 8a**). The Epo concentration range affecting cell size, from 0.01 to 10 Units/ml, covers the entirety of the physiological and stress range that is found for Epo *in vivo*^66,67^. In addition to increasing reticulocyte diameter, higher Epo concentration also increased reticulocyte size distribution, as shown by respective coefficients of variation of the diameters, making them more heterogeneous (**Extended Data Fig 8b**).

**Figure 6.**
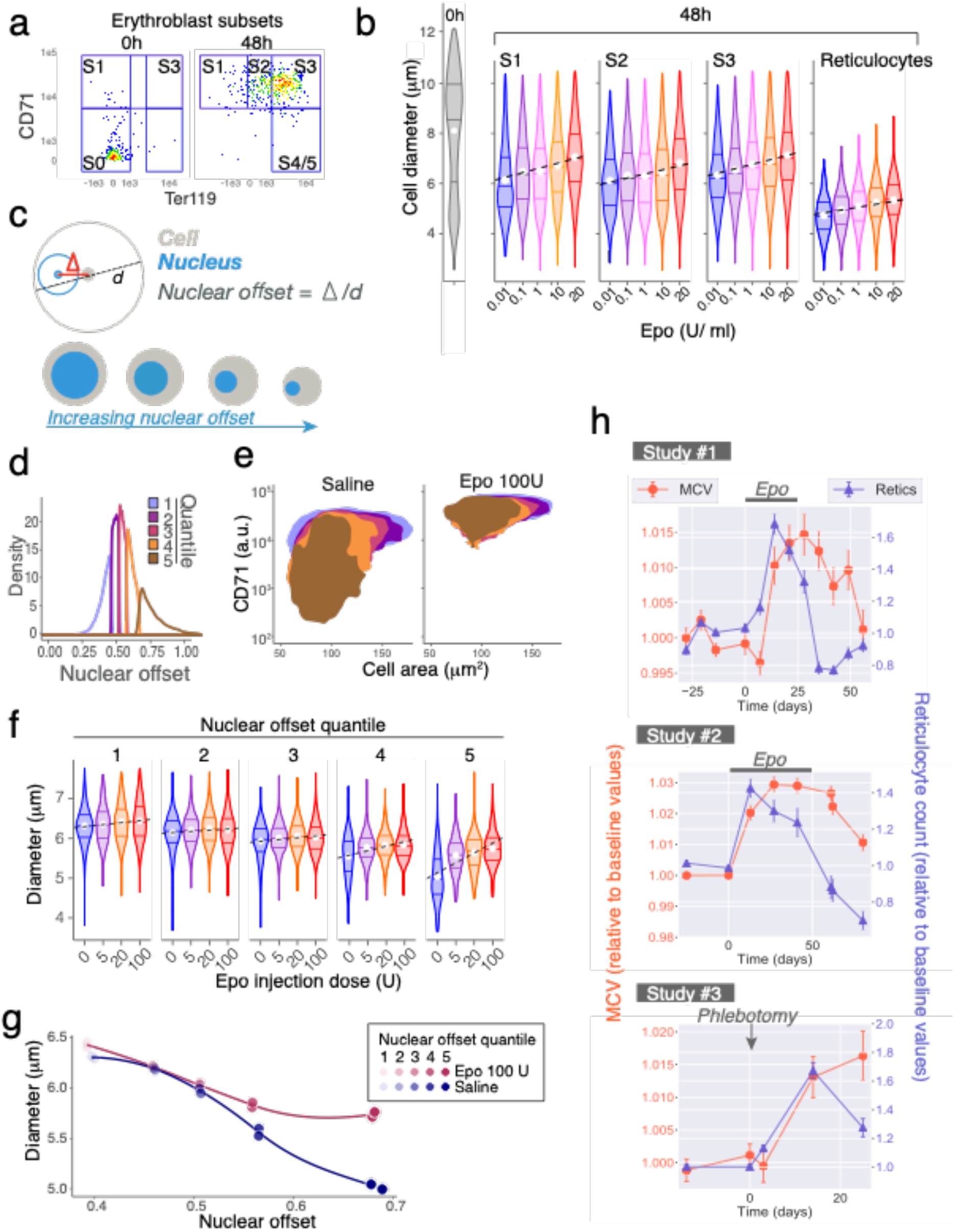
Red cell size is sensitive to Epo concentration in both mice and humans. **a, b** Epo concentration effect on ETD. Wild-type fetal liver cells were enriched for CFU-e progenitors (‘S0’) and differentiated *in vitro*, in a range of Epo concentrations between 0.01 and 20 U/ ml. Cultures were analyzed at 48 h. Data is representative of two independent experiments. **a** S0 cells upregulate cell surface markers CD71 and Ter119 during differentiation. At 48h cells are distributed between erythroblast subsets S1, S2 and S3. The shown example is from a culture in the presence of Epo at 0.1 U/ml. **b** Cell diameter distributions of S0 cells at t= 0h, and of erythroblast subsets S1 to S3 at 48 h, for cultures in the presence of the indicated Epo concentrations. Violin lines mark the 25^th^, 50^th^ and 75^th^ percentile with a white circle marking the mean. **c** Nuclear offset is the ratio of the delta centroid (distance between the centers of the cell and the nucleus, Δ) and the cell diameter. It is a measure of nuclear eccentricity that is independent of cell size. The cartoon indicates how nuclear offset can be used to measure the increasing nuclear eccentricity during erythroid morphological maturation. **d-g** Mice were injected with either saline (n = 2) or Epo (5 U, 20 U or 100 U, n=2 for each Epo dose), and bone marrow was analyzed at 48 h. **d** Ter119^+^ bone marrow erythroblasts in saline-injected mice were divided into 5 maturational stages, by dividing the nuclear offset distribution into quintiles. **e** CD71/ forward-scatter (FSC) histograms for Ter119+ erythroblasts in each of the nuclear offset quintiles. For mice injected with Epo, cells were divided into 5 maturational stages based on the nuclear offset values defined by the control (saline) nuclear offset quintiles. **f** Cell diameter for each of the nuclear offset quintiles in ‘d, e’, for each injected Epo dose. Violin lines mark the 25^th^, 50^th^ and 75^th^ percentile with a white circle marking the mean. Data are representative from one of two mice injected for each Epo dose. **g** Median cell diameter and median nuclear offset values for cells in each nuclear offset quintile, for mice injected with either Epo (100 U) or Saline. Each data point is for one mouse. **h** MCV and reticulocyte count in three independent human intervention studies. In studies #1 & #2, Epo was administered during the period indicated. In study #3, participants were subjected to phlebotomy at the time indicated (see text and methods for details). Because of individual variability between participants in many of the hematological parameters, data is represented as fractional change relative to the baseline values of each participant. All hematological parameters and statistical significance values as well as results for placebo control groups are in **Extended Data Figures 8-10** and **Supplementary statistical analysis**. MCV, mean corpuscular volume; Retics, reticulocyte count.

Epo also caused a dose-dependent lag in differentiation. As expected, at 48 hours of culture, higher Epo resulted in higher cell number at all stages of differentiation (**Extended Data Fig 8c**). However, the distribution of erythroblasts was increasingly skewed in favor of earlier differentiation subsets (**Extended Data Fig 8d**). Thus, at 0.01 U/ml of Epo, 5.4% of erythroblasts were in the earlier, S1 stage, and 62% had attained the more advanced, S3 stage; by contrast, at 10 U/ml, as many as 22% of erythroblasts were still in S1, and only 46% reached S3. Similarly, the intensity of Ter119 expression in erythroblasts pooled from all subsets (S1 to S3, **Extended Data Fig 8e**) decreased at higher Epo concentrations.

We further assessed cell maturation by measuring the nuclear offset, which decreased with increasing Epo concentration at all flow-cytometric stages (**Extended Data Fig 8e, f**). Together these findings suggest that Epo prolongs ETD in a dose-dependent manner. These observations are consistent with our finding that Bcl-x_L_-*Epor^-/-^* erythroblasts, which lack EpoR signaling altogether, have a significantly faster differentiation rate (**Fig 1e, Extended Data Fig 7**).

A skewed distribution in favor of early erythroblasts is also seen in *vivo* following Epo administration (e.g. **Extended Data Figure 3a**), consistent with prolongation of early ETD.

### Epo administration stimulates a dose-dependent increase in erythroblast cell size *in vivo*

To assess whether Epo increases erythroblast cell size *in vivo*, we injected mice with a range of Epo doses, or with saline control. Erythroblast maturational stage is routinely assessed using erythroblast cell size^62,63^ (e.g. **Extended Data Figure 3a**), but this approach precludes an assessment of cell size regulation. Therefore, to determine whether Epo increases erythroblast cell size at a given maturational stage, we used imaging flow cytometry to determine nuclear offset, a size-independent means of assessing maturational stage. Nuclear offset quantitates morphological maturation by measuring increasing nuclear eccentricity (**Fig 6c**). We divided the nuclear offset distribution of Ter119^+^ bonemarrow erythroblasts from saline-injected mice into quintiles (**Fig 6d**). Quintiles with increasing nuclear offset corresponded to increasingly mature erythroblast subsets as judged by the established criteria of decreasing CD71 and cell area, confirming the utility of this approach (**Fig 6e**). We then used the nuclear offset quintile values from these control mice to classify Ter119+ erythroblasts from Epo - injected mice into five maturational stages. We found that for a given nuclear-offset-defined maturational stage, cell diameter increased with increasing Epo dose. This effect was particularly striking in erythroblasts that corresponded to the two most mature quintiles (**Fig 6f, g**), clearly confirming that Epo dose regulates erythroblast cell size.

### Epo administration to healthy human volunteers increases red cell size (MCV) and size variation (RDW) for prolonged periods

To determine whether EpoR regulates cell size *in vivo* in humans, we examined healthy volunteers in three intervention studies. Participants were either given Epo (studies #1 and #2, **Fig 6h, Extended Data Fig 9,10**), or subjected to phlebotomy (study #3, **Fig 6h, Extended Data Fig 11**). In studies #1 and #2, the effect of Epo on athletic performance was examined, and will be either reported elsewhere (study #1) or was previously reported (study #2^68^). Here we present the detailed blood parameters associated with these studies.

In study #1 (**Fig 6h, Extended Data Fig 9**), subjects at sea level had baseline parameters established during four weekly blood samplings. They were then injected with either Epo (20 IU /kg every other day, 25 subjects) or placebo (9 subjects) for 3 weeks. On average, hemoglobin levels increased by 5% over baseline values in the Epo group during the treatment period. Blood sampling continued for an additional five weeks following cessation of treatment. In study #2 (**Fig 6h, Extended Data Fig 10**), baseline measurements were followed by weekly dosing with Epo (24 subjects) or placebo (24 subjects) for 7 weeks. Epo dosing (5000 to 10000 IU) was adjusted for each subject, to achieve an increase of 10 to 15% in hemoglobin over baseline. Follow-up continued for a month after cessation of treatment. In study #3 (**Fig 6h, Extended Data Fig 11**), 21 subjects participated in a randomized double-blind placebo-controlled crossover study in which 900 ml of whole blood was withdrawn from the treatment group by venipuncture. Subjects were then followed for 25 days.

In all three studies, there was a significant increase in MCV in the treatment groups compared with baseline values and with the placebo group, which persisted well beyond the treatment period (**Extended Data Fig 9-11, Supplemental statistical analysis**). There was no correlation between MCV and the reticulocyte count, whose time courses were clearly divergent (r ≤ 0.1 between MCV and reticulocyte count in all three studies, Pearson’s product-moment correlation, **Supplemental statistical analysis**). In studies #1 and #2, the reticulocyte count increased during Epo treatment, but declined sharply below baseline values as soon as Epo treatment ceased. By contrast, MCV values remained high (**Fig 6h**). Similarly, in study #3, MCV values continued to climb at a time when the reticulocyte count was declining (**Fig 6h**). Thus, the increase in MCV is not the result of an increase in the number of reticulocytes. Together with the increase in MCV, there was an increase in red-cell distribution width (RDW-SD, **Extended Data Fig 9,10**; no RDW is available for study #3). There was a significant, positive correlation between MCV and RDW-SD (r = 0.51, p=2 × 10^-28^ for study #1; r = 0.52, 2 × 10^-24^ for study #2).

Although the largest loss in red cell volume occurs during reticulocyte maturation, red cell size continues to decline as a function of red cell age^69–73^. The MCV represents the average volume of all circulating cells; therefore, we considered the possibility that the persistently elevated MCV following Epo administration might be the result of the expected increase in the relative number of younger red cells, rather than an increase in their size. To address this, we simulated the expected increase in MCV that would arise only from an increase in the proportion of younger red cells, assuming no effect of EpoR signaling on red cell size (supplementary materials, ‘Simulation of MCV’). This simulation indicated that an increased proportion of younger red cells cannot fully account for the extent or duration of the observed increase in MCV following Epo administration, consistent with a direct role for EpoR signaling in the regulation of cell size.

## Discussion

Using a genetic model in which we provided *Epor^-/-^* fetal liver cells with exogenous survival signaling, we identified novel non-redundant functions for EpoR during ETD. We found that EpoR signaling determines the number of cell divisions, their speed, as well as the duration of terminal differentiation. While it has little effect on the broad ETD transcriptional program, it drives the formation of qualitatively different, larger red cells. Human intervention studies are consistent with a similar effect of EpoR signaling on red cell size in human erythropoiesis. In the discussion below we integrate these apparently disparate effects into a coherent model for EpoR signaling during terminal differentiation (**Fig 7**). We discuss previously unexplained instances of macrocytic and heterogeneously-sized red cells, now interpretable as the result of EpoR signaling during hypoxic stress.

**Figure 7.**
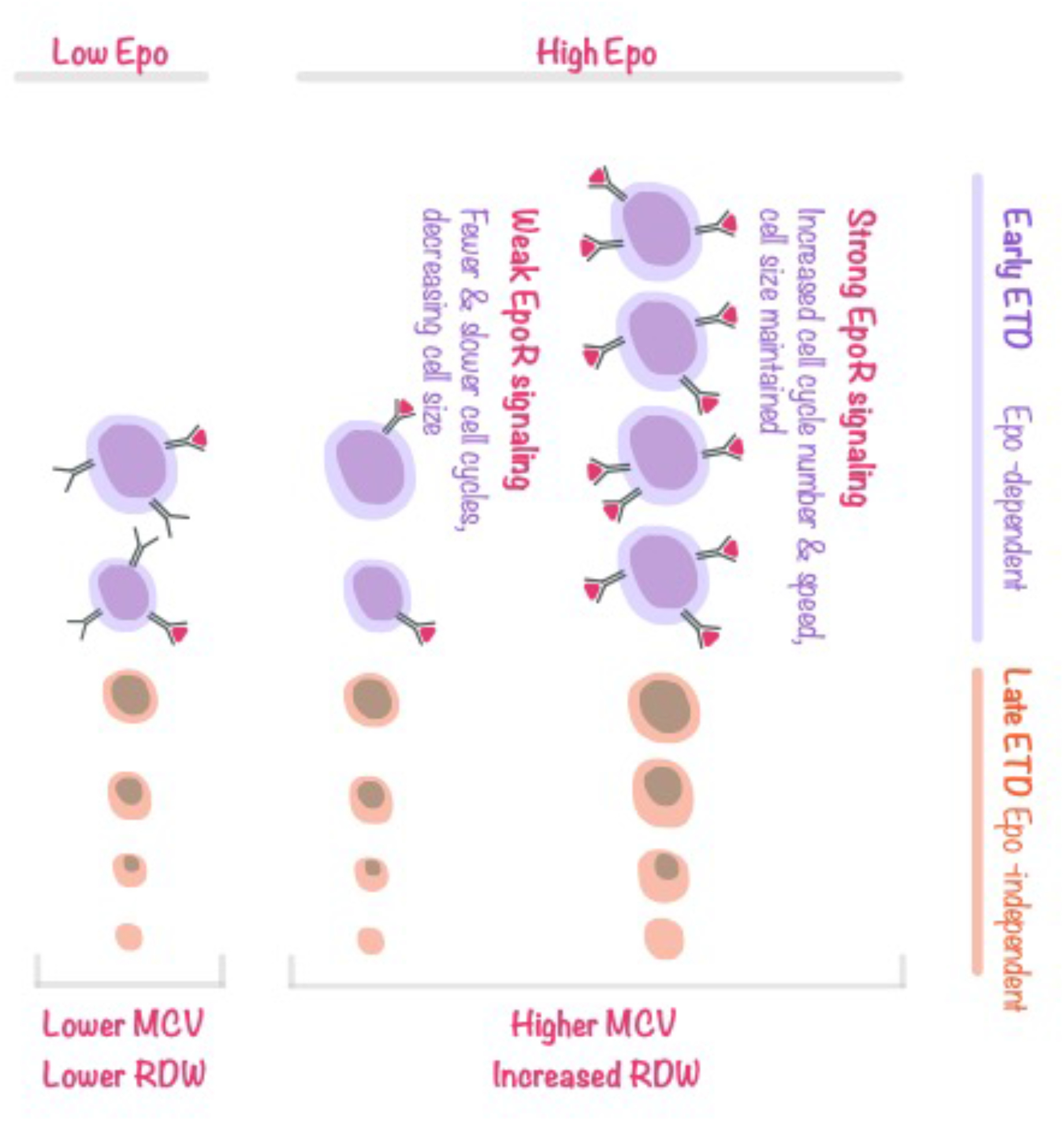
EpoR signaling promotes rapid cycling while maintaining cell size in early erythroblasts. Proposed model explaining EpoR-dependent functions during ETD. Only early erythroblasts express appreciable cell surface EpoR and are sensitive to EpoR signaling. When EpoR signaling is weak or absent, as in late erythroblasts, or in early erythroblasts in the presence of low Epo, cell divisions lead to a loss in cell size. In contrast, strong EpoR signaling, as seen in Epo-sensitive early erythroblasts, can override this default state and simultaneously increase rapid cycling while maintaining cell size. As consequence, high Epo levels increase the duration of the early ETD phase, increase the relative frequency of early erythroblasts, and result in larger erythroblasts at every maturation stage, giving rise to larger red cells. In high Epo, red cell size is also more heterogeneous, a result of the varying sensitivities of early erythroblasts to Epo. Erythroblasts with low sensitivity to Epo, here represented as cells expressing low levels of EpoR, are expected to attain only weak EpoR signaling even in the presence of high Epo, giving rise to smaller red cells.

The ETD is a time of continuous and rapid change in many aspects of the cell, including transcriptional state, susceptibility to apoptosis, cell cycle dynamics, cell size and morphology. Our results strengthen a model in which these changes do not progress uniformly throughout ETD, but rather, are controlled differently in two principal phases: an early, Epo-dependent phase, and an Epo-independent late phase^39^ (**Fig 7**). EpoR expression peaks in early erythroblasts, which are exquisitely sensitive^74^ and dependent on EpoR signaling for survival^8,62,75^ By contrast, late erythroblasts downregulate EpoR expression^13^ and are relatively resistant to apoptosis^62,75^. The functions we identified here for EpoR signaling in ETD are similarly focused on early erythroblasts.

Our work documents five key effects of EpoR signaling in addition to its anti-apoptosis effects: 1) it prolongs the duration of ETD, as determined by delayed expression of cell surface markers and a delayed increase in nuclear offset, compared with cells lacking the EpoR; 2) it increases the number of cell cycles; 3) it skews the distribution of developing erythroblasts in favor of earlier erythroblasts; 4) it increases cell cycle speed; and 5) it increases cell size throughout ETD, generating red cells that are larger and more heterogeneous in size. The prolongation of ETD is consistent with the increase in the number of cycles. Neither informs us directly regarding the stage(s) of ETD that are being prolonged. However, the skewed distribution in favor of early erythroblasts indicates, based on the ergodic principle^76^, that EpoR signaling prolongs early ETD relative to the late ETD phase. Together, these observations suggest that EpoR prolongs the early phase of ETD by increasing the number of early ETD cell cycles. This conclusion is consistent with our data, showing the largest differences in cell cycle number in the first 24 h of ETD; and with the known responsiveness of early ETD to EpoR signaling. In addition, it explains the observation that EpoR increases cell cycle speed, since early ETD cell cycles are unusually fast^27,28^, and much faster than cycles in late ETD^27,28,77^; our observations show that EpoR signaling regulates the speed of these unique cycles.

Therefore, of the five effects of EpoR signaling that we document, the first four are outcomes of an EpoR-driven increase in the number and speed of early ETD cell cycles (**Fig 7**). An increase in the number of these cycles prolongs the early phase of ETD, accounting for the relative increase in the number of early erythroblasts and for the increased duration of ETD as a whole.

One of the factors known to regulate the onset of late ETD is p27^KIP1^, whose induction promotes slower cycling and cell cycle exit^51,78,79^. The expression of cell cycle regulators was largely unchanged in the absence of EpoR signaling, with the exception of p27^KIP1^, which was induced prematurely (**Extended Data Fig 4**). A second CDK inhibitor, p57^KIP2^, was also elevated (**Extended Data Fig 4**). Interestingly, premature p27^KIP1^ expression and morphological maturation were noted in primitive yolk-sac erythroblasts cultured in the absence of Epo^21^. The converse was found in *Klf1^-/-^* erythroblasts, which fail to induce p27^KIP1^ and fail to undergo cell cycle exit; exogenous expression of p27^KIP1^ rescues mitotic exit in these cells^54^, and also promotes terminal differentiation in K562 erythroleukemia cells^80^. It is therefore likely that EpoR signaling extends the rapid cycling phase in early ETD, at least in part, by delaying the induction of p27^KIP1^. This is also consistent with a previous report showing that EpoR downregulates p27^KIP1 53^. As expression of EpoR decreases in the later stage of ETD, its function in maintaining the rate of cell cycle decreases while its anti-apoptotic function is assumed by increased Bcl-x_L_^81^.

The most surprising of our findings was the effect of EpoR signaling on cell size. We found that erythroblasts differentiating in the absence of EpoR gave rise to smaller red cells, in spite of undergoing fewer cell cycles. Further, cell size was sensitive to Epo concentration within the physiological and stress range^67^. These findings appear contrary to the well-established link between the loss in cell size and the number of cell divisions. Thus, deletions of E2F4^29^, cyclin D3^30^, CDK2 or CDK4^31^ each reduce the number of cell divisions during ETD and result in macrocytic red cells. Similarly, macrocytic red cells are seen when nucleotide pools limit DNA synthesis rate, as in patients treated with hydroxyurea^32^, or in B12 or folate deficiencies. We found that the EpoR effect on red cell size was also independent of a second established pathway, in which red cell size is regulated in response to iron status. Iron and heme deficiency activate HRI, which inhibits eIF2α and global protein translation, resulting in small, hypochromic red cells^34,56^. Here we found that erythroblasts lacking the EpoR failed to upregulate the transferrin receptor (CD71), potentially leading to intracellular iron deficiency. However, iron supplementation did not rescue the smaller size of *Epor^-/-^* erythroblasts. Further, the cell size deficit persisted in erythroblasts that were doubly deleted for both *Epor* and *Hri*. While this experiment does not exclude an interaction between HRI and EpoR signaling^35^, it shows conclusively that EpoR signaling regulates red cell size independently of HRI.

Our data therefore suggest that EpoR regulates red cell size through a novel mechanism. The finding that the EpoR-driven increase in cell size begins in early erythroblasts (**Fig 6b, Extended Data Fig 8a**) suggests that it takes place in the very same cells in which EpoR signaling also induces additional rapid cycles. We propose that the well-established coupling of cell size loss with cell divisions is a default state, seen in cells where EpoR signaling is weak or absent. We further suggest that strong EpoR signaling, as may occur in early erythroblasts^74^, can override this default state and maintain cell size in spite of rapid cycling (**Fig 7**). The maintenance of cell size in dividing cell populations is the norm in most tissues^82,83^ and so it is possible that EpoR signaling permits early erythroblasts to employ similar pathways of size control as those found outside ETD. The mechanisms that determine the characteristic size of a cell and that maintain it through cell divisions are not fully understood, but are thought to depend on strong growth factor signaling to promote the metabolic pathways required for building biomass^83^. To maintain their size, cells must attain a size threshold before committing to cell division; in avian erythroblasts and other cell types, a larger size correlates with a longer G1 phase^82,84^. The ability of EpoR signaling to increase cell size in early erythroblasts, which are some of the most rapidly dividing cells *in vivo*^27,28^, predicts that these cells have exceptionally efficient mechanisms for growth. Conversely, this also implies that impairments in growth pathways would have a specifically deleterious effect in early erythroblasts, potentially contributing to the selective damage of ribosomopathies in the erythroid lineage^85^.

Together with an increase in cell size, high Epo levels also increased cell size heterogeneity, in both mouse (**Extended Data Fig 8b**) and human (RDW-SD, **Fig 6d, Extended Data Fig 9, 10**). Unlike low Epo levels, which generate only weak EpoR signals and therefore relatively uniform small cells, high levels of Epo might be expected to support the survival of erythroblasts with varying Epo sensitivities^62,86^, in which the strength of EpoR signaling may vary widely, giving rise to a range of red cell sizes (**Fig 7**).

The relationship between high MCV, high RDW and high levels of Epo may have been overlooked previously by being attributed to an increase in reticulocytes. We have excluded this possibility, finding no correlation between reticulocyte numbers and MCV. High MCV persisted well after Epo and reticulocytes declined. We also found that the extent and duration of increase in MCV following Epo administration cannot be accounted for solely by the skewing in the age distribution of circulating red cells in favor of younger cells (see supplementary data, ‘simulation of MCV’). Indeed, our mouse data show increased cell size throughout terminal differentiation, generating larger than normal reticulocytes, and not simply more numerous reticulocytes.

Recent GWAS and other studies have linked multiple genomic loci to the regulation of MCV^87–90^. These include Epo, EpoR and lnk, all which might be expected to alter EpoR signaling strength^91^. An Epo-mediated increase in MCV in clinical settings might be tempered by iron status or by pathology affecting terminal differentiation. Indeed, Nevertheless, our work predicts that in the absence of erythroid pathology or nutritional deficiencies, Epo levels might be a key determinant of MCV. Indeed, an increase in Epo levels might account for the hitherto unexplained macrocytosis in hypoxemic patients with chronic obstructive pulmonary disease^92,93^ and in iron-replete pregnancy^94,95^. An increase in RDW was recently proposed as a potential longer-term biomarker for brief hypoxemic episodes in conditions such as acute respiratory distress, sepsis or congestive heart failure^96,97^. The regulation of MCV by Epo levels also clarifies previously unexplained changes in red cell volume associated with Kit function. Kit regulates the proliferation of early erythroid progenitors but is downregulated with entry into ETD. Gain of function Kit mutations in mice lead to erythrocytosis as a result of excess progenitors entering ETD; the red cells are microcytic^98^, presumably in response to a compensatory decrease in Epo levels. Conversely, loss of function Kit mutations are associated with an increased MCV, which is in proportion to the severity of anemia^98,99^, and can be explained by a paucity of progenitors entering ETD and the expected compensatory increase in Epo^100^. Transgenic expression of Epo rescues the lethal c-Kit^W/W^ mutation, also resulting in macrocytic red cells^99^. Given the persistence of higher MCV and RDW beyond the period in which Epo is elevated, these markers may be useful in detecting hypoxic stress in the clinic as well as Epo doping by athletes.

The adaptive value, if any, of a higher MCV in erythropoietic stress is not yet clear. Surprisingly, the increase in MCV in our human intervention studies was not associated with increased corpuscular hemoglobin (MCH). On the contrary, we found a statistically significant decrease in mean corpuscular hemoglobin concentration (MCHC) in both Epo intervention studies (**Extended Data Fig 9, 10**), though not in the phlebotomy intervention (**Extended Data Fig 11**). It is of interest, however, that a lower MCHC enhances the action of 2, 3, diphosphoglycerate (2,3 DPG), an allosteric regulator that binds to hemoglobin and lowers its affinity for oxygen. The levels of 2,3 DPG in red cells increase in response to anemia or hypoxia, improving the unloading of oxygen in tissues^101,102^. The affinity of 2,3-DPG to hemoglobin increases significantly at lower MCHC^103^. A lower MCHC may therefore improve the 2,3-DPG-dependent unloading of oxygen in metabolically active tissues during erythropoietic stress. Indeed, a lower MCHC is also an HRI-regulated outcome characteristic of microcytic iron-deficiency anemia, possibly for similar reasons. The EpoR-regulated increase in MCV might therefore provide a mechanism for lowering MCHC and improving oxygen unloading in tissues during hypoxic stress.

## Acknowledgements

The authors would like to thank the UMASS Flow cytometry core and Susanne Pechhold for her help with Imaging flow cytometry. Authors of the human intervention studies in Copenhagen (JB, NBN) wish to thank all participants of the studies as well as Thomas Christian Bonne and Andreas Breenfeldt Andersen (Department of Nutrition, Exercise and Sports, University of Copenhagen, Denmark), Mikkel Gybel-Brask (Section for Transfusion Medicine, Capital Region Blood Bank, Copenhagen University Hospital, Denmark) and Carl-Christian Howard Kitchen (Department of Anesthesiology, Copenhagen University Hospital, Denmark) for excellent assistance throughout the studies. Partnership for Clean Competition and Anti-Doping Denmark funded the phlebotomy trial and World Anti-Doping Agency funded the Copenhagen erythropoietin treatment trial.

This work was funded by NIH R01DK100915, R01DK120639 and R01HL141402 (MS), R25GM113686 (DH) and by R01DK087984 (JJC). JB was funded in part by Partnership for Clean Competition and Anti-Doping Denmark.

## Supplementary materials

1. **Statistical analysis of hematological parameters** in the human studies
2. **Simulations of MCV:** This simulation tests whether the null hypothesis, that EpoR signaling *does not* change red cell size, can account for the observations in the human studies

