## Extended Data Figures, Materials and Methods for "The Erythropoietin Receptor Stimulates Rapid Cycling and Formation of Larger Red Cells During Mouse and Human Erythropoiesis" (from another version of this preprint: version 1, 2020-11-30)

**Extended Data Figures and Figure Legends**

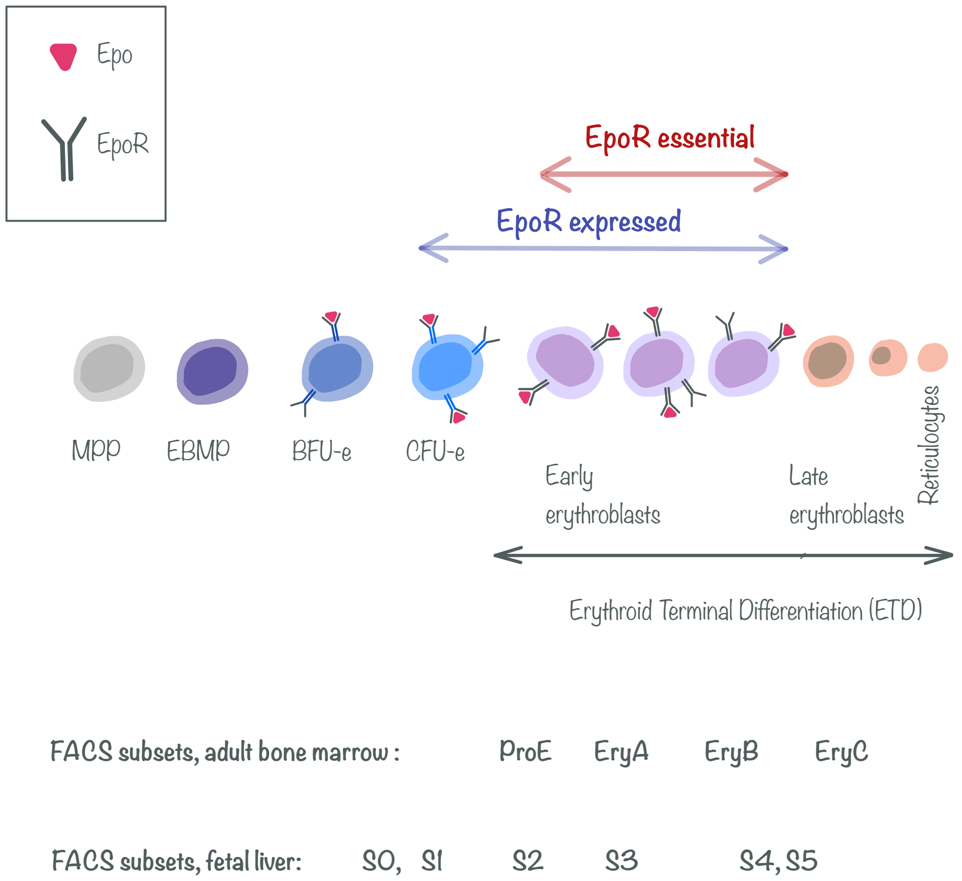


**Extended Data Figure 1 Developmental stages of erythropoiesis when EpoR is expressed or essential.**
The principal stages of the erythroid trajectory [1]. EpoR expression is induced at the BFU-e stage, and is downregulated in late erythroblasts. It is essential for the survival of early erythroblasts. FACS subsets used in the analysis of mouse fetal liver or adult bone marrow are also indicated; their horizontal position corresponds to their approximate developmental stage. MPP= Multipotential progenitors; EBMP = Erythroid, Basophil/mast cell, Megakaryocytic progenitors; BFU-e = Burst -forming unit erythroid; CFU-e = Colony-forming unit erythroid

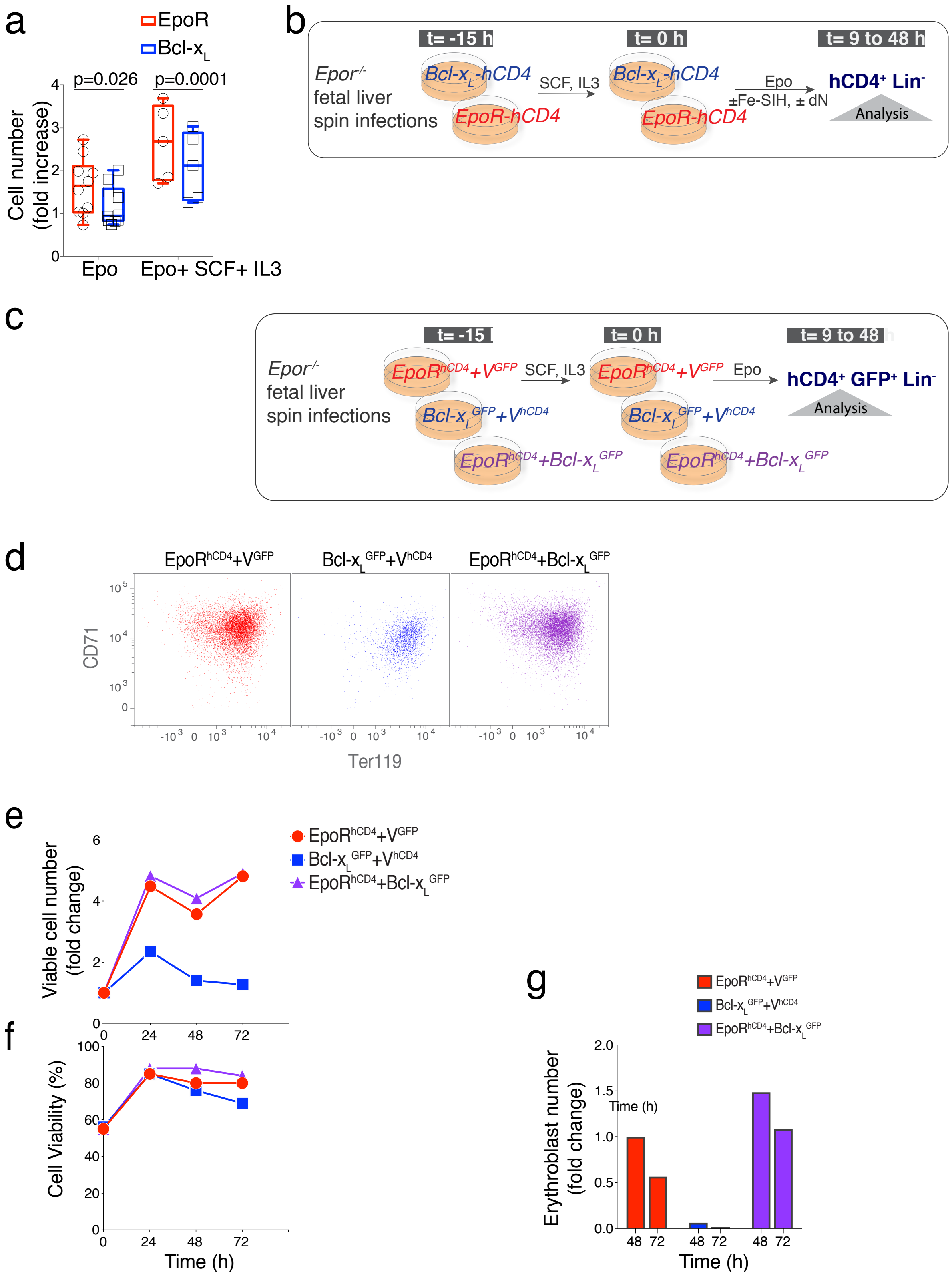


**Extended Data Figure 2** **Reduced growth of Bcl-x$_L$- *Epor$^{-/-}$* erythroblasts and their failure to upregulate CD71 persists in SCF and is not the result of bcl-x$_L$ over-expression**

**a** Effect of IL-3 and SCF on viable cell number. *Epor*$^{-/-}$ fetal livers were transduced as in Fig 1a, and cultured for 19 hours, in the presence of Epo, and either in the presence or absence of IL-3 and SCF. Trypan-blue negative cells were scored. Data points are independent cultures pooled from three independent experiments.

**b** Modified experimental design, used in all subsequent experiments unless otherwise indicated. *Epor*$^{-/-}$ fetal livers were transduced at t= -15 h with bicstronic retroviral vectors encoding Bcl-$x_L$ or EpoR and the hCD4 reporter and cultured in the presence of IL-3 and SCF for 15 hours. At t= 0 h, they were switched to an Epo containing medium, in the presence of additional factors as indicated in each experiment. Analysis was performed on transduced cells expressing hCD4 and negative for non-erythroid lineage markers (CD41, Mac1, Gr1, B220, CD3e).

**c** Experimental design: *Epor*$^{-/-}$ fetal livers were doubly transduced with bicistronic retroviral vectors, in the three shown combinations. Vectors encoded Bcl-$x_L$, EpoR or 'empty vector' (V); reporter constructs (GFP or hCD4) for each vector are shown as a superscript. Transduced cells were cultured in IL-3 and SCF for 15 hours, and switched to an Epo containing medium at t=0. Analysis was performed on cells that were doubly positive for both hCD4 and GFP, and negative for non-erythroid lineage markers (hCD4$^+$GFP$^+$Lin$^-$, where Lin= CD41, Mac1, Gr1, B220, CD3e).

**d** Flow cytometric analysis of *Epor*$^{-/-}$ hCD4$^+$GFP$^+$Lin$^-$ erythroblasts, showing failure to upregulate CD71 in cells lacking the EpoR, but not in cells doubly transduced with both EpoR and Bcl-$x_L$.

**e** Growth curves of doubly transduced *Epor*$^{-/-}$ fetal liver cells as in 'a'. Viable hCD4$^+$Lin$^-$ cells were counted at the indicated time points. Data are from two independent experiments, expressed relative to t=0.

**f** Cell viability, expressed as the fraction (%) of trypan blue negative cells, for the same set of samples shown in 'c'.

**g** Relative cell number of doubly transduced erythroblasts ( Lin-hCD4+GFP+), for each of the indicated retroviral combinations at 48 and 72 hours of differentiation.

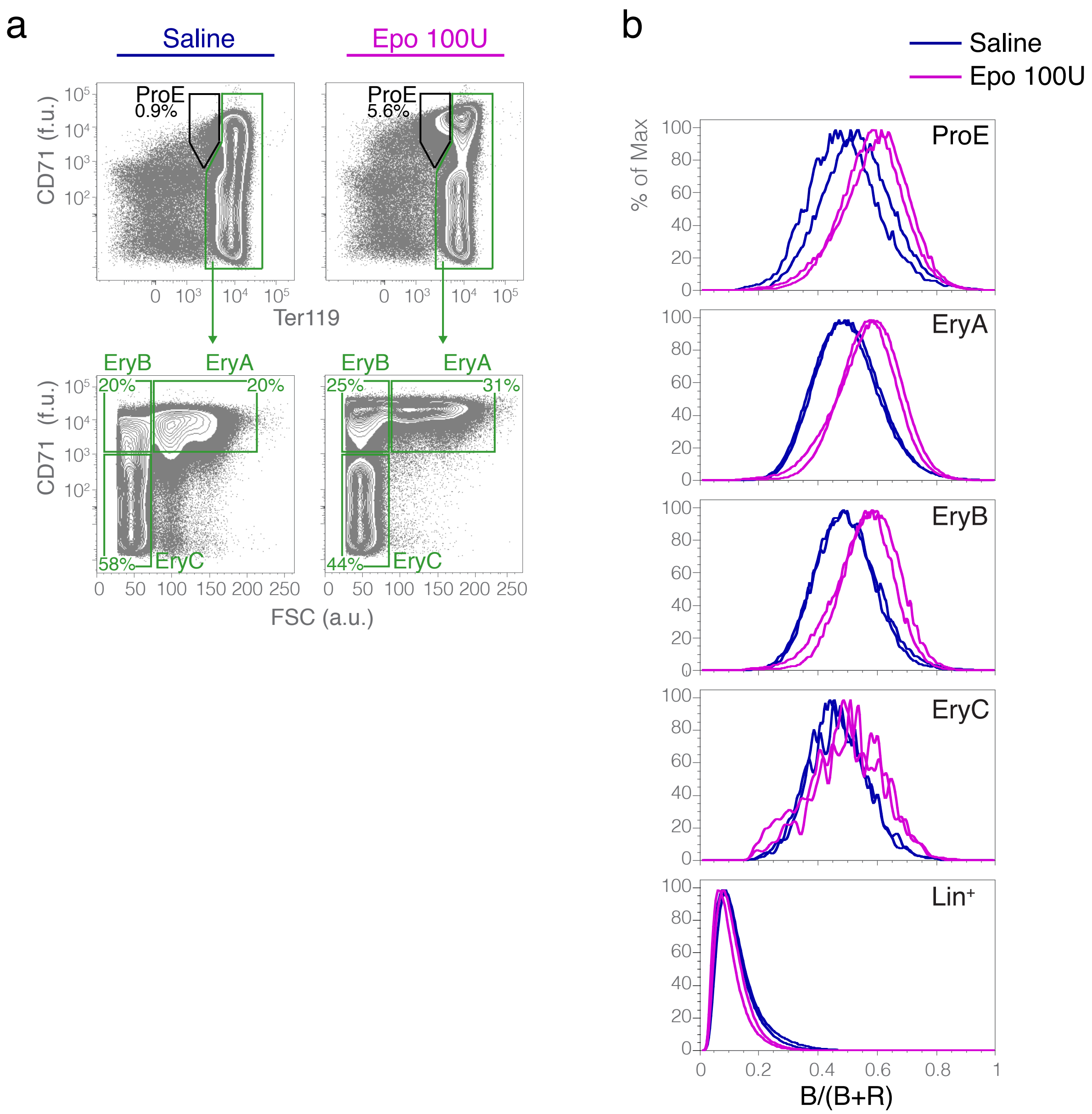


**Extended Data Figure 3 EpoR stimulates cell cycle shortening *in vivo***

**a, b** Mice transgenic for the fluorescent timer protein fusion H2B-FT were injected with either saline or Epo (100 U) and analyzed at 48 h (see Figure 2f for additional detail).

**a** Representative flow cytometric profiles of bone marrow erythroblasts

**b** Histograms of *B/(B+R)* in live cells, for each of the erythroblast subsets gated in ‘a’, and for non-erythroid lineage-positive cells in the same bone-marrow sample (‘Lin+’). Histogram overlays are for 2 mice injected with saline and 2 mice injected with Epo.

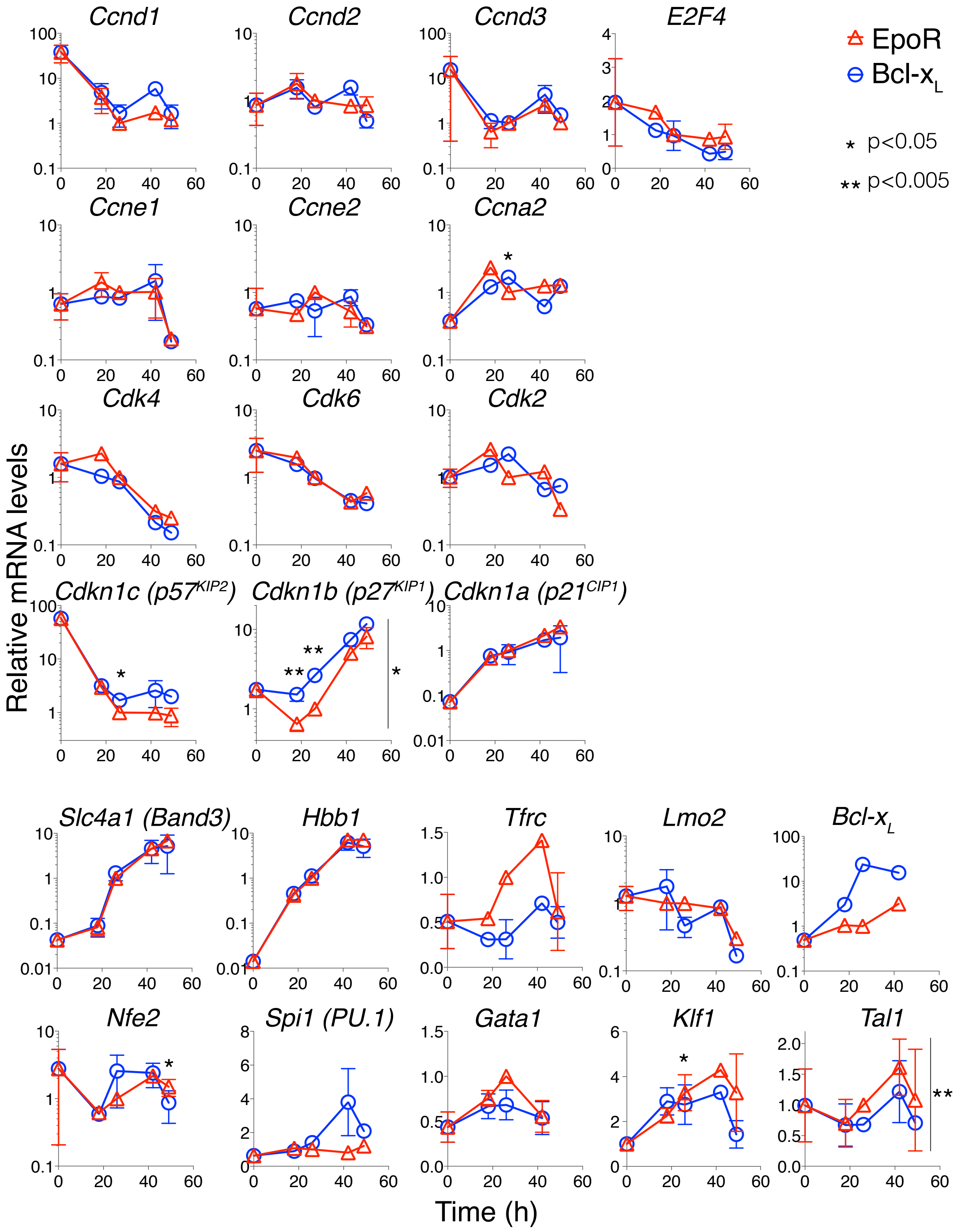


**Extended Data Figure 4** **Premature expression of $p27^{KIP1}$ in the absence of EpoR**

Gene expression in Bcl-$x_L$ or EpoR-transduced *Epor*$^{-/-}$ fetal liver cells during their differentiation in vitro. mRNAs were quantitated with RT-qPCR and are expressed relative to expression of β-actin in the same cells.

In each experiment, RT-qPCR was carried out using a dilution series of the cDNA. Data for each gene is pooled from a number of independent as follows: Ccnd1, n=6; Ccnd2, n=7; Ccnd3, n=6; E2F4, n=5; Ccne1, n=5; Ccne2, n=6; Ccna1, n=5; Cdk4, n=3; Cdk6, n=3; Cdk2, n=3; Cdkn1c ($p57^{KIP2}$), n=7; Cdkn1b ($p27^{KIP1}$), n=7; Cdkn1a ($p21^{CIP1}$), n=2; Slc4a1(Band3), n=6; Hbb1, n=7; Tfrc, n=6; Lmo2, n=6; Bcl-$x_L$, a representative experiment of 3; Gata1, n=7; Spi1 (PU.1), n=7; Tal1, n=6; Nfe2, n=6. p values are paired t-test.

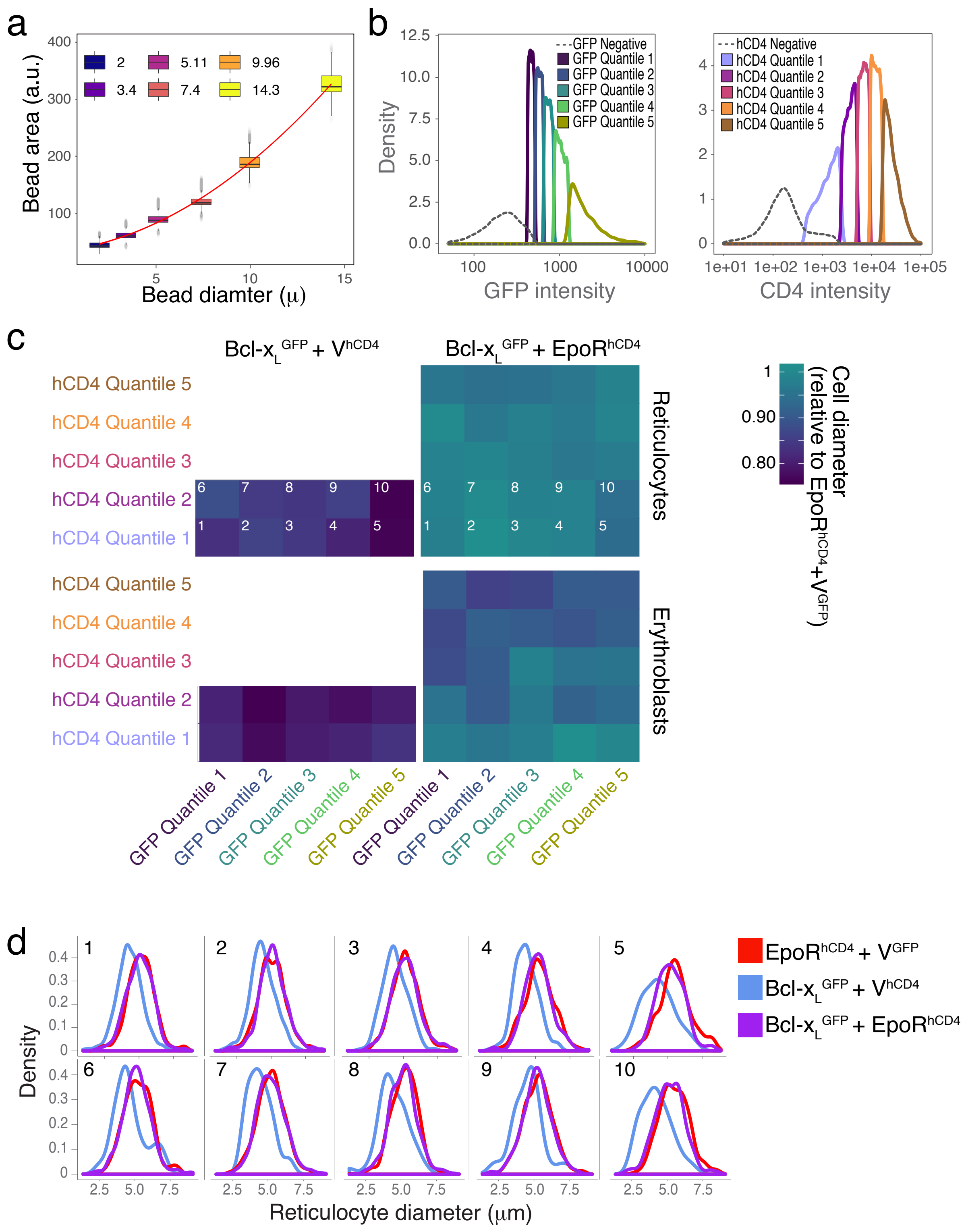


**Extended Data Figure 5 A smaller cell size in the absence of EpoR is not the result of Bcl-x$_L$ over-expression**

**a** A representative calibration of the cell diameter measurement. Imaging flow cytometry was used to measure the area of polystyrene beads (Spherotech) of known diameter (indicated in legend, in micrometers). Data were fitted with a polynomial curve. The box and whiskers mark the $25^{th}$ to $75^{th}$ percentiles and min to max values, respectively, for all data-points with the median indicate. Calibration was performed for each imaging flow cytometry experiment.

**b - c** *Epor$^{-/-}$* fetal livers were doubly transduced with bicistronic retroviral vectors, in three combinations, as described in **ED Figure 2**: EpoR-IRES-hCD4 + 'empty' GFP vector ($EpoR^{hCD4}$ + $V^{GFP}$); Bcl-$x_L$-IRES-GFP + 'empty' hCD4 vector (Bcl-$x_L^{GFP}$ + $V^{hCD4}$); or with Bcl-$x_L$-IRES-GFP + EpoR-IRES-hCD4 (Bcl-$x_L^{GFP}$ + $EpoR^{hCD4}$ ). Transduced cells were cultured in IL-3 and SCF for 15 hours, and switched to an Epo containing medium at t= 0 h. Analysis was performed on cells that were doubly positive for both hCD4 and GFP, and negative for non-erythroid lineage markers ($hCD4^+GFP^+Lin^-$, where Lin= CD41, Mac1, Gr1, B220, CD3e). The analysis below was performed at t= 48 h.

**b** In order to compare cells with similar expression of each of the retroviral constructs, we divided the erythroblast population ($hCD4^+GFP^+Lin^-$) into five fluorescence intensity quantiles for each of the GFP and hCD4 channels. The dashed line indicates the baseline fluorescence of cells that are not transduced with the corresponding vector (that is, cells transduced with only a GFP vector when examining the hCD4 channel, and vice versa).

**c** Erythroblast (upper panel) and reticulocyte (lower panel) cell diameters for cells expressing Bcl-$x_L^{GFP}$ + $V^{hCD4}$, compared with cells expressing Bcl-$x_L^{GFP}$ + $EpoR^{hCD4}$. The comparison is done separately for each quantile combination, and is depicted relative to the diameter of cells expressing $EpoR^{hCD4}$ + $V^{GFP}$ in the same quantile combination. For example, the diameters of cells transduced with Bcl-$x_L^{GFP}$ + $V^{hCD4}$ in quantile combination 10 (containing cells in the $5^{th}$ quantile of the GFP channel and $2^{nd}$ quantile of the hCD4 channel), are compared with diameters in quantile combination 10 of cells transduced with Bcl-$x_L^{GFP}$ + $EpoR^{hCD4}$, and expressed as a ratio to the diameters in quantile combination 10 of cells transduced with $EpoR^{hCD4}$ + $V^{GFP}$. Quantile combinations containing fewer than 10 cells were excluded.

**d** Reticulocyte diameters for each quantile combination, represented as histograms. Data as described in 'c'.

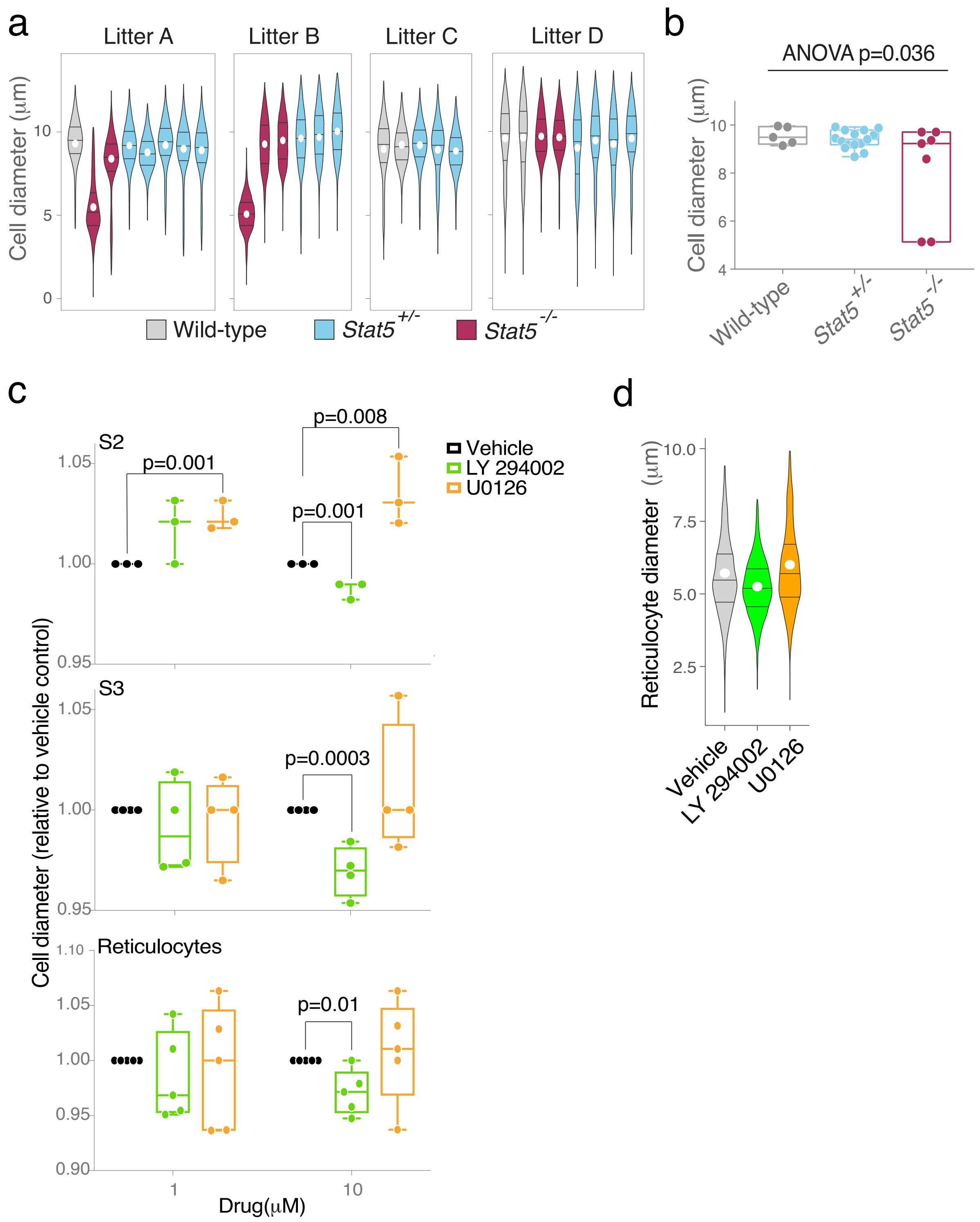


**Extended Data Figure 6 EpoR regulates cell size via multiple signaling pathways**

**a** Cell diameters of circulating red blood cells in E13.5 littermate embryos from matings of *Stat5*$^{+/-}$ x *Stat5*$^{+/-}$ mice. Violin lines mark the 25$^{th}$, 50$^{th}$ and 75$^{th}$ percentile with a white circle marking the mean.

**b** Median cell diameters of circulating red blood cells from panel 'a', by genotype. The box and whiskers mark the 25$^{th}$ to 75$^{th}$ percentiles and min to max values, respectively with the median indicated. p value is one-way ANOVA.

**c** Effect of inhibiting either PI3 kinase (with LY 294002) or MEK1/MEK2 (U0126) on the cell diameter of S2 and S3 erythroblasts and on reticulocytes. S0 cells were isolated from E13.5 wild-type fetal liver and differentiated *in vitro* in the presence of either vehicle (DMSO) or with inhibitors at the indicated concentrations. Diameters were measured at 48 h. Data are population medians from 2-5 experiments for each inhibitor concentration. Box and whiskers mark the 25$^{th}$ to 75$^{th}$ percentiles and min to max values respectively, with the median indicated. p values are for unpaired 2-tailed t-test.

**d** Violin plot of reticulocyte diameters for a representative experiment from 'c', inhibitor concentrations were each 10μM. Violin lines mark the 25$^{th}$, 50$^{th}$ and 75$^{th}$ percentile with a white circle marking the mean.

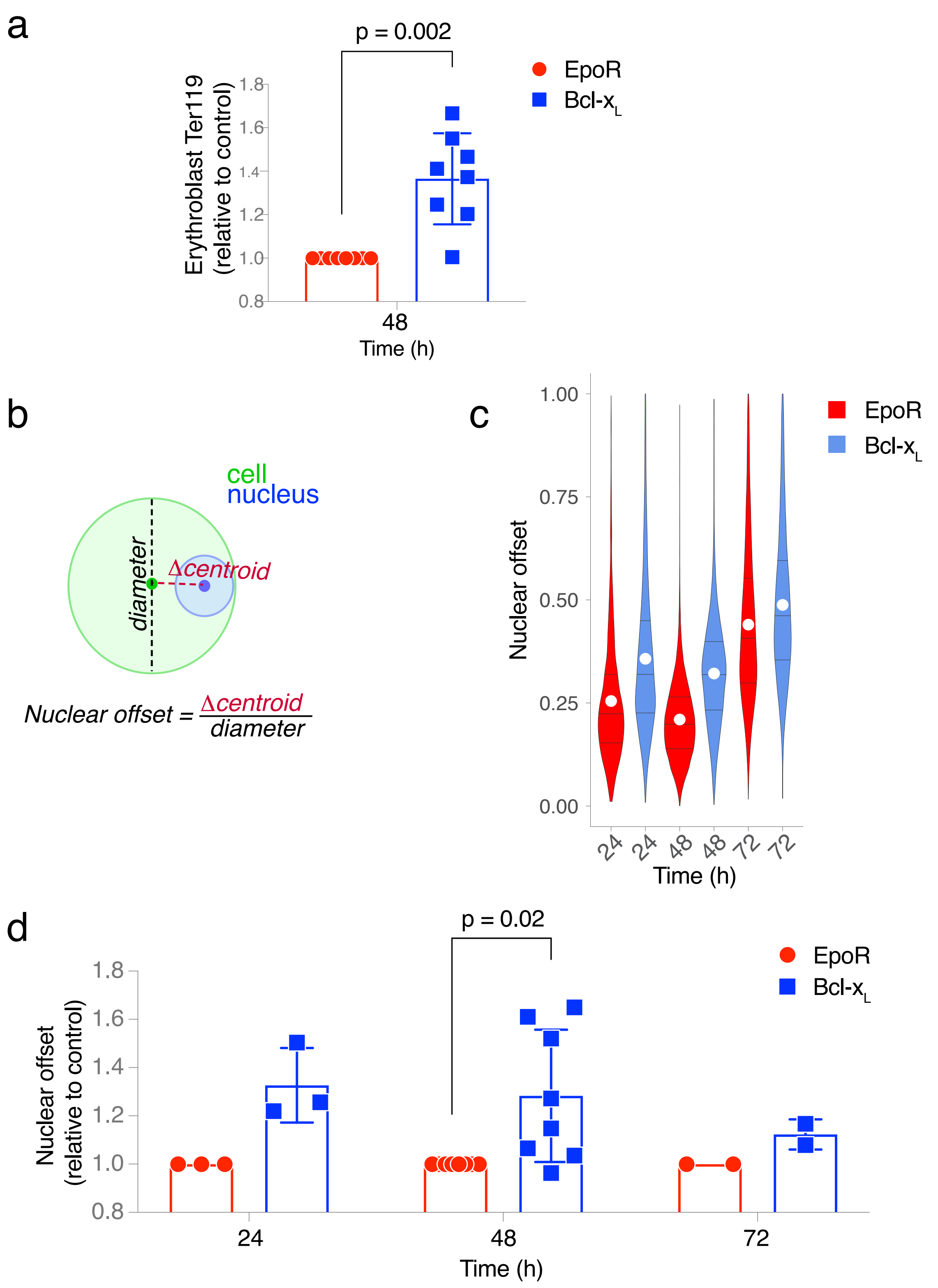


**Extended Data Figure 7** **EpoR signaling prolongs the ETD stage**

**a** Ter119 intensity in *Epor*$^{-/-}$ erythroblasts (Lin$^{-}$ hCD4$^{+}$) transduced with either Bcl-x$_L$ or EpoR, at 48 h. Data are mean Ter119 intensities from 9 independent experiments. Box and error bars are mean ± SD. p values are for paired ttest.

**b** Explanation of the nuclear offset measurement. The geometrical centers (=centroids) of the cell and the nucleus are indicated by green and blue solid circles. The distance between the centroid is the delta centroid (red dashed line, also known as the X,Y delta centroid). The nuclear offset is a dimensionless ratio of the delta centroid to the cell diameter.

**c** Representative nuclear offset measurements of *Epor*$^{-/-}$ erythroblasts (Lin$^{-}$ hCD4$^{+}$) transduced with either Bcl-x$_L$ or EpoR, at the indicated time points. Violin lines mark the 25$^{th}$, 50$^{th}$ and 75$^{th}$ percentile with a white circle marking the mean.

**d** Nuclear offset measurements of *Epor*$^{-/-}$ erythroblasts (Lin$^{-}$ hCD4$^{+}$) transduced with either Bcl-x$_L$ or EpoR. Data are medians pooled from 9 independent experiments. p values are for paired ttest.

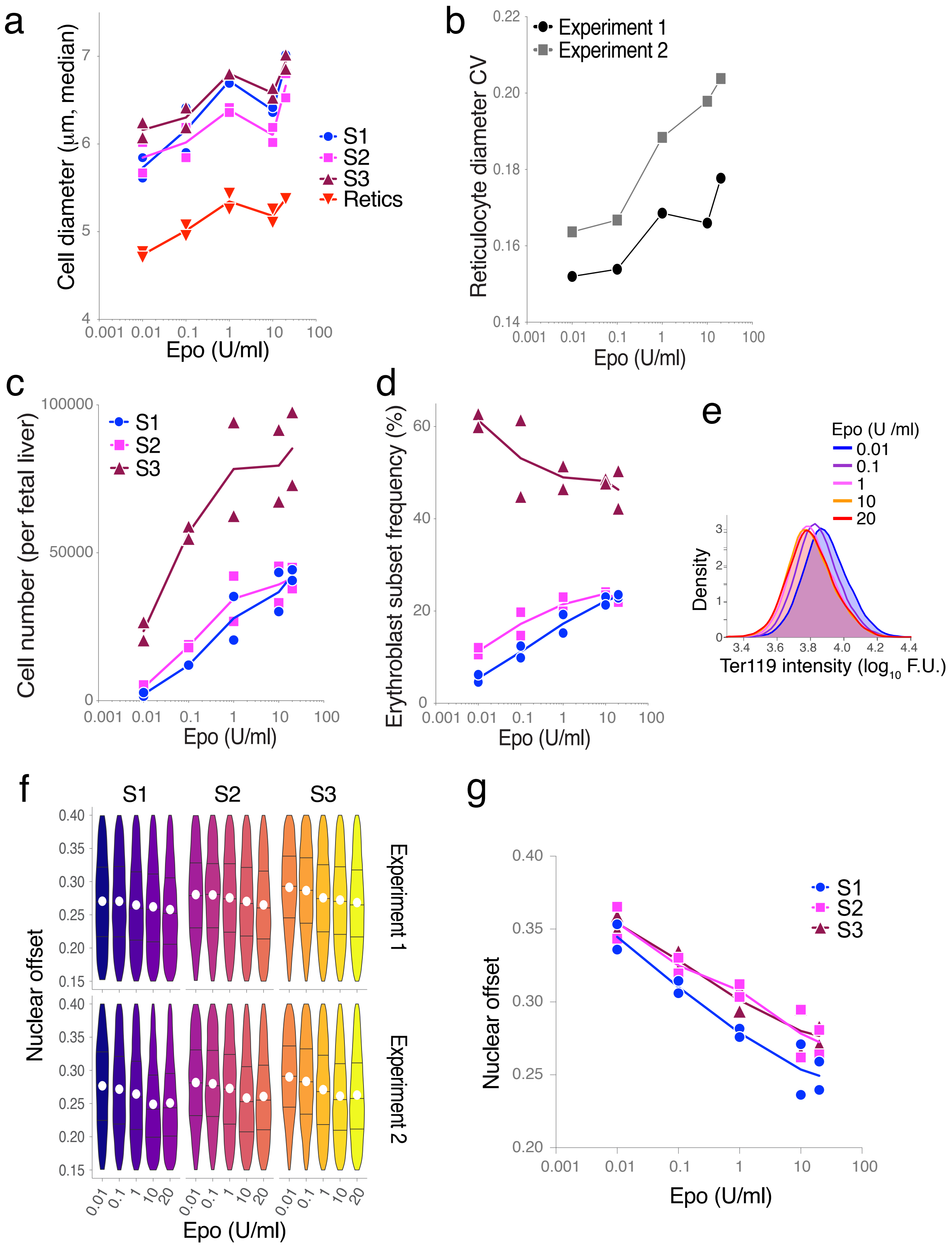


**Extended Data Figure 8 Multiple ETD parameters are sensitive to Epo concentration**

**a - g** Experiment as in **Figure 6a-b**. Wild-type fetal liver cells were enriched for CFU-e progenitors ('S0') and differentiated *in vitro*, in a range of Epo concentrations between 0.01 and 20 U/ ml. Cultures were analyzed at 48 h. Data are pooled from two independent experiments.

**a** Cell diameter (population medians) of erythroblast subsets S1 to S3 and of reticulocytes.

**b** Heterogeneity in reticulocyte diameter. CV, coefficient of variation.

**c, d** Viable cell number ('c') and cell frequency in the culture ('d') for each erythroblast subset.

**e** Ter119 intensity of all nucleated erythroblasts in the culture, for each of the indicated Epo concentrations at 48h.

**f, g** Nuclear offset at each Epo concentration and for each erythroblast subset. Violins ('e') are representative of two independent experiments. Data medians from both experiments are plotted in 'f'. Violin lines are the $25^{th}$, $50^{th}$ and $75^{th}$ percentile with a white circle marking the mean.

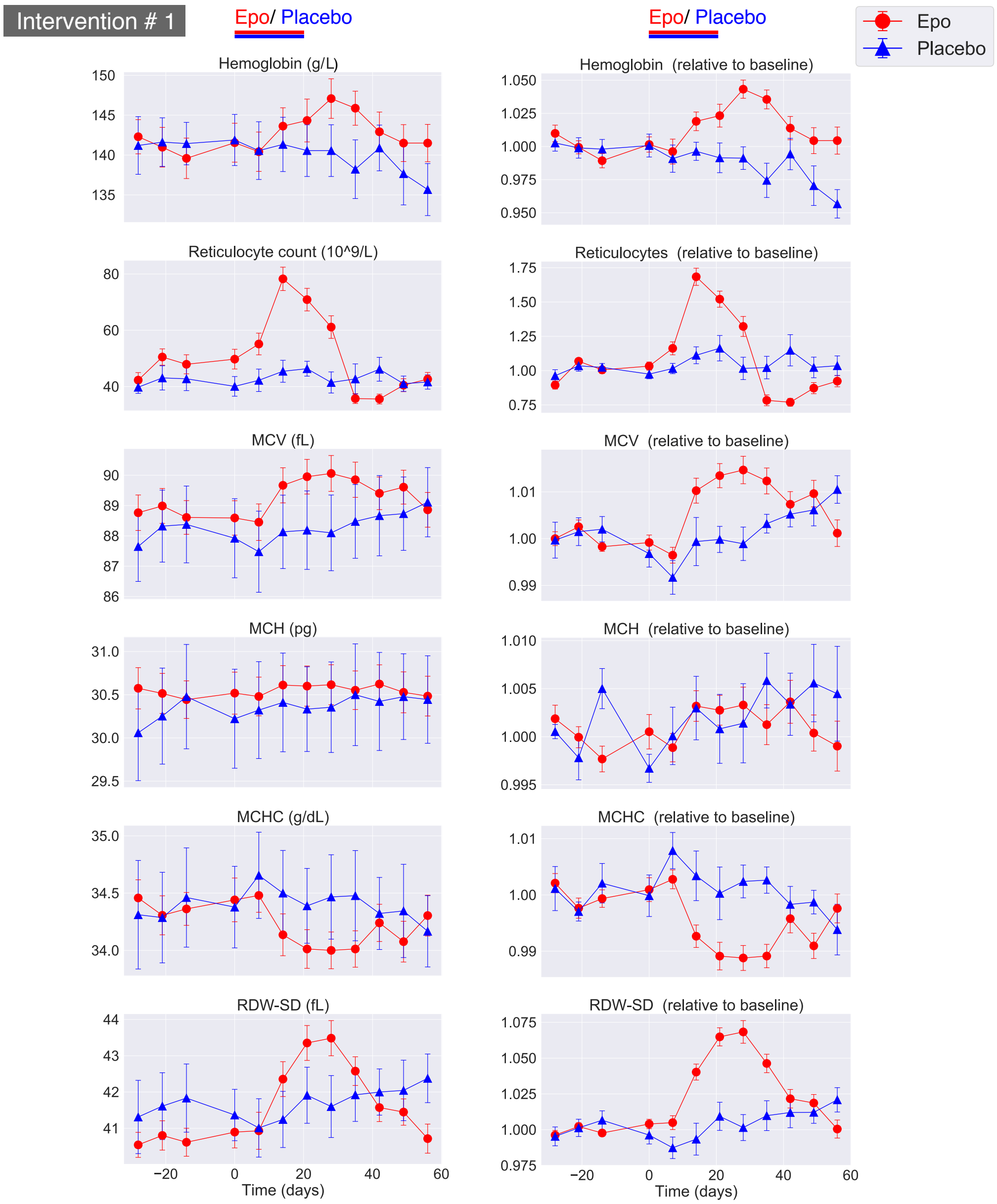


| Corrected p values | | | | | | |
|---|---|---|---|---|---|---|
| Day | Hemoglobin | Reticulocyte count | MCV | MCH | MCHC | RDW-SD |
| 7 | | | | | | |
| 14 | | 0.0001 | | | 0.0289 | 0.00017 |
| 21 | | 0.0005 | 0.0218 | | 0.0289 | <0.0001 |
| 28 | 0.003 | 0.0192 | 0.0097 | | 0.0069 | <0.0001 |
| 35 | 0.00079 | 0.0007 | | | 0.0069 | 0.0044 |
| 42 | | 0.00026 | | | | |
| 49 | | | | | | |
| 56 | 0.006 | | | | | |

significantly higher
significantly lower

**Extended Data Figure 9 Epo increases MCV and RDW independently of reticulocyte counts (University of Copenhagen, Denmark)**

Hematological parameters in intervention study 1. An initial 4 weekly venopuncture sampling period was followed by intravenous injections every 48 hours for 3 weeks, of either Epo (epoetin alpha, 20 IU·kg $bw^{-1}$) or placebo (the treatment period is indicated by red or blue horizontal bars). Venous blood samples were collected weekly during the treatment period and for five weeks following treatment. There were 25 participants in the Epo group (13 male and 12 female) and 9 participants in the placebo group (6 male, 3 female).

Hematological parameters are shown raw (left column) and also relative to baseline values for each participant (right column). Data are mean ± sem. The table shows p values for significant differences between Epo and placebo for each parameter and time point (linear mixed-effect model with Benjamini–Hochberg correction for multiple comparisons, see supplementary statistical analysis).

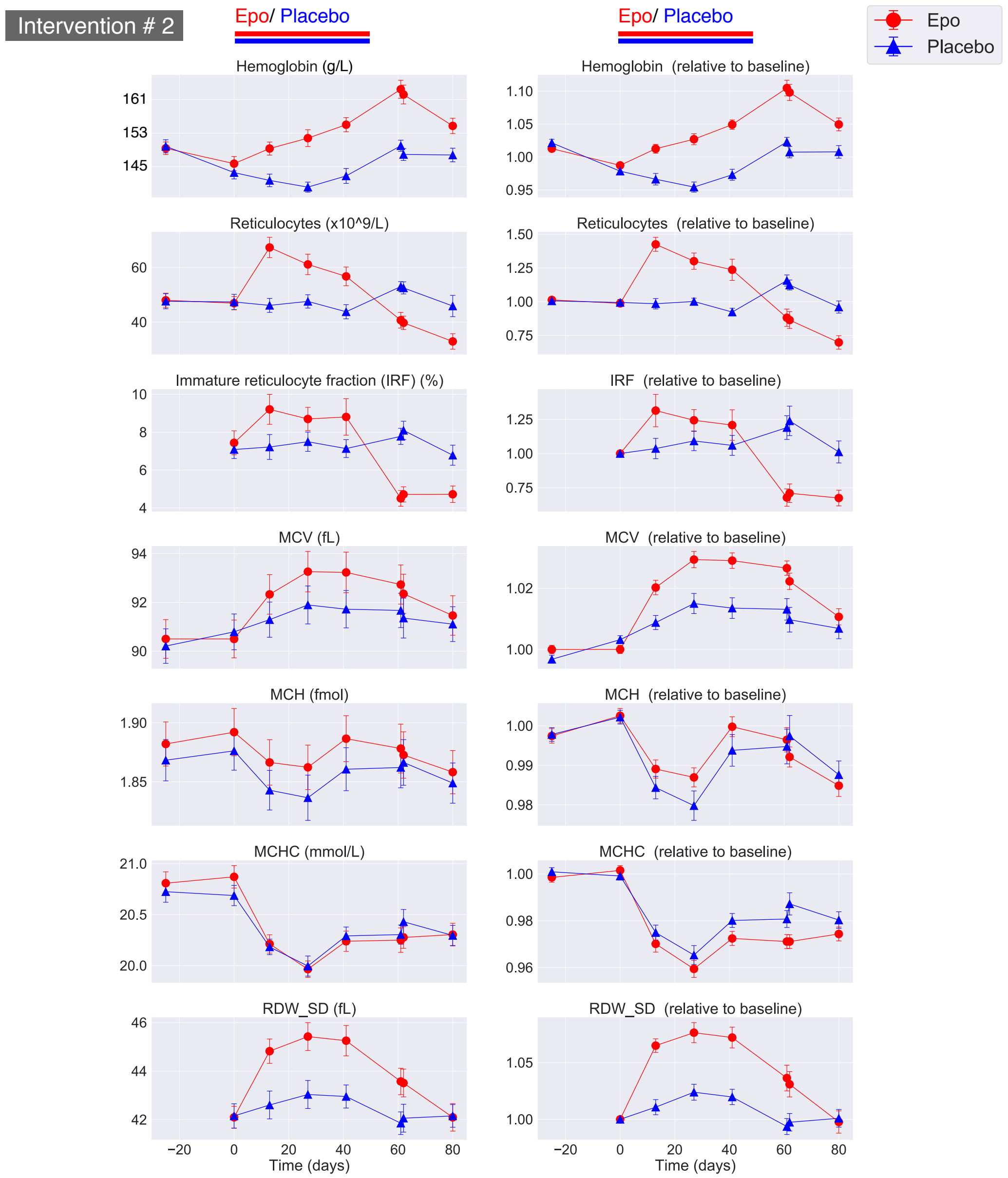


| Corrected p values | | | | | | | |
|---|---|---|---|---|---|---|---|
| Day | Hemoglobin | Reticulocyte count | IRF | MCV | MCH | MCHC | RDW-SD |
| 13 | 0.0002 | <0.0001 | | 0.0071 | | | <0.0001 |
| 27 | <0.0001 | 0.0001 | | 0.003 | | | <0.0001 |
| 41 | <0.0001 | 0.0002 | | 0.0019 | | | <0.0001 |
| 61 | <0.0001 | 0.0004 | 0.0002 | 0.0049 | | | 0.0004 |
| 62 | <0.0001 | 0.0004 | 0.0002 | 0.0071 | | 0.0153 | 0.0032 |
| 80 | 0.006 | 0.0002 | 0.0051 | | | | |

significantly higher
significantly lower

**Extended Data Figure 10 Epo increases MCV and RDW independently of reticulocyte counts (Centre for Human Drug Research, Leiden, Netherlands)**

Hematological parameters in intervention study 2. Two baseline venopuncture sampling measurements were followed by weekly dosing with Epo (24 male subjects) or placebo (24 male subjects) for 7 weeks. Epo dosing (epoetin β, 5000 to 10000 IU) was adjusted for each subject, to achieve an increase of 10 to 15% in hemoglobin over baseline. Follow-up continued for a month after cessation of treatment.

Hematological parameters are shown raw (left column) and also relative to baseline values for each participant (right column). Data are mean ± sem. The table shows p values for significant differences between Epo and placebo for each parameter and time point (linear mixed-effect model with Benjamini–Hochberg correction for multiple comparisons, see supplementary statistical analysis).

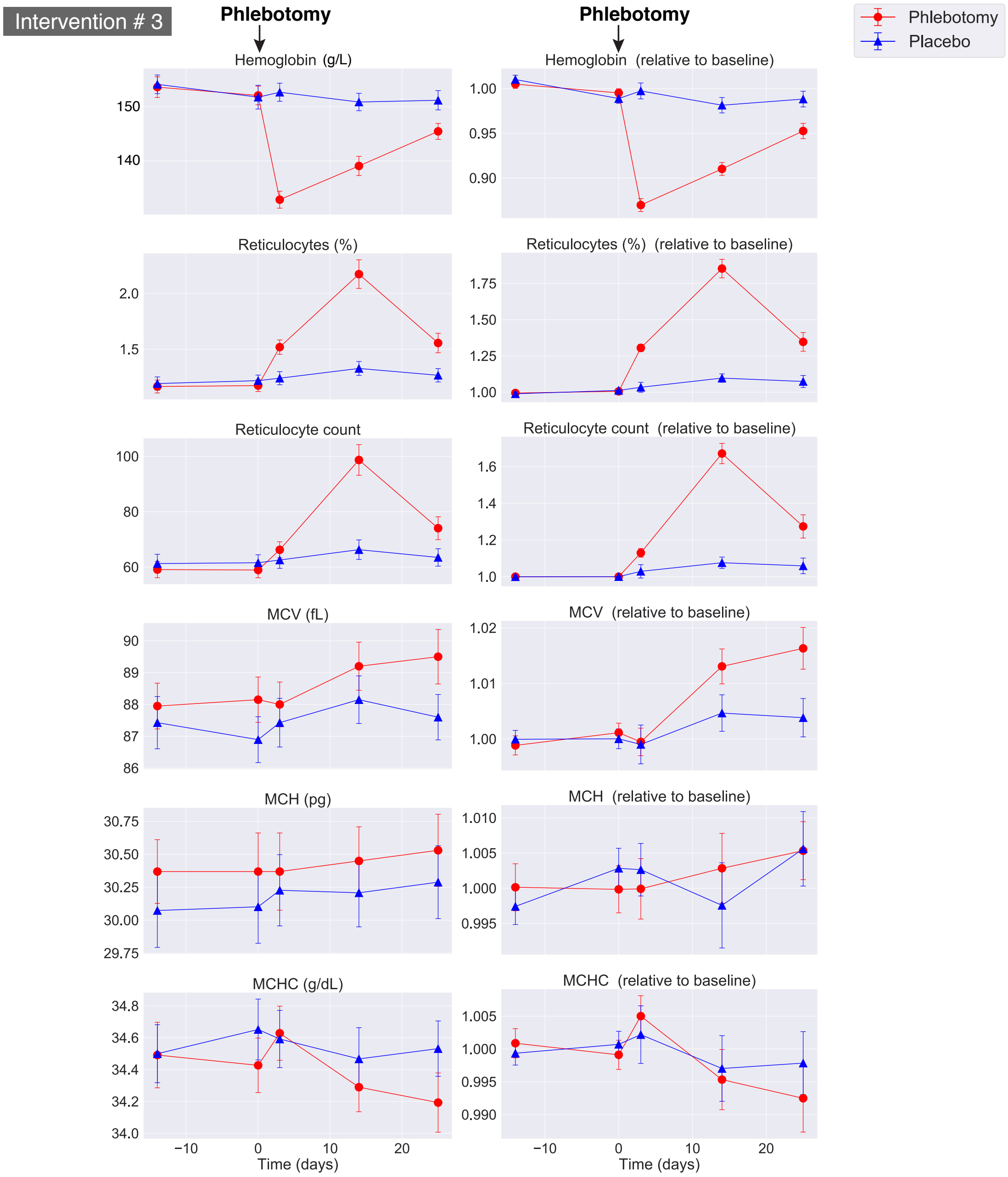


Corrected p values

| Day | Hemoglobin | Reticulocyte count | MCV | MCH | MCHC |
|---|---|---|---|---|---|
| 3 | <0.0001 | 0.03 | | | |
| 14 | <0.0001 | <0.0001 | | | |
| 25 | 0.0105 | 0.0047 | 0.0188 | | |

significantly higher
significantly lower

**Extended Data Figure 11 Phlebotomy results in increased MCV independently of reticulocyte counts (University of Copenhagen, Denmark)**

Hematological parameters in intervention study 3. This was a single blind, placebo-controlled cross-over study, with 21 participants. Two baseline venopuncture sampling measurements 4 days apart were made in the week preceding phlebotomy. At t=0, participants were either phlebotomized of two whole-blood units (900 ml), or sham-phlebotomized. Venous blood was sampled 3, 14 and 25 days later. At least 4 months elapsed before cross-over.

Hematological parameters are shown raw (left column) and also relative to baseline values for each participant (right column). Data are mean ± sem. The table shows p values for significant differences between Epo and placebo for each parameter and time point (linear mixed-effect model with Benjamini–Hochberg correction for multiple comparisons, see supplementary statistical analysis).

### Materials and Methods

#### Mice

*Stat5*$^{-/-}$ mice were obtained from Dr. Lothar Hennighausen (National Institute of Diabetes and Digestive and Kidney Diseases, Bethesda, MD). *Epor*$^{+/-}$ mice were obtained from the Lodish laboratory, Whitehead Institute for Biomedical Research, Cambridge, MA. Balb/C mice were obtained from the Charles River Laboratories, Wilmington, MA.

All experiments were conducted in accordance with animal protocol A-1586 approved by the University of Massachusetts Medical School Institutional Animal Care and Use Committee.

#### Isolation of erythroid progenitors

To isolate wildtype S0 cells, fetal liver cells were depleted of lineage -positive cells by labeling with biotin-conjugated CD71, Ter119, Gr1, Mac1 and CD41 antibodies followed by magnetic separation using either EasySep beads a (StemCell Technologies) or MojoSort™ Streptavidin Nanobeads (BioLegend) according to the manufacturers' instructions.

#### Flow cytometry

Fetal liver cells were analyzed on LSRII (BD Biosciences) cytometers using DIVA software (BD Biosciences). Dead cells were excluded using DAPI (Roche). FACS data was analyzed using FlowJo software (Tree Star Inc., CA).

Antibodies used:
PE Mouse Anti-Human CD4 (RPA-T4) (BD Biosciences)
PE/Cy7 Rat Anti-Mouse CD71 (RI7217) (BioLegend)
APC /Cyanine7 Rat Anti-Mouse Ter119 (Ter119) (BioLegend)
PE Rat Anti-Mouse Ter119 (Ter119) (BD Biosciences)
APC Rat Anti-Mouse Ter119 (Ter119) (BD Biosciences)
biotin Rat Anti-Mouse CD71 (C2) (BD Biosciences)
biotin Rat Anti-Mouse Ter119 (BD Biosciences)
biotin Rat Anti-Mouse Ly-6G and Ly-6C/Gr1 (RB6-8C5) (BD Biosciences)
biotin Rat Anti-Mouse CD11b/Mac1 (M1/70) (BD Biosciences)
biotin Rat Anti-Mouse CD41 (MWReg30) (Thermo Scientific)
FITC Rat Anti-Mouse Ly-6G and Ly-6C/Gr1 (RB6-8C5) (BD Biosciences)
FITC Rat Anti-Mouse CD11b/Mac1 (M1/70) (BD Biosciences)
FITC Rat Anti-Mouse CD41 (MWReg30) (BD Biosciences)
FITC Rat Anti-Mouse CD45R/B220 (RA3-6B2) (BD Biosciences)
FITC Hamster Anti-Mouse CD3e (145-2C11) (BD Biosciences)
PE Rat Anti-Mouse Ly-6G and Ly-6C/Gr1 (RB6-8C5) (BioLegend)
PE Rat Anti-Mouse CD11b/Mac1 (M1/70) (BioLegend)
PE Rat Anti-Mouse CD41 (MWReg30) (BD Biosciences)
PE Rat Anti-Mouse CD45R/B220 (RA3-6B2) (BD Biosciences)
PE Hamster Anti-Mouse CD3e (500A2) (BioLegend)

#### Imaging flow cytometry

Imaging flow cytometry was used to analyze cell fluorescence in conjunction with morphological parameters. It was performed on an Amnis Flowsight cytometer (Luminex Corporation, TX) using INSPIRE software v6.5 (Luminex Corporation, TX). Live nuclear diameter was measured using the cell permeable far-red fluorescent DNA dye, DRAQ5® (Cell Signaling). Amnis data was analyzed using IDEAS software v6.0 (Luminex Corporation, TX). New mask functions were generated to analyze bright field cell area (Definition: Object (M01, Ch01, Tight)) as well as Draq5 fluorescence nuclear area (Definition: Morphology (M11, Ch11)). Raw mean Draq5 pixel intensity feature was generated using Draq5 Morphology mask for nuclear area. Raw flow cytometric feature data were exported and analyzed in R programming language.

*Calibration of nuclear and cell diameters measured by imaging flow cytometry*
Imaging flow cytometry was performed on standardized bead sizes, 2.0µ, 3.4µ, 5.1µ, 7.4µ, 9.96µ, and 14.3µ (Spherotech Inc.). IDEAS bright field cell area mask (Definition: Object(M01, Ch01, Tight)) was fitted to the bead image acquisition. The data was analyzed using R, and within each bead group, values that lie greater or less than 3 standard deviations from the mean were removed (0.9% of events were removed with this threshold). To correct biases in the cell area values calculated by the Amnis software, a linear model (polynomial curve) was fit using the manufacturer bead sizes as a predictor for the Amnis calculated cell area (Stats, base R, degree = 2), with an $R^2$ value of 0.97. This model was then used to predict cell diameters from experimental cell areas.

*Analysis of imaging flow cytometry data*
Further analysis of exported imaging flow cytometry data was done using R Studio Version 1.2.1335, RStudio, Inc. Population distributions were log normalized. Population dataset were filtered by removing outliers that are 3 or more standard deviations from the mean.

*Fluorescence Quantile analysis (extended data Figure 4)*
Events whose cell areas were 3 standard deviations from the mean were removed. For CD4 and GFP intensities, quantiles were calculated across all samples using the quantile function (Stats, base R). To visualize the data, a density plot was drawn using ggplot2 (geom_density) [2] and colors chosen from viridis [3]. Each event was then categorized by which bin it fell into (for CD4 and GFP respectively). After this, each event had 2 associated values, which quantile of GFP and which quantile of CD4 that it belonged to. For each of the 3 samples (Bcl-$x_L^{GFP}$+$V^{hCD4}$, $V^{GFP}$+$EpoR^{hCD4}$ and Bcl-$x_L^{GFP}$+$EpoR^{hCD4}$), the mean cell diameter was then calculated within each of these composite bins (i.e. the mean cell diameter in Sample X, for GFP quantile Y and CD4 quantile Z). Next, the ratio of the mean cell diameters within each CD4/GFP bin were calculated between Bcl-$x_L^{GFP}$+$V^{hCD4}$ and Bcl-$x_L^{GFP}$+$EpoR^{hCD4}$ using $V^{GFP}$+$EpoR^{hCD4}$ as a reference. These data were plotted as a heatmap using ggplot2 and the geom_tile function. To show the distributions of cell diameters for all samples within each composite bin, example density plots were drawn using ggplot2 (geom_density). Example insets were colored by sample, and separate panels were drawn for each composite quantile bin.

*Nuclear offset*
The intensity weighted delta centroid XY feature was used to measure the distance between the centroid features of two images: CD71 fluorescence for the cell image and DRAQ5 fluorescence for the nucleus. To calculate the cell diameter, first the correlation between CD71 area feature and the bright-field-based area feature was obtained by plotting both values for each event. This allowed us to assign a bright-field area value to each event based on the CD71 area, and then use this value, in combination with the bead calibration curve (see '*Calibration of nuclear and cell diameters' above)*, to calculate cell diameter. Nuclear offset was then calculated by dividing the delta centroid by cell diameter.

*Identification of enucleated reticulocytes*

Cells were selected by gating on focused, single cell, live, lineage negative (Gr1, Mac1, CD41, B220, CD3e), hCD4 and GFP positive, and Ter119 positive events. The raw mean pixel intensity of Draq5 (nuclei) was plotted against the total Draq5 intensity (nuclei), giving two clearly distinct populations. We visually confirmed lack of Draq5 signal in the enucleated reticulocyte population.

**Cytospins**
Cells were spun onto coated Shandon™ Cytoslides (Thermo Scientific) using a Shandon™ Cytospin3 at 800 rpm for 5 minutes. The slides were dried, fixed and stained as described [4]. Cytospin preparations were examined using a Zeiss Axioskop 40 microscope using a SPOT Flex Camera (Diagnostic Instruments, Inc.) and imaged using SPOT v.5.6 software (SPOT Imaging).

**Cell Cycle Analysis**
Cell cycle status and S phase speed were analyzed using BrdU incorporation as described [5]. BrdU incorporation was measured by biotin-conjugated anti-BrdU (MOBU-1, BioLegend) followed by a secondary stain with Brilliant Violet 421™ Streptavidin (BioLegend). DNA content was measured by 7AAD (BD Biosciences).

**Retroviral Transduction and *in vitro* differentiation of fetal liver cells**
Epor, Bcl-$x_L$ and Tfrc were subcloned into MSCV-IRES-hCD4 retroviral vector. Bcl-$x_L$ and Tfrc were also subcloned into MSIG-IRES-GFP retroviral vector (MSIG 1.1 SK). High-titer viral supernatants were prepared by co-transfecting the pCL-Eco packaging vector and desired plasmid into Phoenix cells using Lipofectamine 2000 transfection reagent (ThermoFisher Scientific). High-titer virus was collected in 'erythroid medium': IMDM (L-glutamine, 25mM HEPES) (Gibco), 20% fetal calf serum, 1% penicillin/streptomycin, $10^{-4}$ M β-Mercaptoethanol.

Retroviral transduction was done by spin infection of Epor-/- or *Epor-/- Hri-/-* fetal liver cells at 2,000 rpm, 30°C for 1 hour on 50 $\mu$g/ml fibronectin (GIBCO) coated dishes in 4 $\mu$g/ml polybrene (Sigma), supplemented in some experiments with 0.5U/ml Epo (Amgen). Transduced cells were incubated for 15 hours with 100 ng/ml SCF and 10 ng/ml IL3 (Peprotech). Cells were then transferred to differentiation medium: IMDM (L-glutamine, 25mM HEPES) (Gibco), 20% fetal calf serum, 1% penicillin/streptomycin, $10^{-4}$ M β-Mercaptoethanol and 0.5U/ml Epo (Amgen) for the indicated times. In the case of experiments that include *Epor$^{-/-}$ Hri$^{-/-}$*, the media was also supplemented with 1mg/ml iron-saturated human transferrin (Sigma). Where indicated, liquid cultures were also supplemented with Fe-loaded salicylaldehyde isonicotinoyl hydrazone (Fe-SIH, 10$\mu$M, a lipophilic iron chelator, a gift from the late Dr. Prem Ponka (McGill University, Montréal, Québec, Canada), with 0.7$\mu$M deoxyribonucleosides (3'-Deoxythymidine, 2'-Deoxyguanosine monohydrate, 2'-Deoxyadenosine monohydrate, 2'-Deoxycytidine, Sigma),

***In vitro* differentiation of fetal liver cells with PI3K and MEK1/MEK2 inhibitors**
Isolated wildtype S0 cells were cultured in differentiation media (Epo 0.5U/ml) with 1$\mu$M or 10$\mu$M PI3K inhibitor, LY 294002 (EMD Millipore) or MEK1/MEK2 inhibitor, U0126 (EMD Millipore). Inhibitors were replenished every 24hrs.

**Colony-formation assays in methylcellulose**
Retroviral transduction was done by spin infection of *Epor$^{-/-}$* fetal liver cells as described above. From each transduced sample (4 hours post infection), 200,000 cells were mixed with 1 ml MethoCult (M3234, STEMCELL Technologies) supplemented with 2U/ml Epo (Amgen).

Erythroid (CFU-e) were scored from duplicate plates on day 3 of culture. Expression of hemoglobin in erythroid colonies was confirmed by staining with diaminobenzidine (Sigma) in situ before scoring. Colony cell area was measured using ImageJ version: 2.0.0-r-54/1.51h.

**Quantitative RT-PCR Assay**
Total RNA was isolated from *in vitro* cultured fetal liver cells using the AllPrep DNA/RNA Micro Kit (Qiagen) and quantified by Quant-iT RiboGreen RNA reagent kit (Thermo Scientific) on the 3300 NanoDrop Fluorospectrometer. Reverse transcription was done using the SuperScript III first-strand synthesis system (Invitrogen) with random hexamer primers. Quantitative PCR was performed using the ABI 7300 sequence detection system with TaqMan reagents and TaqMan MGB probes (Applied Biosystems). Each reaction was carried out on a dilution series of the template cDNA to ensure linearity of signal.

TagMan MGB probes used: β-actin (Mm02619580_g1), PU.1 (Mm00488140_m1), GATA-1 (Mm01352636_m1), GATA-2 (Mm00492300_m1), Alas2 (Mm01260713_m1), Band3 (Mm01245920_g1), β-globin (Mm01611268_g1), p21 (Mm00432448_m1), p27 (Mm00438168_m1), p57 (Mm01272135_g1), p16ink4a (Mm01257348_m1), p15ink4b (Mm00483241_m1), p18ink4c (Mm00483243_m1), p19ink4d (Mm00486943_m1), CCND1 (Mm00432359_m1), CCND2 (Mm00438071_m1), CCND3 (Mm01612362_m1), CCNE1 (Mm00432367_m1), CCNE2 (Mm00438077_m1), CCNA2 (Mm00438063_m1), CCNA1 (Mm00432337_m1), E2F2 (Mm00624964_m1), E2F4 (Mm00514160_m1), Tfrc (Mm00441950_m1), Bcl-xL (Mm00437783_m1), DNMT1 (Dnmt100599784), Ifitm1 (Mm01279023_m1), Ifitm3 (Mm00847057_s1), Tal1 (Mm00441665_m1), NFE2 (Mm00801891_m1), LMO2 (Mm00493153_m1), cdk6 (Mm00438163_m1), cdk6 (Mm01311342_m1), cdk4 (Mm00726334_s1), cdk2 (Mm00443947_m1), cdc25a (Mm00483162_m1), cdc25b (Mm00499136_m1), cdc25c (Mm00486880_m1), Klf1(Mm00516096_m1).

***in vivo* Epo stimulation**
Epo (Epoetin alfa; Amgen) was injected subcutaneously in a total volume of 300 $\mu$ L in sterile isotonic saline, at the indicated doses and frequencies.

**Human Intervention Studies**

**Human intervention studies 1 and 3** were performed at the University of Copenhagen. In intervention study 1, subjects received recombinant human erythropoietin (rhEPO). In intervention study 3, participants were subjected to phlebotomy. Thirty-four healthy non-smoking Caucasian males (n=19) and females (n=15) (age 25±3 years, height 179±10 cm and weight 70±10 kg) participated in the erythropoietin treatment intervention. Another twenty-one healthy non-smoking Caucasian male subjects (age 29±6 years, 184±7 cm and 77±8 kg) participated in the phlebotomy intervention. No participant had donated blood for at least three months prior to the start of the study or been exposed to high altitude (> 1000 m) for at least two months.
The local ethics committee of Copenhagen, Denmark (H-2-2014-109 & H-17036662) approved the studies, which was performed in accordance with the Declaration of Helsinki. All participants were informed both orally and in writing of potential risks and discomforts associated with participation before written consent was obtained.

<u>*Experimental design*</u>

*rhEPO treatment:* The study used a randomized single-blinded placebo-controlled design. After weekly baseline collection of venous blood for four weeks, the participants received eleven intravenous injections of 20 IU·kg $bw^{-1}$ epoetin alpha (Eprex, Janssen, Birkerød, Denmark) (rhEPO group, 25 participants; 13 male and 12 female) or saline (placebo group, 9 participants; 6 male and 3 female) every second day. Venous blood samples were collected weekly during the treatment and for five weeks following treatment.

*Phlebotomy:* The intervention applied a randomized single-blinded placebo-controlled crossover design. The week before phlebotomy, two baseline venous blood samples were collected with 4 days apart. Next, the participants were phlebotomized of two whole-blood units, corresponding to 900 mL or sham-phlebotomized followed by venous blood collection 3, 14 and 25 days later. A recovery period of >4 months was applied before the participants crossed over and repeated the experiment.

*Blood sample analysis*

All venous blood samples were collected in 2 mL EDTA-anticoagulated vacutainers (Becton Dickinson, New Jersey, USA) after at least 10 min of rest in a seated position and with less than 30 s use of tourniquet. In the rhEPO trial, samples were immediately analyzed for a complete blood count using a Sysmex XN-450 (Sysmex, Kobe, Japan) including mean cell volume, hemoglobin concentration, reticulocyte count, reticulocyte percentage and red cell distribution width. In the phlebotomy trial, samples were stored at 4 °C and analyzed within 2 h of collection for mean cell volume, hemoglobin concentration, reticulocyte count and reticulocyte percentage using a Sysmex XE-2100 (Sysmex, Kobe, Japan).

**Human intervention study 2** was performed at the Centre for Human Drug Research, Leiden, Netherlands. This study was reported elsewhere [6], but reporting did not include MCV and RDW information. Briefly, non-professional well trained male cyclists ages 28 to 50 were randomly assigned to placebo or recombinant human Epo (epoetin β) groups. Baseline measurements were followed by weekly dosing with Epo (24 subjects) or placebo (24 subjects) for 7 weeks. Epo dosing (5000 to 10000 IU) was adjusted for each subject, to achieve an increase of 10 to 15% in hemoglobin over baseline. Follow-up continued for a month after cessation of treatment.

**Human intervention studies- statistical analysis (see supplementary statistical analysis)**

We computed baseline-corrected values at each post-baseline time point for each subject by subtracting the corresponding subject-level mean baseline measurement, which were used to fit linear mixed-effect models using the nlme package [7]. For intervention studies 1 and 2, the model includes subject as random effect, treatment, time, and the interaction of treatment by time as fixed effects. To test whether Epo treatment and placebo differ significantly at each post-baseline time point, a set of pre-defined contrasts were performed using the multcomp package [8] followed by multiplicity adjustment using *Benjamini–Hochberg* procedure [9]. For intervention study 3, each post-baseline time point was analyzed separately with the model that includes subject as random effect, treatment, period, and sequence of treatments as fixed effects.
