## Supplementary material for "The Erythropoietin Receptor Stimulates Rapid Cycling and Formation of Larger Red Cells During Mouse and Human Erythropoiesis": Statistical analysis for human intervention studies #1-3 (from another version of this preprint: version 1, 2020-11-30)

Code ▾

### Data analysis of Epo intervention study # 1, Hidalgo et al.

**By Lihua Julie Zhu**

Citation:


1. Lindstrom, M.J. and Bates, D.M. (1988) "Newton-Raphson and EM Algorithms for Linear Mixed-Effects Models for Repeated-Measures Data", Journal of the American Statistical Association, 83, 1014–1022.
2. Hothorn T, Bretz F, Westfall P (2008). "Simultaneous Inference in General Parametric Models." Biometrical Journal, 50(3), 346–363.
3. Benjamini, Y., and Hochberg, Y. (1995). Controlling the false discovery rate: a practical and powerful approach to multiple testing. Journal of the Royal Statistical Society Series B, 57, 289–300.http://www.jstor.org/stable/2346101.


Data summary

```
[1] "Thu Mar 19 18:32:15 2020"
[1] 407  11
```

| | Treatment <fctr> | Subject <int> | time_point <fctr> | HbgdL <dbl> | MCHpg <dbl> | MCHCgdL <dbl> | MCVfl <dbl> | RDWCV <dbl> | RDWSDfl <dbl> |
|---|---|---|---|---|---|---|---|---|---|
| 1 | EPO | 1 | Baseline 1 | 12.73875 | 31.92750 | 33.70125 | 94.7 | 12.0 | 42.3 |
| 2 | EPO | 1 | Baseline 2 | 12.25500 | 32.18550 | 34.50750 | 93.2 | 12.2 | 42.2 |
| 3 | EPO | 1 | Baseline 3 | 12.73875 | 32.04037 | 33.86250 | 94.6 | 12.1 | 42.4 |
| 4 | EPO | 1 | Baseline 4 | 12.80000 | 31.50000 | 33.10000 | 95.0 | 12.0 | 42.4 |
| 5 | EPO | 1 | EPO wk1 | 13.10000 | 32.00000 | 34.10000 | 94.1 | 12.2 | 42.1 |
| 6 | EPO | 1 | EPO wk2 | 13.00000 | 32.40000 | 34.00000 | 95.5 | 12.3 | 43.7 |

6 rows | 1-10 of 11 columns

```
Correlation matrix among different measurements
                  HbgdL       MCHpg      MCHCgdL       MCVfl       RDWCV     RDWSDfl    Retics
HbgdL       1.000000000  0.10296731 -0.004754063  0.12475968 -0.06211799  0.03608573 0.0754058
MCHpg       0.102967313  1.00000000  0.555489534  0.77843006 -0.47903236  0.08661093 0.2558222
MCHCgdL    -0.004754063  0.55548953  1.000000000 -0.08842794 -0.52550521 -0.54151516 0.1870510
MCVfl       0.124759678  0.77843006 -0.088427942  1.00000000 -0.17906623  0.51124013 0.1649677
RDWCV      -0.062117994 -0.47903236 -0.525505208 -0.17906623  1.00000000  0.73977857 0.1235085
RDWSDfl     0.036085730  0.08661093 -0.541515158  0.51124013  0.73977857  1.00000000 0.1748552
Retics      0.075405804  0.25582219  0.187050951  0.16496774  0.12350851  0.17485524 1.0000000
Retics109L  0.091634764  0.19595026  0.224058174  0.06575297  0.11089097  0.09971701 0.9749060
           Retics109L
HbgdL      0.09163476
MCHpg      0.19595026
MCHCgdL    0.22405817
MCVfl      0.06575297
RDWCV      0.11089097
RDWSDfl    0.09971701
Retics     0.97490601
Retics109L 1.00000000
```

Compute mean of 4 baseline points for each subject

| | Treatment <fctr> | Subject <int> | time_point <fctr> | HbgdL <dbl> | MCHpg <dbl> | MCHCgdL <dbl> | MCVfl <dbl> | RDWCV <dbl> | RDWSDfl <dbl> |
|---|---|---|---|---|---|---|---|---|---|
| 1 | EPO | 1 | Baseline 1 | 12.73875 | 31.92750 | 33.70125 | 94.7 | 12.0 | 42.3 |
| 2 | EPO | 1 | Baseline 2 | 12.25500 | 32.18550 | 34.50750 | 93.2 | 12.2 | 42.2 |
| 3 | EPO | 1 | Baseline 3 | 12.73875 | 32.04037 | 33.86250 | 94.6 | 12.1 | 42.4 |
| 4 | EPO | 1 | Baseline 4 | 12.80000 | 31.50000 | 33.10000 | 95.0 | 12.0 | 42.4 |
| 5 | EPO | 1 | EPO wk1 | 13.10000 | 32.00000 | 34.10000 | 94.1 | 12.2 | 42.1 |
| 6 | EPO | 1 | EPO wk2 | 13.00000 | 32.40000 | 34.00000 | 95.5 | 12.3 | 43.7 |

6 rows | 1-10 of 19 columns

```
[1] 272  19
[1] EPO wk1  EPO wk2  EPO wk3  EPO wk4  post wk1 post wk2 post wk3 post wk4
12 Levels: Baseline 1 Baseline 2 Baseline 3 Baseline 4 EPO wk1 EPO wk2 EPO wk3 ... post wk4
```

| | Treatment <fctr> | Subject <int> | time_point <fctr> | HbgdL <dbl> | MCHpg <dbl> | MCHCgdL <dbl> | MCVfl <dbl> | RDW... <dbl> | RDWSDfl <dbl> |
|---|---|---|---|---|---|---|---|---|---|
| 5 | EPO | 1 | EPO wk1 | 0.466875 | 0.08665625 | 0.3071875 | -0.275 | 0.125 | -0.225 |
| 6 | EPO | 1 | EPO wk2 | 0.366875 | 0.48665625 | 0.2071875 | 1.125 | 0.225 | 1.375 |
| 7 | EPO | 1 | EPO wk3 | 0.266875 | 0.58665625 | 0.5071875 | 0.125 | 0.325 | 1.775 |
| 8 | EPO | 1 | EPO wk4 | 1.166875 | 0.38665625 | 0.0071875 | 1.325 | 0.725 | 2.575 |
| 9 | EPO | 1 | post wk1 | 1.466875 | 0.08665625 | 0.0071875 | 0.425 | 0.125 | 0.875 |
| 10 | EPO | 1 | post wk2 | 0.366875 | -0.11334375 | -0.1928125 | 0.325 | 0.025 | 0.175 |

6 rows | 1-10 of 19 columns

```
########Number of Subjects for each treatment ######
    EPO placebo
     25       9
```

```
##############################################################



##############################################################

########### Repeated measures analysis results for  HbgdL ############

##############################################################
```

| | numDF <int> | denDF <dbl> | F-value <chr> | p-value <chr> |
|---|---|---|---|---|
| (Intercept) | 1 | 217 | 3.475773 | 0.0636 |
| Treatment | 1 | 31 | 12.892589 | 0.0011 |
| time_point | 7 | 217 | 5.433239 | <.0001 |
| Treatment:time_point | 7 | 217 | 2.009929 | 0.0551 |

4 rows

```
############ Compare Epo with Palcebo at each time point using contrasts ###########


##############################################################

######################## Raw p-values #########################


     Simultaneous Tests for General Linear Hypotheses

Multiple Comparisons of Means: User-defined Contrasts


Fit: lme.formula(fixed = thisY ~ -1 + EpoTime, data = y, random = ~1 |
    Subject)

Linear Hypotheses:
                          Estimate Std. Error z value Pr(>|z|)
EpoVsPlacebo.EpoWk1 == 0   0.06244    0.21614   0.289 0.772650
EpoVsPlacebo.EpoWk2 == 0   0.30550    0.21614   1.413 0.157519
EpoVsPlacebo.EpoWk3 == 0   0.45411    0.21614   2.101 0.035637 *
EpoVsPlacebo.EpoWk4 == 0   0.72911    0.21614   3.373 0.000742 ***
EpoVsPlacebo.postWk1 == 0  0.84161    0.21614   3.894 9.86e-05 ***
EpoVsPlacebo.postWk2 == 0  0.27911    0.21614   1.291 0.196576
EpoVsPlacebo.postWk3 == 0  0.45967    0.21614   2.127 0.033441 *
EpoVsPlacebo.postWk4 == 0  0.65967    0.21614   3.052 0.002272 **
---
Signif. codes:  0 ‘***’ 0.001 ‘**’ 0.01 ‘*’ 0.05 ‘.’ 0.1 ‘ ’ 1
(Adjusted p values reported -- none method)

#########################################################


############## BH adjusted p-values ####################
Benjamini, Y., and Hochberg, Y. (1995). Controlling the false discovery rate: a practical and powerful approach t
o multiple testing. Journal of the Royal Statistical Society Series B, 57, 289–300.http://www.jstor.org/stable/23
46101.
##############################################################

     Simultaneous Tests for General Linear Hypotheses

Multiple Comparisons of Means: User-defined Contrasts


Fit: lme.formula(fixed = thisY ~ -1 + EpoTime, data = y, random = ~1 |
    Subject)

Linear Hypotheses:
                          Estimate Std. Error z value Pr(>|z|)
EpoVsPlacebo.EpoWk1 == 0   0.06244    0.21614   0.289 0.772650
EpoVsPlacebo.EpoWk2 == 0   0.30550    0.21614   1.413 0.210026
EpoVsPlacebo.EpoWk3 == 0   0.45411    0.21614   2.101 0.057019 .
EpoVsPlacebo.EpoWk4 == 0   0.72911    0.21614   3.373 0.002970 **
EpoVsPlacebo.postWk1 == 0  0.84161    0.21614   3.894 0.000789 ***
EpoVsPlacebo.postWk2 == 0  0.27911    0.21614   1.291 0.224658
EpoVsPlacebo.postWk3 == 0  0.45967    0.21614   2.127 0.057019 .
EpoVsPlacebo.postWk4 == 0  0.65967    0.21614   3.052 0.006060 **
---
Signif. codes:  0 ‘***’ 0.001 ‘**’ 0.01 ‘*’ 0.05 ‘.’ 0.1 ‘ ’ 1
(Adjusted p values reported -- BH method)


#############################################################
```

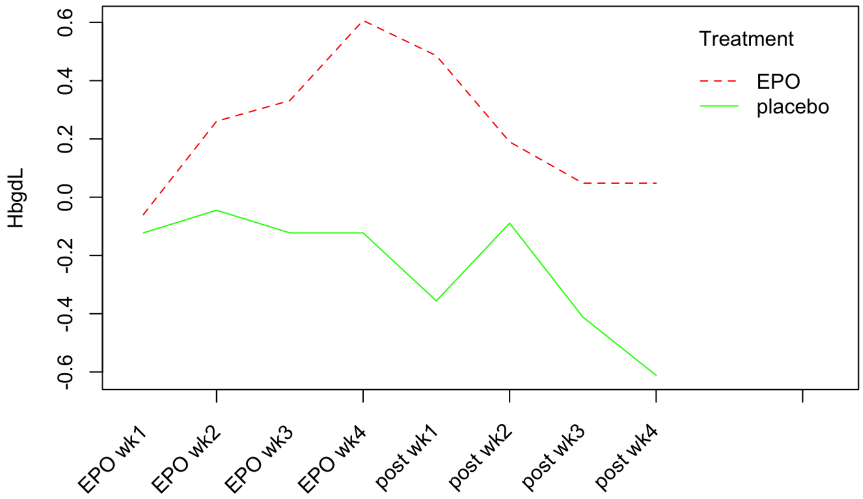


```
################################################################
########### Repeated measures analysis results for  MCHpg ############
#################################################################
```

| | numDF <int> | denDF <dbl> | F-value <chr> | p-value <chr> |
|---|---|---|---|---|
| (Intercept) | 1 | 224 | 2.2165316 | 0.1379 |
| Treatment | 1 | 32 | 0.2185137 | 0.6433 |
| time_point | 7 | 224 | 1.3468076 | 0.2294 |
| Treatment:time_point | 7 | 224 | 1.1691200 | 0.3215 |

4 rows

```
############ Compare Epo with Palcebo at each time point using contrasts ##########


###############################################################

####################### Raw p-values #######################


     Simultaneous Tests for General Linear Hypotheses

Multiple Comparisons of Means: User-defined Contrasts


Fit: lme.formula(fixed = thisY ~ -1 + EpoTime, data = y, random = ~1 |
    Subject)

Linear Hypotheses:
                           Estimate Std. Error z value Pr(>|z|)
EpoVsPlacebo.EpoWk1 == 0  -0.036286   0.119087  -0.305    0.761
EpoVsPlacebo.EpoWk2 == 0   0.006825   0.119087   0.057    0.954
EpoVsPlacebo.EpoWk3 == 0   0.072603   0.119087   0.610    0.542
EpoVsPlacebo.EpoWk4 == 0   0.066381   0.119087   0.557    0.577
EpoVsPlacebo.postWk1 == 0 -0.142064   0.119087  -1.193    0.233
EpoVsPlacebo.postWk2 == 0  0.007714   0.119087   0.065    0.948
EpoVsPlacebo.postWk3 == 0 -0.143842   0.119087  -1.208    0.227
EpoVsPlacebo.postWk4 == 0 -0.154508   0.119087  -1.297    0.194
(Adjusted p values reported -- none method)

##########################################################


############## BH adjusted p-values ####################
Benjamini, Y., and Hochberg, Y. (1995). Controlling the false discovery rate: a practical and powerful approach t
o multiple testing. Journal of the Royal Statistical Society Series B, 57, 289–300.http://www.jstor.org/stable/23
46101.
##############################################################

     Simultaneous Tests for General Linear Hypotheses

Multiple Comparisons of Means: User-defined Contrasts


Fit: lme.formula(fixed = thisY ~ -1 + EpoTime, data = y, random = ~1 |
    Subject)

Linear Hypotheses:
                           Estimate Std. Error z value Pr(>|z|)
EpoVsPlacebo.EpoWk1 == 0  -0.036286   0.119087  -0.305    0.954
EpoVsPlacebo.EpoWk2 == 0   0.006825   0.119087   0.057    0.954
EpoVsPlacebo.EpoWk3 == 0   0.072603   0.119087   0.610    0.924
EpoVsPlacebo.EpoWk4 == 0   0.066381   0.119087   0.557    0.924
EpoVsPlacebo.postWk1 == 0 -0.142064   0.119087  -1.193    0.621
EpoVsPlacebo.postWk2 == 0  0.007714   0.119087   0.065    0.954
EpoVsPlacebo.postWk3 == 0 -0.143842   0.119087  -1.208    0.621
EpoVsPlacebo.postWk4 == 0 -0.154508   0.119087  -1.297    0.621
(Adjusted p values reported -- BH method)


##############################################################
```

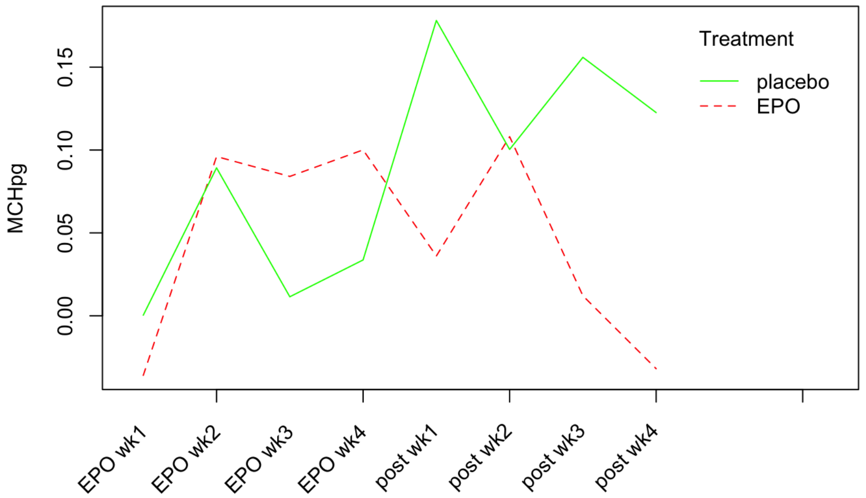


```
##############################################################

########## Repeated measures analysis results for  MCHCgdL ###########

###############################################################
```

| | numDF <int> | denDF <dbl> | F-value <chr> | p-value <chr> |
|---|---|---|---|---|
| (Intercept) | 1 | 224 | 13.550041 | 0.0003 |
| Treatment | 1 | 32 | 6.695523 | 0.0144 |
| time_point | 7 | 224 | 6.931815 | <.0001 |

| | numDF | denDF | F-value | p-value |
| --- | --- | --- | --- | --- |
| | <int> | <dbl> | <chr> | <chr> |
| Treatment:time_point | 7 | 224 | 3.145059 | 0.0034 |

4 rows

```
########### Compare Epo with Palcebo at each time point using contrasts ##########


###############################################################

####################### Raw p-values #######################


     Simultaneous Tests for General Linear Hypotheses

Multiple Comparisons of Means: User-defined Contrasts


Fit: lme.formula(fixed = thisY ~ -1 + EpoTime, data = y, random = ~1 |
    Subject)

Linear Hypotheses:
                          Estimate Std. Error z value Pr(>|z|)
EpoVsPlacebo.EpoWk1 == 0   -0.1742     0.1483  -1.175  0.24004
EpoVsPlacebo.EpoWk2 == 0   -0.3627     0.1483  -2.446  0.01446 *
EpoVsPlacebo.EpoWk3 == 0   -0.3756     0.1483  -2.533  0.01132 *
EpoVsPlacebo.EpoWk4 == 0   -0.4653     0.1483  -3.138  0.00170 **
EpoVsPlacebo.postWk1 == 0  -0.4644     0.1483  -3.132  0.00174 **
EpoVsPlacebo.postWk2 == 0  -0.0809     0.1483  -0.546  0.58540
EpoVsPlacebo.postWk3 == 0  -0.2671     0.1483  -1.801  0.07166 .
EpoVsPlacebo.postWk4 == 0   0.1387     0.1483   0.935  0.34978
---
Signif. codes:  0 '***' 0.001 '**' 0.01 '*' 0.05 '.' 0.1 ' ' 1
(Adjusted p values reported -- none method)

##########################################################


############## BH adjusted p-values ###################
Benjamini, Y., and Hochberg, Y. (1995). Controlling the false discovery rate: a practical and powerful approach t
o multiple testing. Journal of the Royal Statistical Society Series B, 57, 289–300.http://www.jstor.org/stable/23
46101.
##############################################################

     Simultaneous Tests for General Linear Hypotheses

Multiple Comparisons of Means: User-defined Contrasts


Fit: lme.formula(fixed = thisY ~ -1 + EpoTime, data = y, random = ~1 |
    Subject)

Linear Hypotheses:
                          Estimate Std. Error z value Pr(>|z|)
EpoVsPlacebo.EpoWk1 == 0   -0.1742     0.1483  -1.175  0.32005
EpoVsPlacebo.EpoWk2 == 0   -0.3627     0.1483  -2.446  0.02892 *
EpoVsPlacebo.EpoWk3 == 0   -0.3756     0.1483  -2.533  0.02892 *
EpoVsPlacebo.EpoWk4 == 0   -0.4653     0.1483  -3.138  0.00695 **
EpoVsPlacebo.postWk1 == 0  -0.4644     0.1483  -3.132  0.00695 **
EpoVsPlacebo.postWk2 == 0  -0.0809     0.1483  -0.546  0.58540
EpoVsPlacebo.postWk3 == 0  -0.2671     0.1483  -1.801  0.11466
EpoVsPlacebo.postWk4 == 0   0.1387     0.1483   0.935  0.39975
---
Signif. codes:  0 '***' 0.001 '**' 0.01 '*' 0.05 '.' 0.1 ' ' 1
(Adjusted p values reported -- BH method)


##############################################################
```

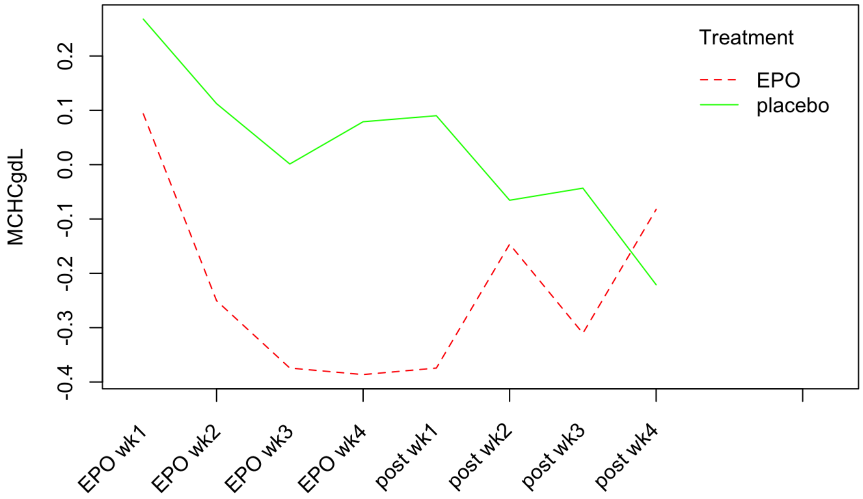

```
##############################################################

########### Repeated measures analysis results for  MCVfl ############

###############################################################
```

| | numDF <int> | denDF <dbl> | F-value <chr> | p-value <chr> |
|---|---|---|---|---|
| (Intercept) | 1 | 224 | 14.637513 | 0.0002 |
| Treatment | 1 | 32 | 2.663040 | 0.1125 |
| time_point | 7 | 224 | 12.843467 | <.0001 |
| Treatment:time_point | 7 | 224 | 6.180984 | <.0001 |

4 rows

```
############ Compare Epo with Palcebo at each time point using contrasts ##########


###############################################################

####################### Raw p-values ########################


     Simultaneous Tests for General Linear Hypotheses

Multiple Comparisons of Means: User-defined Contrasts


Fit: lme.formula(fixed = thisY ~ -1 + EpoTime, data = y, random = ~1 |
    Subject)

Linear Hypotheses:
                          Estimate Std. Error z value Pr(>|z|)
EpoVsPlacebo.EpoWk1 == 0    0.4076     0.4306   0.947  0.34384
EpoVsPlacebo.EpoWk2 == 0    0.9680     0.4306   2.248  0.02456 *
EpoVsPlacebo.EpoWk3 == 0    1.1965     0.4306   2.779  0.00546 **
EpoVsPlacebo.EpoWk4 == 0    1.3934     0.4306   3.236  0.00121 **
EpoVsPlacebo.postWk1 == 0   0.8076     0.4306   1.876  0.06071 .
EpoVsPlacebo.postWk2 == 0   0.1707     0.4306   0.396  0.69178
EpoVsPlacebo.postWk3 == 0   0.3080     0.4306   0.715  0.47437
EpoVsPlacebo.postWk4 == 0  -0.8177     0.4306  -1.899  0.05755 .
---
Signif. codes:  0 ‘***’ 0.001 ‘**’ 0.01 ‘*’ 0.05 ‘.’ 0.1 ‘ ’ 1
(Adjusted p values reported -- none method)

##########################################################


############# BH adjusted p-values ####################
Benjamini, Y., and Hochberg, Y. (1995). Controlling the false discovery rate: a practical and powerful approach t
o multiple testing. Journal of the Royal Statistical Society Series B, 57, 289–300.http://www.jstor.org/stable/23
46101.
##############################################################

     Simultaneous Tests for General Linear Hypotheses

Multiple Comparisons of Means: User-defined Contrasts


Fit: lme.formula(fixed = thisY ~ -1 + EpoTime, data = y, random = ~1 |
    Subject)

Linear Hypotheses:
                          Estimate Std. Error z value Pr(>|z|)
EpoVsPlacebo.EpoWk1 == 0    0.4076     0.4306   0.947   0.4585
EpoVsPlacebo.EpoWk2 == 0    0.9680     0.4306   2.248   0.0655 .
EpoVsPlacebo.EpoWk3 == 0    1.1965     0.4306   2.779   0.0218 *
EpoVsPlacebo.EpoWk4 == 0    1.3934     0.4306   3.236   0.0097 **
EpoVsPlacebo.postWk1 == 0   0.8076     0.4306   1.876   0.0971 .
EpoVsPlacebo.postWk2 == 0   0.1707     0.4306   0.396   0.6918
EpoVsPlacebo.postWk3 == 0   0.3080     0.4306   0.715   0.5421
EpoVsPlacebo.postWk4 == 0  -0.8177     0.4306  -1.899   0.0971 .
---
Signif. codes:  0 ‘***’ 0.001 ‘**’ 0.01 ‘*’ 0.05 ‘.’ 0.1 ‘ ’ 1
(Adjusted p values reported -- BH method)


#############################################################
```

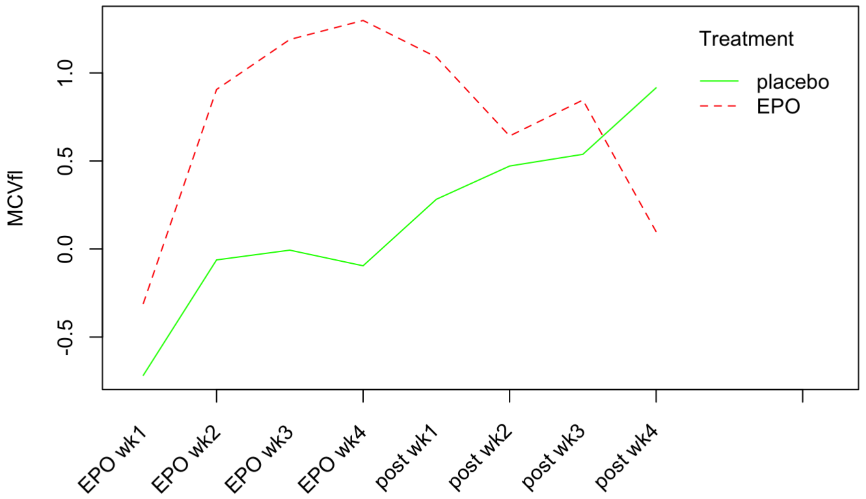


```
################################################################
########### Repeated measures analysis results for  RDWCV ############
#################################################################
```

| | numDF <int> | denDF <dbl> | F-value <chr> | p-value <chr> |
|---|---|---|---|---|
| (Intercept) | 1 | 224 | 35.71804 | <.0001 |
| Treatment | 1 | 32 | 6.79828 | 0.0137 |
| time_point | 7 | 224 | 33.86688 | <.0001 |
| Treatment:time_point | 7 | 224 | 15.77146 | <.0001 |

4 rows

```
############ Compare Epo with Palcebo at each time point using contrasts ##########


###############################################################

####################### Raw p-values #######################


     Simultaneous Tests for General Linear Hypotheses

Multiple Comparisons of Means: User-defined Contrasts


Fit: lme.formula(fixed = thisY ~ -1 + EpoTime, data = y, random = ~1 |
    Subject)

Linear Hypotheses:
                          Estimate Std. Error z value Pr(>|z|)
EpoVsPlacebo.EpoWk1 == 0   0.16689    0.11319   1.474   0.1404
EpoVsPlacebo.EpoWk2 == 0   0.51000    0.11319   4.506 6.61e-06 ***
EpoVsPlacebo.EpoWk3 == 0   0.50422    0.11319   4.455 8.40e-06 ***
EpoVsPlacebo.EpoWk4 == 0   0.61533    0.11319   5.436 5.44e-08 ***
EpoVsPlacebo.postWk1 == 0  0.26689    0.11319   2.358   0.0184 *
EpoVsPlacebo.postWk2 == 0  0.04156    0.11319   0.367   0.7135
EpoVsPlacebo.postWk3 == 0 -0.03844    0.11319  -0.340   0.7341
EpoVsPlacebo.postWk4 == 0 -0.17400    0.11319  -1.537   0.1242
---
Signif. codes:  0 '***' 0.001 '**' 0.01 '*' 0.05 '.' 0.1 ' ' 1
(Adjusted p values reported -- none method)

##########################################################


############## BH adjusted p-values ####################
Benjamini, Y., and Hochberg, Y. (1995). Controlling the false discovery rate: a practical and powerful approach t
o multiple testing. Journal of the Royal Statistical Society Series B, 57, 289–300.http://www.jstor.org/stable/23
46101.
##############################################################

     Simultaneous Tests for General Linear Hypotheses

Multiple Comparisons of Means: User-defined Contrasts


Fit: lme.formula(fixed = thisY ~ -1 + EpoTime, data = y, random = ~1 |
    Subject)

Linear Hypotheses:
                          Estimate Std. Error z value Pr(>|z|)
EpoVsPlacebo.EpoWk1 == 0   0.16689    0.11319   1.474   0.1871
EpoVsPlacebo.EpoWk2 == 0   0.51000    0.11319   4.506 2.24e-05 ***
EpoVsPlacebo.EpoWk3 == 0   0.50422    0.11319   4.455 2.24e-05 ***
EpoVsPlacebo.EpoWk4 == 0   0.61533    0.11319   5.436 4.35e-07 ***
EpoVsPlacebo.postWk1 == 0  0.26689    0.11319   2.358   0.0368 *
EpoVsPlacebo.postWk2 == 0  0.04156    0.11319   0.367   0.7341
EpoVsPlacebo.postWk3 == 0 -0.03844    0.11319  -0.340   0.7341
EpoVsPlacebo.postWk4 == 0 -0.17400    0.11319  -1.537   0.1871
---
Signif. codes:  0 '***' 0.001 '**' 0.01 '*' 0.05 '.' 0.1 ' ' 1
(Adjusted p values reported -- BH method)


##############################################################
```

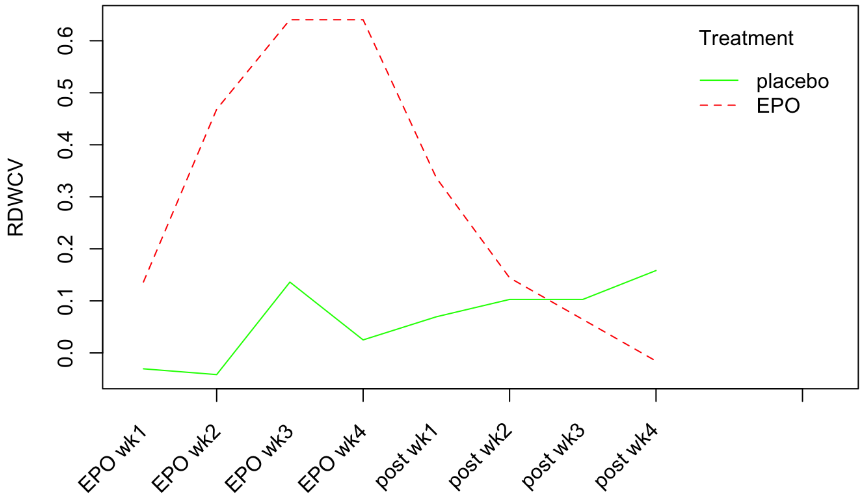


```
##############################################################

########### Repeated measures analysis results for  RDWSDfl ############

##############################################################
```

| | numDF <int> | denDF <dbl> | F-value <chr> | p-value <chr> |
|---|---|---|---|---|
| (Intercept) | 1 | 224 | 39.46842 | <.0001 |

| | numDF <int> | denDF <dbl> | F-value <chr> | p-value <chr> |
|---|---|---|---|---|
| Treatment | 1 | 32 | 8.95499 | 0.0053 |
| time_point | 7 | 224 | 27.83700 | <.0001 |
| Treatment:time_point | 7 | 224 | 12.98759 | <.0001 |

4 rows

```
############ Compare Epo with Palcebo at each time point using contrasts ###########


###############################################################

####################### Raw p-values #######################


     Simultaneous Tests for General Linear Hypotheses

Multiple Comparisons of Means: User-defined Contrasts


Fit: lme.formula(fixed = thisY ~ -1 + EpoTime, data = y, random = ~1 |
    Subject)

Linear Hypotheses:
                          Estimate Std. Error z value Pr(>|z|)
EpoVsPlacebo.EpoWk1 == 0    0.7604     0.4879   1.558  0.11916
EpoVsPlacebo.EpoWk2 == 0    1.9510     0.4879   3.999 6.37e-05 ***
EpoVsPlacebo.EpoWk3 == 0    2.2804     0.4879   4.673 2.96e-06 ***
EpoVsPlacebo.EpoWk4 == 0    2.7235     0.4879   5.582 2.38e-08 ***
EpoVsPlacebo.postWk1 == 0   1.4933     0.4879   3.060  0.00221 **
EpoVsPlacebo.postWk2 == 0   0.4115     0.4879   0.843  0.39906
EpoVsPlacebo.postWk3 == 0   0.2430     0.4879   0.498  0.61842
EpoVsPlacebo.postWk4 == 0  -0.8223     0.4879  -1.685  0.09194 .
---
Signif. codes:  0 '***' 0.001 '**' 0.01 '*' 0.05 '.' 0.1 ' ' 1
(Adjusted p values reported -- none method)

##########################################################


############## BH adjusted p-values ####################
Benjamini, Y., and Hochberg, Y. (1995). Controlling the false discovery rate: a practical and powerful approach t
o multiple testing. Journal of the Royal Statistical Society Series B, 57, 289–300.http://www.jstor.org/stable/23
46101.
##############################################################

     Simultaneous Tests for General Linear Hypotheses

Multiple Comparisons of Means: User-defined Contrasts


Fit: lme.formula(fixed = thisY ~ -1 + EpoTime, data = y, random = ~1 |
    Subject)

Linear Hypotheses:
                          Estimate Std. Error z value Pr(>|z|)
EpoVsPlacebo.EpoWk1 == 0    0.7604     0.4879   1.558  0.15888
EpoVsPlacebo.EpoWk2 == 0    1.9510     0.4879   3.999  0.00017 ***
EpoVsPlacebo.EpoWk3 == 0    2.2804     0.4879   4.673 1.18e-05 ***
EpoVsPlacebo.EpoWk4 == 0    2.7235     0.4879   5.582 1.91e-07 ***
EpoVsPlacebo.postWk1 == 0   1.4933     0.4879   3.060  0.00442 **
EpoVsPlacebo.postWk2 == 0   0.4115     0.4879   0.843  0.45607
EpoVsPlacebo.postWk3 == 0   0.2430     0.4879   0.498  0.61842
EpoVsPlacebo.postWk4 == 0  -0.8223     0.4879  -1.685  0.14711
---
Signif. codes:  0 '***' 0.001 '**' 0.01 '*' 0.05 '.' 0.1 ' ' 1
(Adjusted p values reported -- BH method)


#############################################################
```

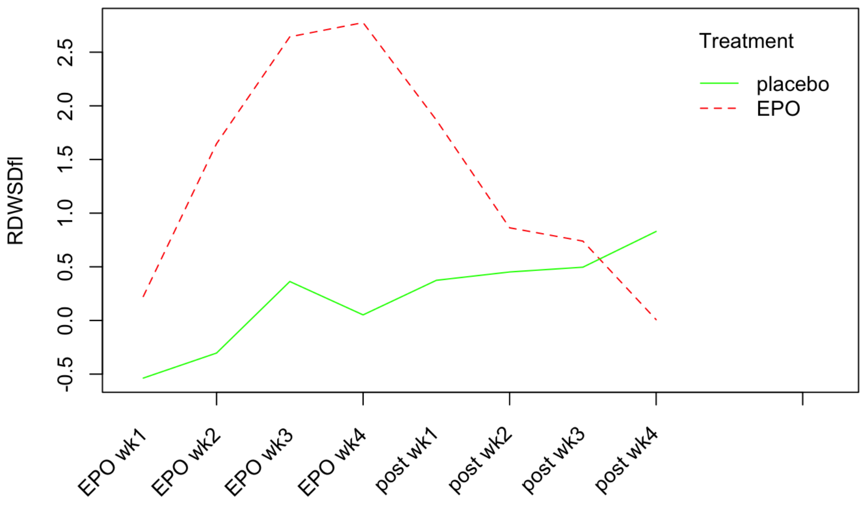

```
##############################################################

########### Repeated measures analysis results for  Retics ############

###############################################################
```

| | numDF <int> | denDF <dbl> | F-value <chr> | p-value <chr> |
|---|---|---|---|---|
| (Intercept) | 1 | 224 | 12.82991 | 0.0004 |
| Treatment | 1 | 32 | 0.32422 | 0.5731 |
| time_point | 7 | 224 | 49.77240 | <.0001 |
| Treatment:time_point | 7 | 224 | 16.77994 | <.0001 |

4 rows

```
############ Compare Epo with Palcebo at each time point using contrasts ##########


###############################################################

####################### Raw p-values ########################


     Simultaneous Tests for General Linear Hypotheses

Multiple Comparisons of Means: User-defined Contrasts


Fit: lme.formula(fixed = thisY ~ -1 + EpoTime, data = y, random = ~1 |
    Subject)

Linear Hypotheses:
                          Estimate Std. Error z value Pr(>|z|)
EpoVsPlacebo.EpoWk1 == 0   0.13632    0.09377   1.454 0.145997
EpoVsPlacebo.EpoWk2 == 0   0.55090    0.09377   5.875 4.23e-09 ***
EpoVsPlacebo.EpoWk3 == 0   0.35037    0.09377   3.737 0.000187 ***
EpoVsPlacebo.EpoWk4 == 0   0.23574    0.09377   2.514 0.011933 *
EpoVsPlacebo.postWk1 == 0 -0.33403    0.09377  -3.562 0.000368 ***
EpoVsPlacebo.postWk2 == 0 -0.37479    0.09377  -3.997 6.42e-05 ***
EpoVsPlacebo.postWk3 == 0 -0.17168    0.09377  -1.831 0.067120 .
EpoVsPlacebo.postWk4 == 0 -0.14937    0.09377  -1.593 0.111176
---
Signif. codes:  0 ‘***’ 0.001 ‘**’ 0.01 ‘*’ 0.05 ‘.’ 0.1 ‘ ’ 1
(Adjusted p values reported -- none method)

##########################################################


############# BH adjusted p-values ####################
Benjamini, Y., and Hochberg, Y. (1995). Controlling the false discovery rate: a practical and powerful approach t
o multiple testing. Journal of the Royal Statistical Society Series B, 57, 289–300.http://www.jstor.org/stable/23
46101.
##############################################################

     Simultaneous Tests for General Linear Hypotheses

Multiple Comparisons of Means: User-defined Contrasts


Fit: lme.formula(fixed = thisY ~ -1 + EpoTime, data = y, random = ~1 |
    Subject)

Linear Hypotheses:
                          Estimate Std. Error z value Pr(>|z|)
EpoVsPlacebo.EpoWk1 == 0   0.13632    0.09377   1.454 0.145997
EpoVsPlacebo.EpoWk2 == 0   0.55090    0.09377   5.875 3.38e-08 ***
EpoVsPlacebo.EpoWk3 == 0   0.35037    0.09377   3.737 0.000498 ***
EpoVsPlacebo.EpoWk4 == 0   0.23574    0.09377   2.514 0.019093 *
EpoVsPlacebo.postWk1 == 0 -0.33403    0.09377  -3.562 0.000735 ***
EpoVsPlacebo.postWk2 == 0 -0.37479    0.09377  -3.997 0.000257 ***
EpoVsPlacebo.postWk3 == 0 -0.17168    0.09377  -1.831 0.089494 .
EpoVsPlacebo.postWk4 == 0 -0.14937    0.09377  -1.593 0.127058
---
Signif. codes:  0 ‘***’ 0.001 ‘**’ 0.01 ‘*’ 0.05 ‘.’ 0.1 ‘ ’ 1
(Adjusted p values reported -- BH method)


#############################################################
```

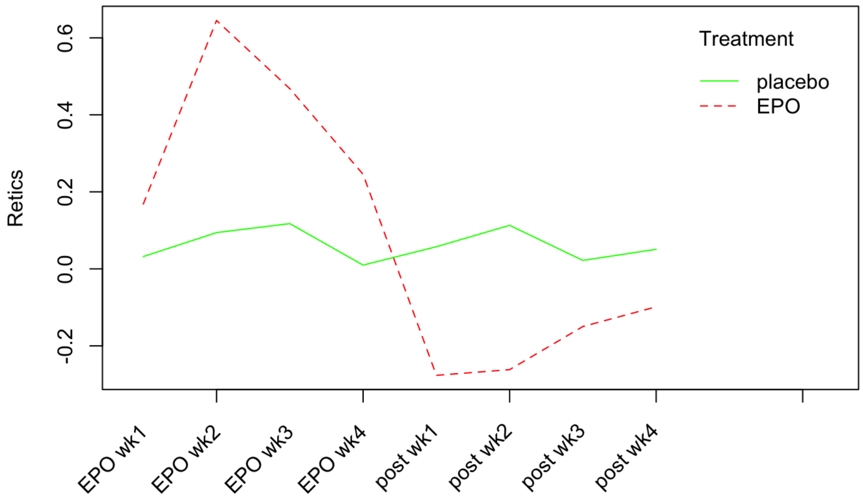


```
###############################################################
########### Repeated measures analysis results for  Retics109L ############
###############################################################
```

| | numDF <int> | denDF <dbl> | F-value <chr> | p-value <chr> |
|---|---|---|---|---|
| (Intercept) | 1 | 224 | 14.67996 | 0.0002 |
| Treatment | 1 | 32 | 1.33941 | 0.2557 |
| time_point | 7 | 224 | 52.94137 | <.0001 |
| Treatment:time_point | 7 | 224 | 16.77202 | <.0001 |

4 rows

```
############ Compare Epo with Palcebo at each time point using contrasts ###########


###############################################################

######################## Raw p-values ########################


     Simultaneous Tests for General Linear Hypotheses

Multiple Comparisons of Means: User-defined Contrasts


Fit: lme.formula(fixed = thisY ~ -1 + EpoTime, data = y, random = ~1 |
    Subject)

Linear Hypotheses:
                          Estimate Std. Error z value Pr(>|z|)
EpoVsPlacebo.EpoWk1 == 0     6.681      4.377   1.527 0.126865
EpoVsPlacebo.EpoWk2 == 0    26.618      4.377   6.082 1.19e-09 ***
EpoVsPlacebo.EpoWk3 == 0    18.323      4.377   4.187 2.83e-05 ***
EpoVsPlacebo.EpoWk4 == 0    13.460      4.377   3.075 0.002102 **
EpoVsPlacebo.postWk1 == 0  -13.196      4.377  -3.015 0.002569 **
EpoVsPlacebo.postWk2 == 0  -16.797      4.377  -3.838 0.000124 ***
EpoVsPlacebo.postWk3 == 0   -6.748      4.377  -1.542 0.123122
EpoVsPlacebo.postWk4 == 0   -5.203      4.377  -1.189 0.234516
---
Signif. codes:  0 '***' 0.001 '**' 0.01 '*' 0.05 '.' 0.1 ' ' 1
(Adjusted p values reported -- none method)

##########################################################


############## BH adjusted p-values ####################
Benjamini, Y., and Hochberg, Y. (1995). Controlling the false discovery rate: a practical and powerful approach t
o multiple testing. Journal of the Royal Statistical Society Series B, 57, 289–300.http://www.jstor.org/stable/23
46101.
##############################################################

     Simultaneous Tests for General Linear Hypotheses

Multiple Comparisons of Means: User-defined Contrasts


Fit: lme.formula(fixed = thisY ~ -1 + EpoTime, data = y, random = ~1 |
    Subject)

Linear Hypotheses:
                          Estimate Std. Error z value Pr(>|z|)
EpoVsPlacebo.EpoWk1 == 0     6.681      4.377   1.527 0.144988
EpoVsPlacebo.EpoWk2 == 0    26.618      4.377   6.082  9.5e-09 ***
EpoVsPlacebo.EpoWk3 == 0    18.323      4.377   4.187 0.000113 ***
EpoVsPlacebo.EpoWk4 == 0    13.460      4.377   3.075 0.004110 **
EpoVsPlacebo.postWk1 == 0  -13.196      4.377  -3.015 0.004110 **
EpoVsPlacebo.postWk2 == 0  -16.797      4.377  -3.838 0.000331 ***
EpoVsPlacebo.postWk3 == 0   -6.748      4.377  -1.542 0.144988
EpoVsPlacebo.postWk4 == 0   -5.203      4.377  -1.189 0.234516
---
Signif. codes:  0 '***' 0.001 '**' 0.01 '*' 0.05 '.' 0.1 ' ' 1
(Adjusted p values reported -- BH method)
```

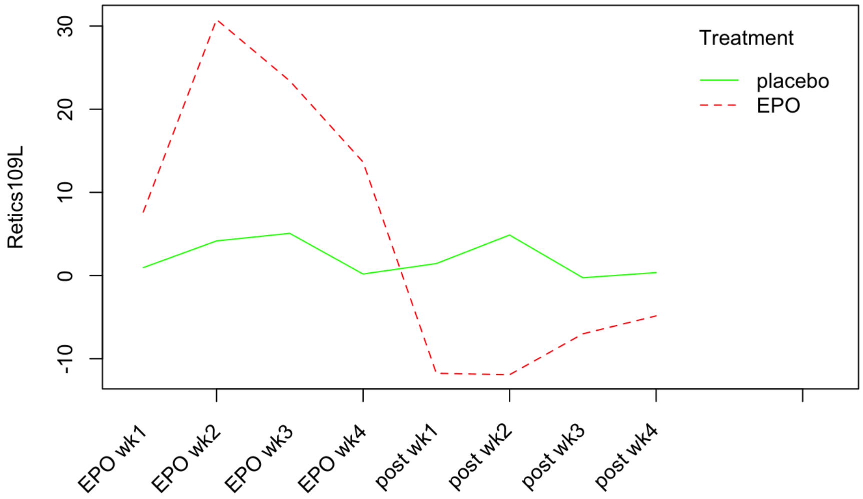


Data visualization

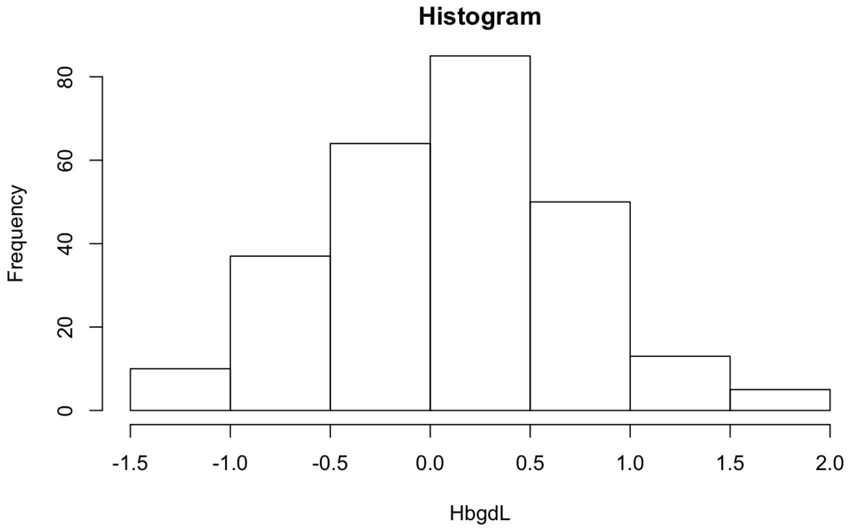
Histogram
Frequency
0
20
40
60
80
-1.5
-1.0
-0.5
0.0
0.5
1.0
1.5
2.0
HbgdL

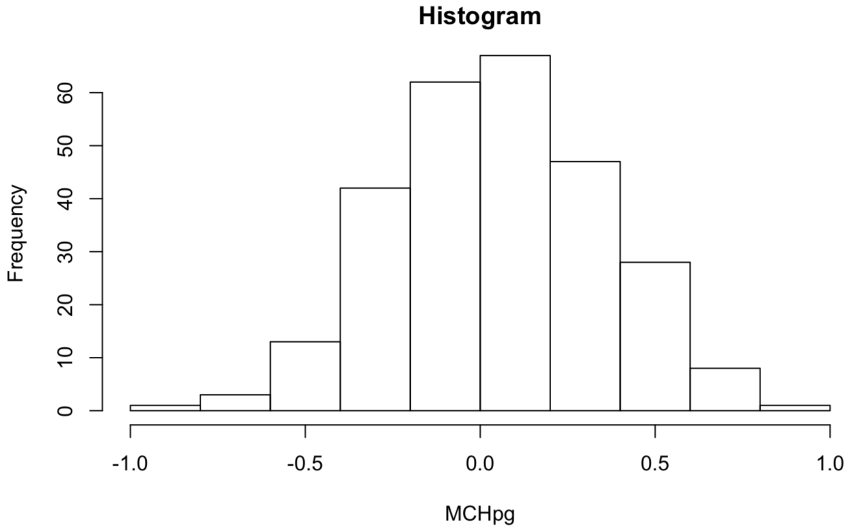
Histogram
Frequency
0
10
20
30
40
50
60
-1.0
-0.5
0.0
0.5
1.0
MCHpg

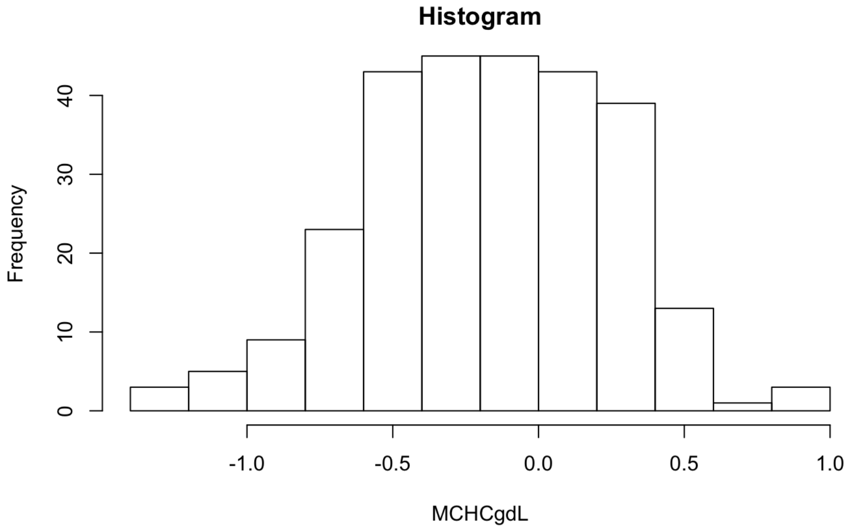
Histogram
Frequency
0
10
20
30
40
-1.0
-0.5
0.0
0.5
1.0
MCHCgdL

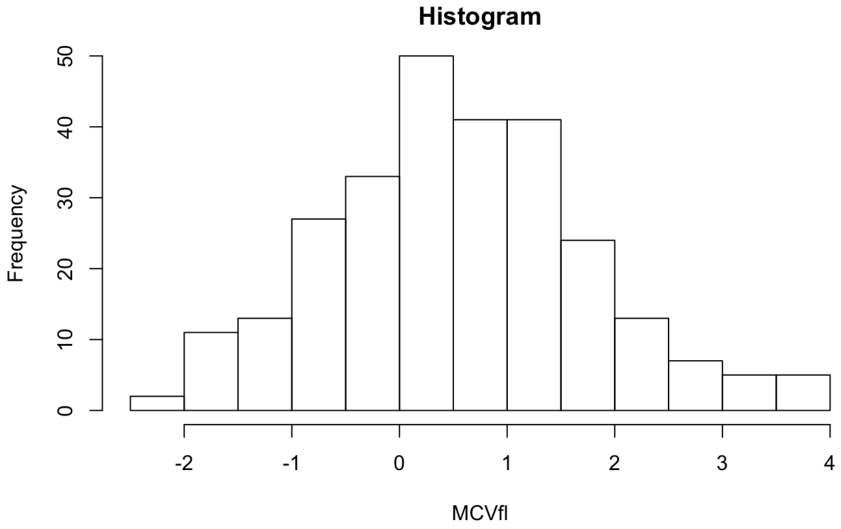
Histogram
Frequency
MCVfl

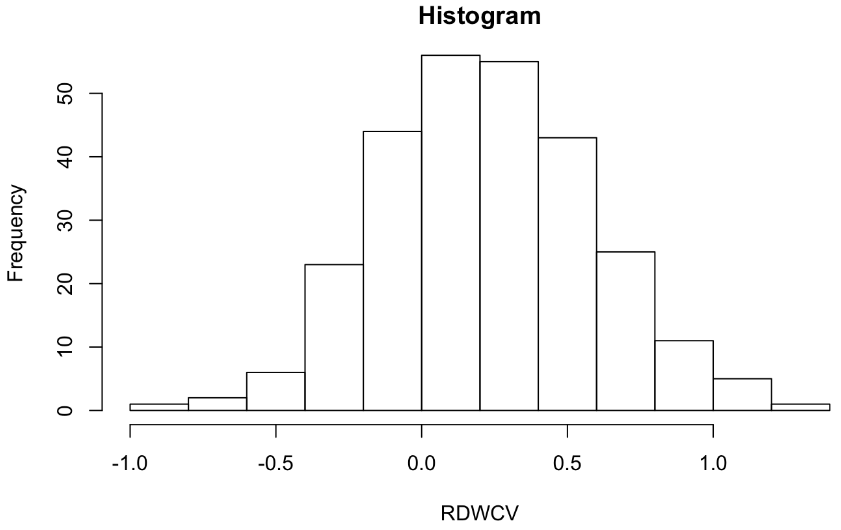
Histogram
Frequency
RDWCV

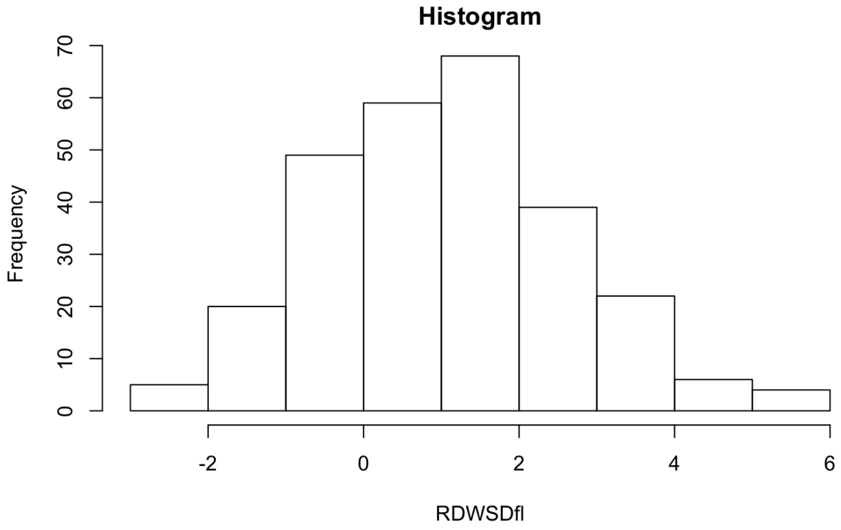
Histogram
Frequency
RDWSDfl

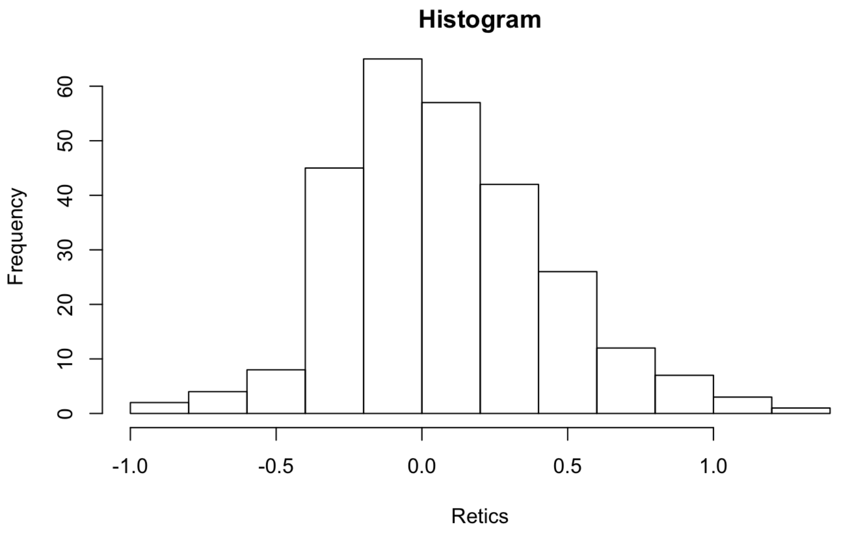
Histogram
Frequency
0
10
20
30
40
50
60
-1.0
-0.5
0.0
0.5
1.0
Retics

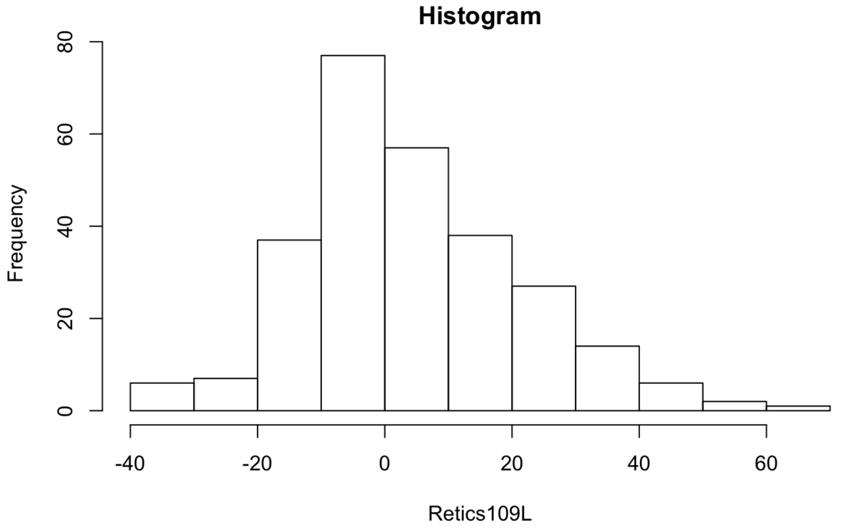
Histogram
Frequency
0
20
40
60
80
-40
-20
0
20
40
60
Retics109L

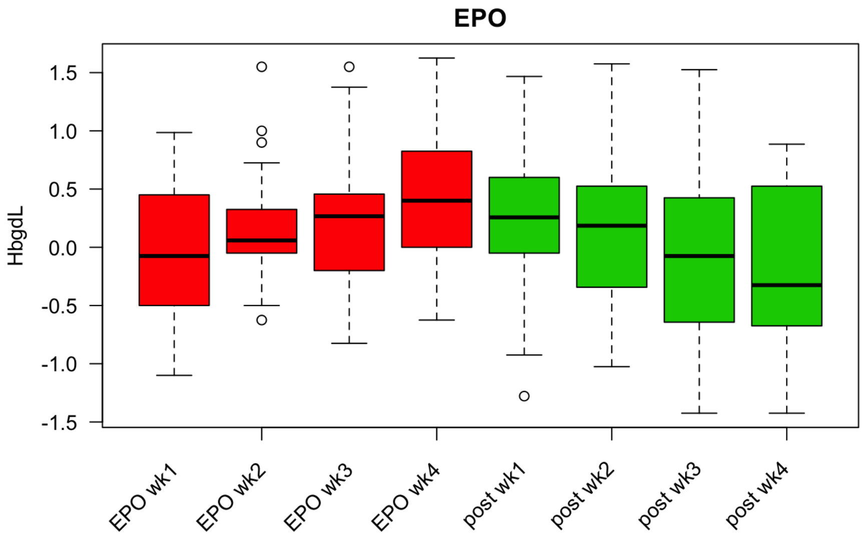
EPO
HbgdL
1.5
1.0
0.5
0.0
-0.5
-1.0
-1.5
EPO wk1
EPO wk2
EPO wk3
EPO wk4
post wk1
post wk2
post wk3
post wk4

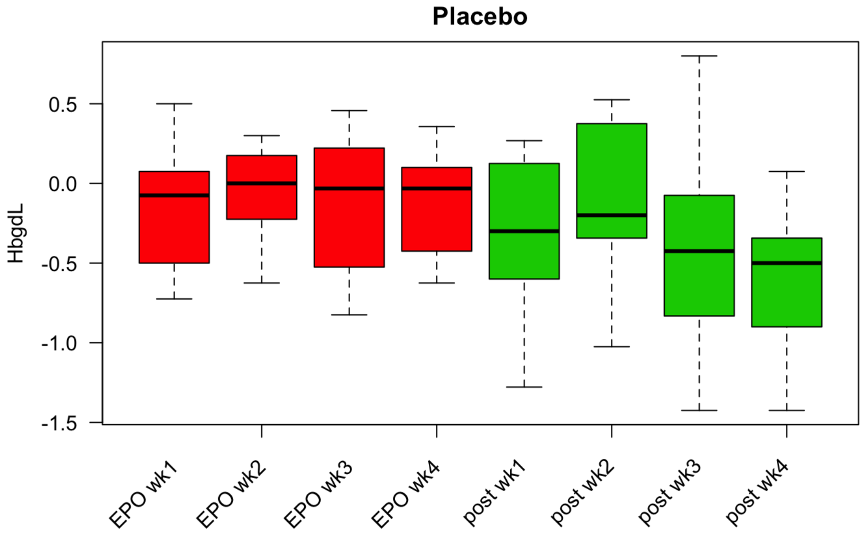
Placebo
HbgdL
0.5
0.0
-0.5
-1.0
-1.5
EPO wk1
EPO wk2
EPO wk3
EPO wk4
post wk1
post wk2
post wk3
post wk4

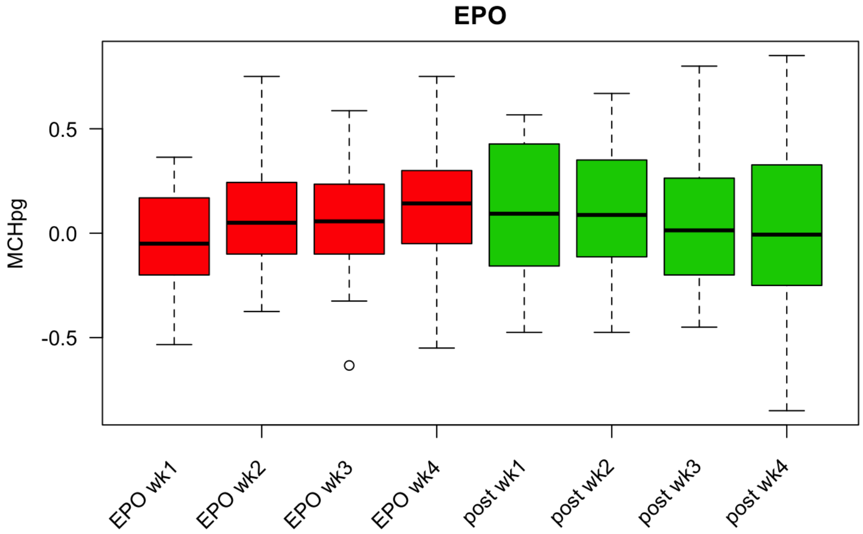
EPO
MCHpg
0.5
0.0
-0.5
EPO wk1
EPO wk2
EPO wk3
EPO wk4
post wk1
post wk2
post wk3
post wk4

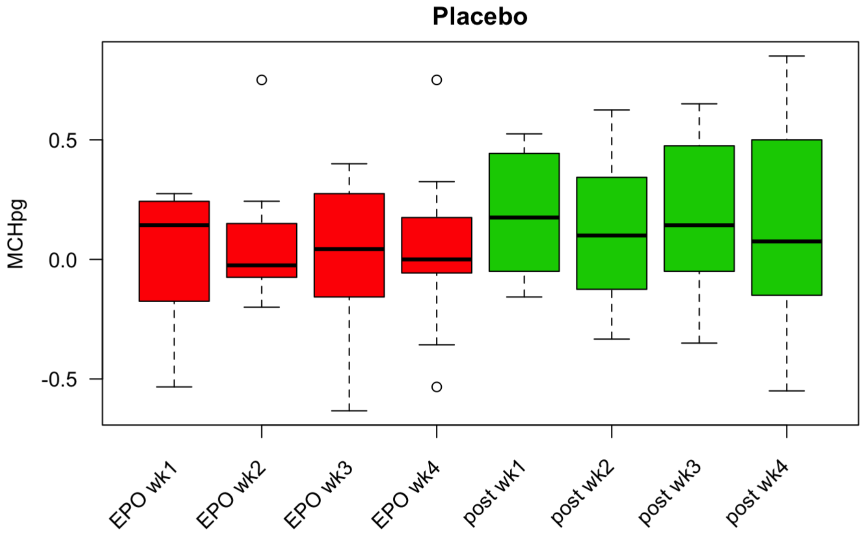
Placebo
MCHpg
0.5
0.0
-0.5
EPO wk1
EPO wk2
EPO wk3
EPO wk4
post wk1
post wk2
post wk3
post wk4

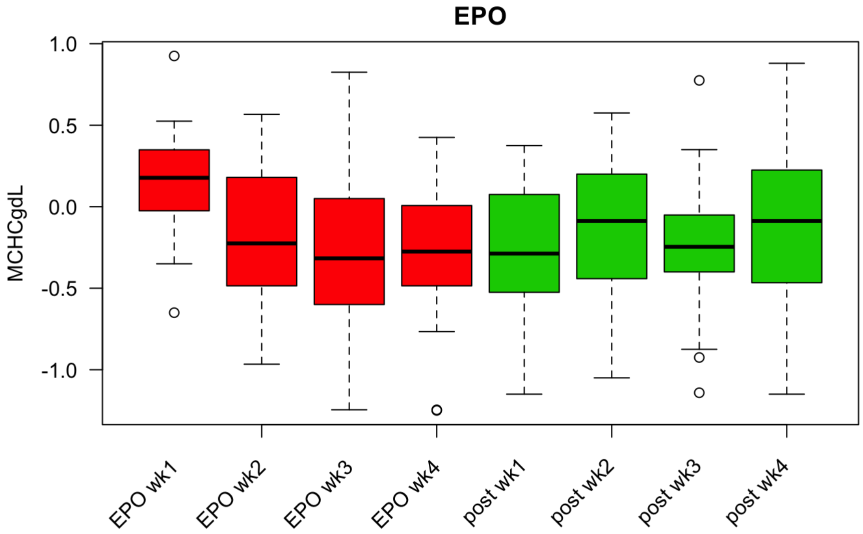
EPO
MCHCgdL
1.0
0.5
0.0
-0.5
-1.0
EPO wk1
EPO wk2
EPO wk3
EPO wk4
post wk1
post wk2
post wk3
post wk4

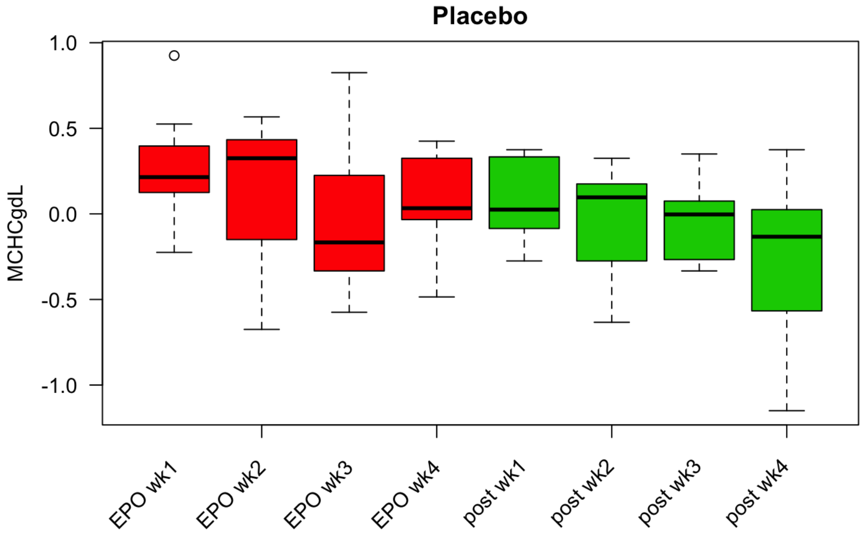
Placebo
MCHCgdL
1.0
0.5
0.0
-0.5
-1.0
EPO wk1
EPO wk2
EPO wk3
EPO wk4
post wk1
post wk2
post wk3
post wk4

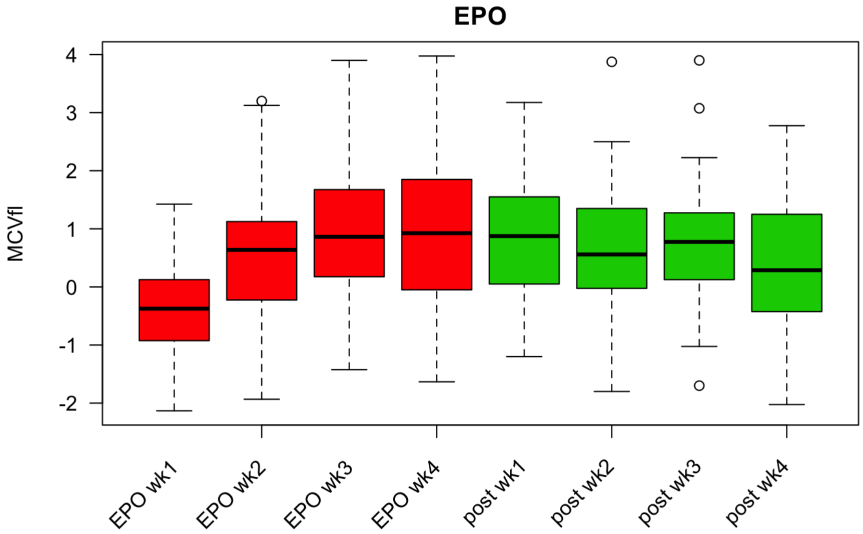
EPO
MCVfl
4
3
2
1
0
-1
-2
EPO wk1
EPO wk2
EPO wk3
EPO wk4
post wk1
post wk2
post wk3
post wk4

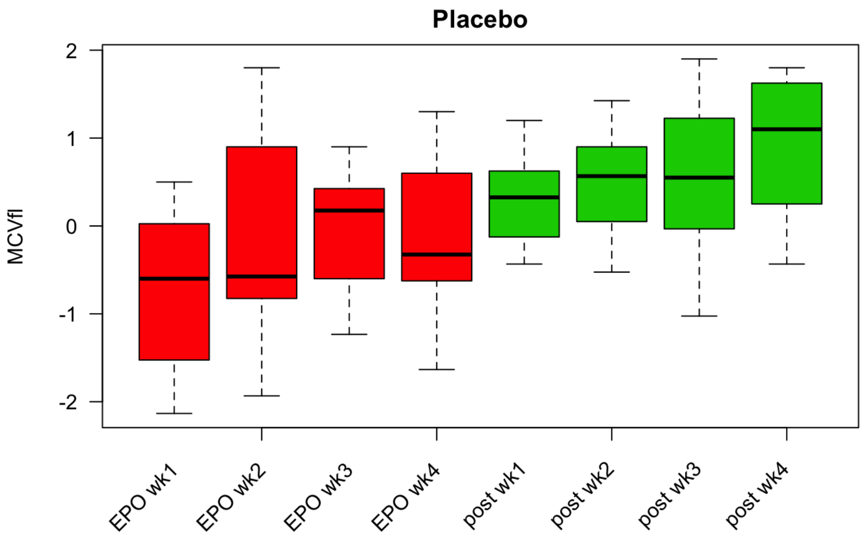
Placebo
MCVfl
2
1
0
-1
-2
EPO wk1
EPO wk2
EPO wk3
EPO wk4
post wk1
post wk2
post wk3
post wk4

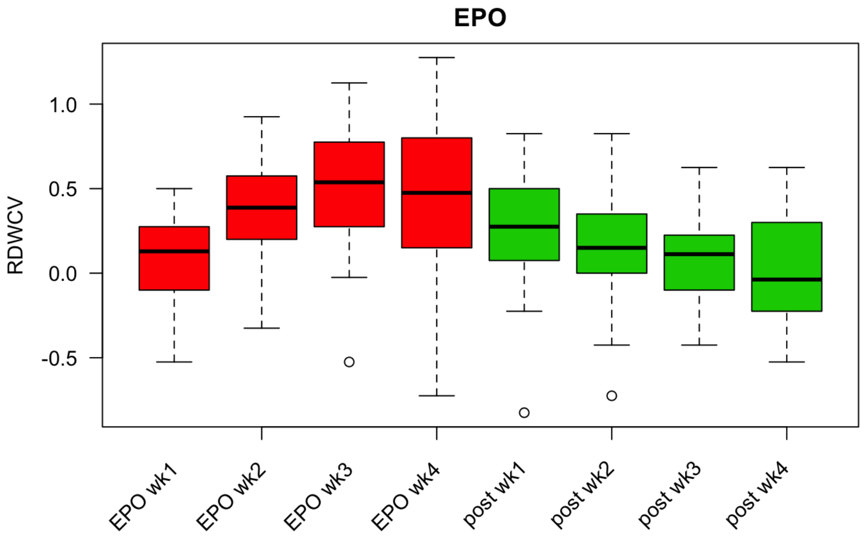
EPO
RDWCV
1.0
0.5
0.0
-0.5
EPO wk1
EPO wk2
EPO wk3
EPO wk4
post wk1
post wk2
post wk3
post wk4

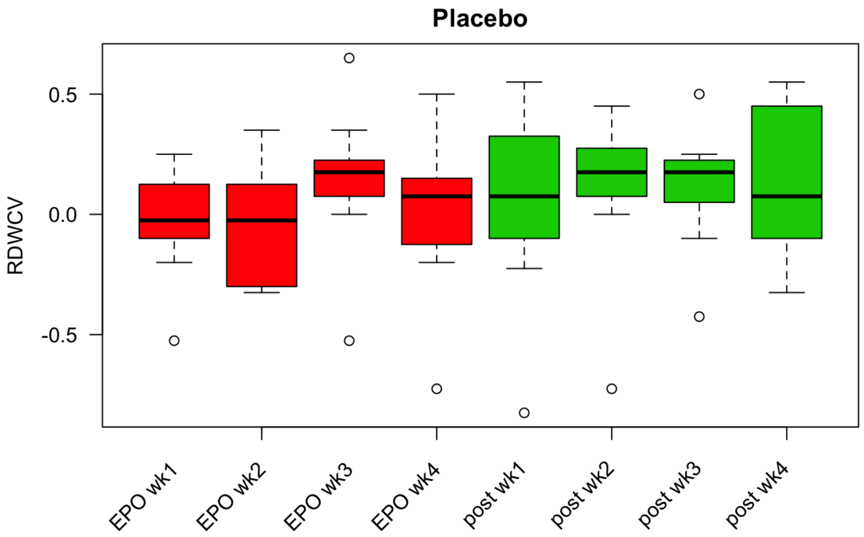
Placebo
RDWCV
0.5
0.0
-0.5
EPO wk1
EPO wk2
EPO wk3
EPO wk4
post wk1
post wk2
post wk3
post wk4

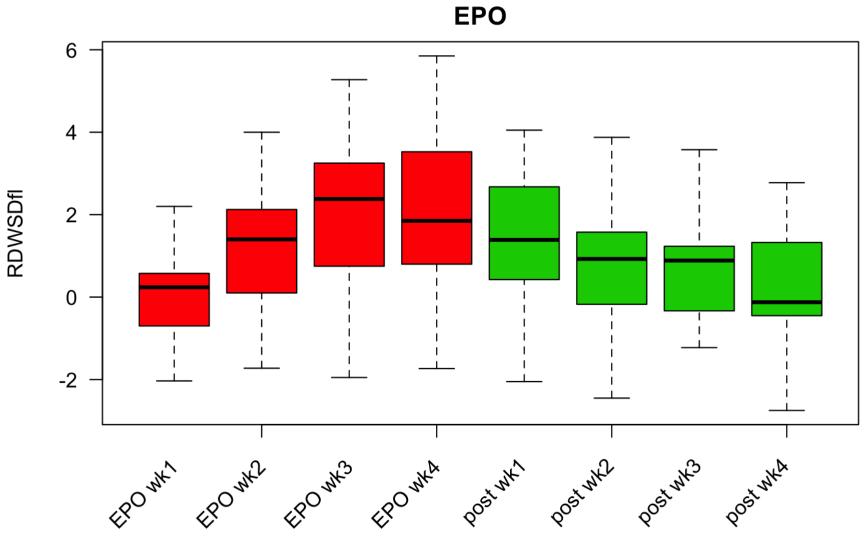
EPO
RDWSDfl
6
4
2
0
-2
EPO wk1
EPO wk2
EPO wk3
EPO wk4
post wk1
post wk2
post wk3
post wk4

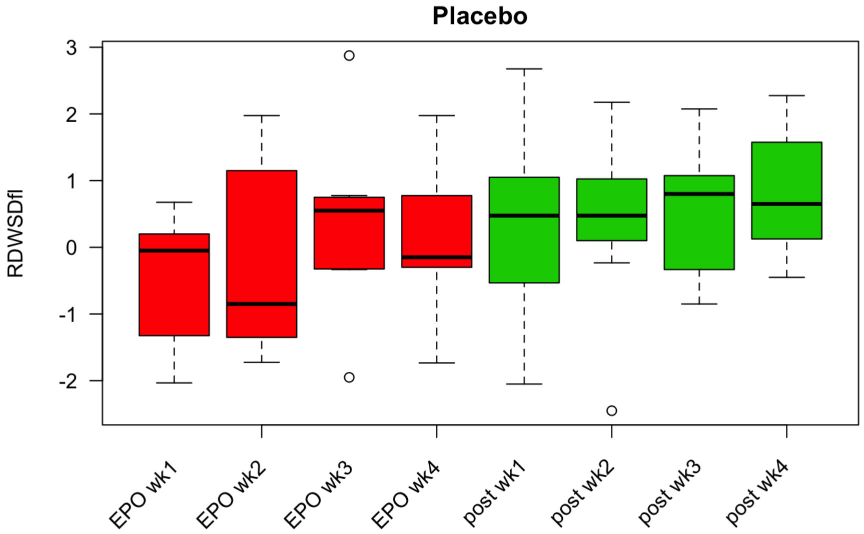
Placebo
RDWSDfl
3
2
1
0
-1
-2
EPO wk1
EPO wk2
EPO wk3
EPO wk4
post wk1
post wk2
post wk3
post wk4

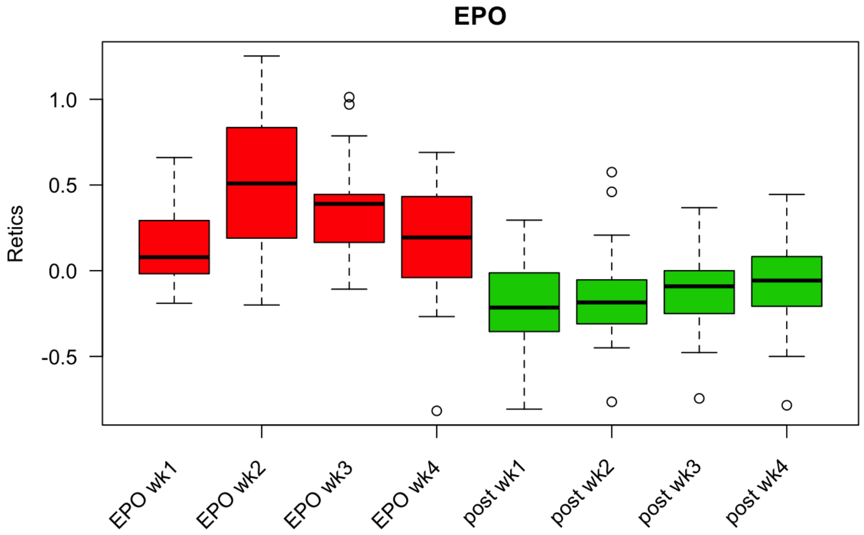
EPO
Retics
1.0
0.5
0.0
-0.5
EPO wk1
EPO wk2
EPO wk3
EPO wk4
post wk1
post wk2
post wk3
post wk4

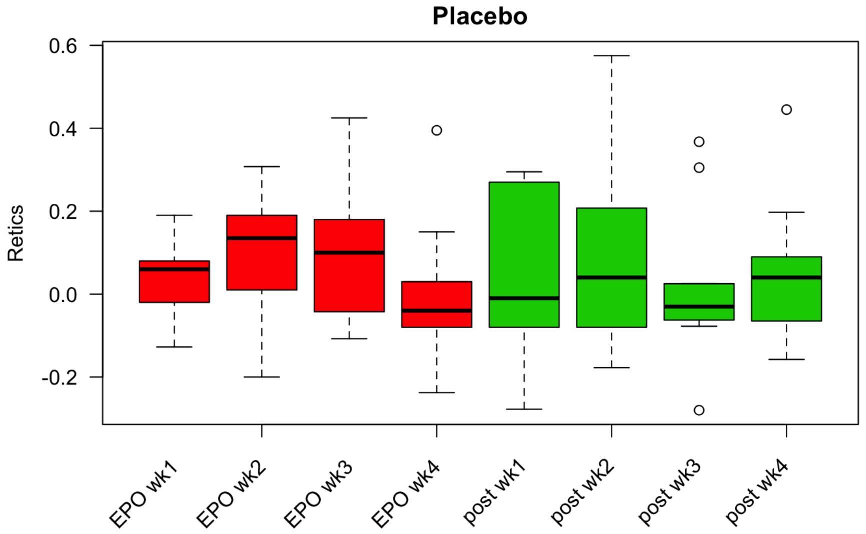
Placebo
Retics
0.6
0.4
0.2
0.0
-0.2
EPO wk1
EPO wk2
EPO wk3
EPO wk4
post wk1
post wk2
post wk3
post wk4

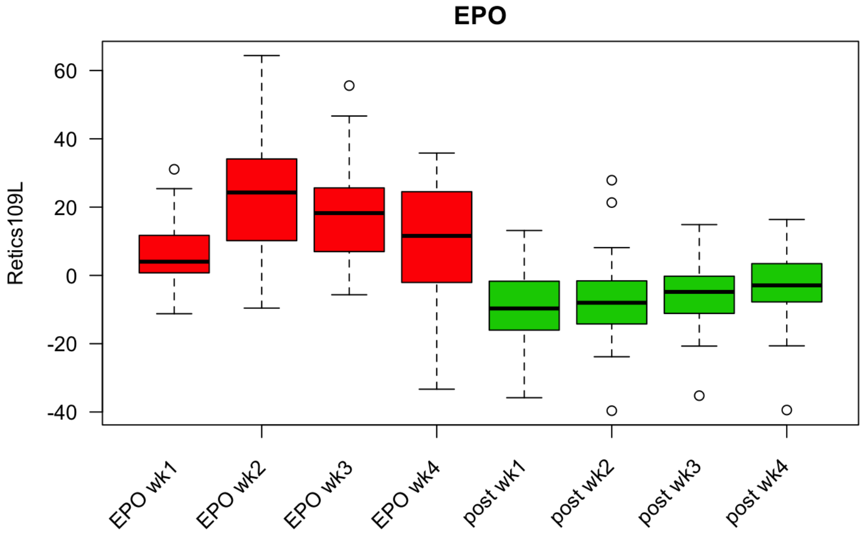
EPO
Retics109L
60
40
20
0
-20
-40
EPO wk1
EPO wk2
EPO wk3
EPO wk4
post wk1
post wk2
post wk3
post wk4

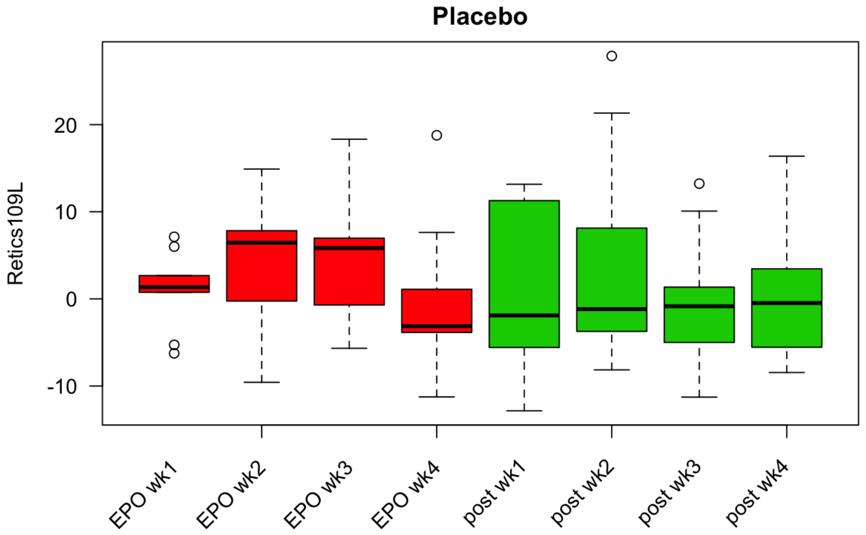
Placebo
Retics109L
20
10
0
-10
EPO wk1
EPO wk2
EPO wk3
EPO wk4
post wk1
post wk2
post wk3
post wk4

```
R version 3.6.1 Patched (2019-09-23 r77210)
Platform: x86_64-apple-darwin15.6.0 (64-bit)
Running under: macOS Mojave 10.14.6

Matrix products: default
BLAS:   /System/Library/Frameworks/Accelerate.framework/Versions/A/Frameworks/vecLib.framework/Versions/A/libBLA
S.dylib
LAPACK: /Library/Frameworks/R.framework/Versions/3.6/Resources/lib/libRlapack.dylib

locale:
[1] en_US.UTF-8/en_US.UTF-8/en_US.UTF-8/C/en_US.UTF-8/en_US.UTF-8

attached base packages:
[1] stats     graphics  grDevices utils     datasets  methods   base

other attached packages:
[1] multcomp_1.4-12 TH.data_1.0-10  MASS_7.3-51.5   survival_3.1-8  mvtnorm_1.0-12  nlme_3.1-143

loaded via a namespace (and not attached):
 [1] Rcpp_1.0.3       knitr_1.27       magrittr_1.5     splines_3.6.1    lattice_0.20-38
 [6] rlang_0.4.2      stringr_1.4.0    tools_3.6.1      grid_3.6.1       xfun_0.12
[11] htmltools_0.4.0  yaml_2.2.0       digest_0.6.23    Matrix_1.2-18    base64enc_0.1-3
[16] codetools_0.2-16 rsconnect_0.8.16 evaluate_0.14    rmarkdown_2.1    sandwich_2.5-1
[21] stringi_1.4.5    compiler_3.6.1   jsonlite_1.6     zoo_1.8-7
```

Code ▾

### Data analysis of Epo intervention study # 2, Hidalgo et al.

##### By Lihua Julie Zhu

Citation:

1. 1. Lindstrom, M.J. and Bates, D.M. (1988) “Newton-Raphson and EM Algorithms for Linear Mixed-Effects Models for Repeated-Measures Data”, Journal of the American Statistical Association, 83, 1014–1022.
2. Hothorn T, Bretz F, Westfall P (2008). “Simultaneous Inference in General Parametric Models.” Biometrical Journal, 50(3), 346–363.
3. Benjamini, Y., and Hochberg, Y. (1995). Controlling the false discovery rate: a practical and powerful approach to multiple testing. Journal of the Royal Statistical Society Series B, 57, 289–300.http://www.jstor.org/stable/2346101.

Data summary

```
[1] "Thu Mar 19 18:28:21 2020"
[1] 384  14
```

| | Treatment <fctr> | SubjectNr <int> | Days_corrected <dbl> | Erythrocytes <dbl> | Hematocrit <dbl> | Hemoglobin <dbl> | IRF <dbl> | MCH <dbl> | M... <dbl> |
|---|---|---|---|---|---|---|---|---|---|
| 1 | Epo | 2 | -25 | 4.90 | 0.430 | 8.7 | NA | 1.780 | 20.2 |
| 2 | Epo | 2 | 0 | 4.98 | 0.436 | 8.7 | 9.7 | 1.750 | 20.0 |
| 3 | Epo | 2 | 13 | 5.11 | 0.452 | 9.0 | 8.7 | 1.761 | 19.9 |
| 4 | Epo | 2 | 27 | 5.18 | 0.462 | 9.1 | 5.9 | 1.757 | 19.7 |
| 5 | Epo | 2 | 41 | 5.23 | 0.469 | 9.2 | 9.8 | 1.759 | 19.6 |
| 6 | Epo | 2 | 61 | 5.07 | 0.455 | 9.0 | 4.1 | 1.775 | 19.8 |

6 rows | 1-10 of 14 columns

```
[1] 384  14
```

| | Treatment <fctr> | Subject <int> | time_point <dbl> | Erythrocytes <dbl> | Hematocrit <dbl> | Hemoglobin <dbl> | IRF <dbl> | MCH <dbl> | MC... <dbl> |
|---|---|---|---|---|---|---|---|---|---|
| 1 | Epo | 2 | -25 | 4.90 | 0.430 | 8.7 | NA | 1.780 | 20.2 |
| 2 | Epo | 2 | 0 | 4.98 | 0.436 | 8.7 | 9.7 | 1.750 | 20.0 |
| 3 | Epo | 2 | 13 | 5.11 | 0.452 | 9.0 | 8.7 | 1.761 | 19.9 |
| 4 | Epo | 2 | 27 | 5.18 | 0.462 | 9.1 | 5.9 | 1.757 | 19.7 |
| 5 | Epo | 2 | 41 | 5.23 | 0.469 | 9.2 | 9.8 | 1.759 | 19.6 |
| 6 | Epo | 2 | 61 | 5.07 | 0.455 | 9.0 | 4.1 | 1.775 | 19.8 |

6 rows | 1-10 of 14 columns

```
Correlation matrix among different measurements
             Erythrocytes  Hematocrit  Hemoglobin         IRF         MCH         MCHC         MCV
Erythrocytes   1.00000000  0.82382229  0.76224650 -0.21111015 -0.46730438 -0.123225940 -0.46685009
Hematocrit     0.82382229  1.00000000  0.92171103 -0.16972995  0.01177281 -0.152100261  0.11359931
Hemoglobin     0.76224650  0.92171103  1.00000000 -0.19673417  0.21254048  0.241721958  0.09190742
IRF           -0.21111015 -0.16972995 -0.19673417  1.00000000  0.04333773 -0.083806217  0.10655835
MCH           -0.46730438  0.01177281  0.21254048  0.04333773  1.00000000  0.520135885  0.83146491
MCHC          -0.12322594 -0.15210026  0.24172196 -0.08380622  0.52013588  1.000000000 -0.04142845
MCV           -0.46685009  0.11359931  0.09190742  0.10655835  0.83146491 -0.041428450  1.00000000
RDW_SD        -0.17643173  0.11537341 -0.01572582  0.28517235  0.25782734 -0.342623100  0.52477730
Reticulo_per  -0.11694348 -0.07618071 -0.04115495  0.56324228  0.12007292  0.096009557  0.07928422
Reticulocyte   0.11219894  0.11718677  0.13663948  0.51440820  0.01013854  0.056001662 -0.02290604
Thrombocytes   0.05332675  0.08754265  0.08085743  0.05574185  0.02160299 -0.007037408  0.03028515
                  RDW_SD Reticulo_per Reticulocyte Thrombocytes
Erythrocytes -0.17643173  -0.11694348   0.11219894  0.053326750
Hematocrit    0.11537341  -0.07618071   0.11718677  0.087542655
Hemoglobin   -0.01572582  -0.04115495   0.13663948  0.080857433
IRF           0.28517235   0.56324228   0.51440820  0.055741847
MCH           0.25782734   0.12007292   0.01013854  0.021602992
MCHC         -0.34262310   0.09600956   0.05600166 -0.007037408
MCV           0.52477730   0.07928422  -0.02290604  0.030285153
RDW_SD        1.00000000   0.15792487   0.12172182 -0.051137552
Reticulo_per  0.15792487   1.00000000   0.97429694  0.307169691
Reticulocyte  0.12172182   0.97429694   1.00000000  0.297618890
Thrombocytes -0.05113755   0.30716969   0.29761889  1.000000000
```

Compute mean of 4 baseline points for each subject

| | Treatment <fctr> | Subject <int> | time_point <dbl> | Erythrocytes <dbl> | Hematocrit <dbl> | Hemoglobin <dbl> | IRF <dbl> | MCH <dbl> | MC... <dbl> |
|---|---|---|---|---|---|---|---|---|---|
| 1 | Epo | 13 | -25 | 5.04 | 0.453 | 9.4 | NA | 1.860 | 20.8 |
| 2 | Epo | 13 | 0 | 4.87 | 0.440 | 9.0 | 13.9 | 1.850 | 20.5 |
| 3 | Epo | 13 | 13 | 5.01 | 0.462 | 9.2 | 13.3 | 1.836 | 19.9 |
| 4 | Epo | 13 | 27 | 5.26 | 0.489 | 9.6 | 15.0 | 1.825 | 19.6 |
| 5 | Epo | 13 | 41 | 5.35 | 0.492 | 9.9 | 11.8 | 1.850 | 20.1 |
| 6 | Epo | 13 | 61 | 5.41 | 0.495 | 9.8 | 8.7 | 1.811 | 19.8 |

6 rows | 1-10 of 25 columns

```
[1] 288  25
[1] 13 27 41 61 62 80
```

| | Treatment <fctr> | Subject <int> | time_point <dbl> | Erythrocytes <dbl> | Hematocrit <dbl> | Hemoglobin <dbl> | IRF <dbl> | MCH <dbl> | MCHC <dbl> |
|---|---|---|---|---|---|---|---|---|---|
| 3 | Epo | 13 | 13 | 0.055 | 0.0155 | 0.0 | -0.6 | -0.019 | -0.75 |

| | Treatment <fctr> | Subject <int> | time_point <dbl> | Erythrocytes <dbl> | Hematocrit <dbl> | Hemoglobin <dbl> | IRF <dbl> | MCH <dbl> | MCHC <dbl> |
|---|---|---|---|---|---|---|---|---|---|
| 4 | Epo | 13 | 27 | 0.305 | 0.0425 | 0.4 | 1.1 | -0.030 | -1.05 |
| 5 | Epo | 13 | 41 | 0.395 | 0.0455 | 0.7 | -2.1 | -0.005 | -0.55 |
| 6 | Epo | 13 | 61 | 0.455 | 0.0485 | 0.6 | -5.2 | -0.044 | -0.85 |
| 7 | Epo | 13 | 62 | 0.375 | 0.0385 | 0.6 | -8.6 | -0.016 | -0.45 |
| 8 | Epo | 13 | 80 | 0.405 | 0.0515 | 0.6 | -3.8 | -0.027 | -0.95 |

6 rows | 1-10 of 25 columns

```
########Number of Subjects for each treatment ######
    Epo Placebo
     24      24
```

```
#############################################################


##############################################################

########### Repeated measures analysis results for  Erythrocytes ############

###############################################################
```

| | numDF <int> | denDF <dbl> | F-value <chr> | p-value <chr> |
|---|---|---|---|---|
| (Intercept) | 1 | 221 | 55.57089 | <.0001 |
| Treatment | 1 | 46 | 59.51913 | <.0001 |
| time_point | 5 | 221 | 30.48662 | <.0001 |
| Treatment:time_point | 5 | 221 | 3.86877 | 0.0022 |

4 rows

```
############ Compare Epo with Palcebo at each time point using contrasts ###########


###############################################################

####################### Raw p-values #######################


     Simultaneous Tests for General Linear Hypotheses

Multiple Comparisons of Means: User-defined Contrasts


Fit: lme.formula(fixed = thisY ~ -1 + EpoTime, data = y, random = ~1 |
    Subject)

Linear Hypotheses:
                            Estimate Std. Error z value Pr(>|z|)
EpoVsPlacebo.EpoDay13 == 0   0.20542    0.05979   3.435 0.000591 ***
EpoVsPlacebo.EpoDay27 == 0   0.32784    0.06079   5.393 6.94e-08 ***
EpoVsPlacebo.EpoDay41 == 0   0.34185    0.06032   5.667 1.45e-08 ***
EpoVsPlacebo.EpoDay61 == 0   0.39138    0.06085   6.432 1.26e-10 ***
EpoVsPlacebo.postDay62 == 0  0.45753    0.06187   7.395 1.42e-13 ***
EpoVsPlacebo.postDay80 == 0  0.22375    0.05979   3.742 0.000182 ***
---
Signif. codes:  0 '***' 0.001 '**' 0.01 '*' 0.05 '.' 0.1 ' ' 1
(Adjusted p values reported -- none method)

##########################################################


############# BH adjusted p-values ####################
Benjamini, Y., and Hochberg, Y. (1995). Controlling the false discovery rate: a practical and powerful approach t
o multiple testing. Journal of the Royal Statistical Society Series B, 57, 289–300.http://www.jstor.org/stable/23
46101.
##############################################################

     Simultaneous Tests for General Linear Hypotheses

Multiple Comparisons of Means: User-defined Contrasts


Fit: lme.formula(fixed = thisY ~ -1 + EpoTime, data = y, random = ~1 |
    Subject)

Linear Hypotheses:
                            Estimate Std. Error z value Pr(>|z|)
EpoVsPlacebo.EpoDay13 == 0   0.20542    0.05979   3.435 0.000591 ***
EpoVsPlacebo.EpoDay27 == 0   0.32784    0.06079   5.393 1.04e-07 ***
EpoVsPlacebo.EpoDay41 == 0   0.34185    0.06032   5.667 2.90e-08 ***
EpoVsPlacebo.EpoDay61 == 0   0.39138    0.06085   6.432 3.77e-10 ***
EpoVsPlacebo.postDay62 == 0  0.45753    0.06187   7.395 8.51e-13 ***
EpoVsPlacebo.postDay80 == 0  0.22375    0.05979   3.742 0.000219 ***
---
Signif. codes:  0 '***' 0.001 '**' 0.01 '*' 0.05 '.' 0.1 ' ' 1
(Adjusted p values reported -- BH method)


#############################################################
```

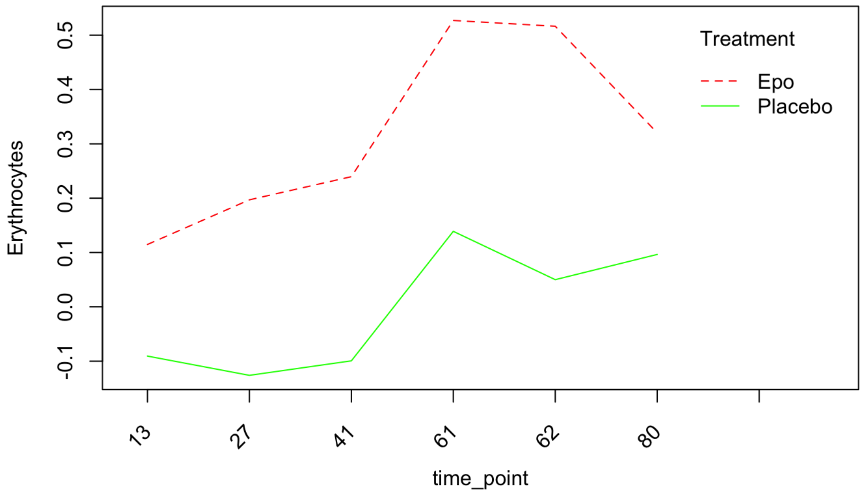


```
################################################################
########### Repeated measures analysis results for  Hematocrit ############
#################################################################
```

| | numDF <int> | denDF <dbl> | F-value <chr> | p-value <chr> |
|---|---|---|---|---|
| (Intercept) | 1 | 221 | 160.33697 | <.0001 |
| Treatment | 1 | 46 | 100.41793 | <.0001 |
| time_point | 5 | 221 | 30.58785 | <.0001 |
| Treatment:time_point | 5 | 221 | 5.44892 | 1e-04 |

4 rows

```
############ Compare Epo with Palcebo at each time point using contrasts ##########


###############################################################

####################### Raw p-values #######################


     Simultaneous Tests for General Linear Hypotheses

Multiple Comparisons of Means: User-defined Contrasts


Fit: lme.formula(fixed = thisY ~ -1 + EpoTime, data = y, random = ~1 |
    Subject)

Linear Hypotheses:
                            Estimate Std. Error z value Pr(>|z|)
EpoVsPlacebo.EpoDay13 == 0  0.023521   0.005168   4.551 5.33e-06 ***
EpoVsPlacebo.EpoDay27 == 0  0.036135   0.005260   6.870 6.42e-12 ***
EpoVsPlacebo.EpoDay41 == 0  0.037982   0.005216   7.282 3.29e-13 ***
EpoVsPlacebo.EpoDay61 == 0  0.041754   0.005265   7.931 2.22e-15 ***
EpoVsPlacebo.postDay62 == 0 0.047576   0.005359   8.877  < 2e-16 ***
EpoVsPlacebo.postDay80 == 0 0.021521   0.005168   4.164 3.12e-05 ***
---
Signif. codes:  0 ‘***’ 0.001 ‘**’ 0.01 ‘*’ 0.05 ‘.’ 0.1 ‘ ’ 1
(Adjusted p values reported -- none method)

##########################################################


############## BH adjusted p-values ####################
Benjamini, Y., and Hochberg, Y. (1995). Controlling the false discovery rate: a practical and powerful approach t
o multiple testing. Journal of the Royal Statistical Society Series B, 57, 289–300.http://www.jstor.org/stable/23
46101.
##############################################################

     Simultaneous Tests for General Linear Hypotheses

Multiple Comparisons of Means: User-defined Contrasts


Fit: lme.formula(fixed = thisY ~ -1 + EpoTime, data = y, random = ~1 |
    Subject)

Linear Hypotheses:
                            Estimate Std. Error z value Pr(>|z|)
EpoVsPlacebo.EpoDay13 == 0  0.023521   0.005168   4.551 6.40e-06 ***
EpoVsPlacebo.EpoDay27 == 0  0.036135   0.005260   6.870 9.62e-12 ***
EpoVsPlacebo.EpoDay41 == 0  0.037982   0.005216   7.282 6.59e-13 ***
EpoVsPlacebo.EpoDay61 == 0  0.041754   0.005265   7.931 6.66e-15 ***
EpoVsPlacebo.postDay62 == 0 0.047576   0.005359   8.877  < 2e-16 ***
EpoVsPlacebo.postDay80 == 0 0.021521   0.005168   4.164 3.12e-05 ***
---
Signif. codes:  0 ‘***’ 0.001 ‘**’ 0.01 ‘*’ 0.05 ‘.’ 0.1 ‘ ’ 1
(Adjusted p values reported -- BH method)


#############################################################
```

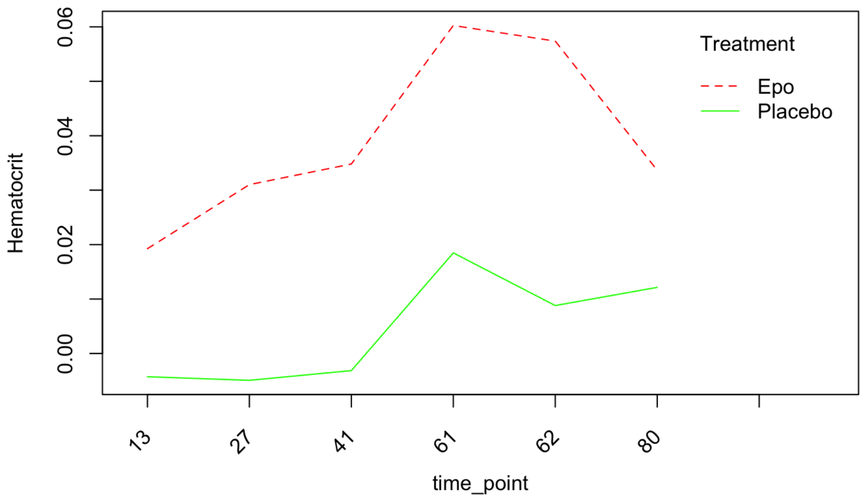


```
##############################################################

########### Repeated measures analysis results for  Hemoglobin ###########

###############################################################
```

| | numDF <int> | denDF <dbl> | F-value <chr> | p-value <chr> |
|---|---|---|---|---|
| (Intercept) | 1 | 221 | 24.81583 | <.0001 |
| Treatment | 1 | 46 | 60.47537 | <.0001 |
| time_point | 5 | 221 | 38.73160 | <.0001 |

| | numDF <int> | denDF <dbl> | F-value <chr> | p-value <chr> |
|---|---|---|---|---|
| Treatment:time_point | 5 | 221 | 3.92180 | 0.002 |

4 rows

```
########### Compare Epo with Palcebo at each time point using contrasts ##########


##############################################################

####################### Raw p-values #######################


     Simultaneous Tests for General Linear Hypotheses

Multiple Comparisons of Means: User-defined Contrasts


Fit: lme.formula(fixed = thisY ~ -1 + EpoTime, data = y, random = ~1 |
    Subject)

Linear Hypotheses:
                            Estimate Std. Error z value Pr(>|z|)
EpoVsPlacebo.EpoDay13 == 0    0.4229     0.1129   3.745 0.000181 ***
EpoVsPlacebo.EpoDay27 == 0    0.6827     0.1148   5.947 2.73e-09 ***
EpoVsPlacebo.EpoDay41 == 0    0.6953     0.1139   6.104 1.04e-09 ***
EpoVsPlacebo.EpoDay61 == 0    0.7571     0.1149   6.589 4.41e-11 ***
EpoVsPlacebo.postDay62 == 0   0.8204     0.1168   7.024 2.16e-12 ***
EpoVsPlacebo.postDay80 == 0   0.3812     0.1129   3.376 0.000736 ***
---
Signif. codes:  0 '***' 0.001 '**' 0.01 '*' 0.05 '.' 0.1 ' ' 1
(Adjusted p values reported -- none method)

#########################################################


############# BH adjusted p-values ###################
Benjamini, Y., and Hochberg, Y. (1995). Controlling the false discovery rate: a practical and powerful approach t
o multiple testing. Journal of the Royal Statistical Society Series B, 57, 289–300.http://www.jstor.org/stable/23
46101.
#############################################################

     Simultaneous Tests for General Linear Hypotheses

Multiple Comparisons of Means: User-defined Contrasts


Fit: lme.formula(fixed = thisY ~ -1 + EpoTime, data = y, random = ~1 |
    Subject)

Linear Hypotheses:
                            Estimate Std. Error z value Pr(>|z|)
EpoVsPlacebo.EpoDay13 == 0    0.4229     0.1129   3.745 0.000217 ***
EpoVsPlacebo.EpoDay27 == 0    0.6827     0.1148   5.947 4.10e-09 ***
EpoVsPlacebo.EpoDay41 == 0    0.6953     0.1139   6.104 2.07e-09 ***
EpoVsPlacebo.EpoDay61 == 0    0.7571     0.1149   6.589 1.32e-10 ***
EpoVsPlacebo.postDay62 == 0   0.8204     0.1168   7.024 1.30e-11 ***
EpoVsPlacebo.postDay80 == 0   0.3812     0.1129   3.376 0.000736 ***
---
Signif. codes:  0 '***' 0.001 '**' 0.01 '*' 0.05 '.' 0.1 ' ' 1
(Adjusted p values reported -- BH method)


#############################################################
```

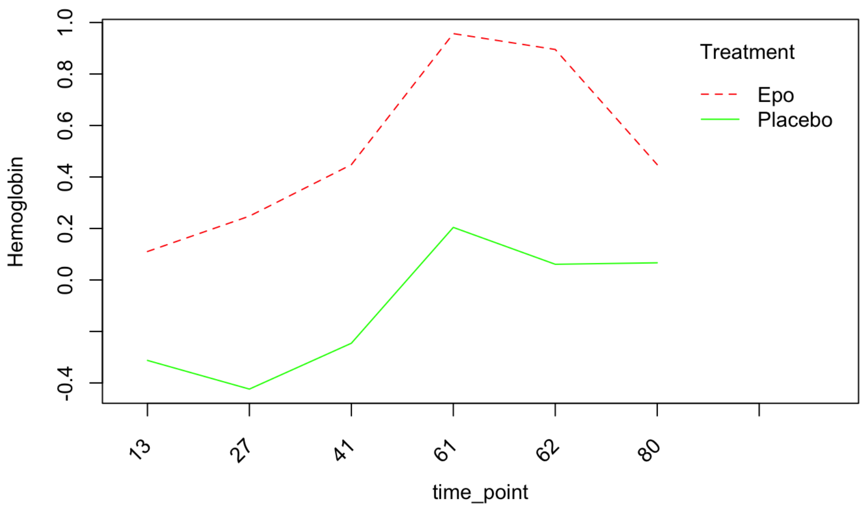


```
#############################################################

########## Repeated measures analysis results for  IRF ###########

##############################################################
```

| | numDF <int> | denDF <dbl> | F-value <chr> | p-value <chr> |
|---|---|---|---|---|
| (Intercept) | 1 | 221 | 0.049419 | 0.8243 |
| Treatment | 1 | 46 | 1.536417 | 0.2214 |
| time_point | 5 | 221 | 14.500704 | <.0001 |
| Treatment:time_point | 5 | 221 | 18.468077 | <.0001 |

4 rows

```
############ Compare Epo with Palcebo at each time point using contrasts ###########


################################################################

####################### Raw p-values #######################


     Simultaneous Tests for General Linear Hypotheses

Multiple Comparisons of Means: User-defined Contrasts


Fit: lme.formula(fixed = thisY ~ -1 + EpoTime, data = y, random = ~1 |
    Subject)

Linear Hypotheses:
                            Estimate Std. Error z value Pr(>|z|)
EpoVsPlacebo.EpoDay13 == 0    1.6333     0.8013   2.038  0.04153 *
EpoVsPlacebo.EpoDay27 == 0    0.9957     0.8117   1.227  0.21996
EpoVsPlacebo.EpoDay41 == 0    1.4390     0.8069   1.783  0.07454 .
EpoVsPlacebo.EpoDay61 == 0   -3.2588     0.8123  -4.012 6.03e-05 ***
EpoVsPlacebo.postDay62 == 0  -3.4052     0.8228  -4.139 3.49e-05 ***
EpoVsPlacebo.postDay80 == 0  -2.4167     0.8013  -3.016  0.00256 **
---
Signif. codes:  0 '***' 0.001 '**' 0.01 '*' 0.05 '.' 0.1 ' ' 1
(Adjusted p values reported -- none method)

###########################################################


############## BH adjusted p-values ####################
Benjamini, Y., and Hochberg, Y. (1995). Controlling the false discovery rate: a practical and powerful approach t
o multiple testing. Journal of the Royal Statistical Society Series B, 57, 289–300.http://www.jstor.org/stable/23
46101.
##############################################################

     Simultaneous Tests for General Linear Hypotheses

Multiple Comparisons of Means: User-defined Contrasts


Fit: lme.formula(fixed = thisY ~ -1 + EpoTime, data = y, random = ~1 |
    Subject)

Linear Hypotheses:
                            Estimate Std. Error z value Pr(>|z|)
EpoVsPlacebo.EpoDay13 == 0    1.6333     0.8013   2.038 0.062289 .
EpoVsPlacebo.EpoDay27 == 0    0.9957     0.8117   1.227 0.219965
EpoVsPlacebo.EpoDay41 == 0    1.4390     0.8069   1.783 0.089447 .
EpoVsPlacebo.EpoDay61 == 0   -3.2588     0.8123  -4.012 0.000181 ***
EpoVsPlacebo.postDay62 == 0  -3.4052     0.8228  -4.139 0.000181 ***
EpoVsPlacebo.postDay80 == 0  -2.4167     0.8013  -3.016 0.005127 **
---
Signif. codes:  0 '***' 0.001 '**' 0.01 '*' 0.05 '.' 0.1 ' ' 1
(Adjusted p values reported -- BH method)


##############################################################
```

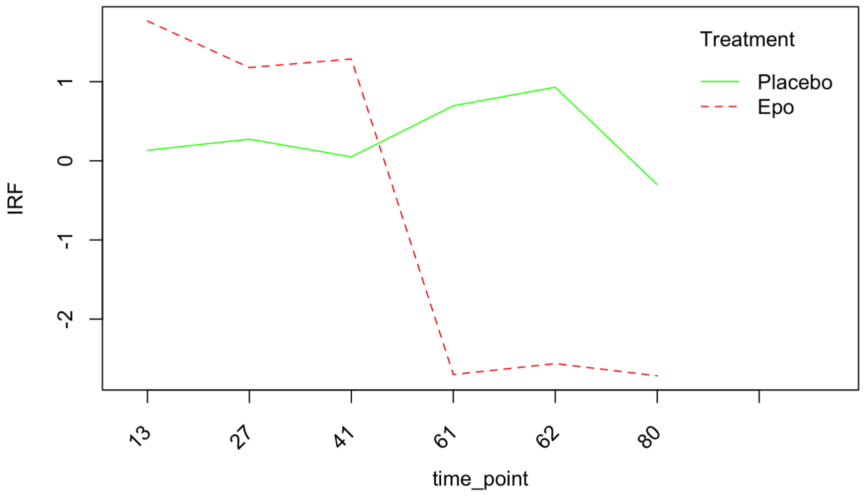

```
################################################################
########### Repeated measures analysis results for  MCH ############
################################################################
```

| | numDF <int> | denDF <dbl> | F-value <chr> | p-value <chr> |
|---|---|---|---|---|
| (Intercept) | 1 | 220 | 24.770706 | <.0001 |
| Treatment | 1 | 46 | 0.196088 | 0.6600 |
| time_point | 5 | 220 | 15.091237 | <.0001 |
| Treatment:time_point | 5 | 220 | 2.595238 | 0.0264 |

4 rows

```
############ Compare Epo with Palcebo at each time point using contrasts ###########


################################################################
######################## Raw p-values ########################


     Simultaneous Tests for General Linear Hypotheses

Multiple Comparisons of Means: User-defined Contrasts


Fit: lme.formula(fixed = thisY ~ -1 + EpoTime, data = y, random = ~1 |
    Subject)

Linear Hypotheses:
                             Estimate Std. Error z value Pr(>|z|)
EpoVsPlacebo.EpoDay13 == 0   0.008750   0.008861   0.987    0.323
EpoVsPlacebo.EpoDay27 == 0   0.013783   0.008970   1.537    0.124
EpoVsPlacebo.EpoDay41 == 0   0.010579   0.008924   1.186    0.236
EpoVsPlacebo.EpoDay61 == 0   0.003309   0.008975   0.369    0.712
EpoVsPlacebo.postDay62 == 0 -0.009398   0.009075  -1.036    0.300
EpoVsPlacebo.postDay80 == 0 -0.006019   0.008924  -0.674    0.500
(Adjusted p values reported -- none method)

##########################################################


############## BH adjusted p-values ####################
Benjamini, Y., and Hochberg, Y. (1995). Controlling the false discovery rate: a practical and powerful approach t
o multiple testing. Journal of the Royal Statistical Society Series B, 57, 289–300.http://www.jstor.org/stable/23
46101.
##############################################################

     Simultaneous Tests for General Linear Hypotheses

Multiple Comparisons of Means: User-defined Contrasts


Fit: lme.formula(fixed = thisY ~ -1 + EpoTime, data = y, random = ~1 |
    Subject)

Linear Hypotheses:
                             Estimate Std. Error z value Pr(>|z|)
EpoVsPlacebo.EpoDay13 == 0   0.008750   0.008861   0.987    0.485
EpoVsPlacebo.EpoDay27 == 0   0.013783   0.008970   1.537    0.485
EpoVsPlacebo.EpoDay41 == 0   0.010579   0.008924   1.186    0.485
EpoVsPlacebo.EpoDay61 == 0   0.003309   0.008975   0.369    0.712
EpoVsPlacebo.postDay62 == 0 -0.009398   0.009075  -1.036    0.485
EpoVsPlacebo.postDay80 == 0 -0.006019   0.008924  -0.674    0.600
(Adjusted p values reported -- BH method)


##############################################################
```

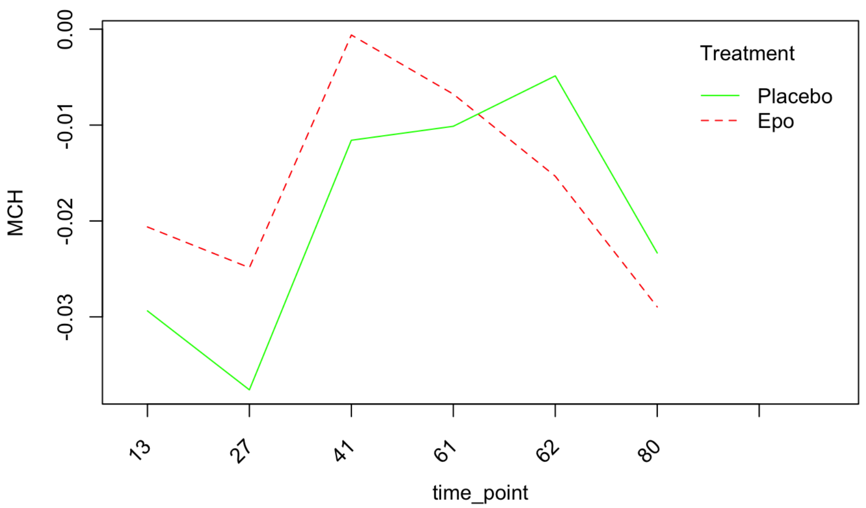

```
##############################################################

########### Repeated measures analysis results for  MCHC ############

###############################################################
```

| | numDF <int> | denDF <dbl> | F-value <chr> | p-value <chr> |
|---|---|---|---|---|
| (Intercept) | 1 | 221 | 225.14362 | <.0001 |
| Treatment | 1 | 46 | 5.45011 | 0.0240 |
| time_point | 5 | 221 | 10.71895 | <.0001 |
| Treatment:time_point | 5 | 221 | 0.94348 | 0.4537 |

4 rows

```
############ Compare Epo with Palcebo at each time point using contrasts ##########


###############################################################

####################### Raw p-values ########################


	 Simultaneous Tests for General Linear Hypotheses

Multiple Comparisons of Means: User-defined Contrasts


Fit: lme.formula(fixed = thisY ~ -1 + EpoTime, data = y, random = ~1 | 
    Subject)

Linear Hypotheses:
                            Estimate Std. Error z value Pr(>|z|)   
EpoVsPlacebo.EpoDay13 == 0   -0.1042     0.1017  -1.024  0.30580   
EpoVsPlacebo.EpoDay27 == 0   -0.1126     0.1033  -1.089  0.27598   
EpoVsPlacebo.EpoDay41 == 0   -0.1607     0.1026  -1.567  0.11710   
EpoVsPlacebo.EpoDay61 == 0   -0.1896     0.1034  -1.833  0.06677 . 
EpoVsPlacebo.postDay62 == 0  -0.3171     0.1051  -3.018  0.00255 **
EpoVsPlacebo.postDay80 == 0  -0.1250     0.1017  -1.229  0.21911   
---
Signif. codes:  0 ‘***’ 0.001 ‘**’ 0.01 ‘*’ 0.05 ‘.’ 0.1 ‘ ’ 1
(Adjusted p values reported -- none method)

##########################################################


############# BH adjusted p-values ####################
Benjamini, Y., and Hochberg, Y. (1995). Controlling the false discovery rate: a practical and powerful approach t
o multiple testing. Journal of the Royal Statistical Society Series B, 57, 289–300.http://www.jstor.org/stable/23
46101.
##############################################################

	 Simultaneous Tests for General Linear Hypotheses

Multiple Comparisons of Means: User-defined Contrasts


Fit: lme.formula(fixed = thisY ~ -1 + EpoTime, data = y, random = ~1 | 
    Subject)

Linear Hypotheses:
                            Estimate Std. Error z value Pr(>|z|)  
EpoVsPlacebo.EpoDay13 == 0   -0.1042     0.1017  -1.024   0.3058  
EpoVsPlacebo.EpoDay27 == 0   -0.1126     0.1033  -1.089   0.3058  
EpoVsPlacebo.EpoDay41 == 0   -0.1607     0.1026  -1.567   0.2342  
EpoVsPlacebo.EpoDay61 == 0   -0.1896     0.1034  -1.833   0.2003  
EpoVsPlacebo.postDay62 == 0  -0.3171     0.1051  -3.018   0.0153 *
EpoVsPlacebo.postDay80 == 0  -0.1250     0.1017  -1.229   0.3058  
---
Signif. codes:  0 ‘***’ 0.001 ‘**’ 0.01 ‘*’ 0.05 ‘.’ 0.1 ‘ ’ 1
(Adjusted p values reported -- BH method)


#############################################################
```

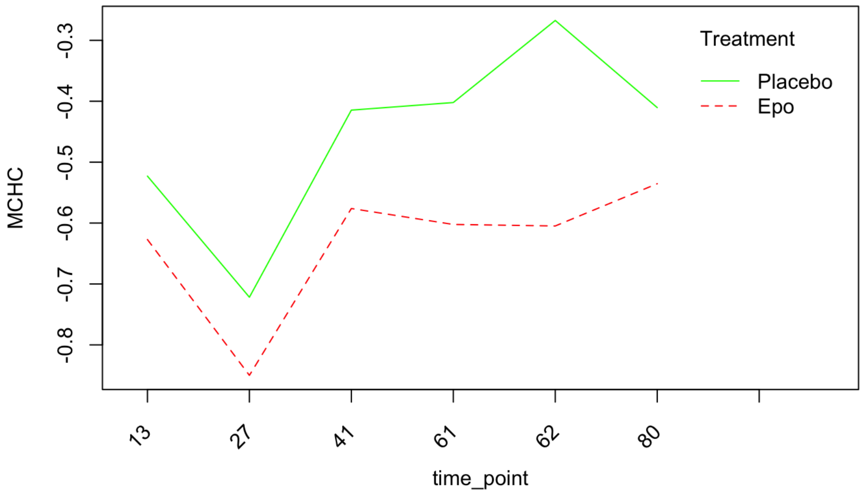


```
################################################################
########### Repeated measures analysis results for  MCV ############
#################################################################
```

| | numDF <int> | denDF <dbl> | F-value <chr> | p-value <chr> |
|---|---|---|---|---|
| (Intercept) | 1 | 221 | 105.96797 | <.0001 |
| Treatment | 1 | 46 | 11.91240 | 0.0012 |
| time_point | 5 | 221 | 11.83883 | <.0001 |
| Treatment:time_point | 5 | 221 | 1.87568 | 0.0996 |

4 rows

```
############ Compare Epo with Palcebo at each time point using contrasts ##########


###############################################################

####################### Raw p-values #######################


     Simultaneous Tests for General Linear Hypotheses

Multiple Comparisons of Means: User-defined Contrasts


Fit: lme.formula(fixed = thisY ~ -1 + EpoTime, data = y, random = ~1 |
    Subject)

Linear Hypotheses:
                            Estimate Std. Error z value Pr(>|z|)
EpoVsPlacebo.EpoDay13 == 0    1.0375     0.3742   2.773 0.005559 **
EpoVsPlacebo.EpoDay27 == 0    1.2458     0.3789   3.288 0.001010 **
EpoVsPlacebo.EpoDay41 == 0    1.3542     0.3767   3.595 0.000325 ***
EpoVsPlacebo.EpoDay61 == 0    1.1483     0.3792   3.028 0.002462 **
EpoVsPlacebo.postDay62 == 0   1.0576     0.3840   2.754 0.005886 **
EpoVsPlacebo.postDay80 == 0   0.3542     0.3742   0.947 0.343884
---
Signif. codes:  0 '***' 0.001 '**' 0.01 '*' 0.05 '.' 0.1 ' ' 1
(Adjusted p values reported -- none method)

##########################################################


############## BH adjusted p-values ####################
Benjamini, Y., and Hochberg, Y. (1995). Controlling the false discovery rate: a practical and powerful approach t
o multiple testing. Journal of the Royal Statistical Society Series B, 57, 289–300.http://www.jstor.org/stable/23
46101.
##############################################################

     Simultaneous Tests for General Linear Hypotheses

Multiple Comparisons of Means: User-defined Contrasts


Fit: lme.formula(fixed = thisY ~ -1 + EpoTime, data = y, random = ~1 |
    Subject)

Linear Hypotheses:
                            Estimate Std. Error z value Pr(>|z|)
EpoVsPlacebo.EpoDay13 == 0    1.0375     0.3742   2.773  0.00706 **
EpoVsPlacebo.EpoDay27 == 0    1.2458     0.3789   3.288  0.00303 **
EpoVsPlacebo.EpoDay41 == 0    1.3542     0.3767   3.595  0.00195 **
EpoVsPlacebo.EpoDay61 == 0    1.1483     0.3792   3.028  0.00492 **
EpoVsPlacebo.postDay62 == 0   1.0576     0.3840   2.754  0.00706 **
EpoVsPlacebo.postDay80 == 0   0.3542     0.3742   0.947  0.34388
---
Signif. codes:  0 '***' 0.001 '**' 0.01 '*' 0.05 '.' 0.1 ' ' 1
(Adjusted p values reported -- BH method)


##############################################################
```

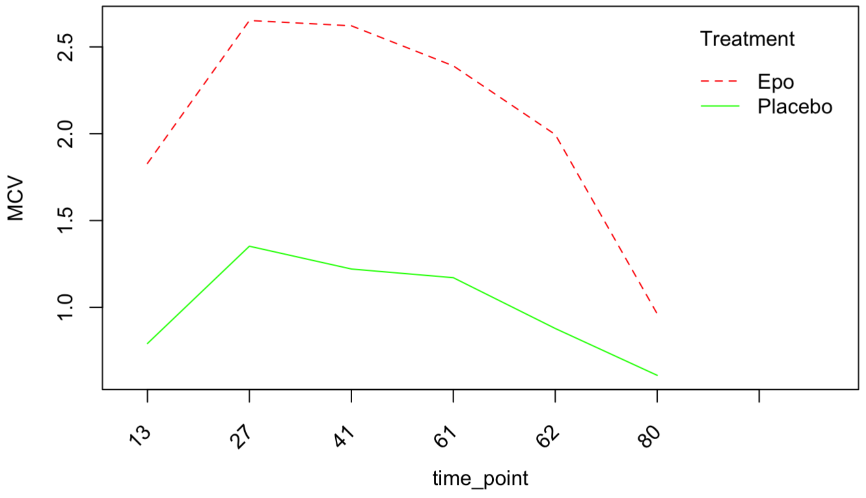


```
##############################################################

########### Repeated measures analysis results for  RDW_SD ###########

###############################################################
```

| | numDF <int> | denDF <dbl> | F-value <chr> | p-value <chr> |
|---|---|---|---|---|
| (Intercept) | 1 | 219 | 32.04582 | <.0001 |
| Treatment | 1 | 46 | 17.70673 | 1e-04 |
| time_point | 5 | 219 | 30.09056 | <.0001 |

| | numDF <int> | denDF <dbl> | F-value <chr> | p-value <chr> |
|---|---|---|---|---|
| Treatment:time_point | 5 | 219 | 9.09271 | <.0001 |

4 rows

```
############ Compare Epo with Palcebo at each time point using contrasts ###########


################################################################

######################## Raw p-values ########################


     Simultaneous Tests for General Linear Hypotheses

Multiple Comparisons of Means: User-defined Contrasts


Fit: lme.formula(fixed = thisY ~ -1 + EpoTime, data = y, random = ~1 |
    Subject)

Linear Hypotheses:
                            Estimate Std. Error z value Pr(>|z|)
EpoVsPlacebo.EpoDay13 == 0    2.2750     0.4841   4.700 2.61e-06 ***
EpoVsPlacebo.EpoDay27 == 0    2.1992     0.4925   4.466 7.97e-06 ***
EpoVsPlacebo.EpoDay41 == 0    2.1981     0.4874   4.510 6.49e-06 ***
EpoVsPlacebo.EpoDay61 == 0    1.7946     0.4901   3.661 0.000251 ***
EpoVsPlacebo.postDay62 == 0   1.4846     0.4954   2.997 0.002728 **
EpoVsPlacebo.postDay80 == 0  -0.1585     0.4874  -0.325 0.745109
---
Signif. codes:  0 '***' 0.001 '**' 0.01 '*' 0.05 '.' 0.1 ' ' 1
(Adjusted p values reported -- none method)

##########################################################


############## BH adjusted p-values ####################
Benjamini, Y., and Hochberg, Y. (1995). Controlling the false discovery rate: a practical and powerful approach t
o multiple testing. Journal of the Royal Statistical Society Series B, 57, 289–300.http://www.jstor.org/stable/23
46101.
##############################################################

     Simultaneous Tests for General Linear Hypotheses

Multiple Comparisons of Means: User-defined Contrasts


Fit: lme.formula(fixed = thisY ~ -1 + EpoTime, data = y, random = ~1 |
    Subject)

Linear Hypotheses:
                            Estimate Std. Error z value Pr(>|z|)
EpoVsPlacebo.EpoDay13 == 0    2.2750     0.4841   4.700 1.56e-05 ***
EpoVsPlacebo.EpoDay27 == 0    2.1992     0.4925   4.466 1.59e-05 ***
EpoVsPlacebo.EpoDay41 == 0    2.1981     0.4874   4.510 1.59e-05 ***
EpoVsPlacebo.EpoDay61 == 0    1.7946     0.4901   3.661 0.000376 ***
EpoVsPlacebo.postDay62 == 0   1.4846     0.4954   2.997 0.003274 **
EpoVsPlacebo.postDay80 == 0  -0.1585     0.4874  -0.325 0.745109
---
Signif. codes:  0 '***' 0.001 '**' 0.01 '*' 0.05 '.' 0.1 ' ' 1
(Adjusted p values reported -- BH method)


##############################################################
```

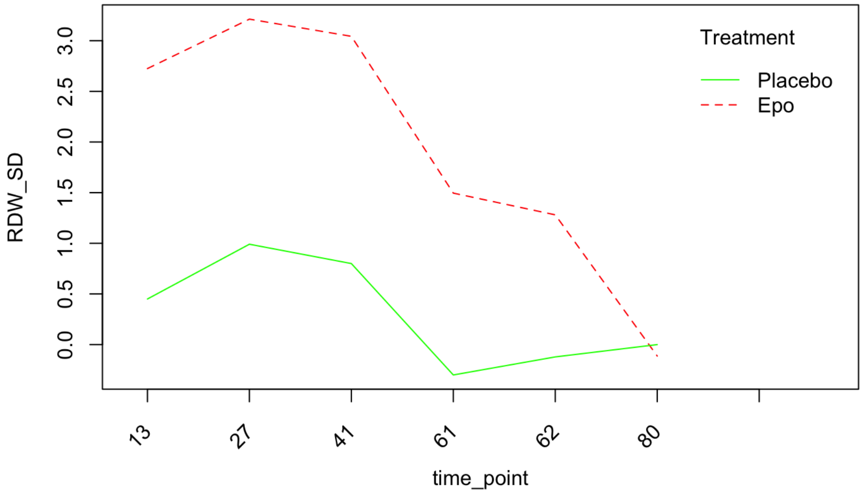


```
##############################################################

########### Repeated measures analysis results for  Reticulo_per ############

###############################################################
```

| | numDF <int> | denDF <dbl> | F-value <chr> | p-value <chr> |
|---|---|---|---|---|
| (Intercept) | 1 | 220 | 0.004686 | 0.9455 |
| Treatment | 1 | 46 | 0.083848 | 0.7735 |
| time_point | 5 | 220 | 24.837678 | <.0001 |
| Treatment:time_point | 5 | 220 | 30.711048 | <.0001 |

4 rows

```
############ Compare Epo with Palcebo at each time point using contrasts ###########


################################################################

####################### Raw p-values #######################


     Simultaneous Tests for General Linear Hypotheses

Multiple Comparisons of Means: User-defined Contrasts


Fit: lme.formula(fixed = thisY ~ -1 + EpoTime, data = y, random = ~1 |
    Subject)

Linear Hypotheses:
                            Estimate Std. Error z value Pr(>|z|)
EpoVsPlacebo.EpoDay13 == 0   0.40417    0.07003   5.772 7.85e-09 ***
EpoVsPlacebo.EpoDay27 == 0   0.22739    0.07179   3.167 0.001538 **
EpoVsPlacebo.EpoDay41 == 0   0.23532    0.07064   3.331 0.000865 ***
EpoVsPlacebo.EpoDay61 == 0  -0.27589    0.07126  -3.872 0.000108 ***
EpoVsPlacebo.postDay62 == 0 -0.29188    0.07245  -4.029 5.61e-05 ***
EpoVsPlacebo.postDay80 == 0 -0.26042    0.07003  -3.719 0.000200 ***
---
Signif. codes:  0 '***' 0.001 '**' 0.01 '*' 0.05 '.' 0.1 ' ' 1
(Adjusted p values reported -- none method)

##########################################################


############## BH adjusted p-values ####################
Benjamini, Y., and Hochberg, Y. (1995). Controlling the false discovery rate: a practical and powerful approach t
o multiple testing. Journal of the Royal Statistical Society Series B, 57, 289–300.http://www.jstor.org/stable/23
46101.
##############################################################

     Simultaneous Tests for General Linear Hypotheses

Multiple Comparisons of Means: User-defined Contrasts


Fit: lme.formula(fixed = thisY ~ -1 + EpoTime, data = y, random = ~1 |
    Subject)

Linear Hypotheses:
                            Estimate Std. Error z value Pr(>|z|)
EpoVsPlacebo.EpoDay13 == 0   0.40417    0.07003   5.772 4.71e-08 ***
EpoVsPlacebo.EpoDay27 == 0   0.22739    0.07179   3.167 0.001538 **
EpoVsPlacebo.EpoDay41 == 0   0.23532    0.07064   3.331 0.001038 **
EpoVsPlacebo.EpoDay61 == 0  -0.27589    0.07126  -3.872 0.000216 ***
EpoVsPlacebo.postDay62 == 0 -0.29188    0.07245  -4.029 0.000168 ***
EpoVsPlacebo.postDay80 == 0 -0.26042    0.07003  -3.719 0.000300 ***
---
Signif. codes:  0 '***' 0.001 '**' 0.01 '*' 0.05 '.' 0.1 ' ' 1
(Adjusted p values reported -- BH method)


##############################################################
```

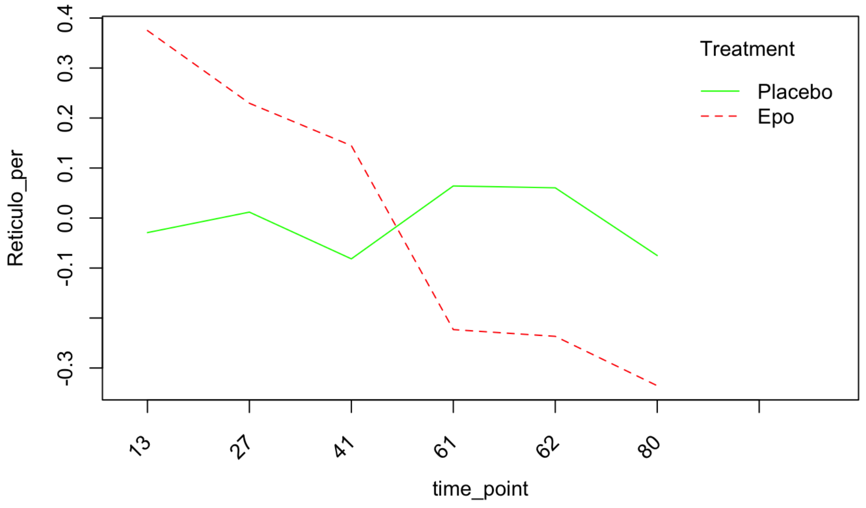

```
################################################################

########### Repeated measures analysis results for  Reticulocyte ############

################################################################
```

| | numDF <int> | denDF <dbl> | F-value <chr> | p-value <chr> |
|---|---|---|---|---|
| (Intercept) | 1 | 221 | 2.294845 | 0.1312 |
| Treatment | 1 | 46 | 1.017214 | 0.3185 |
| time_point | 5 | 221 | 17.398805 | <.0001 |
| Treatment:time_point | 5 | 221 | 28.143622 | <.0001 |

4 rows

```
############ Compare Epo with Palcebo at each time point using contrasts ##########


################################################################

######################## Raw p-values ########################


     Simultaneous Tests for General Linear Hypotheses

Multiple Comparisons of Means: User-defined Contrasts


Fit: lme.formula(fixed = thisY ~ -1 + EpoTime, data = y, random = ~1 |
    Subject)

Linear Hypotheses:
                            Estimate Std. Error z value Pr(>|z|)
EpoVsPlacebo.EpoDay13 == 0    21.098      3.398   6.209 5.33e-10 ***
EpoVsPlacebo.EpoDay27 == 0    14.097      3.462   4.071 4.67e-05 ***
EpoVsPlacebo.EpoDay41 == 0    13.324      3.431   3.883 0.000103 ***
EpoVsPlacebo.EpoDay61 == 0   -12.174      3.466  -3.513 0.000444 ***
EpoVsPlacebo.postDay62 == 0  -12.491      3.533  -3.535 0.000407 ***
EpoVsPlacebo.postDay80 == 0  -13.052      3.398  -3.841 0.000122 ***
---
Signif. codes:  0 '***' 0.001 '**' 0.01 '*' 0.05 '.' 0.1 ' ' 1
(Adjusted p values reported -- none method)

##########################################################


############## BH adjusted p-values ####################
Benjamini, Y., and Hochberg, Y. (1995). Controlling the false discovery rate: a practical and powerful approach t
o multiple testing. Journal of the Royal Statistical Society Series B, 57, 289–300.http://www.jstor.org/stable/23
46101.
###############################################################

     Simultaneous Tests for General Linear Hypotheses

Multiple Comparisons of Means: User-defined Contrasts


Fit: lme.formula(fixed = thisY ~ -1 + EpoTime, data = y, random = ~1 |
    Subject)

Linear Hypotheses:
                            Estimate Std. Error z value Pr(>|z|)
EpoVsPlacebo.EpoDay13 == 0    21.098      3.398   6.209  3.2e-09 ***
EpoVsPlacebo.EpoDay27 == 0    14.097      3.462   4.071 0.000140 ***
EpoVsPlacebo.EpoDay41 == 0    13.324      3.431   3.883 0.000184 ***
EpoVsPlacebo.EpoDay61 == 0   -12.174      3.466  -3.513 0.000444 ***
EpoVsPlacebo.postDay62 == 0  -12.491      3.533  -3.535 0.000444 ***
EpoVsPlacebo.postDay80 == 0  -13.052      3.398  -3.841 0.000184 ***
---
Signif. codes:  0 '***' 0.001 '**' 0.01 '*' 0.05 '.' 0.1 ' ' 1
(Adjusted p values reported -- BH method)


##############################################################
```

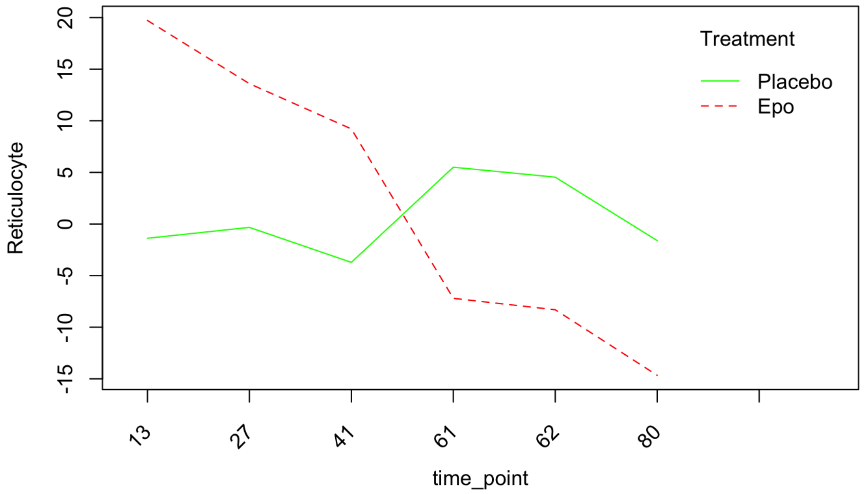


```
################################################################

########### Repeated measures analysis results for  Thrombocytes ############

################################################################
```

| | numDF <int> | denDF <dbl> | F-value <chr> | p-value <chr> |
|---|---|---|---|---|
| (Intercept) | 1 | 221 | 8.443376 | 0.0040 |
| Treatment | 1 | 46 | 0.000478 | 0.9827 |
| time_point | 5 | 221 | 6.081569 | <.0001 |
| Treatment:time_point | 5 | 221 | 2.263850 | 0.0492 |

4 rows

```
############ Compare Epo with Palcebo at each time point using contrasts ###########


################################################################

######################## Raw p-values ########################


     Simultaneous Tests for General Linear Hypotheses

Multiple Comparisons of Means: User-defined Contrasts


Fit: lme.formula(fixed = thisY ~ -1 + EpoTime, data = y, random = ~1 |
    Subject)

Linear Hypotheses:
                            Estimate Std. Error z value Pr(>|z|)
EpoVsPlacebo.EpoDay13 == 0     8.687      7.673   1.132    0.258
EpoVsPlacebo.EpoDay27 == 0    12.476      7.814   1.597    0.110
EpoVsPlacebo.EpoDay41 == 0     4.498      7.746   0.581    0.561
EpoVsPlacebo.EpoDay61 == 0   -11.817      7.821  -1.511    0.131
EpoVsPlacebo.postDay62 == 0   -9.326      7.968  -1.170    0.242
EpoVsPlacebo.postDay80 == 0   -5.396      7.673  -0.703    0.482
(Adjusted p values reported -- none method)

##########################################################


############## BH adjusted p-values ####################
Benjamini, Y., and Hochberg, Y. (1995). Controlling the false discovery rate: a practical and powerful approach t
o multiple testing. Journal of the Royal Statistical Society Series B, 57, 289–300.http://www.jstor.org/stable/23
46101.
###############################################################

     Simultaneous Tests for General Linear Hypotheses

Multiple Comparisons of Means: User-defined Contrasts


Fit: lme.formula(fixed = thisY ~ -1 + EpoTime, data = y, random = ~1 |
    Subject)

Linear Hypotheses:
                            Estimate Std. Error z value Pr(>|z|)
EpoVsPlacebo.EpoDay13 == 0     8.687      7.673   1.132    0.386
EpoVsPlacebo.EpoDay27 == 0    12.476      7.814   1.597    0.386
EpoVsPlacebo.EpoDay41 == 0     4.498      7.746   0.581    0.561
EpoVsPlacebo.EpoDay61 == 0   -11.817      7.821  -1.511    0.386
EpoVsPlacebo.postDay62 == 0   -9.326      7.968  -1.170    0.386
EpoVsPlacebo.postDay80 == 0   -5.396      7.673  -0.703    0.561
(Adjusted p values reported -- BH method)
```

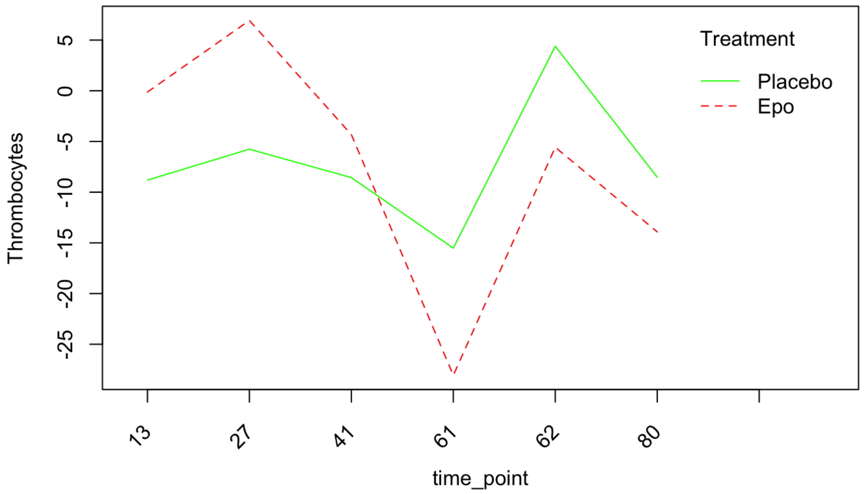


Data visualization

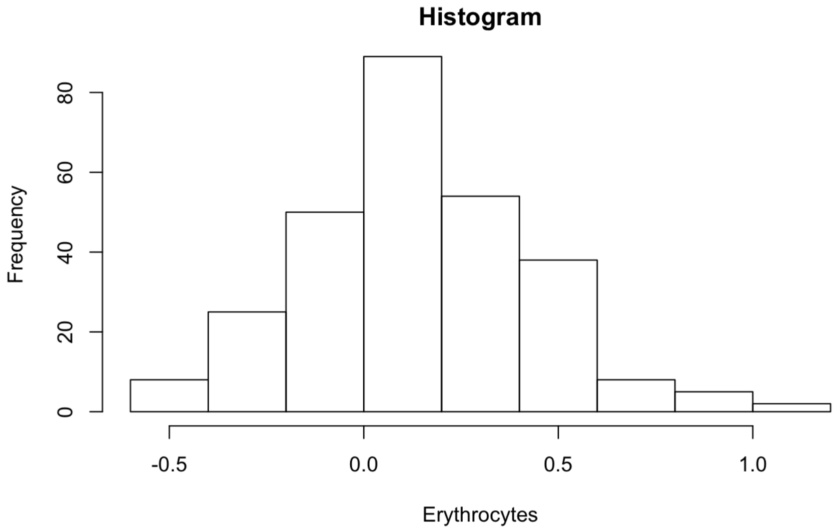


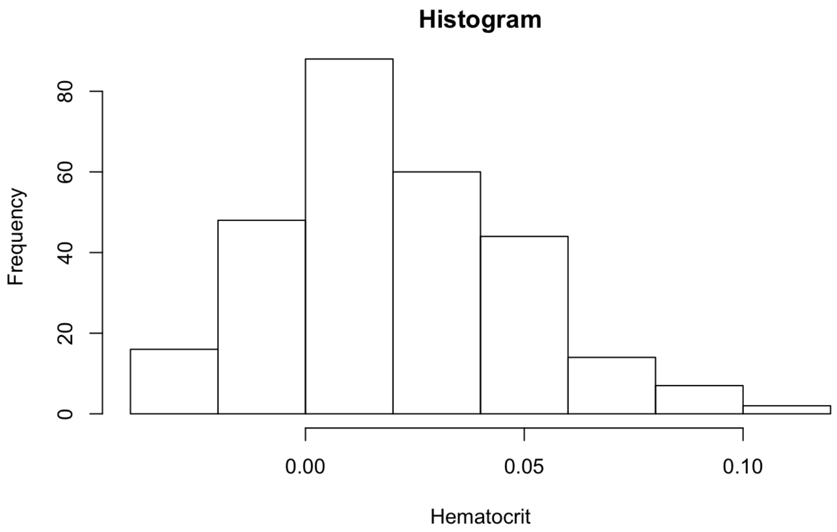

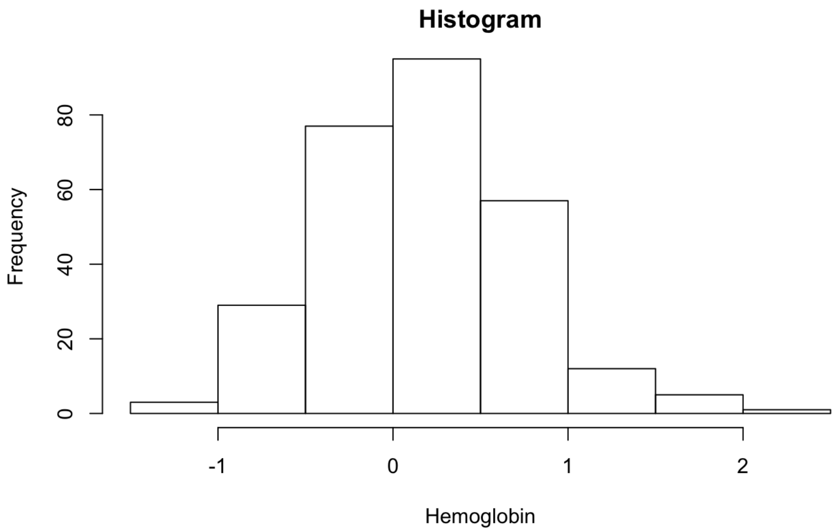
Histogram
Frequency
0
20
40
60
80
-1
0
1
2
Hemoglobin

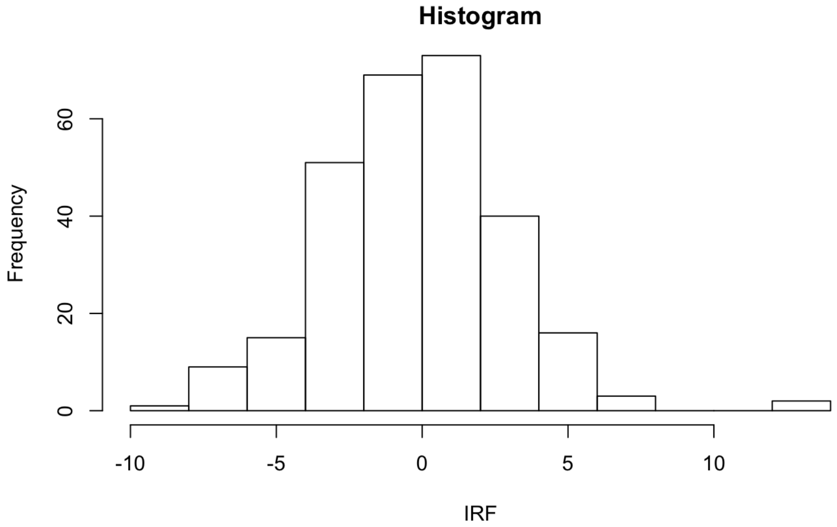
Histogram
Frequency
0
20
40
60
-10
-5
0
5
10
IRF

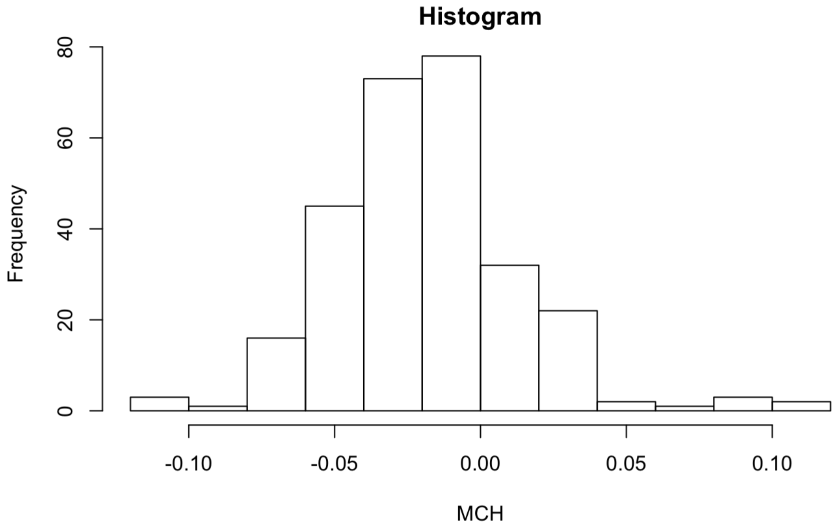
Histogram
Frequency
0
20
40
60
80
-0.10
-0.05
0.00
0.05
0.10
MCH

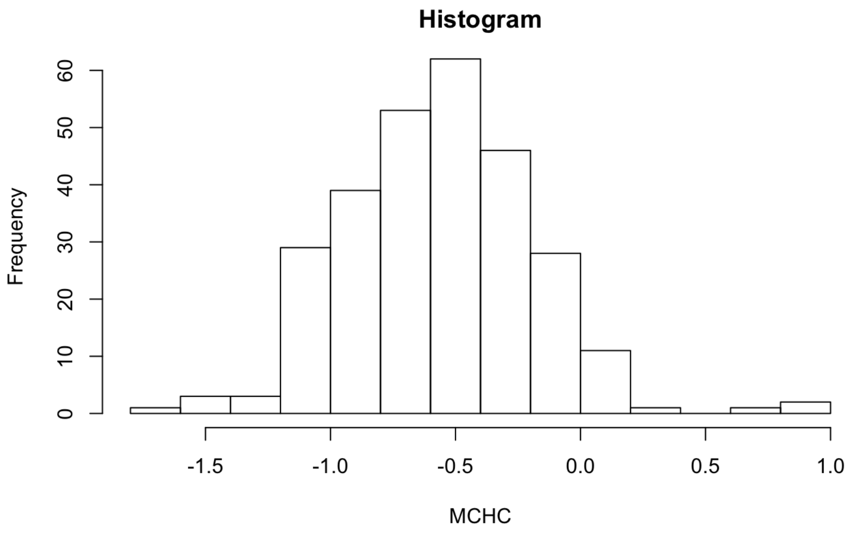
Histogram
Frequency
0
10
20
30
40
50
60
-1.5
-1.0
-0.5
0.0
0.5
1.0
MCHC

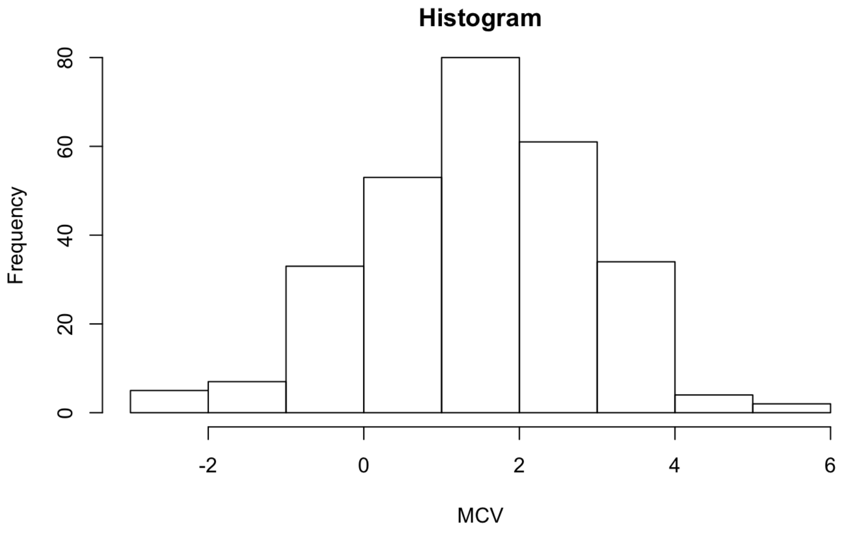
Histogram
Frequency
0
20
40
60
80
-2
0
2
4
6
MCV

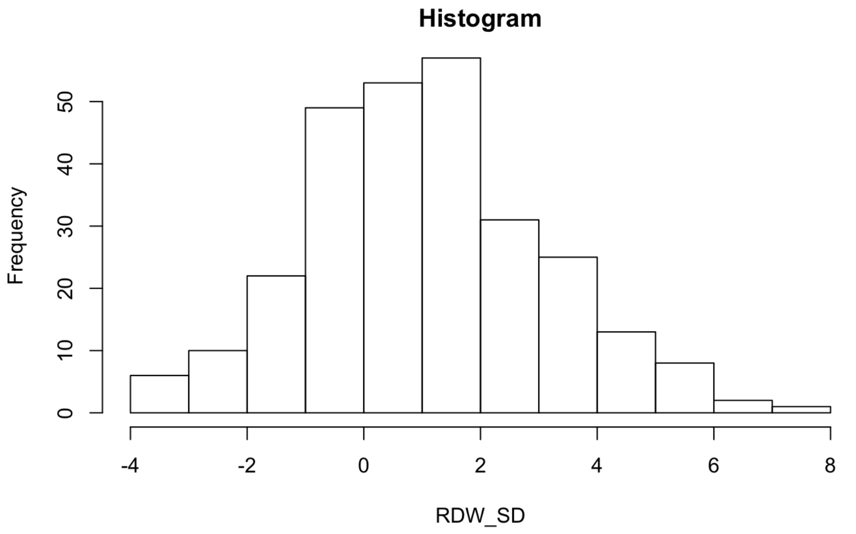
Histogram
Frequency
0
10
20
30
40
50
-4
-2
0
2
4
6
8
RDW_SD

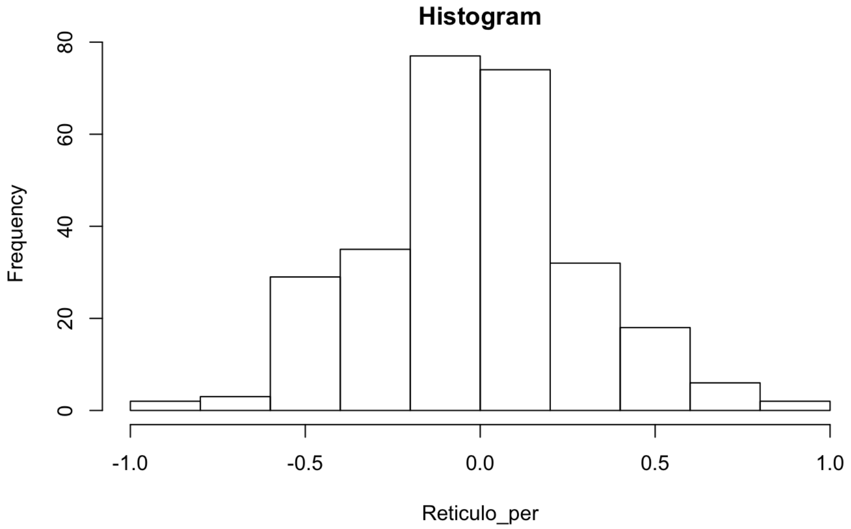
Histogram
Frequency
0
20
40
60
80
-1.0
-0.5
0.0
0.5
1.0
Reticulo_per

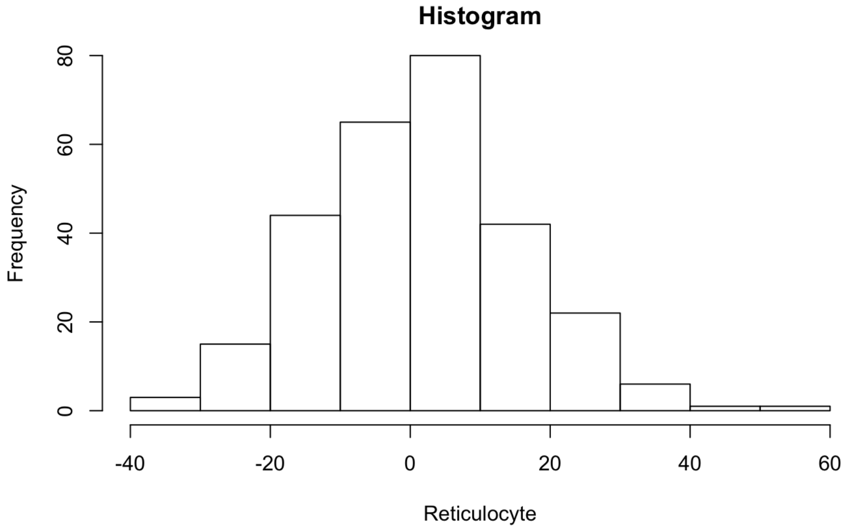
Histogram
Frequency
0
20
40
60
80
-40
-20
0
20
40
60
Reticulocyte

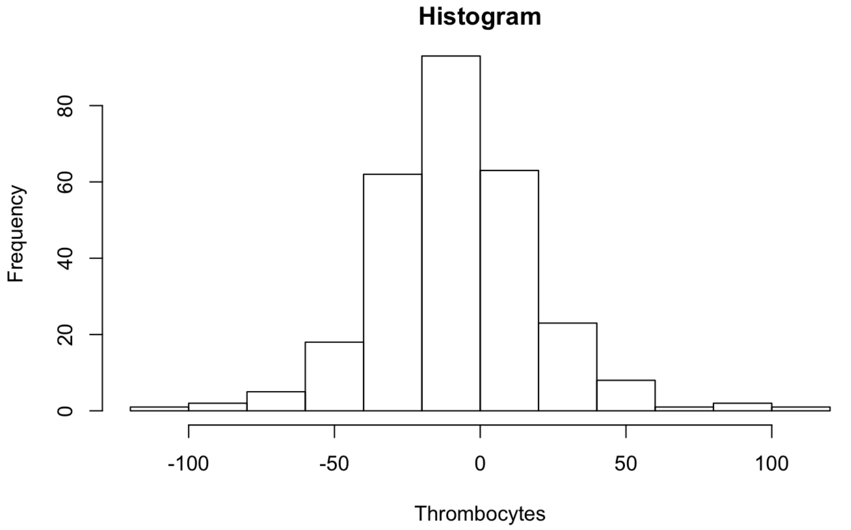
Histogram
Frequency
0
20
40
60
80
-100
-50
0
50
100
Thrombocytes

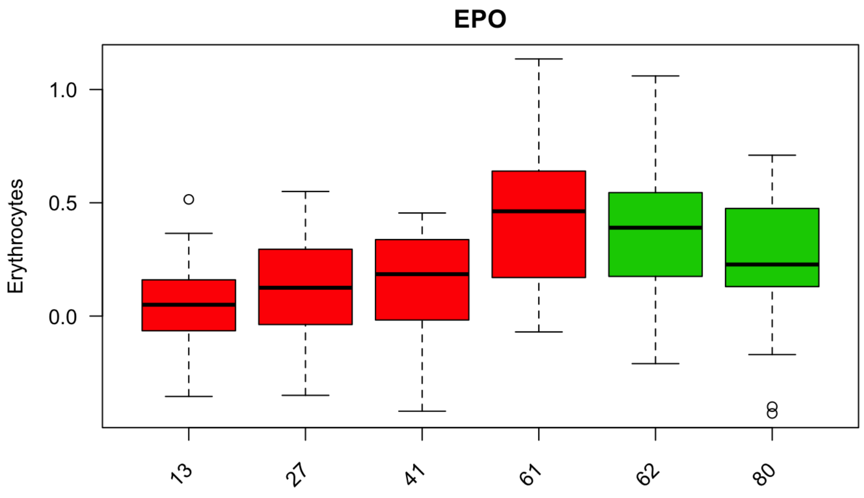
EPO
Erythrocytes
1.0
0.5
0.0
13
27
41
61
62
80

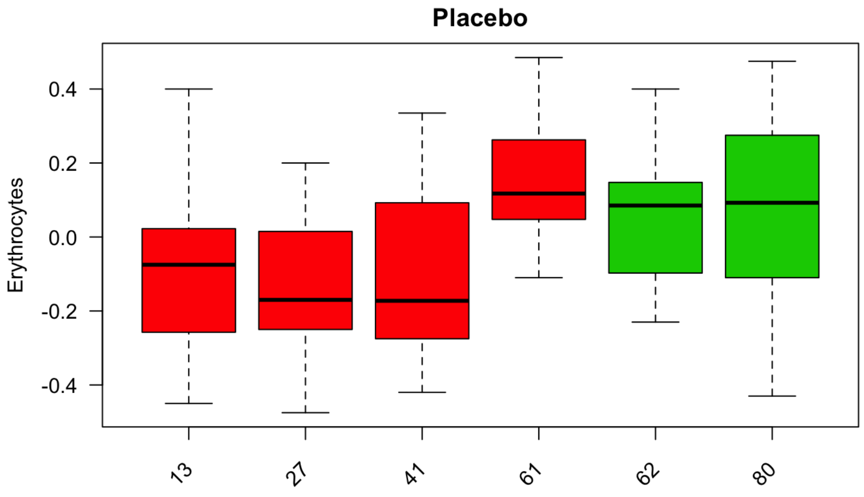
Placebo
Erythrocytes
0.4
0.2
0.0
-0.2
-0.4
13
27
41
61
62
80

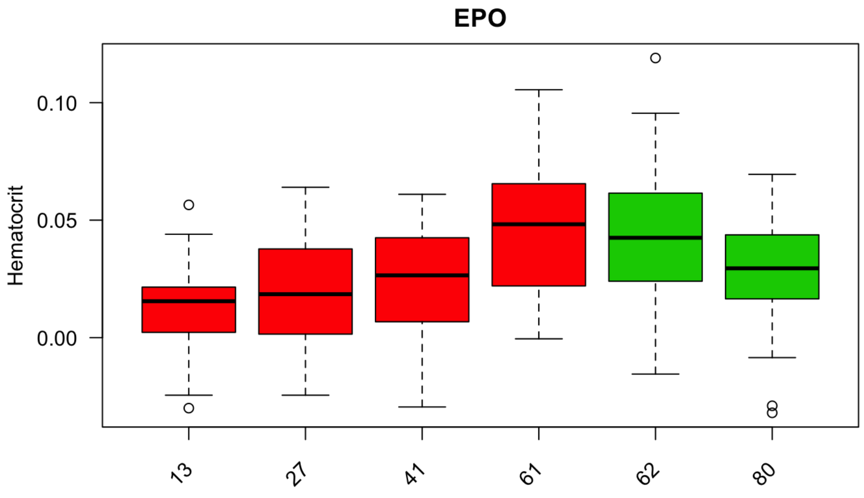
EPO
Hematocrit
0.10
0.05
0.00
13
27
41
61
62
80

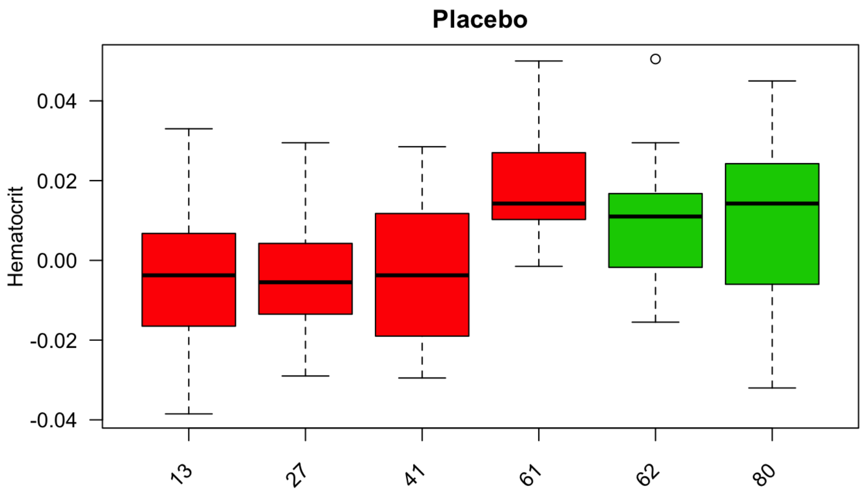

Placebo
Hematocrit
0.04
0.02
0.00
-0.02
-0.04
13
27
41
61
62
80


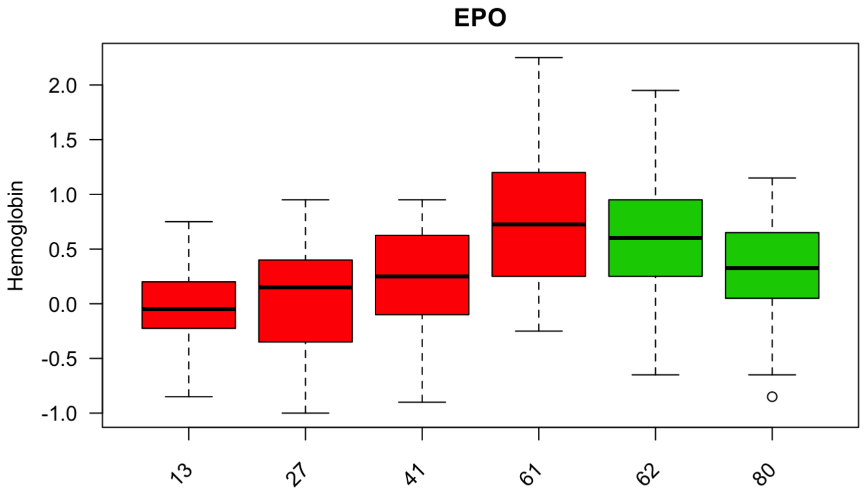

EPO
Hemoglobin
2.0
1.5
1.0
0.5
0.0
-0.5
-1.0
13
27
41
61
62
80


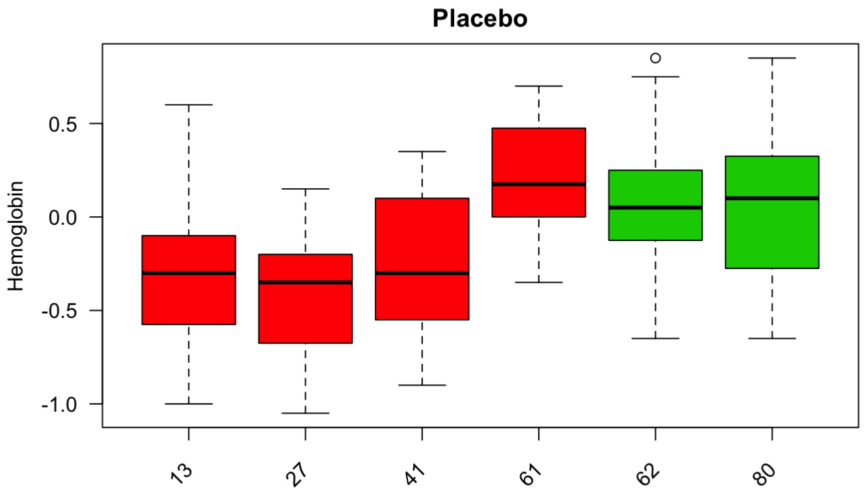

Placebo
Hemoglobin
0.5
0.0
-0.5
-1.0
13
27
41
61
62
80

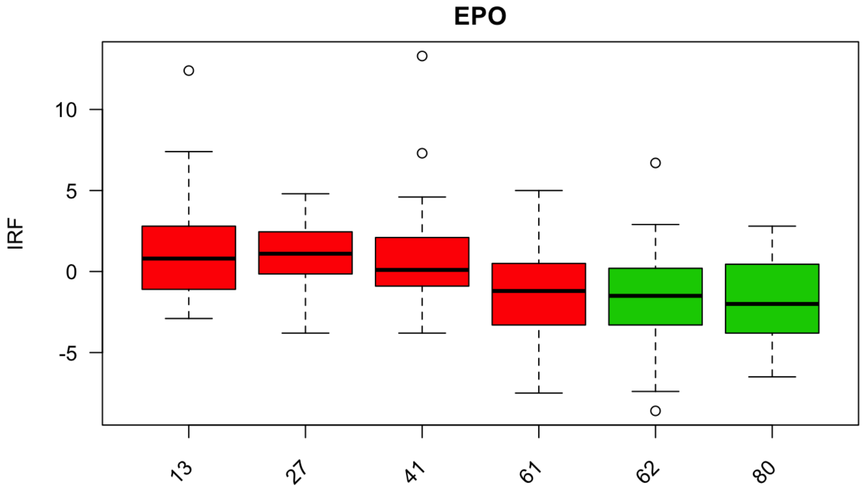
EPO
IRF
10
5
0
-5
13
27
41
61
62
80

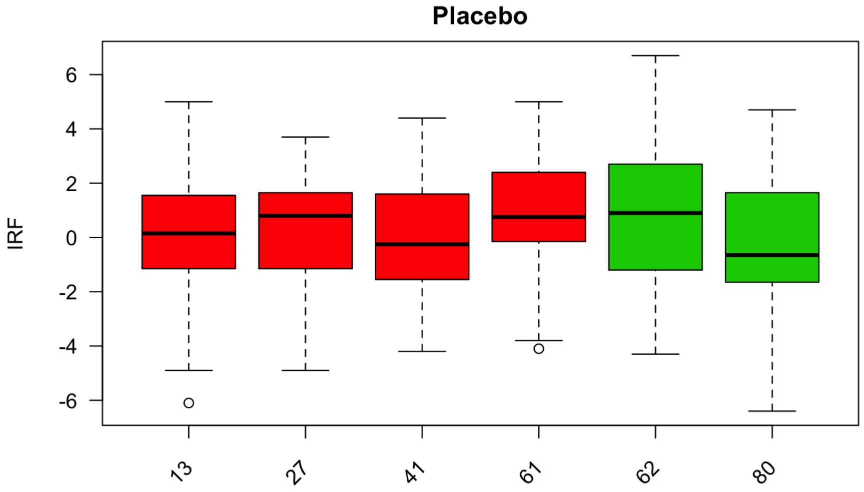
Placebo
IRF
6
4
2
0
-2
-4
-6
13
27
41
61
62
80

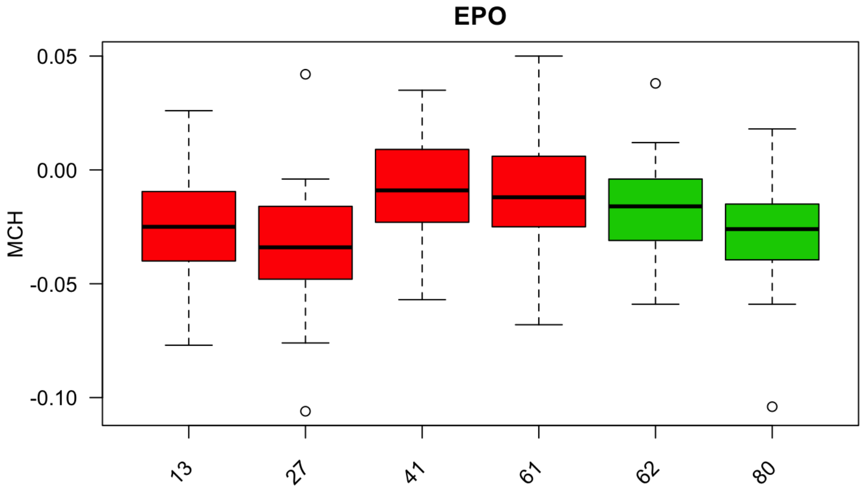
EPO
MCH
0.05
0.00
-0.05
-0.10
13
27
41
61
62
80

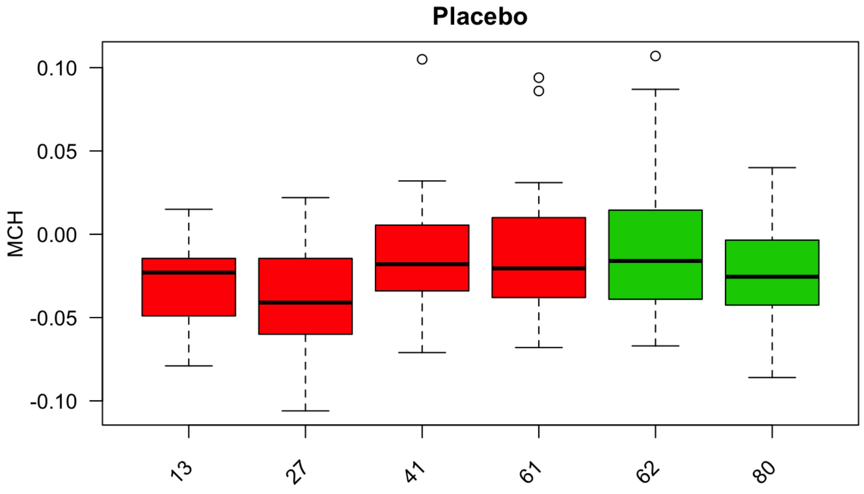
Placebo
MCH
0.10
0.05
0.00
-0.05
-0.10
13
27
41
61
62
80

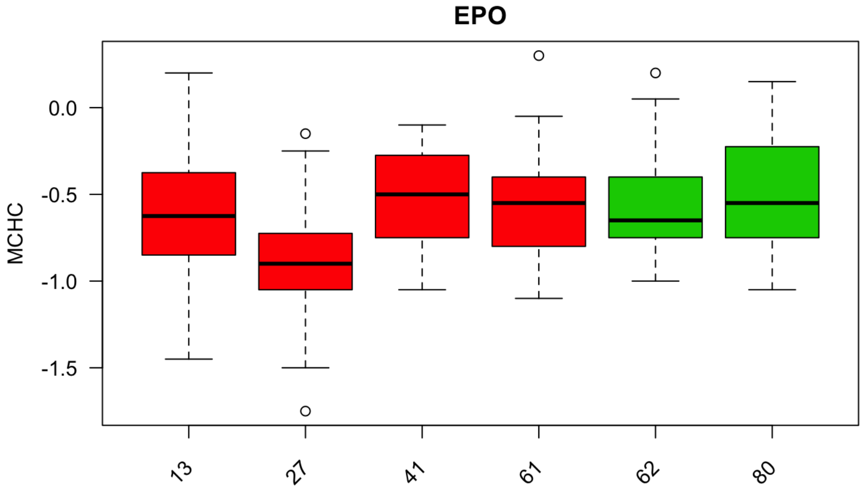
EPO
MCHC
0.0
-0.5
-1.0
-1.5
13
27
41
61
62
80

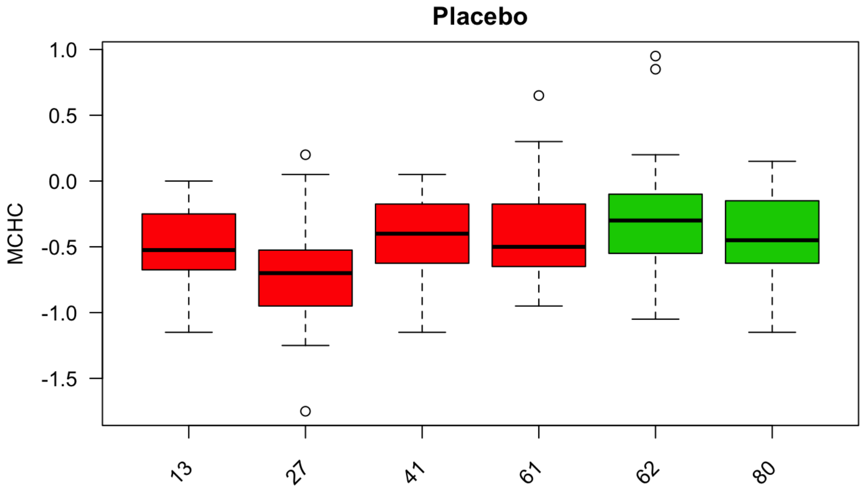
Placebo
MCHC
1.0
0.5
0.0
-0.5
-1.0
-1.5
13
27
41
61
62
80

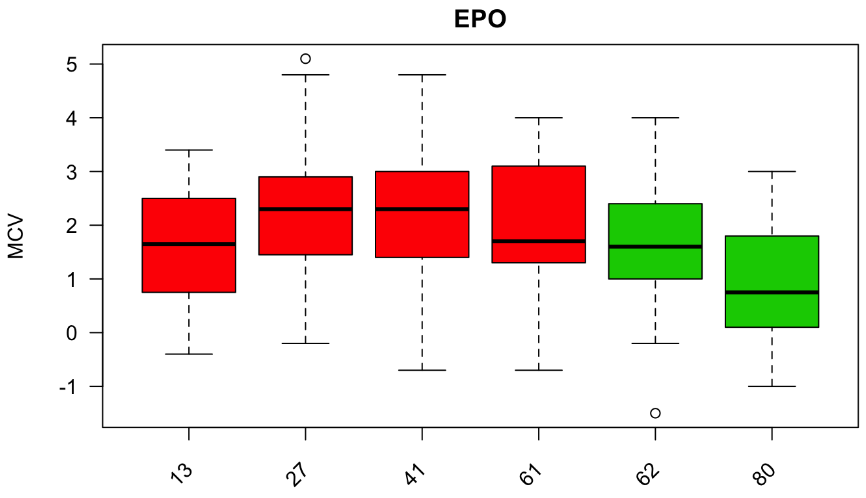
EPO
MCV
5
4
3
2
1
0
-1
13
27
41
61
62
80

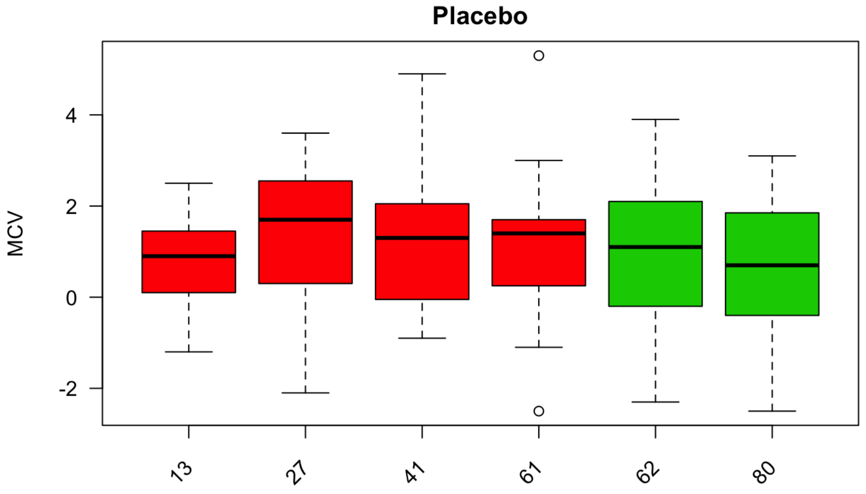
Placebo
MCV
4
2
0
-2
13
27
41
61
62
80

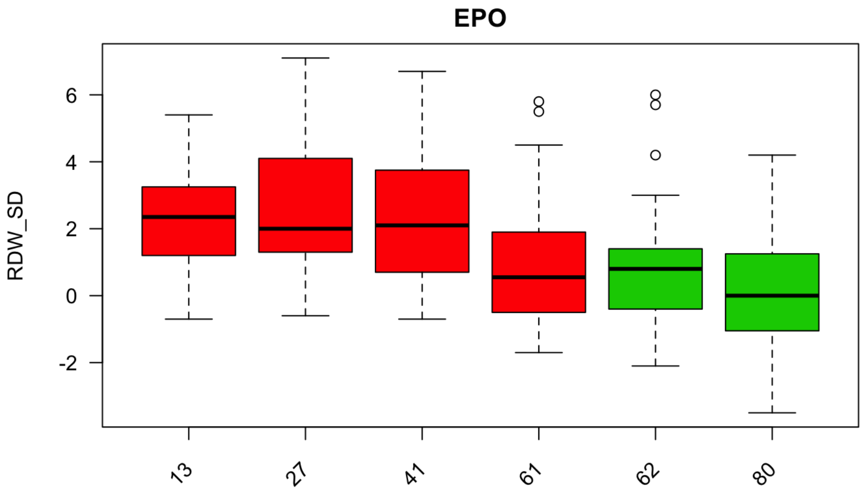
EPO
RDW_SD
6
4
2
0
-2
13
27
41
61
62
80

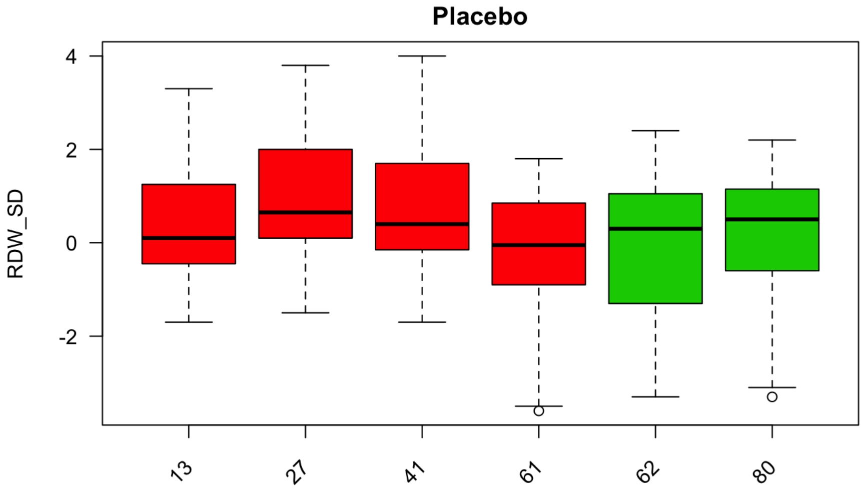
Placebo
RDW_SD
4
2
0
-2
13
27
41
61
62
80

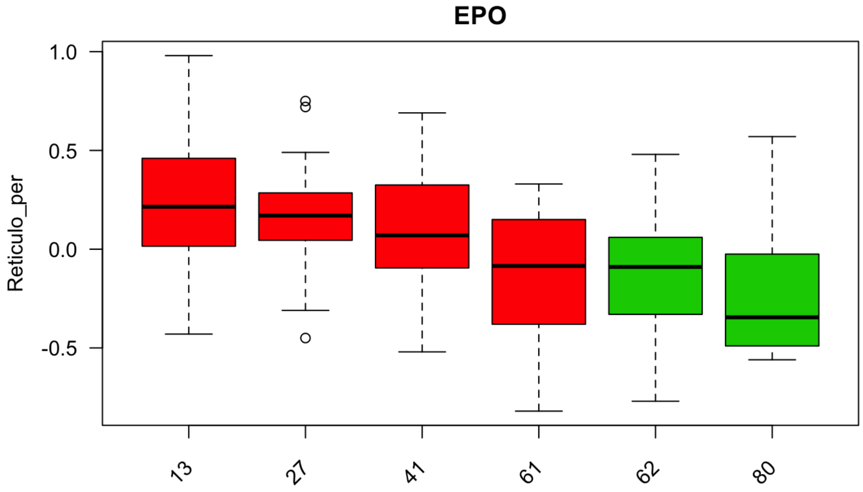
EPO
Reticulo_per
1.0
0.5
0.0
-0.5
13
27
41
61
62
80

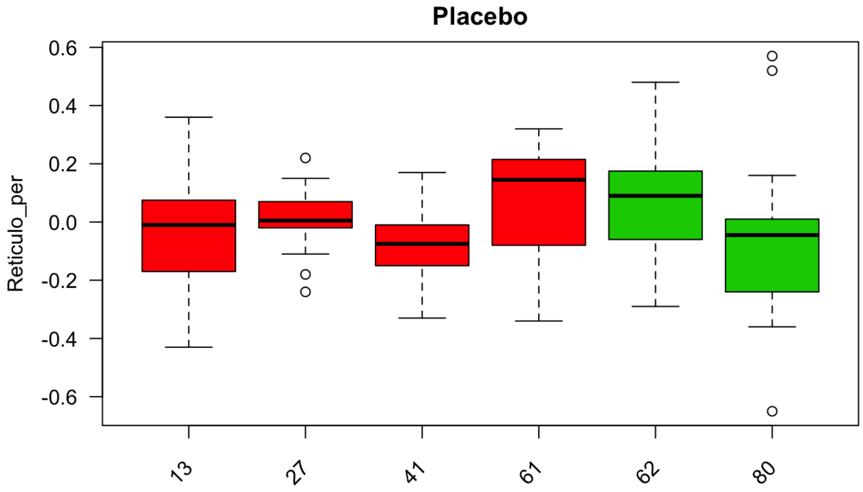
Placebo
Reticulo_per
0.6
0.4
0.2
0.0
-0.2
-0.4
-0.6
13
27
41
61
62
80

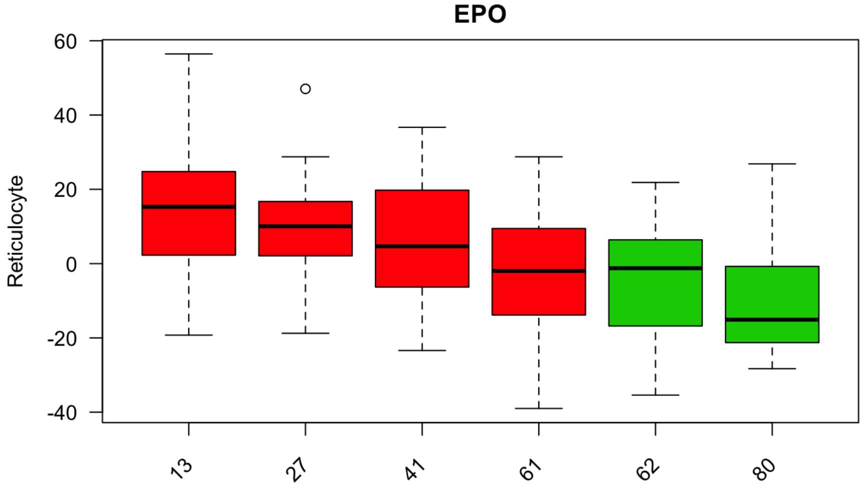
EPO
Reticulocyte
60
40
20
0
-20
-40
13
27
41
61
62
80

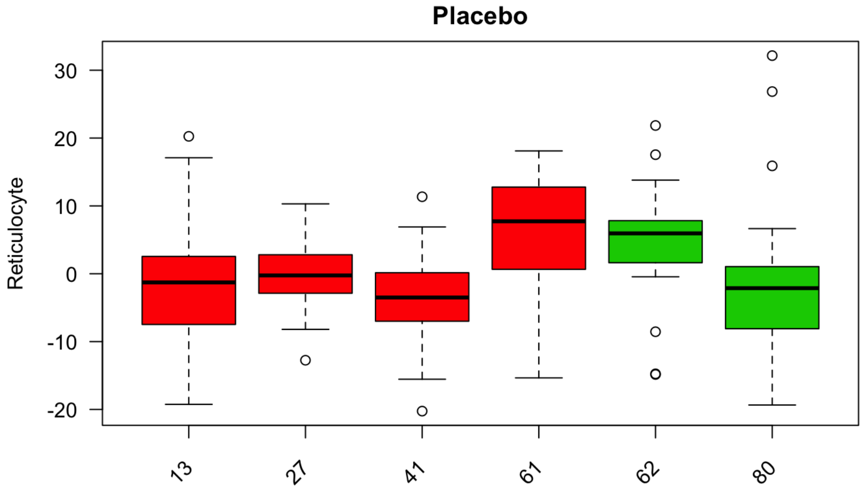
Placebo
Reticulocyte
30
20
10
0
-10
-20
13
27
41
61
62
80

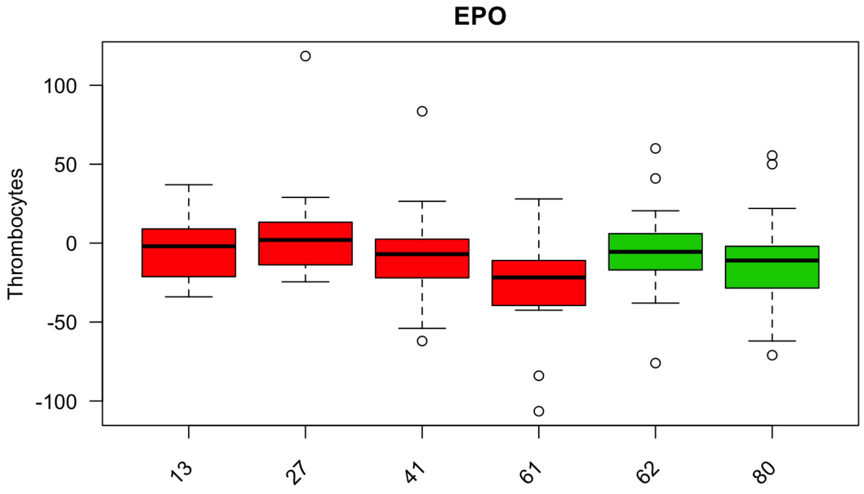
EPO
Thrombocytes
100
50
0
-50
-100
13
27
41
61
62
80

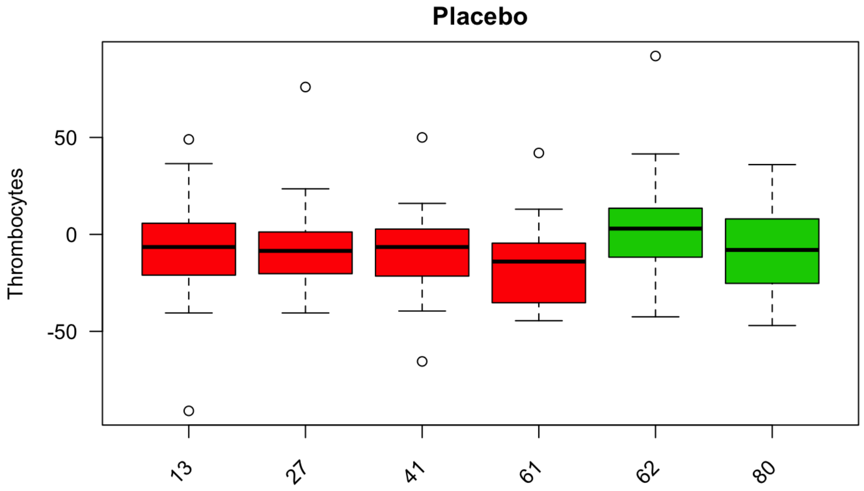


```
R version 3.6.1 Patched (2019-09-23 r77210)
Platform: x86_64-apple-darwin15.6.0 (64-bit)
Running under: macOS Mojave 10.14.6

Matrix products: default
BLAS:   /System/Library/Frameworks/Accelerate.framework/Versions/A/Frameworks/vecLib.framework/Versions/A/libBLA
S.dylib
LAPACK: /Library/Frameworks/R.framework/Versions/3.6/Resources/lib/libRlapack.dylib

locale:
[1] en_US.UTF-8/en_US.UTF-8/en_US.UTF-8/C/en_US.UTF-8/en_US.UTF-8

attached base packages:
[1] stats     graphics  grDevices utils     datasets  methods   base

other attached packages:
[1] multcomp_1.4-12 TH.data_1.0-10  MASS_7.3-51.5   survival_3.1-8  mvtnorm_1.0-12  nlme_3.1-143

loaded via a namespace (and not attached):
 [1] Rcpp_1.0.3       knitr_1.27       magrittr_1.5     splines_3.6.1    lattice_0.20-38
 [6] rlang_0.4.2      stringr_1.4.0    tools_3.6.1      grid_3.6.1       xfun_0.12
[11] htmltools_0.4.0  yaml_2.2.0       digest_0.6.23    Matrix_1.2-18    base64enc_0.1-3
[16] codetools_0.2-16 rconnect_0.8.16 evaluate_0.14    rmarkdown_2.1    sandwich_2.5-1
[21] stringi_1.4.5    compiler_3.6.1   jsonlite_1.6     zoo_1.8-7
```

### Data analysis of phlebotomy intervention study # 3, Hidalgo et al.

Code ▾

##### By Lihua Julie Zhu

Citation:Lindstrom, M.J. and Bates, D.M. (1988) “Newton-Raphson and EM Algorithms for Linear Mixed-Effects Models for Repeated-Measures Data”, Journal of the American Statistical Association, 83, 1014–1022.

Data summary

```
[1] 201   9
```

| | Treatment <fctr> | Subject <int> | time_point <fctr> | Hb..g.dL. <dbl> | MCH..pg. <dbl> | MCHC..g.dL. <dbl> | MCV..fl. <dbl> | Retic.... <dbl> | Retic.count <dbl> |
|---|---|---|---|---|---|---|---|---|---|
| 1 | Phlebotomy | 20 | Baseline 1 | 14.67375 | 30.61060 | 33.37875 | 90 | 0.8606557 | 42 |
| 2 | Phlebotomy | 20 | Baseline 2 | 14.35125 | 30.61060 | 33.70125 | 90 | 1.0403397 | 49 |
| 3 | Phlebotomy | 20 | 3 days | 12.41625 | 30.61060 | 33.37875 | 92 | 1.4285714 | 58 |
| 4 | Phlebotomy | 20 | 14 days | 12.73875 | 30.61060 | 33.37875 | 91 | 2.1615202 | 91 |
| 5 | Phlebotomy | 20 | 25 days | 14.35125 | 30.61060 | 33.86250 | 90 | 1.7094017 | 80 |
| 6 | Phlebotomy | 21 | Baseline 1 | 16.28625 | 28.99952 | 35.63625 | 81 | 0.9769094 | 55 |

6 rows

| | Subject <int> | Round.1 <fctr> | Round.2 <fctr> |
|---|---|---|---|
| 1 | 20 | Transfusion | Placebo |
| 2 | 21 | Placebo | Transfusion |
| 3 | 22 | Placebo | Transfusion |
| 4 | 23 | Placebo | Transfusion |
| 5 | 24 | Transfusion | Placebo |
| 6 | 25 | Transfusion | Placebo |
| 7 | 26 | Placebo | Transfusion |
| 8 | 27 | Transfusion | Placebo |
| 9 | 28 | Placebo | Transfusion |
| 10 | 29 | Placebo | Transfusion |

1-10 of 21 rows Previous **1** 2 3 Next

```
[1] Transfusion Placebo
Levels:  Placebo Transfusion
[1] Placebo          Transfusion      DID NOT COMPLETE
Levels:  DID NOT COMPLETE Placebo Transfusion
, , time_point = 14 days

            Subject
Treatment    1 2 3 4 5 6 7 8 9 20 21 22 23 24 25 26 27 28 29 30 31
  Phlebotomy 1 1 1 1 1 1 1 0 1  1  1  1  1  1  1  1  1  1  1  1  1
  placebo    1 1 1 1 0 1 1 1 1  1  1  1  1  1  1  1  1  1  1  1  1

, , time_point = 25 days

            Subject
Treatment    1 2 3 4 5 6 7 8 9 20 21 22 23 24 25 26 27 28 29 30 31
  Phlebotomy 1 1 1 1 1 1 1 0 1  1  1  1  1  1  1  1  1  1  1  1  1
  placebo    0 1 1 1 1 1 1 1 1  1  1  1  1  1  1  1  1  1  1  1  1

, , time_point = 3 days

            Subject
Treatment    1 2 3 4 5 6 7 8 9 20 21 22 23 24 25 26 27 28 29 30 31
  Phlebotomy 1 1 1 1 1 1 1 0 1  1  1  1  1  1  1  1  1  1  1  1  1
  placebo    1 1 1 1 1 1 1 1 1  1  1  1  1  1  1  1  1  1  1  1  1

, , time_point = Baseline 1

            Subject
Treatment    1 2 3 4 5 6 7 8 9 20 21 22 23 24 25 26 27 28 29 30 31
  Phlebotomy 1 1 1 1 1 1 1 0 1  1  1  1  1  1  1  1  1  1  1  1  1
  placebo    1 1 1 1 1 1 1 1 1  1  1  1  1  1  1  1  1  1  1  1  1

, , time_point = Baseline 2

            Subject
Treatment    1 2 3 4 5 6 7 8 9 20 21 22 23 24 25 26 27 28 29 30 31
  Phlebotomy 1 1 1 1 1 1 1 0 1  1  1  1  1  1  1  1  1  1  1  1  1
  placebo    0 1 0 1 1 1 1 1 1  1  1  1  1  1  1  1  1  1  1  1  1

[1] "Treatment"   "Subject"     "time_point"  "Hb..g.dL."   "MCH..pg."    "MCHC..g.dL."
[7] "MCV..fl."    "Retic...."   "Retic.count"


Correlation matrix for the dependent varilables
                Hb..g.dL.    MCH..pg. MCHC..g.dL.    MCV..fl.  Retic.... Retic.count
Hb..g.dL.    1.000000000 -0.06580095   0.2300736 -0.23750662 -0.2420295 0.007253484
MCH..pg.    -0.065800953  1.00000000   0.2607208  0.75688745  0.2619209 0.123775019
MCHC..g.dL.  0.230073640  0.26072085   1.0000000 -0.31399836  0.1030669 0.123685146
MCV..fl.    -0.237506619  0.75688745  -0.3139984  1.00000000  0.1938071 0.025301734
Retic....   -0.242029523  0.26192087   0.1030669  0.19380706  1.0000000 0.955760979
Retic.count  0.007253484  0.12377502   0.1236851  0.02530173  0.9557610 1.000000000
[1] Baseline 1 Baseline 2 3 days     14 days    25 days
Levels: 14 days 25 days 3 days Baseline 1 Baseline 2
```

Cross-over design

```
##########Use the mean of Baseline 1 and 2 as Baseline value
################[1] 162  11
[1] 3 days   14 days  25 days  Baseline
Levels: 14 days 25 days 3 days Baseline 1 Baseline 2 Baseline


################ time point:  14 days measurement:  Hb..g.dL. ###############
```

| | numDF <int> | denDF <dbl> | F-value <chr> | p-value <chr> |
|---|---|---|---|---|
| (Intercept) | 1 | 18 | 91.73061 | <.0001 |
| Treatment | 1 | 17 | 55.81482 | <.0001 |
| period | 1 | 17 | 4.12047 | 0.0583 |
| tseq | 1 | 18 | 3.23582 | 0.0888 |

4 rows

```
##########################################################################
##########################


################ time point:  25 days measurement:  Hb..g.dL. ###############
```

| | numDF <int> | denDF <dbl> | F-value <chr> | p-value <chr> |
|---|---|---|---|---|
| (Intercept) | 1 | 18 | 24.595951 | 0.0001 |
| Treatment | 1 | 17 | 8.268737 | 0.0105 |
| period | 1 | 17 | 0.007883 | 0.9303 |
| tseq | 1 | 18 | 2.284370 | 0.1480 |

4 rows

```
##########################################################################
##########################


################ time point:  3 days measurement:  Hb..g.dL. ###############
```

| | numDF <int> | denDF <dbl> | F-value <chr> | p-value <chr> |
|---|---|---|---|---|
| (Intercept) | 1 | 18 | 111.83610 | <.0001 |
| Treatment | 1 | 18 | 184.52548 | <.0001 |
| period | 1 | 18 | 1.56737 | 0.2266 |
| tseq | 1 | 18 | 3.32853 | 0.0847 |

4 rows

```
##########################################################################
##########################
```

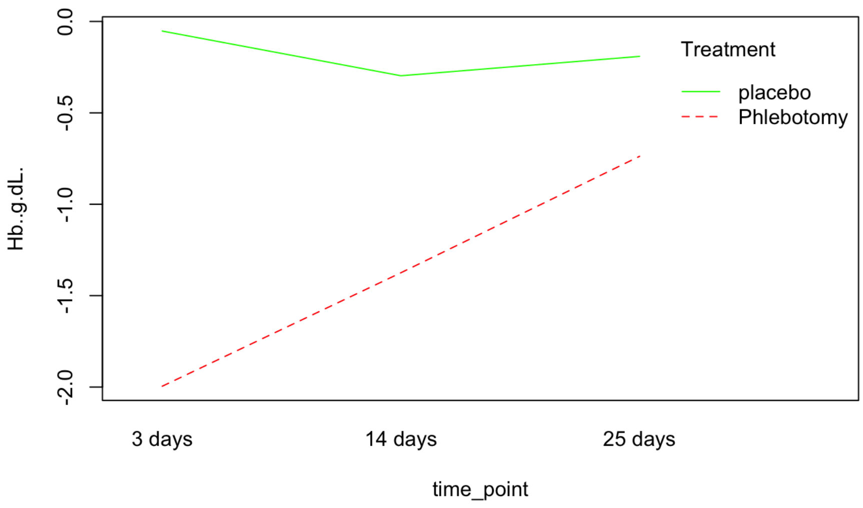


```
################ time point:  14 days measurement:  MCH..pg. ###############
```

| | numDF <int> | denDF <dbl> | F-value <chr> | p-value <chr> |
|---|---|---|---|---|
| (Intercept) | 1 | 18 | 0.0290347 | 0.8666 |

| | numDF <int> | denDF <dbl> | F-value <chr> | p-value <chr> |
|---|---|---|---|---|
| Treatment | 1 | 17 | 0.7338136 | 0.4036 |
| period | 1 | 17 | 1.5821467 | 0.2255 |
| tseq | 1 | 18 | 0.0226521 | 0.8820 |

4 rows

```
############################################################################
###########################


############### time point:  25 days measurement:  MCH..pg. ##############
```

| | numDF <int> | denDF <dbl> | F-value <chr> | p-value <chr> |
|---|---|---|---|---|
| (Intercept) | 1 | 18 | 1.9810073 | 0.1763 |
| Treatment | 1 | 17 | 0.0272362 | 0.8709 |
| period | 1 | 17 | 2.1342341 | 0.1623 |
| tseq | 1 | 18 | 0.2755240 | 0.6061 |

4 rows

```
############################################################################
###########################


############### time point:  3 days measurement:  MCH..pg. ##############
```

| | numDF <int> | denDF <dbl> | F-value <chr> | p-value <chr> |
|---|---|---|---|---|
| (Intercept) | 1 | 18 | 0.0583115 | 0.8119 |
| Treatment | 1 | 18 | 0.0583115 | 0.8119 |
| period | 1 | 18 | 0.5660340 | 0.4616 |
| tseq | 1 | 18 | 2.9691754 | 0.1020 |

4 rows

```
############################################################################
###########################
```

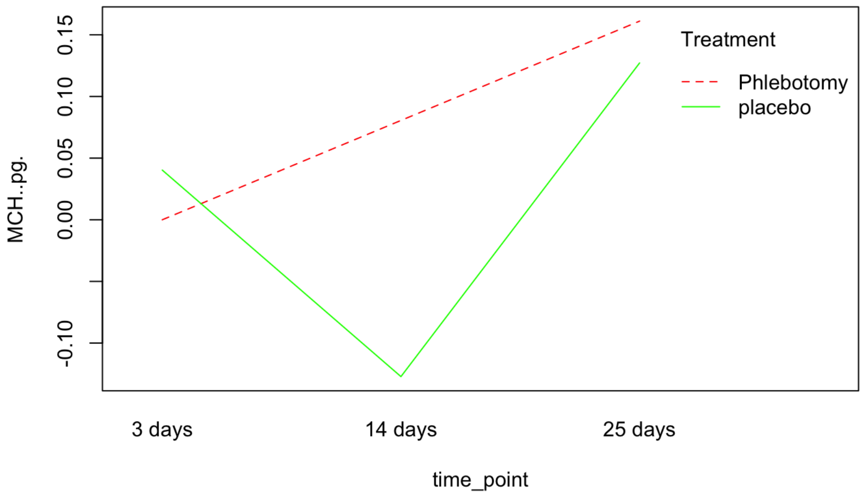


```
############### time point:  14 days measurement:  MCHC..g.dL. ##############
```

| | numDF <int> | denDF <dbl> | F-value <chr> | p-value <chr> |
|---|---|---|---|---|
| (Intercept) | 1 | 18 | 2.0187549 | 0.1725 |
| Treatment | 1 | 17 | 0.0004147 | 0.9840 |
| period | 1 | 17 | 1.2044672 | 0.2877 |
| tseq | 1 | 18 | 0.0683320 | 0.7967 |

4 rows

```
###########################################################################
##########################


############### time point:  25 days measurement:  MCHC..g.dL. ##############
```

| | numDF <int> | denDF <dbl> | F-value <chr> | p-value <chr> |
|---|---|---|---|---|
| (Intercept) | 1 | 18 | 1.4746692 | 0.2403 |
| Treatment | 1 | 17 | 0.8481687 | 0.3700 |
| period | 1 | 17 | 0.0047336 | 0.9460 |
| tseq | 1 | 18 | 0.0615946 | 0.8068 |

4 rows

```
###########################################################################
##########################


############### time point:  3 days measurement:  MCHC..g.dL. ##############
```

| | numDF <int> | denDF <dbl> | F-value <chr> | p-value <chr> |
|---|---|---|---|---|
| (Intercept) | 1 | 18 | 1.3797674 | 0.2554 |
| Treatment | 1 | 18 | 0.3338669 | 0.5705 |
| period | 1 | 18 | 0.1374699 | 0.7151 |
| tseq | 1 | 18 | 1.3085410 | 0.2676 |

4 rows

```
###########################################################################
##########################
```

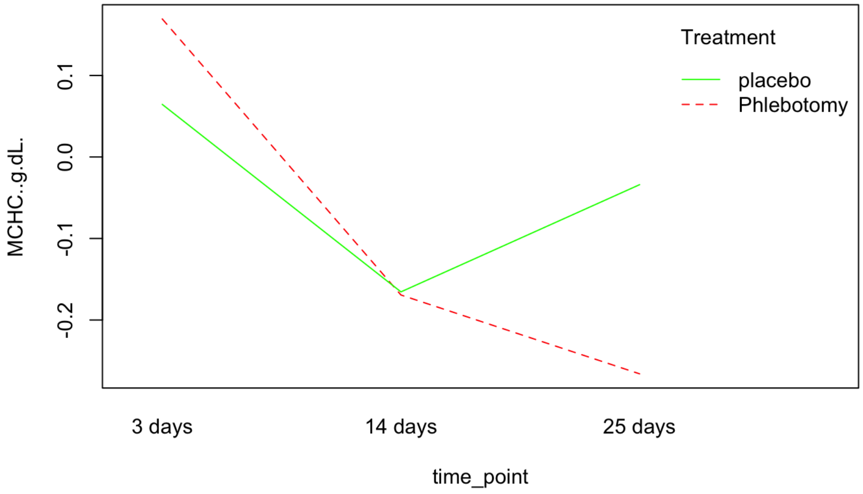


```
############### time point:  14 days measurement:  MCV..fl. ##############
```

| | numDF <int> | denDF <dbl> | F-value <chr> | p-value <chr> |
|---|---|---|---|---|
| (Intercept) | 1 | 18 | 14.437097 | 0.0013 |
| Treatment | 1 | 17 | 3.364034 | 0.0842 |
| period | 1 | 17 | 0.679356 | 0.4212 |
| tseq | 1 | 18 | 0.841953 | 0.3710 |

4 rows

```
###########################################################################
##########################


############### time point:  25 days measurement:  MCV..fl. ##############
```

| | numDF <int> | denDF <dbl> | F-value <chr> | p-value <chr> |
|---|---|---|---|---|
| (Intercept) | 1 | 18 | 16.901658 | 0.0007 |

| | numDF <int> | denDF <dbl> | F-value <chr> | p-value <chr> |
|---|---|---|---|---|
| Treatment | 1 | 17 | 6.744724 | 0.0188 |
| period | 1 | 17 | 1.023332 | 0.3259 |
| tseq | 1 | 18 | 1.672958 | 0.2122 |

4 rows

```
##########################################################################
##########################

############### time point:  3 days measurement:  MCV..fl. ##############
```

| | numDF <int> | denDF <dbl> | F-value <chr> | p-value <chr> |
|---|---|---|---|---|
| (Intercept) | 1 | 18 | 0.1562853 | 0.6972 |
| Treatment | 1 | 18 | 0.0173650 | 0.8966 |
| period | 1 | 18 | 2.4839018 | 0.1324 |
| tseq | 1 | 18 | 0.2401286 | 0.6300 |

4 rows

```
##########################################################################
##########################
```

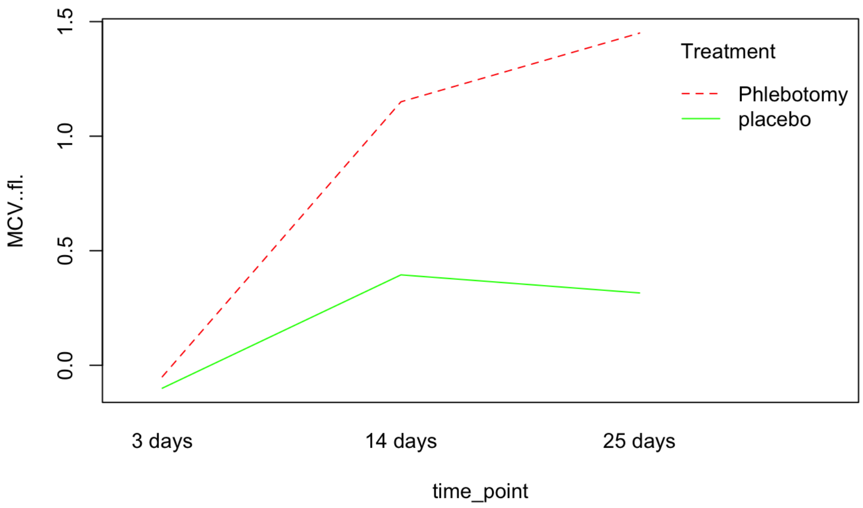


```
############### time point:  14 days measurement:  Retic.... ##############
```

| | numDF <int> | denDF <dbl> | F-value <chr> | p-value <chr> |
|---|---|---|---|---|
| (Intercept) | 1 | 18 | 125.71963 | <.0001 |
| Treatment | 1 | 17 | 76.59931 | <.0001 |
| period | 1 | 17 | 0.71310 | 0.4101 |
| tseq | 1 | 18 | 1.45510 | 0.2433 |

4 rows

```
##########################################################################
##########################

############### time point:  25 days measurement:  Retic.... ##############
```

| | numDF <int> | denDF <dbl> | F-value <chr> | p-value <chr> |
|---|---|---|---|---|
| (Intercept) | 1 | 18 | 24.552824 | 0.0001 |
| Treatment | 1 | 17 | 15.699562 | 0.0010 |
| period | 1 | 17 | 0.112146 | 0.7418 |
| tseq | 1 | 18 | 0.842159 | 0.3709 |

4 rows

```
##########################################################################
##########################

############### time point:  3 days measurement:  Retic.... ##############
```

| | numDF <int> | denDF <dbl> | F-value <chr> | p-value <chr> |
|---|---|---|---|---|
| (Intercept) | 1 | 18 | 51.63309 | <.0001 |
| Treatment | 1 | 18 | 35.08276 | <.0001 |
| period | 1 | 18 | 0.00583 | 0.9400 |
| tseq | 1 | 18 | 0.00277 | 0.9586 |

4 rows

```
##########################################################################
##########################
```

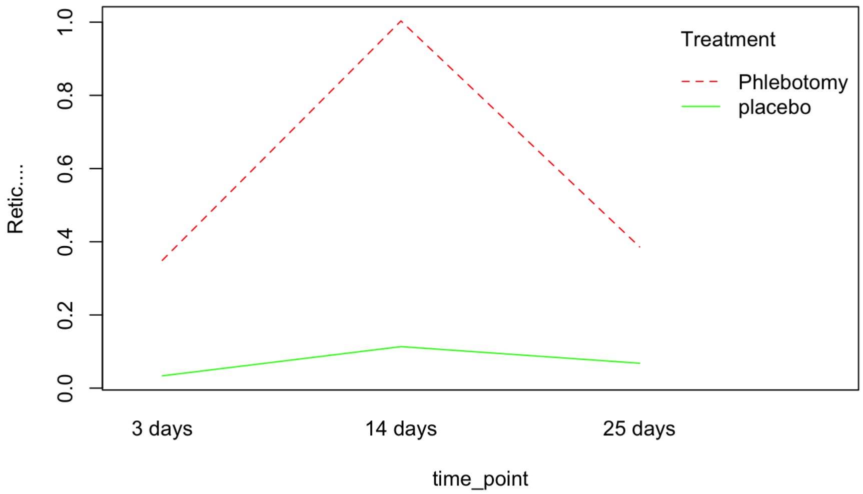


```
############### time point:  14 days measurement:  Retic.count ##############
```

| | numDF <int> | denDF <dbl> | F-value <chr> | p-value <chr> |
|---|---|---|---|---|
| (Intercept) | 1 | 18 | 115.28009 | <.0001 |
| Treatment | 1 | 17 | 70.15011 | <.0001 |
| period | 1 | 17 | 2.06103 | 0.1693 |
| tseq | 1 | 18 | 1.17870 | 0.2919 |

4 rows

```
##########################################################################
##########################

############### time point:  25 days measurement:  Retic.count ##############
```

| | numDF <int> | denDF <dbl> | F-value <chr> | p-value <chr> |
|---|---|---|---|---|
| (Intercept) | 1 | 18 | 14.198772 | 0.0014 |
| Treatment | 1 | 17 | 10.587278 | 0.0047 |
| period | 1 | 17 | 0.125420 | 0.7276 |
| tseq | 1 | 18 | 0.242599 | 0.6283 |

4 rows

```
##########################################################################
##########################

############### time point:  3 days measurement:  Retic.count ##############
```

| | numDF <int> | denDF <dbl> | F-value <chr> | p-value <chr> |
|---|---|---|---|---|
| (Intercept) | 1 | 18 | 9.186449 | 0.0072 |

| | numDF <int> | denDF <dbl> | F-value <chr> | p-value <chr> |
|---|---|---|---|---|
| Treatment | 1 | 18 | 5.519200 | 0.0304 |
| period | 1 | 18 | 0.088983 | 0.7689 |
| tseq | 1 | 18 | 0.410961 | 0.5296 |

4 rows

```
##########################################################################
##########################
```

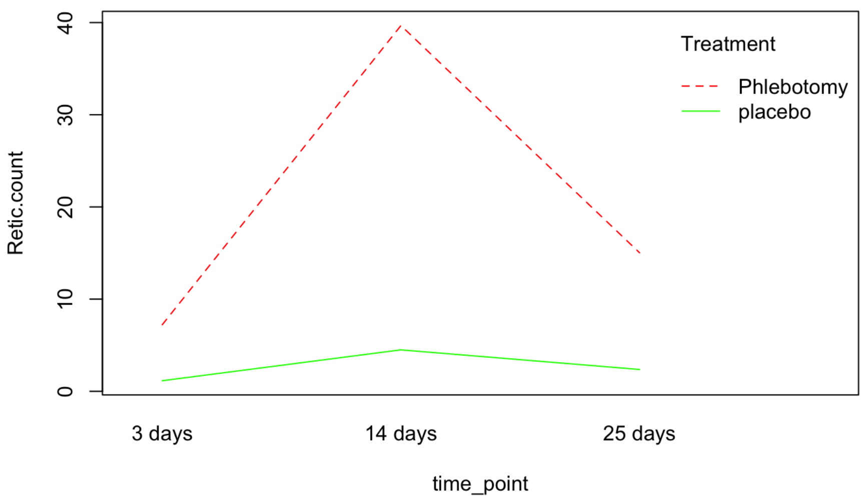


Data visualization

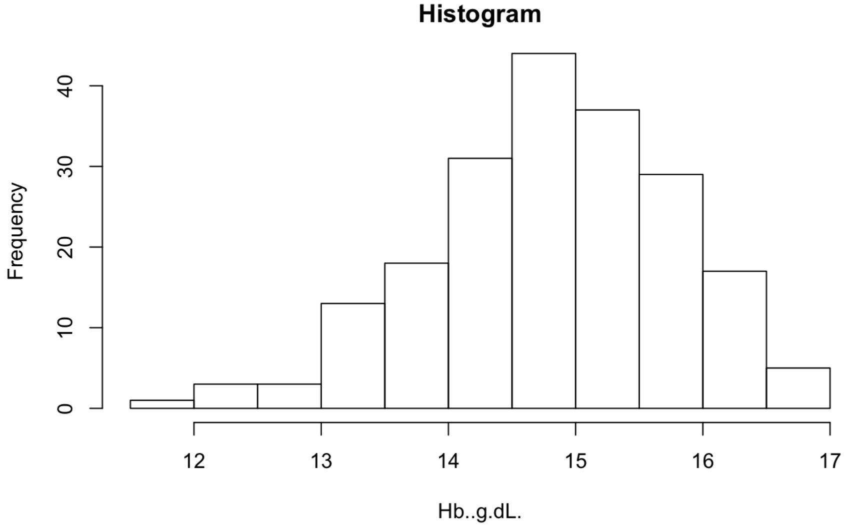

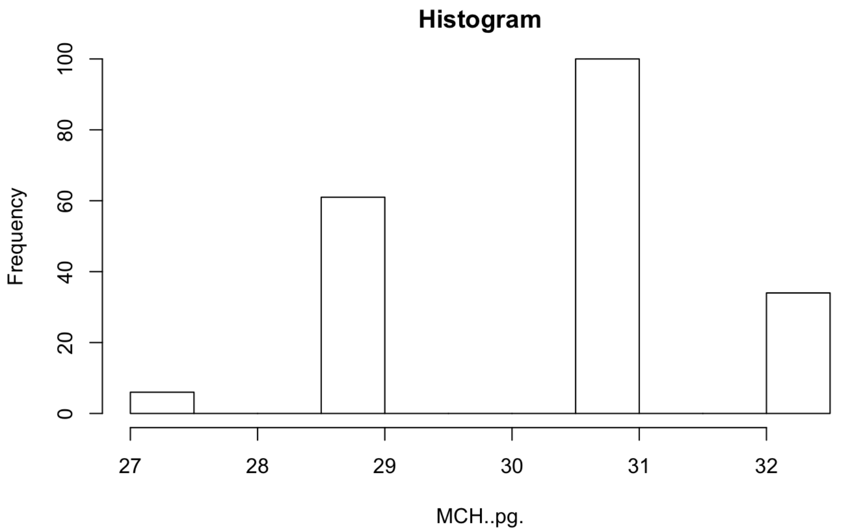
Histogram
Frequency
0
20
40
60
80
100
27
28
29
30
31
32
MCH..pg.

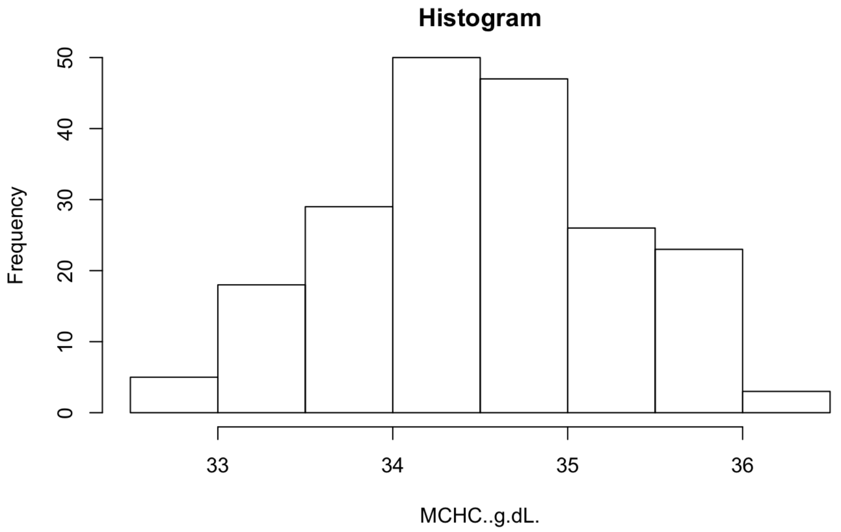
Histogram
Frequency
0
10
20
30
40
50
33
34
35
36
MCHC..g.dL.

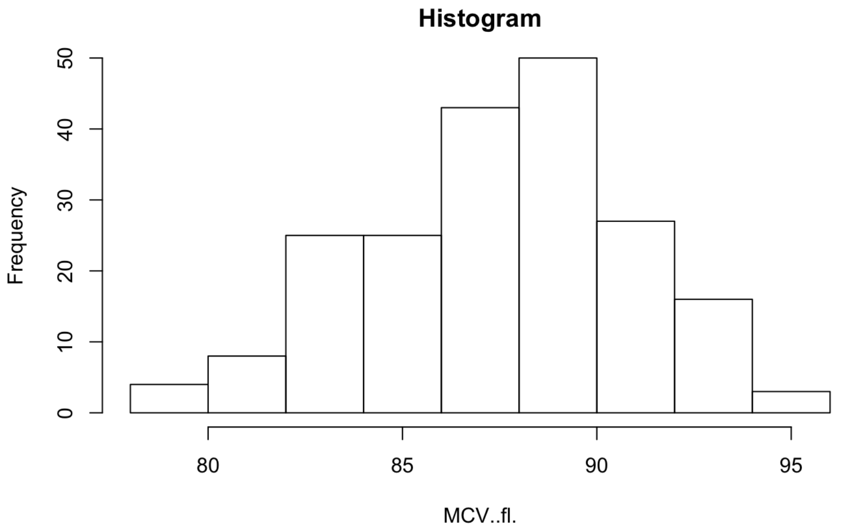
Histogram
Frequency
0
10
20
30
40
50
80
85
90
95
MCV..fl.

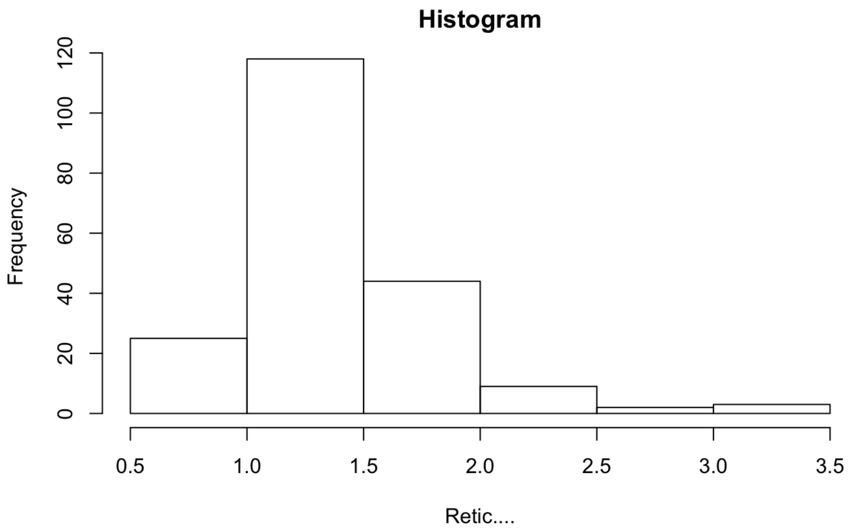
Histogram
Frequency
Retic....
0
20
40
60
80
100
120
0.5
1.0
1.5
2.0
2.5
3.0
3.5

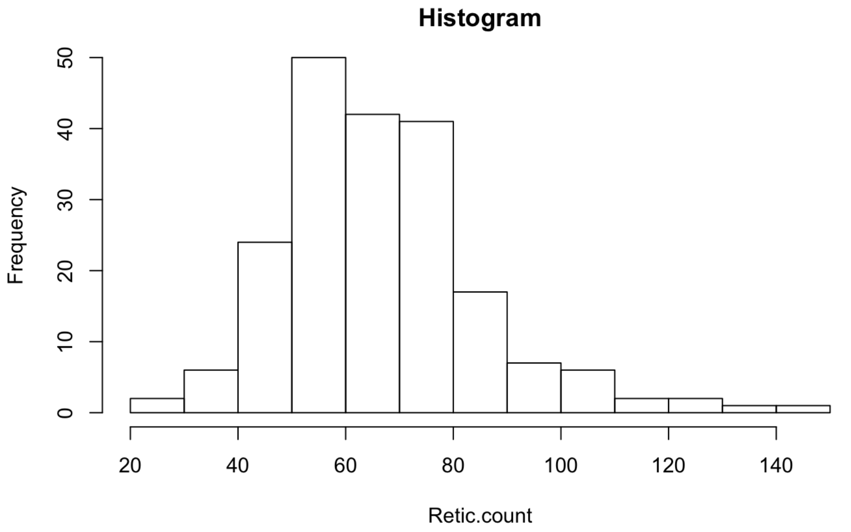
Histogram
Frequency
Retic.count
0
10
20
30
40
50
20
40
60
80
100
120
140
