## Supplementary material for "The Erythropoietin Receptor Stimulates Rapid Cycling and Formation of Larger Red Cells During Mouse and Human Erythropoiesis": Simulation analysis of MCV (from another version of this preprint: version 1, 2020-11-30)

### Simulations of MCV time course during and following Epo administration, assuming EpoR signaling *does not* change red cell size

#### I Summary

In human intervention studies, we found an increase in MCV that does not correlate with reticulocyte count (Extended Data Figures 9,10 and Figure 6; statistical analysis supplement). Two hypotheses might account for the increase in MCV.

The first, or null hypothesis, is based on the known negative correlation between red cell (RC) age and volume. Although the largest loss in RC volume occurs during reticulocyte maturation, RCs continue to lose volume throughout their life (in part as a result of vesiculation)[1-4]. In the steady state, the MCV is the average of the volume of an equal number of RCs of every age. Immediately following Epo administration, in addition to an increase in reticulocytes, there is an increase in the relative number of mature, younger RCs. This skew in the age distribution of mature RCs is expected to increase the MCV and may be sufficient to account for the observed increase that we see in the human studies.

In the alternative hypothesis, the observed increase in MCV cannot be accounted for solely by the skewing of the RC age distribution, and is also the result of EpoR-signaling -mediated increase in cell size. This is based on our experimental findings in the mouse, which show that EpoR signaling increases cell size in an Epo-dose dependent manner at every stage of erythroid terminal differentiation (ETD), leading to the production of larger reticulocytes and RCs.

Below, we simulated the first, or null hypothesis, and then determined whether it is sufficient to account for the observations in the human studies. We simulated the MCV time course during and following Epo treatments of varying lengths and strengths. We looked at treatment durations from 10 to 100 days, that increased erythropoietic rate ('ery. rate') by 1.25 to 10 fold the basal rate. We found that the increased MCV predicted by the null hypothesis accounts for 30 to 50% of the observed increase in the human studies. Therefore, additional factors must contribute to the increase in MCV. These findings are consistent with our experiments in the mouse, which show an EpoR signaling- mediated increase in cell size.

#### II Input variables

##### A Reticulocyte parameters during the basal steady state

*Input:*

We set the MCV in the basal state to 1, initial circulating reticulocyte volume (Vr) to 1.25, and the reticulocyte circulation life to 1 day.

*Justification:*

- Reticulocytes take 3.5 -4 days to mature, with the last 24 h of this process taking place in the circulation. The mean volume of circulating reticulocytes is 24% larger than the MCV [5, 6].

##### B Reticulocyte parameters during Epo administration

Increased erythropoietic rate is associated with early release of immature, larger reticulocytes from the marrow [7-9].

*Input:*

We assume linear models for the increase in reticulocyte volumes and circulation times with increasing erythropoietic rate (Fig.1).

*Justification:*

Fig. 1: Reticulocyte volumes and circulation times during stress

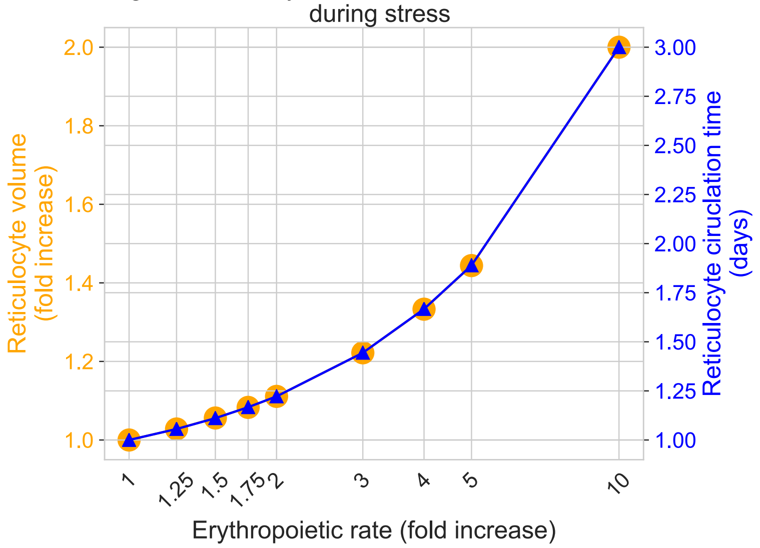

- In extreme stress (aplastic anemia, phenylhydrazine treatment in animals), there is a doubling of Vr [6].
- Administration of 1200 IU/kg over a 10 day period, which resulted in a 2.5 fold increase in the absolute reticulocyte count, was associated with a 20% increased in mean reticulocyte volume [10,11]. The linear model that we assume gives a close approximation to this, with a 22% increase in Vr for a 3 -fold increase in erythropoietic rate.
- Reticulocyte maturation time in the circulation increases approximately linearly with falling hematocrit [7,12].

##### C RC volume distribution as a function of RC age in the basal state

*Input:*

We assumed that reticulocytes have an initial volume Vr that is 1.25 fold larger than the MCV (see 'IIA' above). Following an initial rapid volume loss corresponding to reticulocyte maturation, volume continues to decrease linearly with cell age, but at a slower pace. We assigned this loss to the variable 'd', which denotes the fraction by which the newest mature (non-reticulocyte) RC is larger than the oldest RC. The total volume loss over the entire reticulocyte and mature RC life can be calculated for varying values of 'd'. Setting 'd' in the range of 0.15 to 0.2 gives a total RC volume loss of 27% to 29%, including an initial loss in reticulocyte volume of 11% to 14%. We chose to use d = 0.2 for the simulations below, in order to make the null hypothesis as likely as possible.

Fig. 2: Calculating RC volumes as a function of RC age

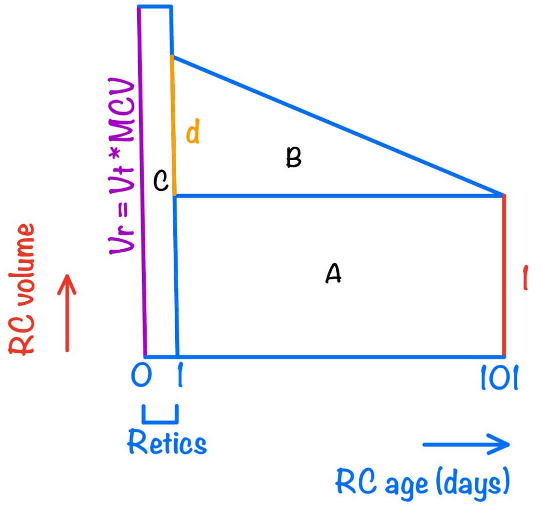


In the steady state, there are equal numbers of cells at each age. We assume retics circulate during day 1, and RCs circulate for 100 additional days.

MCV = mean cell volume.
Volume of oldest RC = 1 (arbitrary units)
Vr = reticulocyte volume
Vt = 1.25 factor by which Vr is larger than MCV
d = loss in mature red cell volume during circulation life

MCV = (A+B+C)/101
A = 100
B = 100d/2 = 50d
C = Vt * MCV

$$MCV = \frac{100 + 50d}{101 - Vt}$$

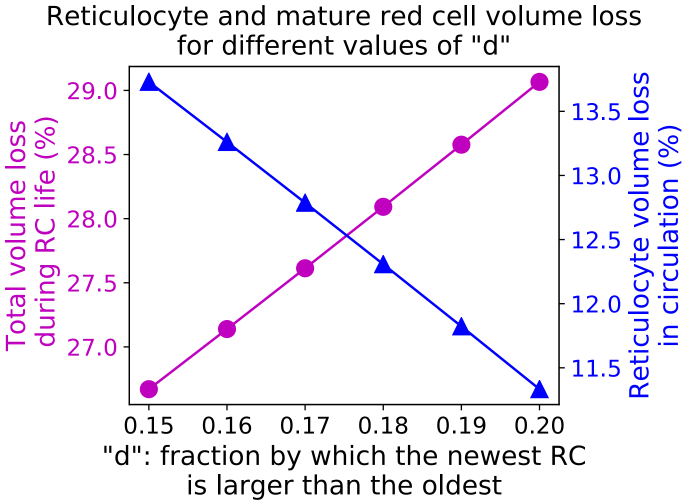

*Justification:*

- There is a linear decline in volume with RC Hemoglobin A1c (an indicator of RC age), of ~30% over the entire span of circulation life [1]
- There is an overall ~30% decrease in RC volume with aging [2, 13].
- There is a linear decline in Forward Scatter (flow cytometry) with RC age [4].
- There is a loss of 10 to 14% of cell surface area during the reticulocyte stage (~1 day), and an additional 16-17% during the rest of RC life [3]. The surface area/volume ratio remain constant [3].

#### III RC volume as a function of RC age, in basal and increased erythropoietic rates

The null hypothesis suggests that increased erythropoietic rate does not change mature RC volumes. Only reticulocyte size and time in the circulation increase, based on the input variables in 'II B' above. Figure 3 shows the volume distribution with age for increasing erythropoietic rates (for plotting purposes, but not for MCV calculations, the reticulocyte circulating time in Fig.3 is rounded to nearest integer). Other than for reticulocytes, RC age is expressed as an integer number of days.

Fig. 3: RC volume distribution as a function of RC age, for the indicated increase in erythropoietic rate (MCV calculated for a steady state increase in erythropoietic rate)

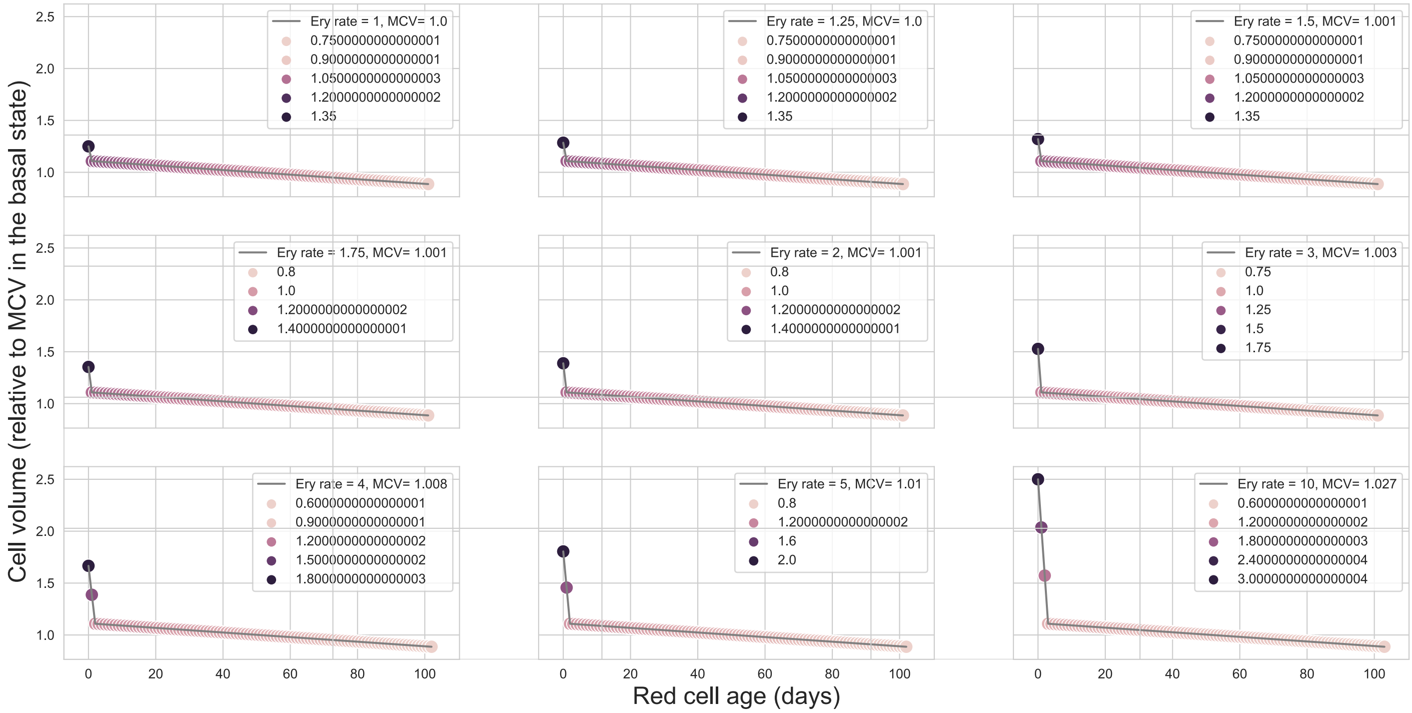


#### IV RC volume distributions in response to continuous stimulation with Epo

We simulated the RC volume distributions for various durations and levels of Epo administration (Figure 4A,B).

We examined 9 different levels of increased erythropoietic rate: 1.25, 1.5, 1.75, 2, 3, 4, 5, & 10 fold the basal rate. For each of these, we determined the volume distribution by increasing the number of RCs for the appropriate RC ages, and in addition, by increasing the number, size, and circulation times of reticulocytes.

NB all cells of a given age were considered to have the same volume.

Fig. 4A: RC volume distributions for 1.25 to 2 fold increase in ery. rate, of 10 to 90 day duration

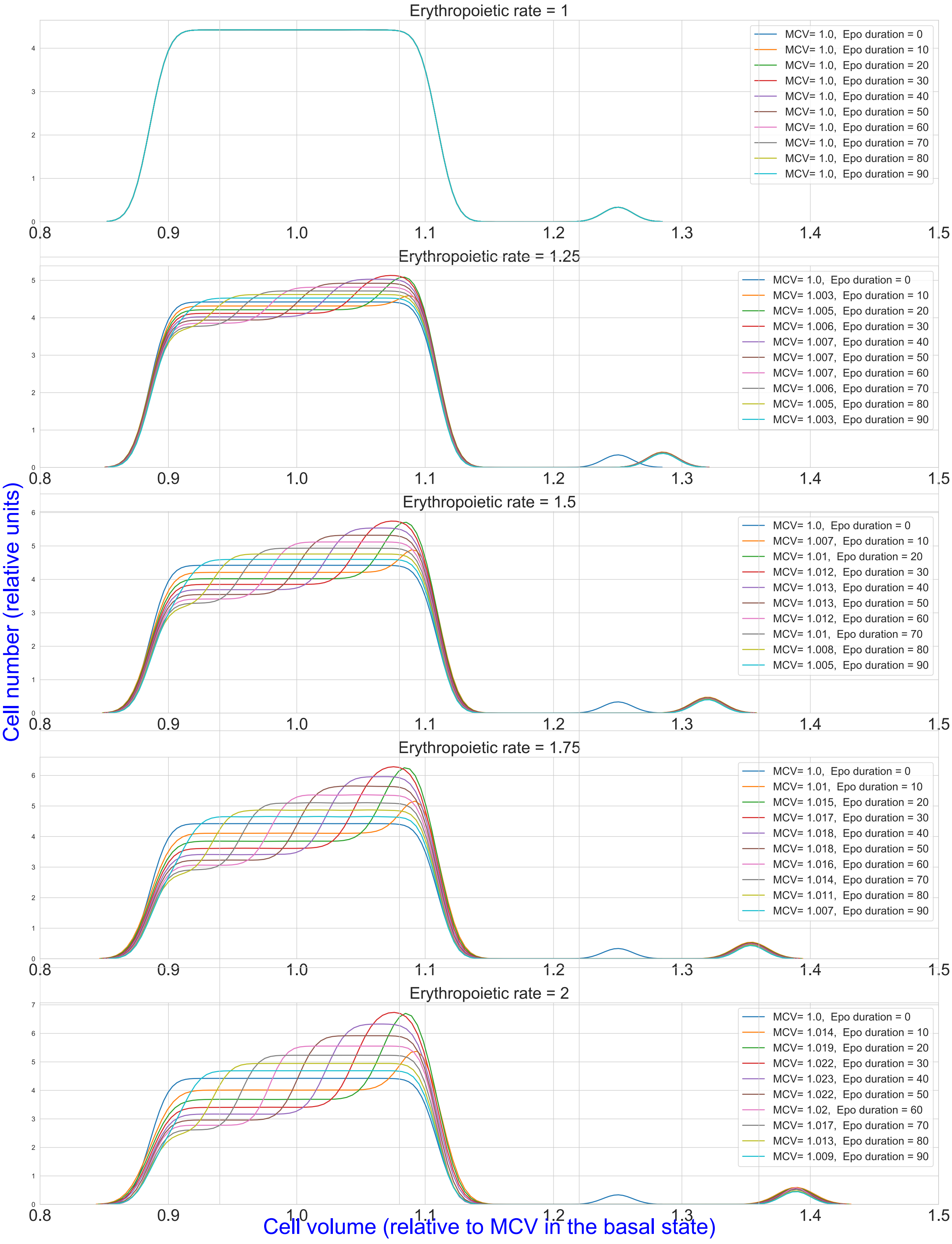

Fig. 4B: RC volume distributions for 3 to 10 fold increase in ery. rate, of 10 to 90 day duration

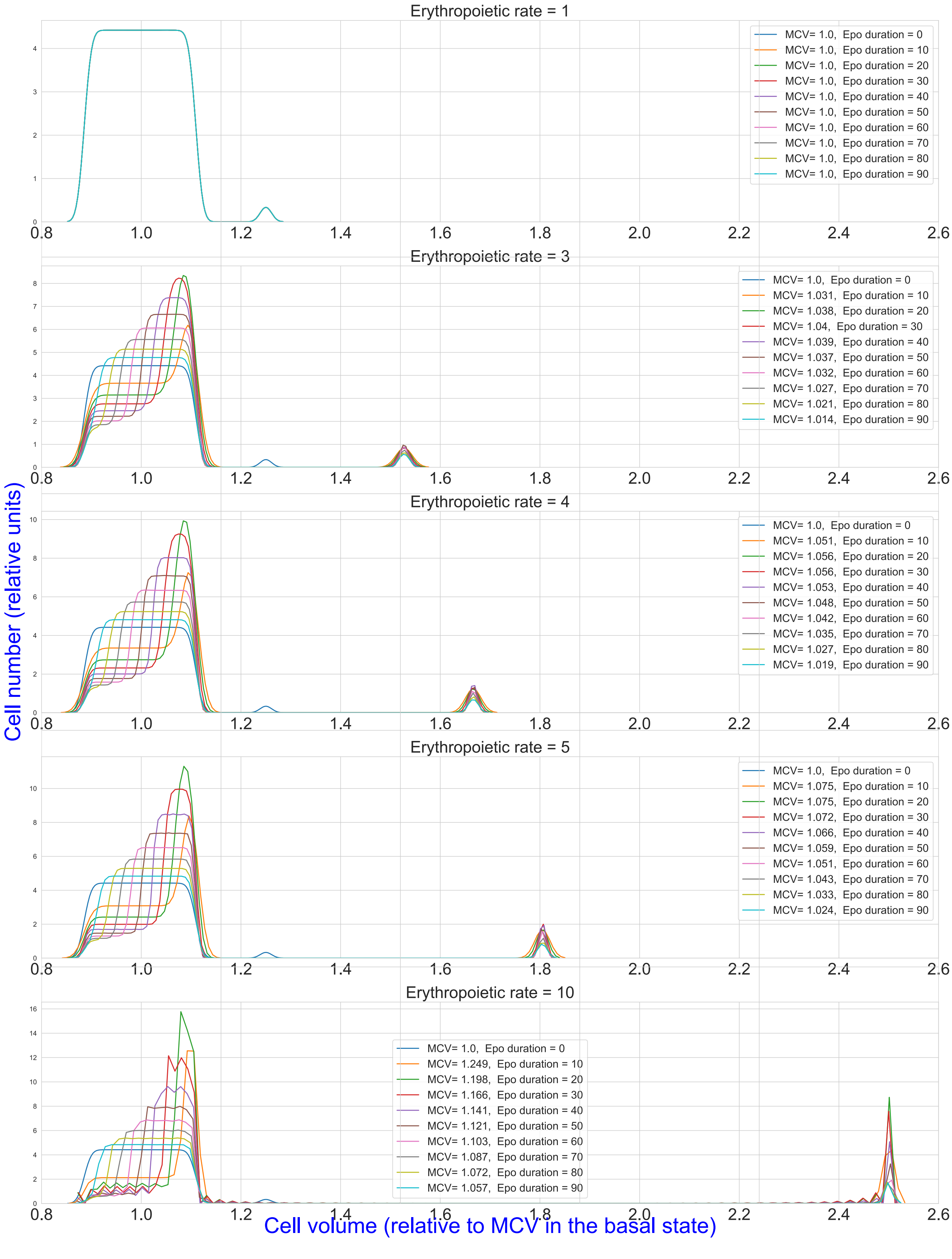

#### V Prediction of MCV time course during continuous stimulation with Epo

The MCV values computed from each of the volume distributions in ‘IV’ were used to plot the time course of MCV changes that occur with continuous Epo stimulation (Fig. 5A, B):

Fig 5A: MCV time course during continuous Epo treatment stimulating up to a 2-fold increase in erythropoietic rate

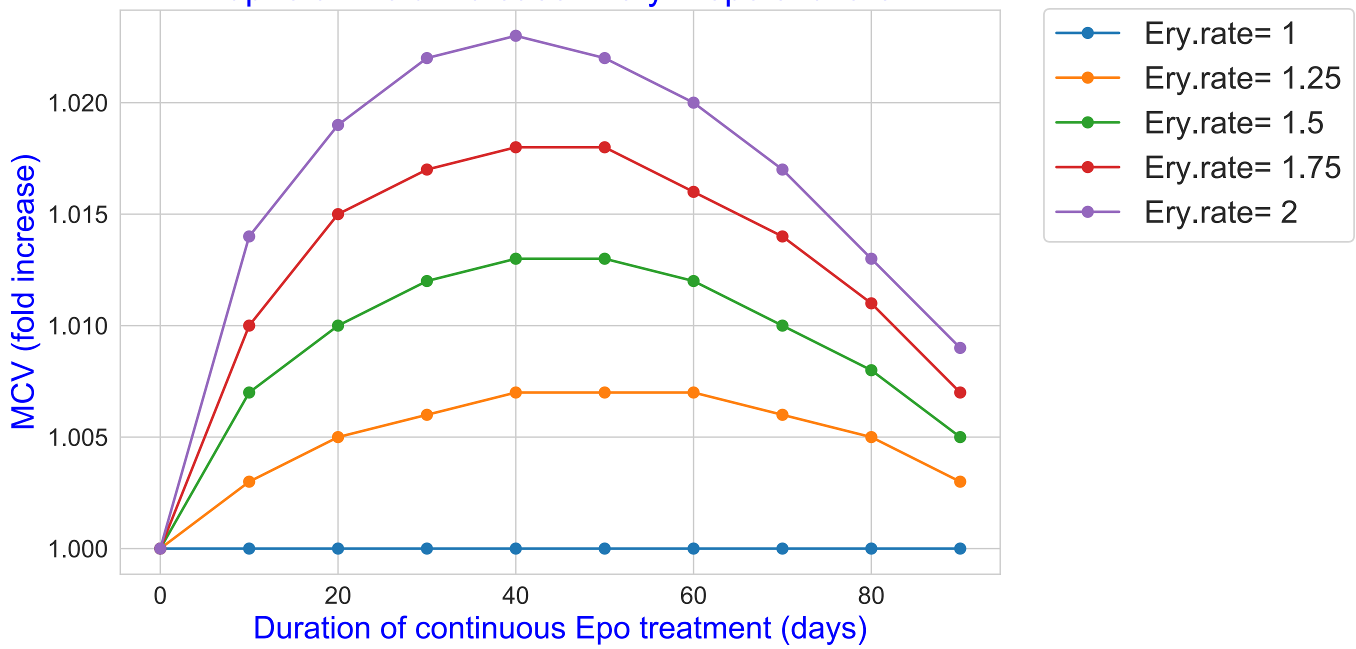


Fig 5B: MCV time course during continuous Epo treatment stimulating a 3 to 10-fold increase in erythropoietic rate

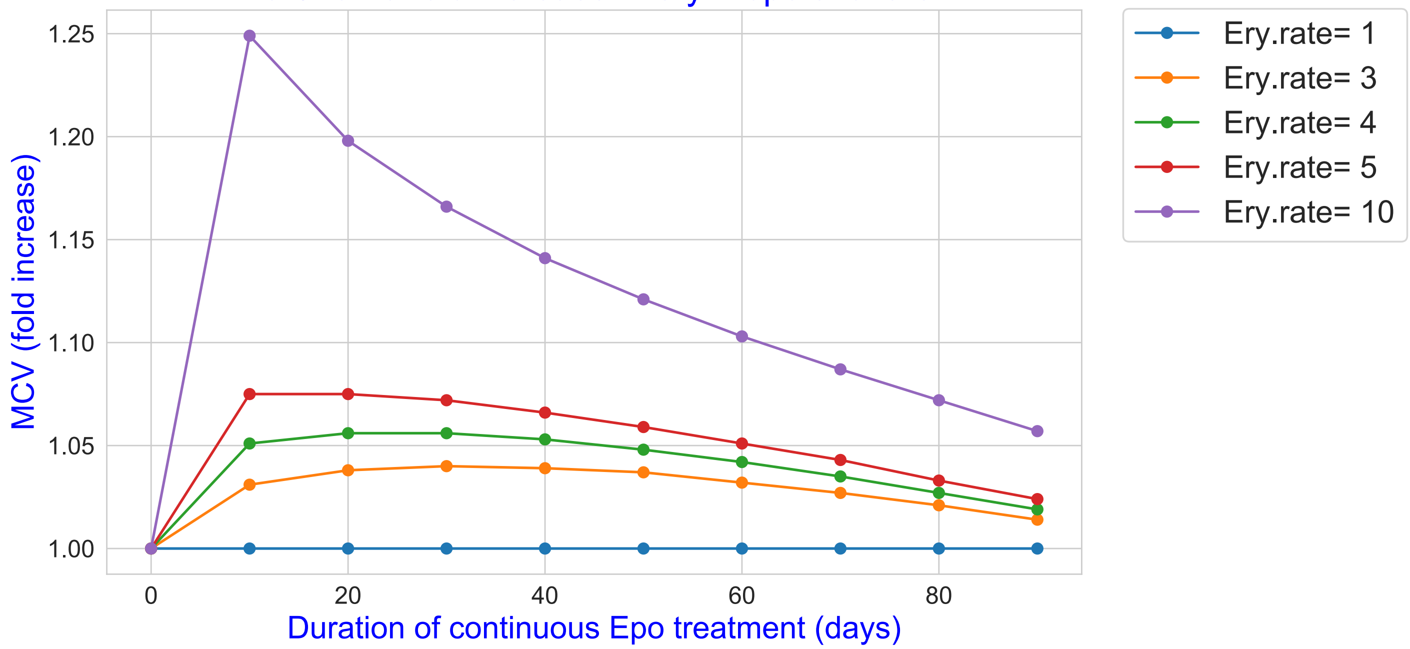

#### VI Predicted RC volume distributions: incorporating the post-treatment period

Figure 6A,B, show examples of RC volume distributions for specific Epo treatment durations. We overlay the volume distributions during and following the treatment at 10 day intervals. These show a transient increase in young (larger) RC number during the treatment; this cohort becomes gradually older and smaller in volume over time. The MCV is the arithmetic mean of RC volumes for each time point.

Fig. 6A: RC volume distributions during and following Epo treatment.
Treatment durations: 10 to 40 days; ery rate increase: 1.5 fold

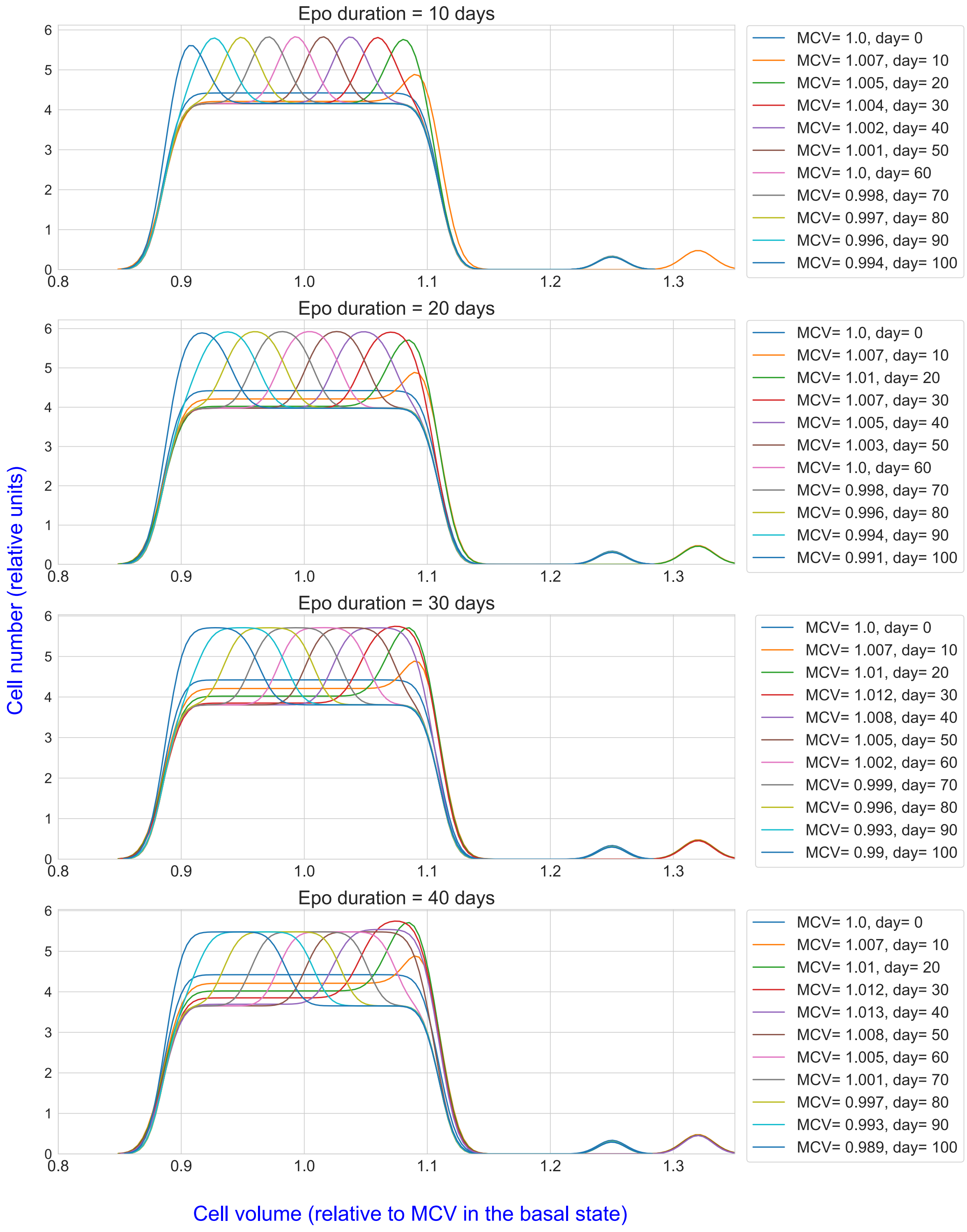

Fig. 6B: RC volume distributions during and following Epo treatment.
Treatment durations: 30 to 60 days; ery rate increase: 1.25 fold

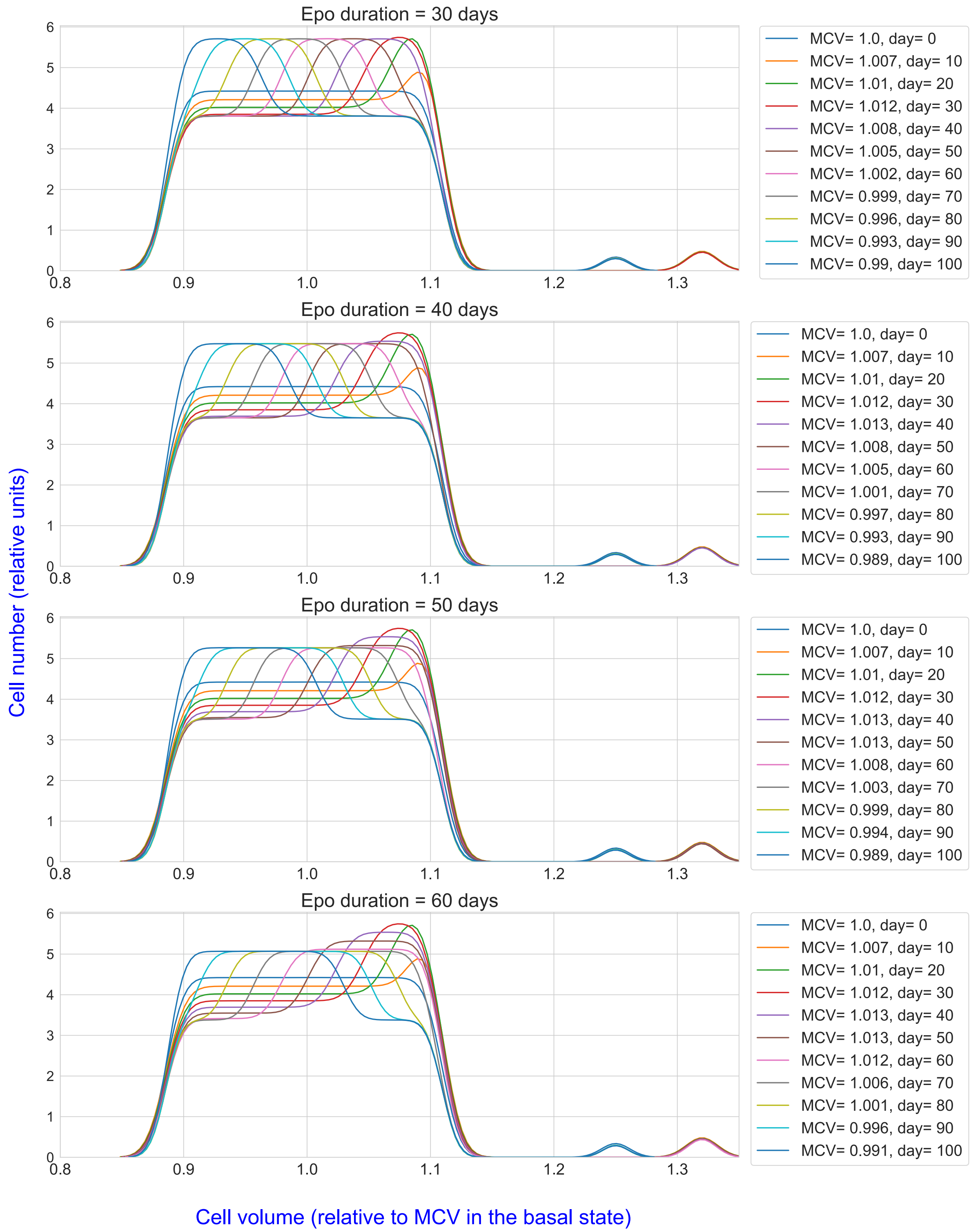

#### VII Predicted MCV time course during and following Epo treatment

In Figure 7A, B we simulated an increase in erythropoietic rate for varying durations, and plot the resulting time course of the MCV and reticulocyte counts d. The MCV at each time point is calculated from the distributions as shown in Figure 6 above. We overlay the time course plots for all treatment durations for a particular increase in ery. rate.

Fig 7A: Overlays of MCV and reticulocyte time course plots, for a 1.25 to 2 fold increase in ery. rate of varying durations

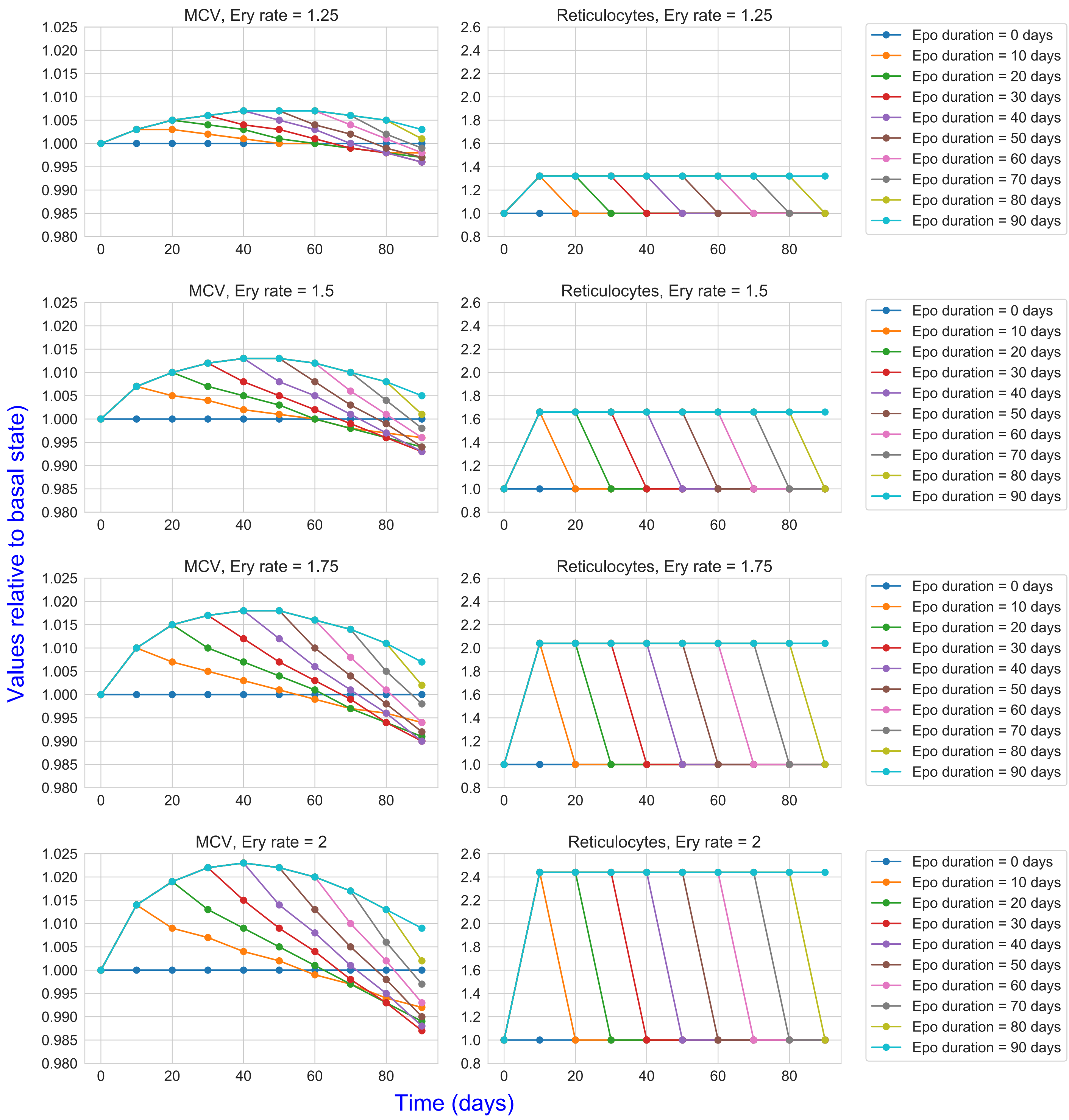

#### Fig 7B: Overlays of MCV and reticulocyte time course plots, for a 3 to 10 fold increase in ery. rate of varying durations

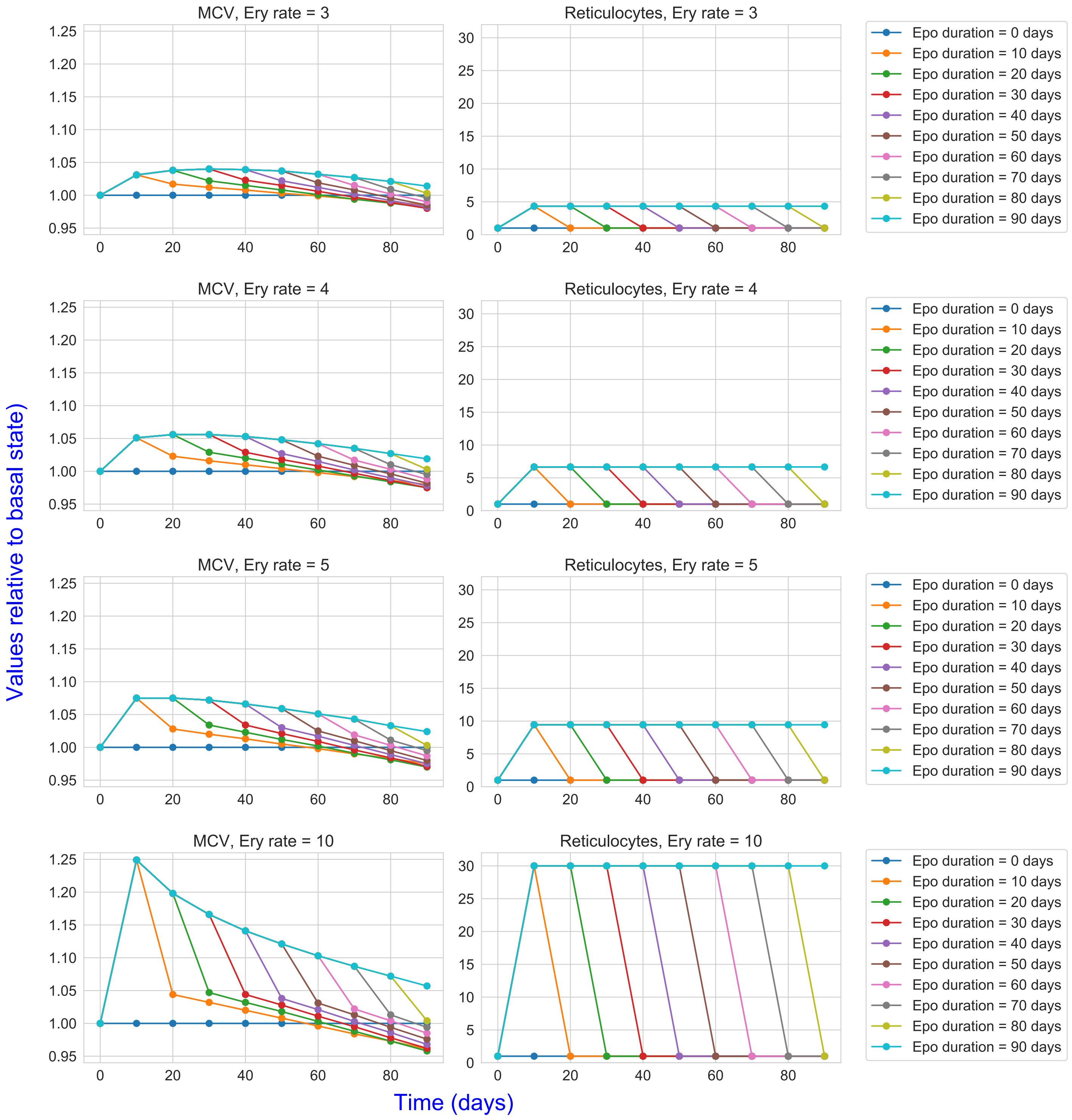

#### VIII Effect of reticulocyte count following Epo treatment on MCV time course

Following a period of Epo administration to healthy volunteers, erythropoietic rate (as reflected by the reticulocyte count) often declines below baseline. This decline is expected to lower the MCV. We had no clear data for the relationship between the drop in reticulocyte count, and the preceding increase in erythropoietic rate. For that reason, the simulations in sections VI and VII err on the side of caution and assume a return of reticulocyte count to baseline (which would tend to result in a higher MCV).

In the following section we use simulations that mimic the time course of the human interventions studies. We matched the post-treatment reticulocyte counts to the study data, where, in both studies, reticulocyte count dropped to 80% of the baseline value following treatment.

Here we examined the extent to which a drop below baseline affects the simulated MCV time course. In the example in Figure 8, we simulate Epo treatment for 50 days, which increased erythropoietic rate 1.5 fold. Following treatment, reticulocyte count dropped either to baseline, or to 50% the baseline value. A drop of reticulocyte count below baseline had a very small effect on the MCV. The same was found for other durations of Epo treatment and for higher or lower erythropoietic rates.

Fig. 8: Effect of reticulocyte count following Epo treatment on MCV
Epo duration = 50 day, ery. rate during Epo = 1.5

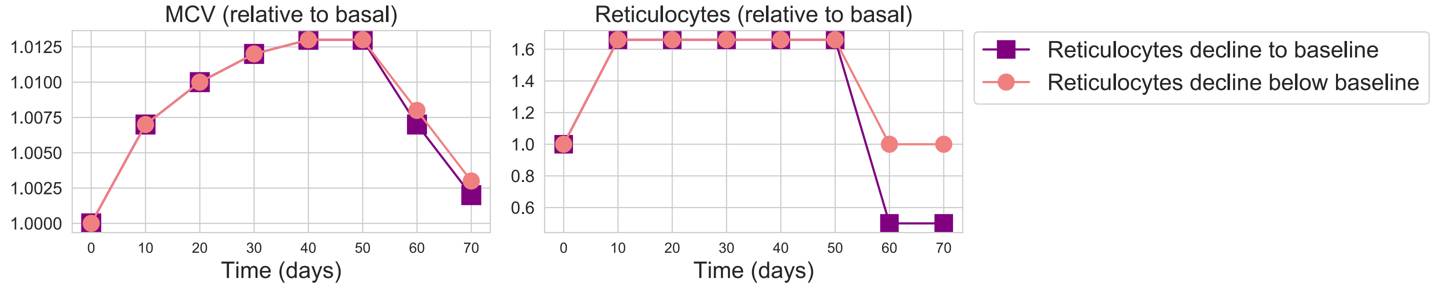

#### IX Comparison of human study data with predicted values:

##### A Human intervention study #1 (Extended Data Fig 9):

Participants were administered with 20 IU/kg Epoetin alpha every 48 hours for 3 weeks (~70 IU/kg Epoetin alpha / week). Figure 9A compares the observed MCV and reticulocyte time courses, with the simulated time courses based on the null hypothesis. We include simulations that assume an increase in erythropoietic rate of 1.25, 1.375 and 1.5 fold, which best match the observed increase in reticulocytes (lower panel). The numerical values for time points marked with a star are shown in the table below. The simulations suggest that the null hypothesis can account for up to 50% of the increase in MCV seen during and following Epo treatment in this study.

Fig 9A: Observation versus simulation, assuming the null hypothesis for human intervetion study #1

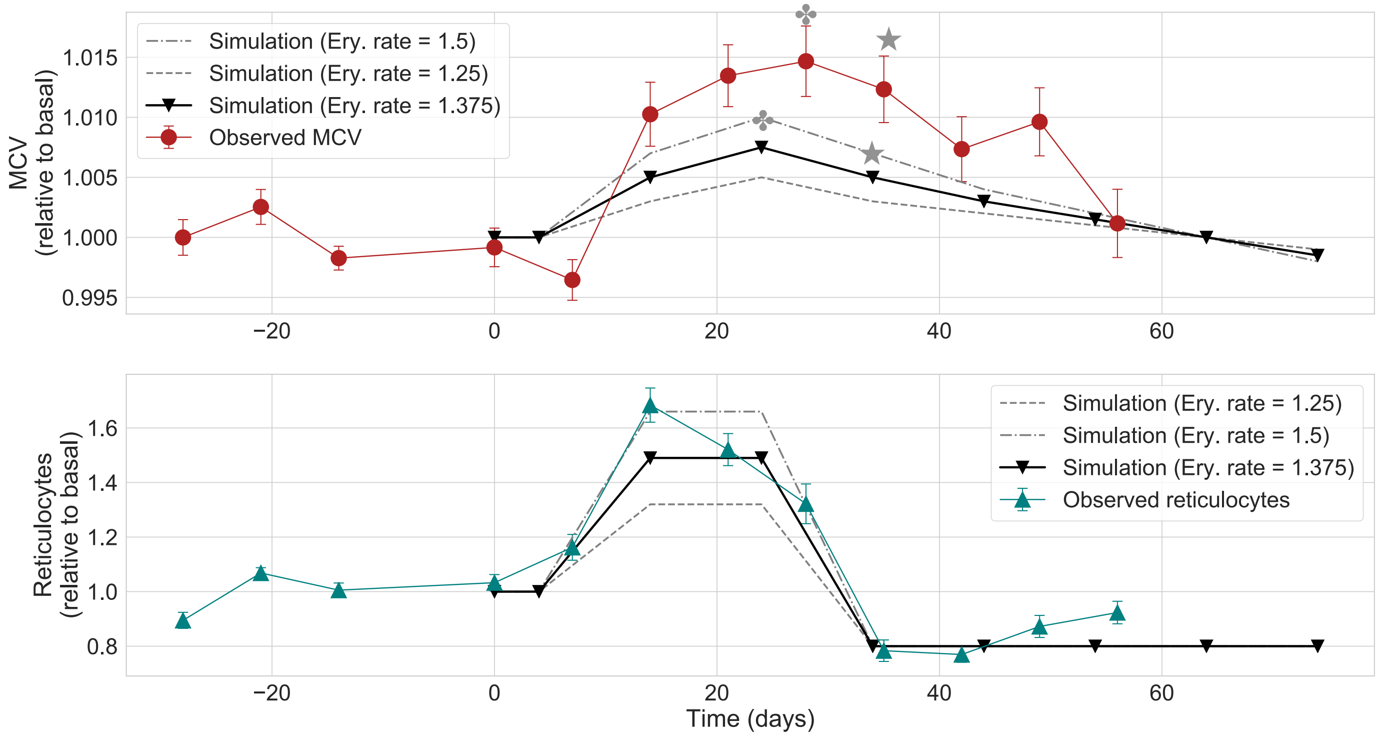


| | Increase in observed MCV | Simulated increase in MCV Ery rate = 1.375 (range: 1.25 -1.5) |
|---|---|---|
| **peak** ✣ | 1.47% | 0.75% (0.5% - 1.0%) |
| **post-treatment** ★ | 1.23% | 0.5% (0.3%-0.7%) |

**B Human intervention study #2 ( Extended Data Fig 10):**

Participants were administered weekly for 7 weeks with 5000 to 10000 IU/ 77kg (average participant weight), ~100 IU/kg Epoetin β/week. Figure 9B compares the observed MCV and reticulocyte time courses, with simulations based on the null hypothesis. We include simulations that assume an increase in erythropoietic rate of 1.25, 1.375 and 1.5 fold, which best match the increase in reticulocytes. The numerical values for time points marked with stars are shown in the table below. These simulations suggest that the null hypothesis can account for up to 30% the observed increase in MCV, either during or following Epo treatment.

Figure 9B: Observation versus simulation, assuming the null hypothesis for human intervetion study #2

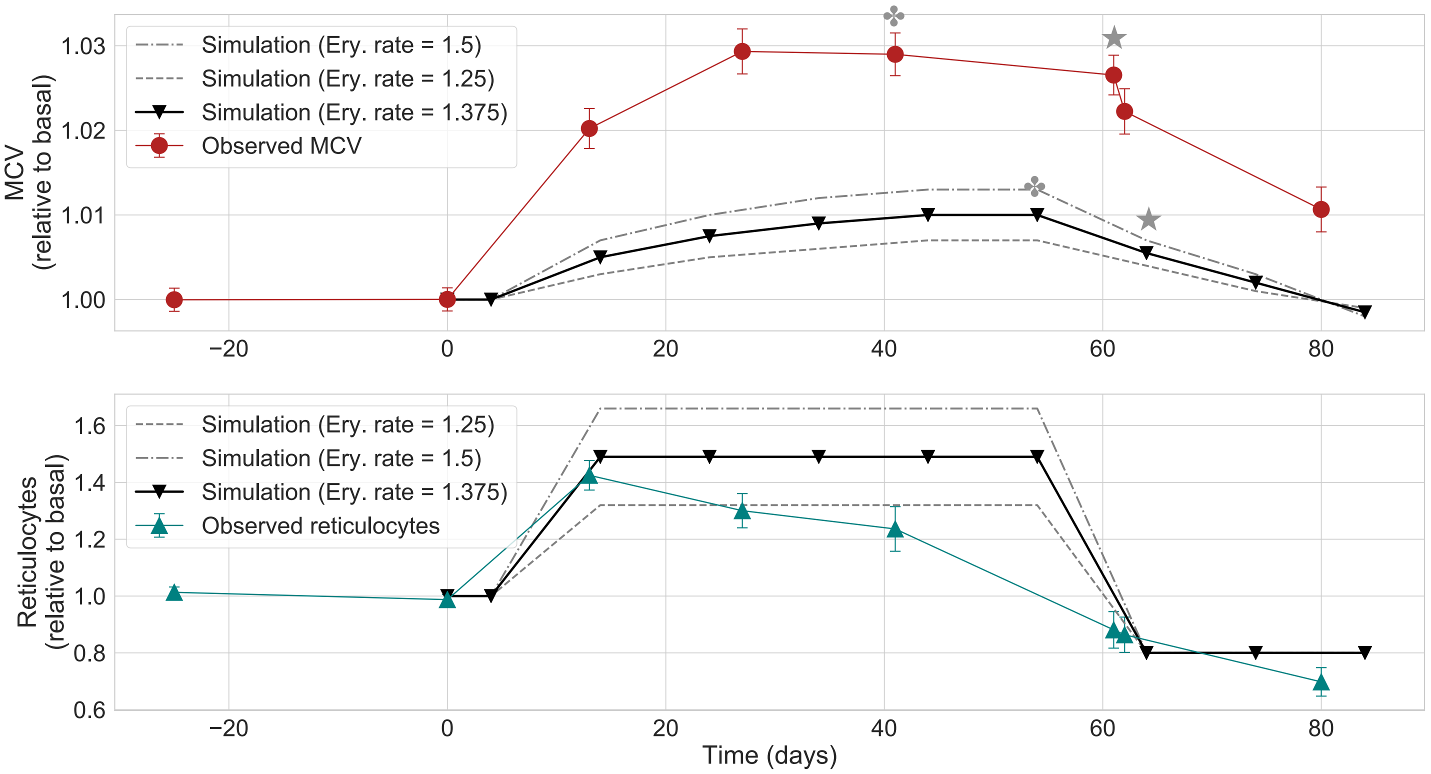


| | Increase in observed MCV | Simulated increase in MCV Ery rate = 1.375 (range: 1.25 -1.5 ) |
|---|---|---|
| **peak** ✤ | 2.93% | 1.0% (0.7% - 1.3%) |
| **post-treatment** ★ | 2.65% | 0.55% (0.4%-0.7%) |

**C Effect of decreasing RC life span from 100 days to 95 days**

We repeated the entire simulation, setting RC life to 95 days, while maintaining the same rate of cell size loss with age. As indicated by the graphs below, the results are closely similar to those obtained with RC life of 100 days.

Fig 10A: Observation versus simulation, assuming the null hypothesis for human intervention study #1 (RC life = 95 days)

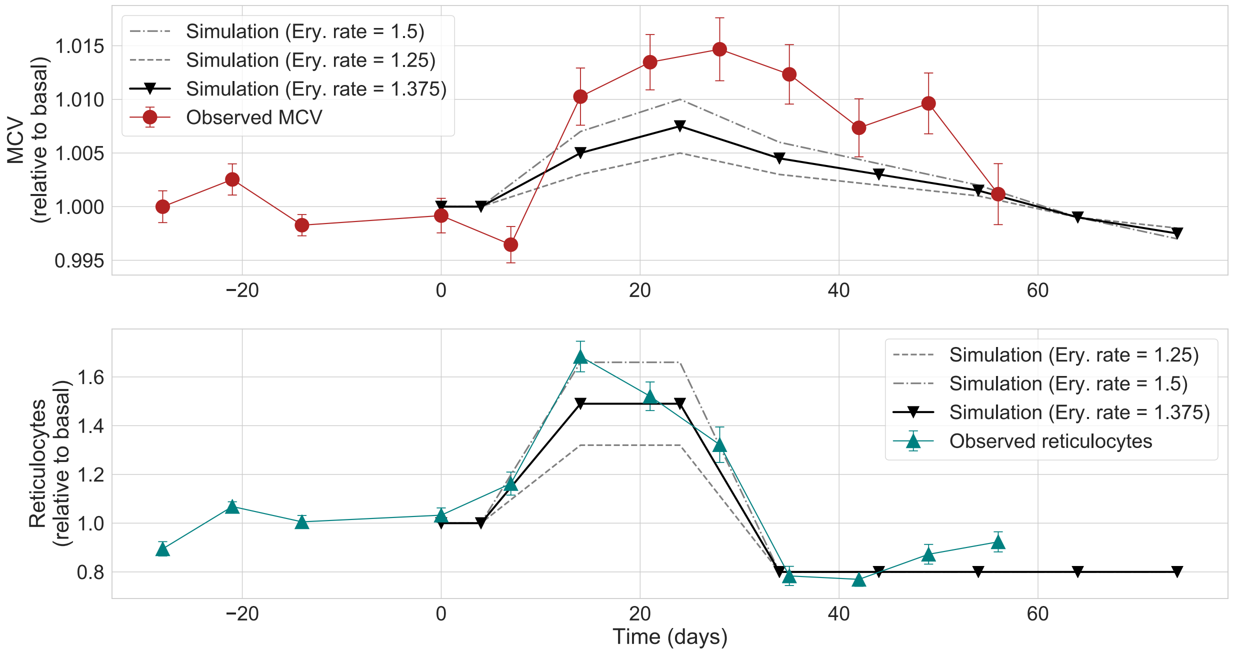


Figure 10B: Observation versus simulation, assuming the null hypothesis for human intervention study #2 (RC life = 95 days)

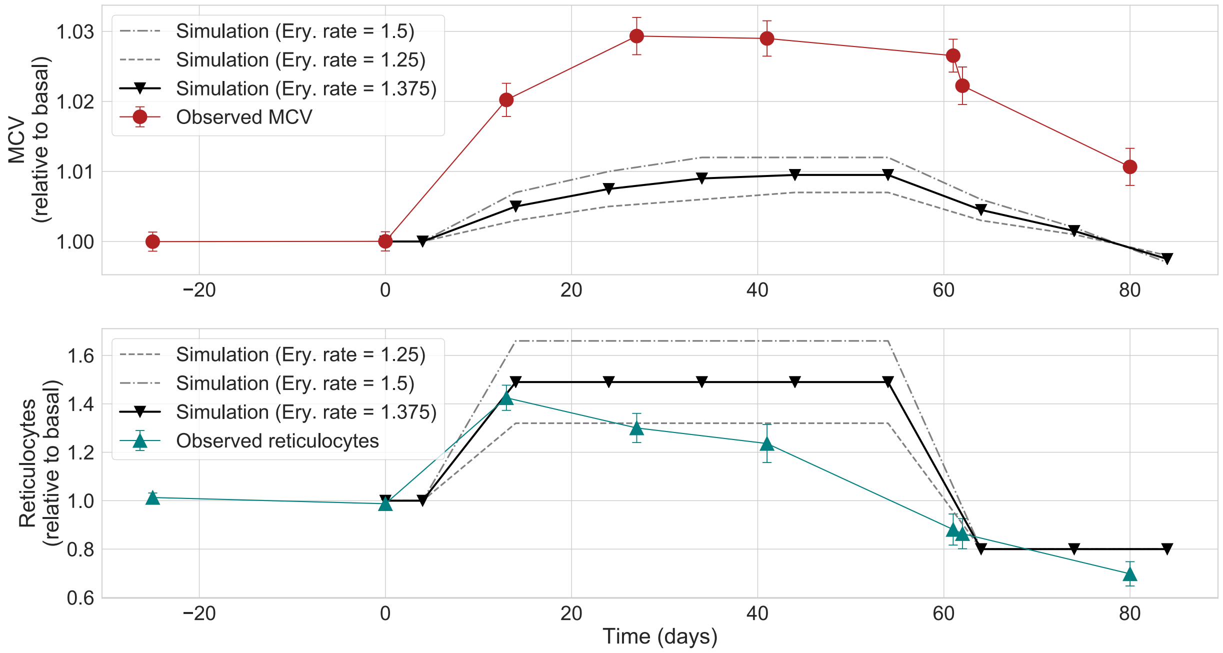

#### X Conclusions

We simulated the change in in MCV over time during and after Epo treatment. The simulated change in MCV results from increased reticulocyte count, the early release of larger reticulocyte with extended circulation time, and from a skewed age distribution in the population of mature RCs.

We found that the simulated MCV accounted for 30 to 50% of the increase in the observed MCV in the 2 human intervention studies that we report.

These findings suggest additional factors increase the MCV during Epo administration. Based on our experimental findings in the mouse, these additional factors are likely to include the generation of larger red cells in response to EpoR signaling.
